# How Varenicline Works: Identifying Critical Receptor and Ligand-based Interactions

**DOI:** 10.1101/2025.06.14.659675

**Authors:** Sheenagh G. Aiken, Daniele Fiorito, Matthew Harper, Grzegorz Pikus, Juno Underhill, Jacob Murray, Joshua Rawlinson, AnnMarie C. O’Donoghue, Cecilia Gotti, Sarah C. R. Lummis, Teresa Minguez Viñas, Franco Viscarra, Isabel Bermudez, Timothy Gallagher, A. Sofia F. Oliveira

**Affiliations:** School of Chemistry, University of Bristol, Bristol BS8 1TS (United Kingdom); Department of Chemistry, Durham University, South Road, Durham DH1 3LE, (United Kingdom); CNR, Institute of Neuroscience, University of Milan, I-20129 Milan (Italy); Department of Biochemistry, University of Cambridge, Cambridge, CB2 1QW (United Kingdom); Department of Biological and Medical Sciences, Oxford Brookes University, Oxford OX3 0BP (United Kingdom); Computational Chemistry Centre, School of Chemistry, University of Bristol, Bristol BS8 1TS (United Kingdom)

## Abstract

Approved by the FDA in 2006, varenicline became the first nicotinic-based therapeutic for smoking cessation and has since been used by tens of millions of smokers worldwide. Varenicline works by targeting the α4β2 nicotinic acetylcholine receptor (nAChR), the primary focus for nicotine addiction, where ligand recognition by the receptor triggers ion channel opening. While widely recognized that varenicline’s development was rooted in the well-established pharmacology of cytisine, the two compounds display notably different profiles, not only at nAChRs, but also at key off-target sites such as the 5-HT_3_ serotonin receptor. Despite varenicline’s widespread use and proven efficacy as a smoking cessation aid, our knowledge of the precise molecular mechanism underlying its action, particularly the specific receptor-ligand interactions that underpin its functional specificity, remains incomplete.

Through a multidisciplinary approach that integrates complementary fields of research, this study reveals the critical receptor-ligand interactions that distinguish varenicline from related nAChR agonists, such as cytisine and nicotine. Our findings reveal previously unrecognized, critical hydrogen bonding interactions within the α4β2 binding sites, specifically involving α4T139, α4T183, and β2S133, that are uniquely and selectively engaged by varenicline. Of these, β2S133 emerged as the pivotal determinant of varenicline’s function, with substitution by valine significantly impairing the ligand efficacy. Furthermore, the design and synthesis of novel varenicline analogues shed new light into the functional importance of the ligand’s quinoxaline moiety, revealing that not just the presence but also the precise positioning of this hydrogen bond acceptor are critical for receptor activation by varenicline. Together, these findings uncover a previously uncharacterized interaction network essential for varenicline’s function at α4β2, offering a deeper and more comprehensive framework for understanding its distinct pharmacological profile while expanding our broader understanding of how ligand binding is translated into function in these receptors.

**GRAPHICAL ABSTRACT:** 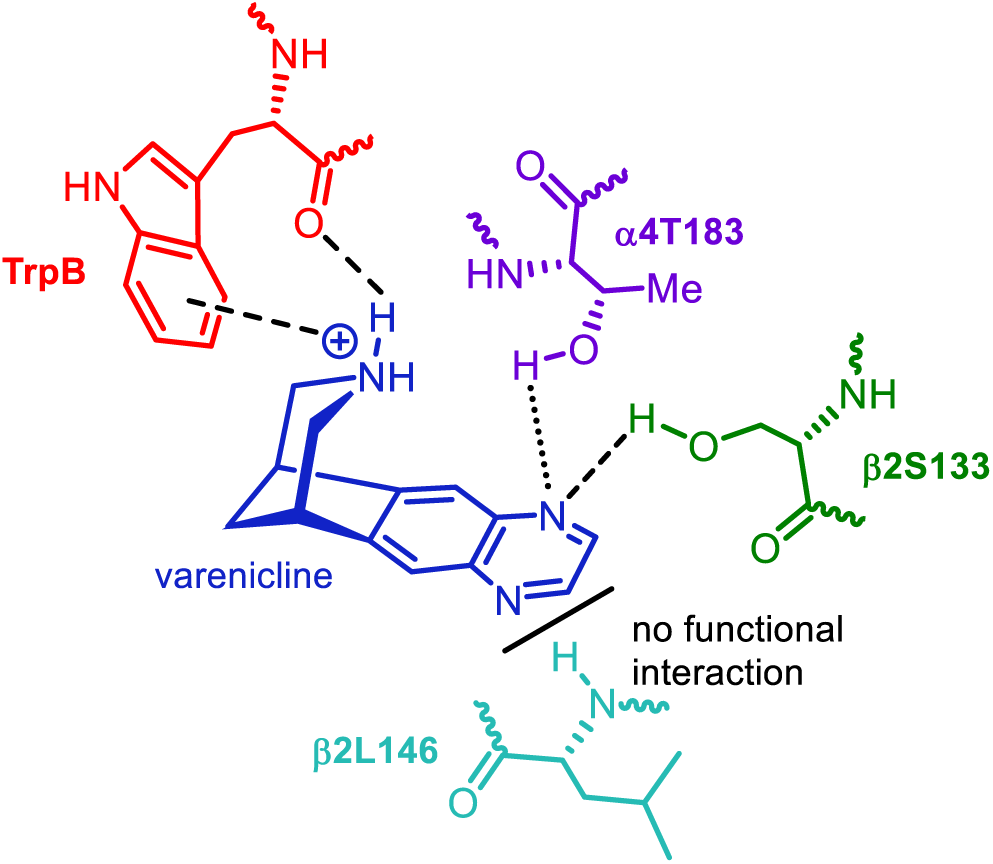

## INTRODUCTION

Tobacco consumption, with the World Health Organization (WHO) estimating >8 million deaths annually, is a leading cause of preventable disease and death worldwide. Of this, 7 million deaths are attributable to direct smoking and approximately 1.3 million additionally to second-hand smoke exposure.^1–3^ As a result, smoking cessation efforts have come to represent a major but frustratingly challenging and increasingly ephemeral global health objective.^4–8^ Further, recent declines in the prevalence of tobacco consumption have slowed, legislative activities to limit tobacco sales have failed to keep pace (or been reversed), and the underlying issue has been exacerbated by a sharp increase in nicotine consumption via electronic vapes^9–12^ and (more recently) nicotine pouches.^13, 14^ As a result, this rapidly increasing public health threat should be viewed as one of nicotine, as opposed to solely tobacco, addiction.^15–18^

A key part of the toolkit of smoking cessation agents is varenicline **1** (Figure 1A). Marketed as Chantix® (in the United States) and Champix® (in Europe),^19^ this drug was launched in 2006 to support smoking cessation.^18–21^ Now (since 2022) available in generic form, varenicline **1** has been used by >24 million smokers and represents the first nicotinic acetylcholine receptor (nAChR) partial agonist^22^ FDA-approved therapeutic.^23^

**Figure 1.**
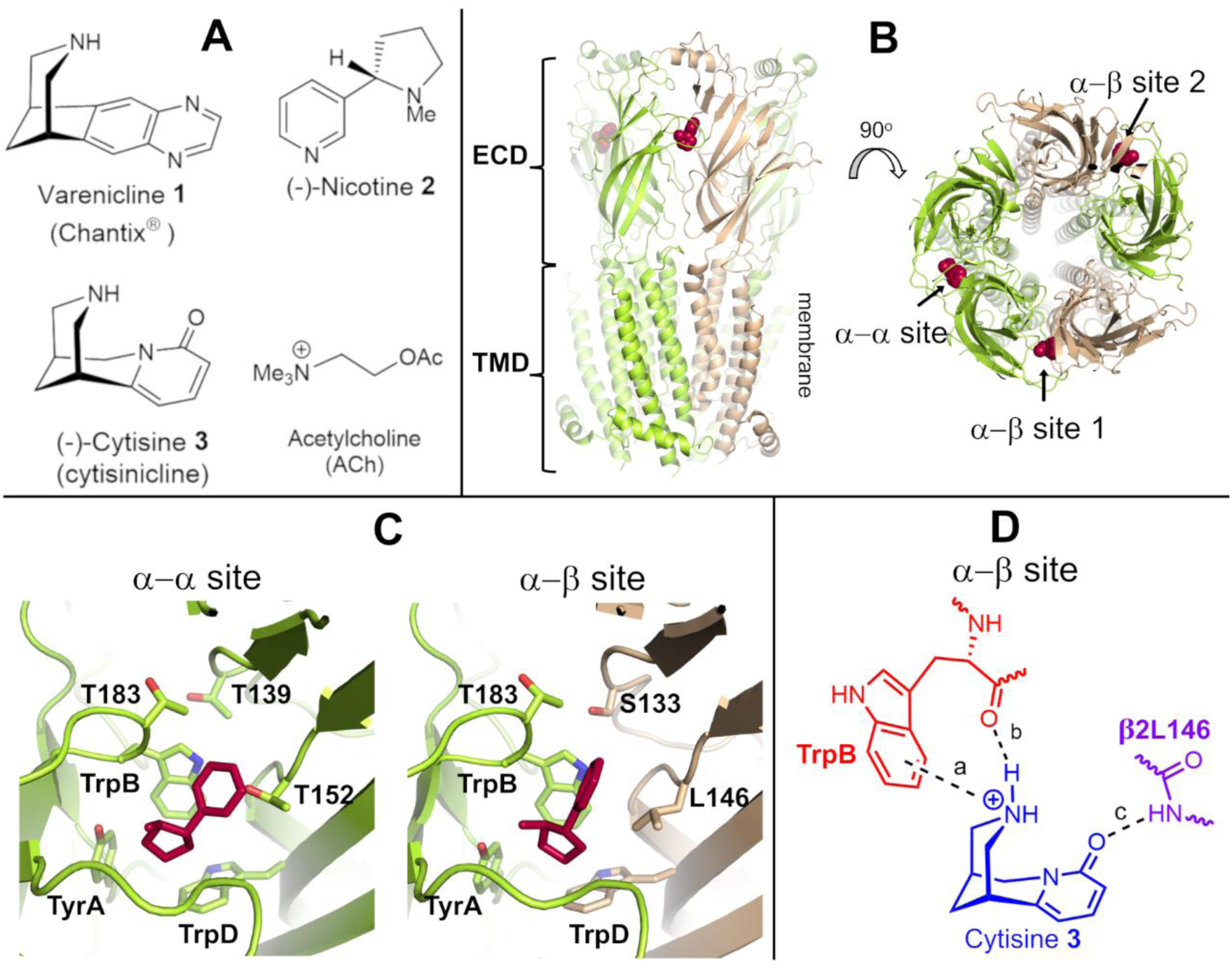
The α4β2 nAChR: structure, agonists and key agonist-receptor interactions. **A**) Chemical structure of the agonists varenicline **1**, nicotine **2**, cytisine **3**, and ACh. **B**) Cryo-EM structure of the human (α4)_3_(β2)_2_ nAChR with nicotine bound (PDB code: 6CNK^47^). NAChRs are composed of three domains: an extracellular domain (ECD), a transmembrane domain (TMD) and an intracellular domain (ICD). Please note that the ICD is absent in the cryo-EM 6CNK structure.^47^ The (α4)_3_(β2)_2_ nAChR contains one α−α pocket (formed by two α4 subunits) and two α-β sites (formed by a α4 and a β2 subunit). **C**) Close-up view of the α-α and α-β binding pockets in the cryo-EM structure of the human (α4)_3_(β2)_2_ nAChR with nicotine bound (PDB code: 6CNK^47^). The side-chains of the several conserved residues, namely TyrA (Y126 in the principal α4 side), TrpB (W182 in the principal α4 side) and TrpD (W88 in the complementary α4 side of the α-α pocket and W82 in the complementary β2 side of the α-β pocket) as well as the residues that are the focus of the current work, notably α4T183 (principal α4 side), α4T139 (complementary α4 side of the α-α pocket), β2S133 (complementary β2 side of the α-β pocket), α4T152 (complementary α4 side of the α-α pocket) and β2L146 (complementary β2 side of the α-β pocket) are shown with sticks. The residue numbers refer to Uniprot sequences P43681 and P17787 for the human α4 and β2 subunits, respectively. In panels A and C, the α4 and β2 subunits are colored in yellow and light brown, respectively. Nicotine **2** is highlighted in red. **D**) Dougherty-Lester nAChR functionally-important binding model for the α-β pocket illustrated for cytisine **3** showing the three key functional interactions identified: (*a*) cation-π, and (*b*) backbone C=O as H-bond acceptor associated with TrpB within the α4 subunit and (*c*) backbone NH as H-bond donor associated with b2L146 (in the complementary β2 subunit).^51–54^

As a partial agonist, varenicline **1** targets the α4β2 subtype of the wider set of nAChRs found in the nervous system.^20, 24–26^ This subtype, because it has a high affinity for nicotine **2**, emerged as the primary focus for nicotine addiction^15, 27, 28^ and, consequently, the target receptor for smoking cessation.^15, 27, 28^ There are, however, competing nAChR subtypes present in the nervous system, and varenicline **1** also binds to the α7 subtype where it acts as a full agonist,^26, 29, 30^ although the role of the α7 receptor in smoking cessation remains undefined. Significantly, varenicline **1** also activates other receptors besides nAChRs, namely the 5-HT_3_ serotonin receptor,^31^ which is a structurally related member (together with GAABA and glycine receptors) of the Cys loop superfamily.^16, 32^

The genesis of varenicline **1** was based on the known profile of cytisine (**3**; cytisinicline^33^) (Figure 1A).^34, 35^ Cytisine **3**, isolated from laburnum seed, has been used in Eastern Europe (under the brand name Tabex®) as a smoking cessation agent since the 1960s.^36–40^ Cytisine **3**, whose cholinergic properties were first studied by Dale *et al.*^41^ over 100 years ago, is both a high-affinity partial agonist for the α4β2 nAChR and a full agonist at α7 nAChR.^26, 29, 42, 43^ In this context, cytisine **3** served as an excellent structural and pharmacological lead for the discovery and development of a novel (and consequently patentable) smoking cessation agent, such as varenicline **1**. However, importantly, varenicline **1** and cytisine **3** diverge in terms of two facets of their pharmacology, namely their functional profile at the α4β2 and 5-HT_3_ receptors. Firstly, varenicline **1** and cytisine **3** have substantially different functional profiles in α4β2 nAChR,^26, 29^ which presents in two receptor stoichiometries: (α4)_2_(β2)_3_, the high sensitivity (HS) complex (Figure 1B), which has high-sensitivity to activation by ACh (EC_50_ 3 μM; Table 1) and (α4)_3_(β2)_2_, the low sensitivity (LS) complex, which has lower ACh sensitivity (EC_50_ 99 μM; Table 1).^44–46^ Varenicline **1** is efficacious in both α4β2 isoforms, while cytisine **3** only activates the LS (α4)_3_(β2)_2_ stoichiometry. Note that the two α4β2 stoichiometries have different numbers of agonist binding sites: the HS isoform contains two binding sites at the interface between the α4 and β2 subunits (hereafter labelled as α-β sites), while the LS isoform has these two α-β sites plus an additional α-α pocket at the interface between the two α4 subunits (Figures 1B and 1C).^47, 48^

**Table 1.** Potency and relative efficacy of nicotinic ligands at wild type (WT) and targeted mutants of both HS (α4)_2_(β2)_3_ and LS (α4)_3_(β2)_2_ of α4β2 nAChR. EC50 values were estimated as described in the Supporting Information. Relative efficacy (RE) was determined by normalizing the maximal current responses elicited by varenicline **1** to the maximal current response to ACh (I/I_max_ACh). Data shown represent the mean +/- SEM of n= 8-10 experiments in 6 to 8 different batches of *Xenopus* oocytes. Statistical differences between wild-type (WT) and mutant receptors were determined by one-way ANOVA followed by a post hoc Dunnett’s test and/or a post hoc Bonferroni multiple comparison test to determine the level of significance between wild type and mutants. * denotes a statistically significant difference (p < 0.05) between mutant and wild-type receptors. ND, not determined due to low levels of functional expression (less than 50 nA of ACh maximal currents).

| Mutation | ACh |  | Varenicline 1 |  | Nicotine 2 |  | Cytisine 3 |  |
| --- | --- | --- | --- | --- | --- | --- | --- | --- |
| | EC<sub>50</sub> μM | RE | EC<sub>50</sub> μM | RE | EC<sub>50</sub> μM | RE | EC<sub>50</sub> μM | RE |
| WT | 3.2 ±0.7 | 1 | 0.090 ±0.01 | 0.18 ±0.02 | 1.77 ±0.4 | 0.31 ±0.01 | ND | 0.025 ±0.004 |
| $\beta 2$S133V | 4 ±1.3 | 1 | 3.10 ±0.4* | 0.059 ±0.017* | 3.0 ±0.5 | 0.23 ±0.02* | ND | 0.016 ±0.002 |
| $\alpha 4$T183V | 3.4 ±0.4 | 1 | 0.57 ±0.01* | 0.099 ±0.004* | 2.9 ±0.5 | 0.29 ±0.01 | ND | 0.021±0.003 |
| $\alpha 4$T139V | 2.8 ±0.4 | 1 | 0.051 ±0.001 | 0.18 ±0.02 | 1.5 ±0.4 | 0.29 ±0.02 | ND | 0.023 ±0.001 |
| $\beta 2$S133V$\alpha 4$T183V | 3.3 ±1.0 | 1 | 2.47 ±0.6* | 0.07 5±0.0031* | 3.2 ±0.5 | 0.20 ±0.01* | ND | 0.018 ±0.003 |
| $\beta 2$S133V$\alpha 4$T139V | 3.5 ±0.9 | 1 | 3.58 ±0.4* | 0.067 ±0.018* | 2.8 ±0.9 | 0.22 ±0.02* | ND | 0.022 ±0.009 |

( $\alpha 4$ )<sub>3</sub>( $\beta 2$ )<sub>2</sub> (LS isoform)
| Mutation | ACh |  | Varenicline 1 |  | Nicotine 2 |  | Cytisine 3 |  |
| --- | --- | --- | --- | --- | --- | --- | --- | --- |
| | EC<sub>50</sub> μM | RE | EC<sub>50</sub> μM | RE | EC<sub>50</sub> μM | RE | EC<sub>50</sub> μM | RE |
| WT | 99± 10 | 1 | 1.15 ±6 | 0.41 ±0.02 | 7.5±1.6 | 0.53 ±0.05 | 2.9 ±0.8 | 0.23 ±0.08 |
| $\beta 2$S133V | 105 ±15 | 1 | 33 ±1.4* | 0.02 ±0.003* | 11.2±2 | 0.48±0.06* | 5.1 ±1 | 0.21 ±0.01 |
| $\alpha 4$T183V | 95 ±10 | 1 | 8.0 ±0.3* | 0.20 ±0.02* | 6.9±2 | 0.52 ±0.03 | 4.6 ±1 | 0.22 ±0.02 |
| $\alpha 4$T139V | 88 ±9 | 1 | 8.4 ±5.4* | 0.20 ±0.02 | 7.4±1 | 0.54 ±0.05 | 2.5 ±0.7 | 0.19 ±0.02 |
| $\beta 2$S133V$\alpha 4$T183V | 101 ±4 | 1 | 33.0 ±5* | 0.033 ±0.005* | 10.2±2 | 0.47 ±0.01* | 6.9 ±1.5 | 0.19 ±0.01 |
| $\beta 2$S133V$\alpha 4$T139V | 98 ±10 | 1 | 41.34 ±2* | 0.031 ±0.005* | 11.4±2 | 0.46 ±0.02* | 5.8 ±1.4 | 0.18 ±0.01 |

*(c) Exploring the structural features of varenicline **1** that contribute to binding and function: a focus on the role of the heteroaryl moiety.*

Secondly, cytisine **3** is a weak antagonist at the 5-HT_3_ receptor, unlike varenicline **1**, which is an agonist at this receptor.^31, 49^ This pharmacological distinction has been linked to reduced side-effects experienced with cytisine **3** compared to those associated with varenicline **1** therapy.^50^ The differing profiles of these two closely related compounds, particularly given that cytisine **3** is often considered the progenitor of varenicline **1**, raise several fundamental questions about their underlying mechanisms of action at the α4β2 nAChR. Firstly, what are the molecular details of varenicline **1** binding to the α4β2 nAChR in relation to the key protein-ligand interactions that both mediate and differentiate varenicline **1** and cytisine **3** functional effects? Secondly, how does varenicline **1** binding translate into efficacy, i.e. receptor activation? Clearly, this then begs the question, that if varenicline **1** differs from cytisine **3** in terms of *how* it binds to the receptor, does that difference translate to function? Further, if the specifics of the α4β2 receptor-ligand interactions are different for varenicline **1** and cytisine **3**, does this then help to shed light on why varenicline **1** is an agonist while cytisine is an antagonist at the human 5-HT_3_ receptor. This directs towards the need to enhance our understanding of the network of receptor-ligand interactions that govern function, with the ultimate goal of achieving precise control over its activity.

The seminal work of Dougherty and Lester,^51–56^ together with structural studies of ligands bound to different soluble acetylcholine binding proteins (AChBP),^57–60^ and recent high-resolution structural data for complete nAChRs,^61–65^ including the α4β2-nicotine, acetylcholine and varenicline complexes (PDB codes: 5KXI, 6UR8, 6USF 6CNJ, 6CNK, 8SSZ, 8ST0, 8ST1 and 8ST2),^47, 66–68^ has defined a binding model for important nicotinic agonists (Figure 1D).

The salient features of the Dougherty-Lester model consist of a cation-π and H-bond donor (except for acetylcholine) associated with the piperidine ammonium center and a highly conserved tryptophan residue in loop B (TrpB) in the principal face of the agonist binding site,^51, 52^ and an H-bond acceptor component within the ligand (C=O, in the case of cytisine **3**) that interacts with a donor group within the complementary face (Figure 1D).^53^ For most agonists bound to the α4β2 nAChR, the H-bond with the complementary face involves an interaction with the backbone NH of β2L146 (which corresponds to β2L119 in the work of Blum *et al.*^53^) in the complementary face.^53, 54, 56^ However, while this interaction mediates ACh, nicotine **2**, cytisine **3,** sazetidine-A, carbamylcholine, and epibatidine function,^53, 54, 56^ varenicline **1** was shown not to engage with β2L146 at least in terms of linking to receptor *function* in the α4β2 subtype.^56^ Dougherty and co-workers used changes in EC_50_ as the primary metric to determine the relevance of a ligand-receptor interaction,^55^ and mutation of β2L146 (to its α-hydroxy analogue) did not affect varenicline **1** *function* in either of the LS and HS α4β2 isoforms.^56^ Note, however, that their findings do not exclude the involvement of β2L146 in varenicline **1** *recognition*/*binding*, as agonism relates to function. Moreover, it is worth pointing out that other additional (ligand-receptor or/and receptor-receptor) interactions within the agonist binding sites can contribute to subtype differentiation. For instance, we have previously uncovered how specific interactions (such as those involving α7R101 and β2R106 within β3 strand) modulate functional outcomes across various human nAChR subtypes.^29, 69^ We have also explored how agonist-induced structural and dynamics changes are transmitted from the binding sites in the α4β2 and α7 nAChRs to the ion channel,^70, 71^ ultimately allowing gating to occur.

Given that varenicline **1** retains the features associated with cation-π and H-bond donor components (shown in Figure 1D) in the α4β2 nAChR,^54^ but lacks interaction with β2L146, we have focused here on clarifying the role of the ligand-based H-bond acceptor interaction in shaping the functional profile of this ligand and uncovering the specific residue(s) involved in this interaction. To investigate the variation of receptor-ligand network that mediates *function* across different nicotinic agonists, including varenicline **1**, we employed a multidisciplinary approach, integrating computational and experimental methods, to identify and explore the role of alternative *receptor* networks of H-bonding interactions within both the primary and complementary faces (of both α-β and the α-α binding sites) of α4β2 nAChR. In addition, we have also designed a unique set of varenicline **1** variants that allow exploration of key *ligand* features that mediate binding to and function of the human α4β2 nAChR. Two considerations guided this latter set of ligand designs: *(i)* we sought to maintain both ligand size and molecular volume; and, *(ii)* we required an ability to vary both the presence/absence and location of the H-bond *acceptor* moiety within the ligand scaffold. These two complementary lines of investigation converged to offer new insights and a nuanced explanation of how and why the profile of varenicline **1** deviates from that of other nicotinic ligands, including from its closely related compound cytisine **3,** in the α4β2 nAChR. Our results demonstrate that varenicline **1** engages in functionally relevant interactions with β2S133 on the complementary β2 side and (to a lesser extent) with α4T183 on the principal α4 side of the agonist pocket. Notably, the interaction with β2S133 plays a substantial role in shaping the functional profile of varenicline **1**, emerging as a unique and (so far) distinguishing feature of its association with the receptor.

## RESULTS AND DISCUSSION

### (a) Uncovering alternative H-bond networks within the agonist binding sites of α4β2 nAChR

Molecular dynamics (MD) simulations were performed for several α4β2 nAChR-agonist complexes to identify potential alternative H-bonding donor networks within the agonist binding sites of this receptor (Table S1). The complexes between the extracellular domain (ECD) of the two stoichiometries of the human α4β2 nAChR (i.e. the LS (α4)_3_(β2)_2_ and HS (α4)_2_(β 2)_3_ isoforms) and varenicline **1**, nicotine **2**, cytisine **3**, and ACh were constructed (hereafter named as wild-type complexes, WT). The optimized binding mode of all four ligands within the α-α and the two α-β binding pockets for the two α4β2 stoichiometries is shown in Figures S2-S3 (please refer to the Supporting Information for a detailed description of the construction and optimization of the complexes). While ACh is a full agonist of the α4β2 nAChR, varenicline **1**, nicotine **2,** and cytisine **3** are all partial agonists of this receptor, with an efficacy relative to ACh of 0.18, 0.31 and 0.025 for the HS isoform and 0.41, 0.53 and 0.23 for the LS one, respectively (Table 1). Note that efficacy is not an inherent ligand property, with some molecules acting as full agonists for one nAChR subtype and partial agonists for another.^22^

In the WT-varenicline **1** models, the nitrogen atoms of the quinoxaline moiety within the ligand, which is the putative H-bond acceptor, do not directly engage with the backbone amide nitrogen of either α4T152 (in the α-α pocket) or β2L146 (in the α-β pockets) in any of the three agonist binding pockets (Figure 2A). This is consistent with Dougherty and co-workers’ observation that β2L146 is not involved in modulating the functional profile of varenicline **1**.^56^

**Figure 2.**
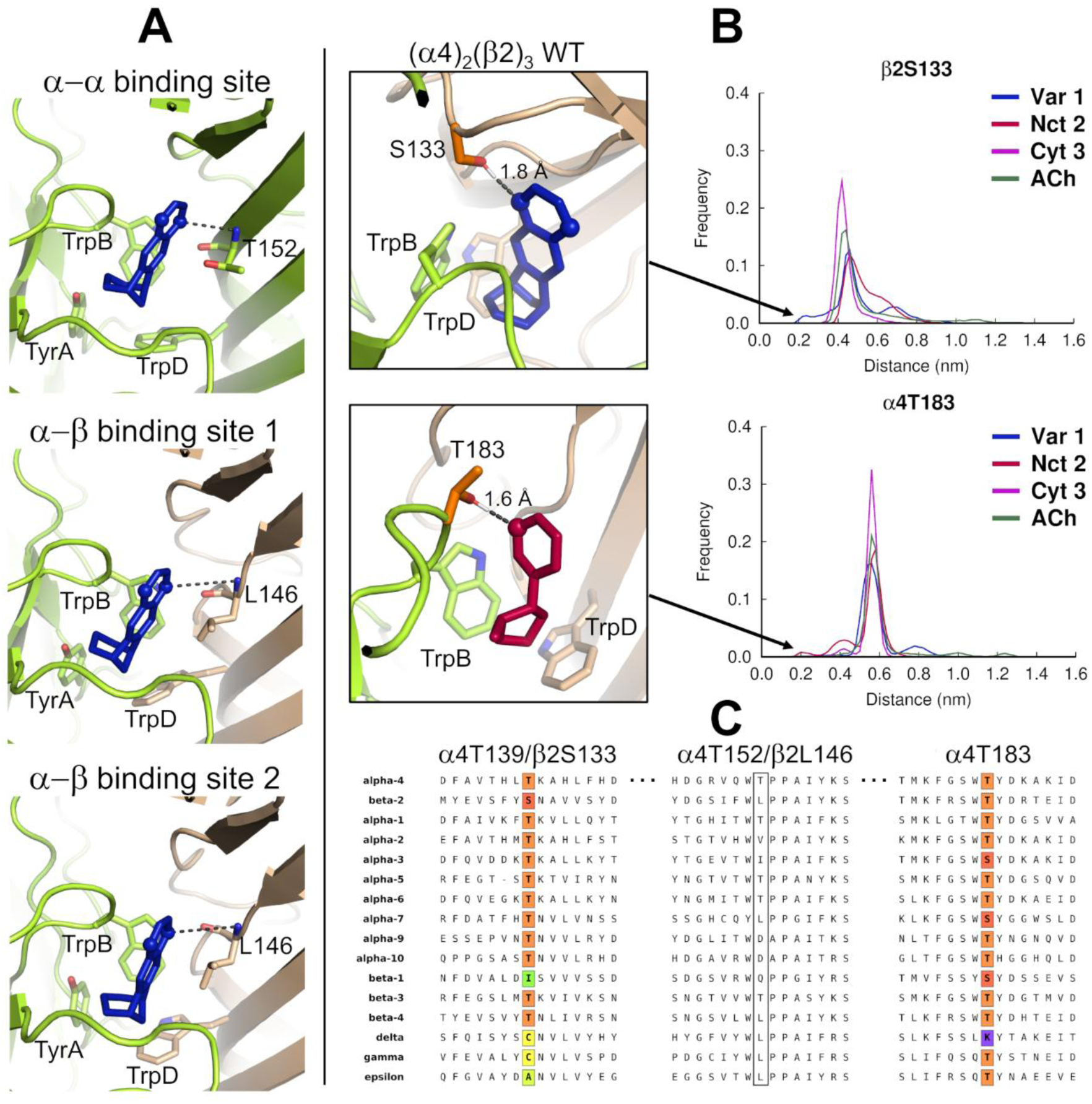
Varenicline interactions in the α-α and α-β binding sites in the wild-type complexes. **A)** Optimized binding mode for varenicline **1** in the α-α and α-β binding sites in the LS wild-type complex. The α4 and β2 subunits are colored in yellow and brown, respectively. TyrA, TrpB, TrpD, α4T152 and β2L146 are shown with sticks. Varenicline **1** is colored in blue, with the nitrogen atoms of the quinoxaline moiety highlighted with spheres. Note that the distance between the nearest H-bond acceptor in the quinoxaline ring and the backbone NH group of α4T152 and β2L146 (indicated by dashed lines) exceeds 3.8 Å (distances >3.8 Å are also observed for the α-β binding sites in the HS complex), thus suggesting that no H-bond is formed between these two groups. **B)** Distribution of the minimum distance between the agonist (specifically, the closer pyrazine nitrogen in the quinoxaline moiety of varenicline **1**, the pyridine nitrogen of nicotine **2**, the pyridone carbonyl oxygen of cytisine **3** and the closest oxygen in the ester group of ACh) and α4T183 and β2S133 in the α-β binding pockets of the HS wild-type complex (right panels). The left panels illustrate examples of conformations where a hydrogen bond between varenicline **1** and β2S133 and nicotine **2** and α4T183 are present (as indicated by the dashed lines). The α4 and β2 subunits are colored in yellow and brown, whereas varenicline **1** and nicotine **2** are highlighted in blue and red. The side-chain of TrpB, TrpD, α4T183 and β2S133 are represented with sticks. The nitrogen atoms of the quinoxaline ring of varenicline **1** and the pyridine nitrogen of nicotine **2** are highlighted with spheres. **C)** Sequence alignment for the α4T183, α4T152/β2L146 and α4T139/β2S133 regions of various human nAChR subunits. The colored boxes highlight the locations of α4T183, α4T139 and β2S133 (the residues mutated in this work), with threonine, serine, isoleucine, cysteine, alanine, and lysine residues represented by orange, red, green, yellow, light green, and purple, respectively. The white box marks the location of α4T152 and β2L146, which were not mutated here. The sequences shown correspond to the P43681 (human α4), P17787 (human β2), P02708 (human α1), Q15822 (human α2), P32297 (human α3), P30532 (human α5), Q15825 (human α6), P36544 (human α7), Q9U6MI (human α9), Q9GZZ6 (human α10), P11230 (human β1), Q05901 (human β3), Q07001 (human δ), P07510 (human γ), and Q04844 (human ε) UniProt codes. The sequence alignments were performed using the Muscle server.^72^

MD simulations were performed for the HS and LS α4β2 wild-type complexes to investigate the dynamics of the different ligands and reveal potentially relevant differences in their H-bond patterns with the receptor. All trajectories remained stable during the simulation time, showing structural convergence after ∼20 ns and exhibiting minimal secondary structure loss (Figures S6-S7). Principal component analysis was performed to assess the sampling of the individual replicates, showing that they generally explored different regions of the conformational space, thereby enhancing the overall sampling for each system (Figure S8). All agonists remained stably bound to the wild-type receptor (Figures S11-S13), with their protonated nitrogen (i.e. piperidine nitrogen in varenicline **1** and cytisine **3**, the pyrrolidine nitrogen in nicotine **2**, and quaternary ammonium nitrogen of ACh) making persistent cation-π interactions with TrpB (Trp182 located in loop B in the principal α4 face of the pockets) and occasionally with TyrA (Tyr126 in loop A in the principal α4 face of the pockets), and TrpD (Trp88 and Trp82 in loop D in the complementary α4 face of the α-α pocket and the complementary β2 face of the α-β pocket, respectively) (Figures S18-S20). These simulations also revealed distinct dynamic behaviors between agonists (Figures S14-S17): ACh exhibited high positional and conformational variability, adopting many binding modes within the pockets; in contrast, the bulkier ligands, probably due to their more rigid structures and, in some cases, additional interactions with the protein compared to ACh,^51–54, 56^ showed reduced mobility and generally maintained an orientation closer to the initial one throughout the simulation time (Figures S11-S13). Differences in agonist dynamics were also observed between the α-α and α-β pockets, mainly for nicotine **2** and cytisine **3**, with these ligands showing increased mobility when bound to the α-α pocket (Figures S11-S13).

Analysis of the distances between the H-bond acceptor in varenicline **1**, nicotine **2**, cytisine **3** and ACh, and H-bond donors within the binding pockets further identified three alternative residues that can transiently interact with some of the agonists (Figure S21). These alternative H-bonds involved T183 in the principal face of the pockets (hereafter identified as α4T183) and S133 and T139 in the complementary face (hereafter labelled as β2S133 and α4T139, respectively) of the α-β and α-α pockets. As can be seen in Figure S21, the H-bond acceptor groups of the agonists (i.e. the pyrazine nitrogen atoms in the quinoxaline moiety of varenicline **1**, the pyridyl nitrogen of nicotine **2**, the pyridone oxygen of cytisine **3** and the carbonyl oxygen of ACh) can, in some cases, closely approach the H-bond donors located in the side-chains of α4T183, β2S133 and α4T139, thus suggesting that transient (either direct or mediated by a close water molecule) interactions between the groups are feasible.

Within the α-β binding sites, the residues identified as potential H-bond partners for the agonists were α4T183, located in loop B on the principal face of the pocket, and β2S133, situated in the segment connecting β−sheets 4 and 5 on the complementary face. While α4T183 is positioned in the side of the pocket immediately after the key TrpB residue, β2S133 lines the back of the orthosteric site (Figure 1C). In our simulations, (and given the meso nature of varenicline **1**),^73^ only one of the quinoxaline nitrogen atoms of varenicline **1** was able to closely contact with the hydroxyl group of β2S133, while the pyridine nitrogen of nicotine **2** was able to interact with the hydroxyl group of α4T183 (Figures 2B and S21). In the LS (α4)_3_(β2)_2_ stoichiometry of the α4β2 nAChR, in addition to the α-β agonist pockets, there is an additional agonist site on the α-α subunit interface.^47, 48^ In the α-α site, the new interaction is associated with α4T139, which is located in the complementary face of the pocket in a position analogous to that of β2S133 on the α-β interface (Figure S21). Note that the interactions described above involving α4T139/β2S133 and α4T183 do not use backbone NH donors (as in the case of β2L146 in Figure 1D) but rather involve the side chain hydroxyl groups of the residues, which are potentially synergistic in terms of the network that they offer. Further, the side-chain of these residues are also conformationally more flexible and, as a result, more accommodating to ligand “demands”. Sequence alignment of various nAChR subunits indicates that the presence of an H-bond donor group at positions equivalent to α4T139/β2S133 and α4T183 is highly conserved across human receptors (Figures 2C and S22). All human neuronal subunits possess a residue with a hydroxyl-containing side-chain at the α4T139/β2S133 position, with the α2-α7, α9-α10 and β3-β4 subunits featuring a threonine and the β2 subunit a serine. In contrast to the human neuronal subunits, the muscle subunits (i.e. α1, β1, γ, δ, and ε) exhibit greater diversity in the residue at position α4T139/β2S133, with some having apolar residues that cannot participate in H-bonding, or a cysteine, which generally forms weaker hydrogen bonds compared to those involving hydroxyl (–OH) or amido/amino (–NH₂) groups (Figures 2C and S22).^74, 75^

At the α4T183 position, all human neuronal and muscle subunits have either a threonine or a serine, except for the δ subunit, which contains a lysine, which has a positively charged side-chain at physiological pH (Figures 2C and S22). The strongly conserved nature of residues at the α4T139/β2S133 and α4T183 positions in neuronal subunits suggests that the side chain hydroxyl group, acting as an H-bond donor, may play a key role in defining the action of certain ligands.

In the wild-type simulations, varenicline **1** exhibits strong, positively concerted motions with α4T139, α4T183 and β2S133 in both the α-α and α-β sites, whereas the correlation profiles for nicotine **2**, cytisine **3** and ACh vary between pockets, generally showing weaker correlations with the α-α pocket (Figure S23). The dynamics and preferred positions of α4T183 and β2S133 in the α-β binding pockets are generally similar across all wild-type complexes, with the side-chain of the two residues showing reduced flexibility (Figures S24-S25). In contrast, within the α-α binding pocket, the side-chain of α4T183 and α4T139 exhibit greater mobility, allowing them to adopt different conformations over the course of the simulations (Figure S24). Previous work, including by Dougherty and co-workers, comparing the α-α and α-β sites has shown that their differences stem primarily from the chemical characteristics of three residues located on the complementary face of the sites.^48, 56, 76, 77^ In the α-β site, hydrophobic residues β2V136, β2F144 and β2L146 are present on the complementary side, whereas, in the α-α site, these are replaced by the polar α4H142, α4Q150 and α4T152.^48, 56, 76, 77^ These substitutions alter the pocket properties, increasing its hydrophilicity and influencing agonist affinity and functional profile.

The distance between the H-bond acceptor in varenicline **1**, nicotine **2**, cytisine **3** and ACh, and the backbone NH donor of β2L146 (located in the complementary β2 face of the α-β pockets) and α4T152 (found in the complementary α4 face of the α-α pocket) was also determined (Figure S26). Note that α4T152 in the α-α site occupies an equivalent position to β2L146 in the α-β pocket as shown in Figure S26A. Generally, during the simulations in both the α-β and α-α pockets, only two of the agonists (notably varenicline **1** and ACh) were able to closely approach the backbone NH donor associated with these residues. Among them, only ACh showed distance values indicative of direct hydrogen bond formation (Figure S26).

Additionally, in the α-α pocket, α4T152, besides having a backbone amide NH donor, also possesses a hydroxyl group that can participate in H-bonding. Given this, the distance profile between the agonist H-bond acceptor(s) group (see above) and the α4T152 hydroxyl donor was also determined (Figure S26B). These profiles clearly demonstrate that cytisine **3** (and, to a lesser extent nicotine **2** and ACh) can directly interact with the α4T152 side-chain. Additionally, residues α4H142 and α4Q150, both located on the complementary side of the α-α pocket, contain H-bond donor groups in their side-chains, specifically the NH group in the imidazole ring of α4H142 and the amide NH_2_ group in α4Q150. However, none of the ligands formed H-bonds with these two residues during the simulation (Figure S26). The interaction observed with the side-chain of α4T152, specific to the α-α pocket, highlights the distinct nature of this pocket compared to the α-β one and the uniqueness of the interactions this region can form with agonists. The presence of an α-α pocket in the α4β2 nAChR and its unique interactions lowers the overall receptor sensitivity to agonists like varenicline **1**, nicotine **2** and cytisine **3** (Table 1) but increases their relative efficacy compared to the HS isoform.^46, 78^ Also, the α-α pocket provides binding sites for allosteric modulators, such as NS9283, that target the LS (α4)_3_(β2)_2_ stoichiometry, promoting conformational changes that increase agonist efficacy.^79–81^ The binding mode of allosteric modulators in the α-α interface overlaps with the agonist binding site, albeit the modulators interact mostly with residues on the complementary side of the pockets,^80, 81^ which may explain their inability to activate (α4)_3_(β2)_2_ receptors.^79–81^

The mobility of β2L146 and α4T152 backbone NH groups, as well as the OH group of α4T152, was also analyzed, with no noticeable differences observed across the various wild-type agonist-receptor systems (Figures S27-S28).

### (b) Probing unexplored H-bond interactions within the agonist binding sites of α4β2 nAChR

The MD simulations performed for the wild-type complexes uncovered that β2S133, α4T183 and α4T139 can (albeit to varying extents) sustain hydrogen bonding interactions with certain agonists (Figures 2 and S21). To test whether these interactions are not only present but crucially also are of functional relevance, we ablated the hydrogen bonding potential of these three residues by replacing them with valine, thereby creating the corresponding β2S133V, α4T183V, and α4T139V receptor mutants.

We posited that removal of the hydroxyl donor(s) in β2S133V, α4T183V, and α4T139V would affect agonist binding (particularly for varenicline **1**) to the α4β2 nAChR, resulting in a decrease in their functional potency (EC_50_) and/or relative efficacy (RE). To assess the impact of the threonine/serine-to-valine mutations on the functional profile of varenicline **1**, we obtained activation-concentration curves using oocytes expressing heterologously the HS and LS α4β2 receptors. For comparison purposes, the mutations were also assessed for their impact on the agonist profiles of nicotine **2**, cytisine **3** and ACh (Figures S41-S43). As shown in Figures 3 and S39-S40, with data summarized in Table 1, β2S133V significantly reduced the EC_50_ and RE of varenicline **1** at the α4β2 nAChR, with a decrease in potency at both receptor isoforms of approximately 30-fold. Changes in RE were more pronounced at the LS stoichiometry (with 95% of efficacy loss in LS relative to wild-type compared to a 67% reduction in the HS), although this difference between isoforms could be due to the challenges of accurately measuring the very low efficacy of varenicline **1** in the β2S133V mutant. The α4T183V mutation also affected the EC_50_ and RE of varenicline **1** at both stoichiometries (Figures 3 and S39-S40, Table 1). The effect was less marked than the β2S133V mutation above, with a potency and RE decrease of only 2-fold. Note that the β2S133V mutation is only present in the (two) α-β binding pockets, whereas α4T183V affects both the α-β and α-α sites.

**Figure 3.**
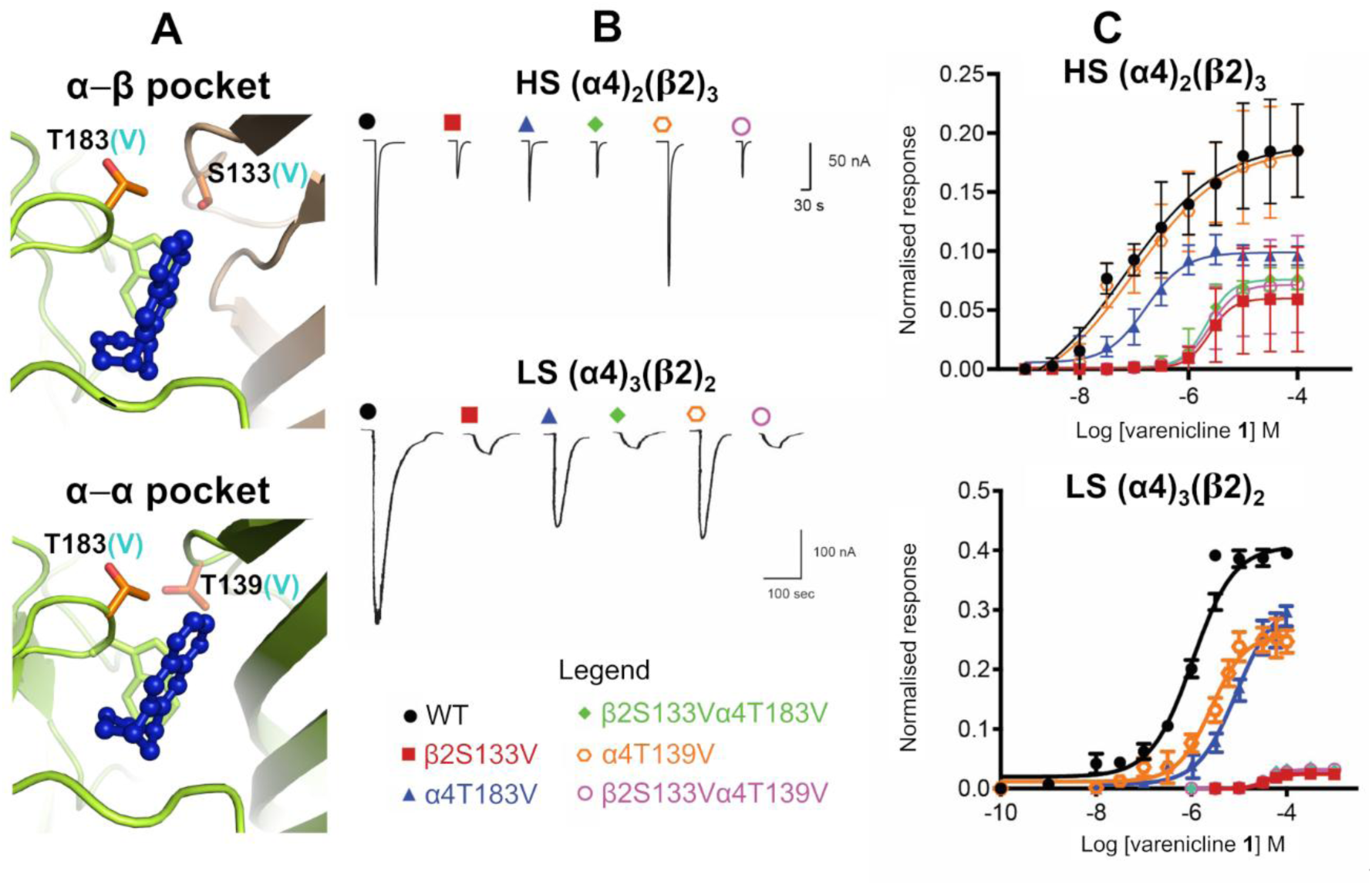
Functional effect of threonine-to-valine mutation (with subsequent side-chain hydroxyl removal) on varenicline 1 agonism at α4β2 nAChR. **A**) Location of α4T183, α4T139, and β2S133 in the α-β (top panel) and α-α (bottom panel) binding pockets of the wild-type α4β2 receptor. Residues α4T183, α4T139, and β2S133 were replaced with valine, as indicated by the cyan-colored “(V)” in the residue labels, resulting in the three single mutants β2S133V, α4T183V, and α4T139V, and the two double mutants β2S133Vα4T183V and β2S133Vα4T139V. The α4 and β2 subunits are colored in yellow and light brown, respectively. Varenicline **1** is highlighted in dark blue. The side-chains of α4T183, α4T139 and β2S133 are represented with orange sticks, whereas TrpB is shown with yellow sticks. **B**) Representative current traces of the current responses elicited by varenicline 2 in oocytes expressing heterologously wild-type or mutant HS or LS α4β2 receptors. The current responses were concentration-dependent. Full concentration-responses curves are shown in panel C. **C**) Varenicline **1** concentration-response curves for wild-type (WT) HS (α4)_2_(β2)_3_ and LS (α4)_2_(β2)_3_ nAChRs (data shown in black) and corresponding mutants β2S133V (data shown in red), α4T183V (data shown in blue), β2S133Vα4T183V (data shown in green), α4T139V (data shown in orange), and β2S133V α4T139V (data shown in magenta). Data points in the concentration-response curves represent the mean ± SEM of 8-10 experiments using 6-8 different *Xenopus* donors. Current responses were measured using two-electrode voltage-clamping from *Xenopus* oocytes heterologously expressing wild-type or mutant (α4)_2_(β2)_3_/(α4)_3_(β2)_2_ nAChRs. Peak current amplitudes for all agonists were normalized to the maximal ACh response (1 mM), as described in the Supporting Information. Estimated potency (EC_50_) and maximal relative efficacy (RE) parameters are shown in Table 1.

Within the α-α binding site, α4T139 occupies a position in the complementary face homologous to the position of β2S133 in the α-β agonist site (Figure 1C). The α4T139V mutant reduced the potency (by approximately 7-fold) and RE (by 50%) of varenicline **1** at the (α-α site-containing) LS (α4)_3_(β2)_2_ receptor (Table 1). However, this mutation has no effect on agonist binding at the HS (α4)_2_(β2)_3_ stoichiometry i.e. plays no role in the function of the orthosteric α-β site (Figure 3, Table 1).

When β2S133V was co-expressed with α4T183V or α4T39V, the changes in varenicline **1** potency and RE were generally no different from those of the single mutant β2S133 (Figures 3 and S39-S40, Table 1), indicating the essential functional role of the β2S133-varenicline **1** interaction in α4β2 receptors.

As shown in Table 1 and Figures S41-S43, none of β2S133V, α4T183V, or α4T139V mutations had a significant impact on the agonism profiles of nicotine **2**, cytisine **3** and ACh at either the HS or LS isoforms of the α4β2 receptor. This is consistent with our MD simulations showing that varenicline **1** is the only agonist that can approach closely the hydroxyl group of β2S133 and make a direct H-bond with this residue (Figures 2B and S21).

To understand the structural and dynamic effect of the serine/threonine-to-valine mutations above, MD simulations for the LS (α4)_3_(β2)_2_ isoform incorporating the β2S133V, α4T183V, α4T139V, β2S133Vα4T183V and β2S133Vα4T139V mutations with varenicline **1**, nicotine **2**, cytisine **3** and ACh were carried out (see Supporting Information for full details), with all complexes remaining stable throughout the simulation(Figures S6-S7). To probe how the mutations affect the dynamics of the receptor, C_α_ atom fluctuations were determined for both wild-type and mutant simulations. The wild-type and mutants C_α_ fluctuation profiles are generally similar, thus indicating comparable dynamics (Figures S9-S10). Despite this, some regions, such as, for example, the Cys loop of the β2S133V complexes, show discernible differences compared to wild-type; however, these differences are not statistically significant, as evidenced by their high *p*-values (Figure S10).

We have also examined the impact of the β2S133V, α4T183V and α4T139V mutations on a range of other relevant interactions associated with ligand-receptor interactions (Figures S18-S21, S26 and S29-S31). In our mutant complexes, all agonists remained in their respective binding sites (Figures S11-S17), forming interactions with TrpB, TyrA, and TrpD, with varying frequencies depending on the mutant (Figures S18-S20). The only exception was the ACh molecule in the second α-β pocket of one β2S133Vα4T139V-ACh replicate, which exited the pocket after approximately 112 ns (Figure S13G).

An analysis of the distance between the H-bond acceptor groups in the agonists and H-bond donors present within the mutants binding pockets was performed to evaluate how the mutations alter the pattern of interactions with the receptor. As anticipated, the threonine-to-valine mutations introduced, and subsequent removal of the residue hydroxyl side-chain, redefined the agonist H-bond network within the binding pockets to varying extents (Figures S29-S31). For instance, in the β2S133V mutant, changes in H-bonding profiles were observed in the α-β pockets, with an increase in the frequency of interaction between α4T183 and cytisine **3** (Figure S29). In the α4T183V and α4T139V mutants, a significant increase in the interactions between the ligands and the backbone NH and side-chain OH group of α4T152 in the α-α pocket is observed, with this increase being especially pronounced for varenicline **1** and cytisine **2** (Figure S30). Furthermore, mutations involving α4T183 and α4T139 within the α-α pocket generally resulted in enhanced interactions between varenicline **1** and the side-chain of α4Q150 but not α4H142 (Figure S31).

### (c) Exploring the structural features of varenicline **1** that contribute to binding and function: a focus on the role of the heteroaryl moiety

In addition to the well-characterized cation-π and H-bond donor contacts associated with the protonated piperidine moiety of varenicline **1**,^54^ potential interactions involving the heteroaryl (quinoxaline) moiety of this ligand have also been identified from structural studies involving soluble AChBPs^49, 57, 58^ and the α4β2 nAChR.^67^ These have, for example, demonstrated a potential for the quinoxaline unit of varenicline **1** to interact as an H-bond acceptor (e.g., with water) in the binding site. However, given the very weakly basic nature of a quinoxaline (p*K*_a_=0.6, which is approx. 3 p*K* units *less* basic than the pyridyl moiety of nicotine **2**; p*K*_a_=3.1),^82^ the functional significance of this interaction remains unclear. To gain a more complete picture of the molecular-level interactions made by the quinoxaline moiety of varenicline **1** at native nAChRs, we have explored the (hetero)aryl-based structural interactions involved in both receptor recognition (binding) and function (gating). Others, in particular the Pfizer medicinal chemistry program, have reported numerous variants on the heteroaryl portion of varenicline **1**.^25, 35, 83–85^ However, these variants generally differ significantly in geometry/size and may incorporate distinct aryl/heteroaryl core features as well as peripheral substituents capable of forming new ligand-receptor interactions. This structural diversity complicates, if not entirely hinders, an accurate assessment of the functional role played by the quinoxaline-involving H-bond(s). We aimed to simplify this issue by varying only the heteroaryl region of varenicline **1** while retaining both overall ligand size and shape, as well as the crucial cation-π/H-bond donor components within the protonated piperidine moiety (Figure 1D). Accordingly, we have synthesized three novel varenicline variants: C_2_ varenicline **4**, isovarenicline **5** and N_2_ varenicline **6**. These new ligands retain a common scaffold - tetracyclic core containing a basic (protonatable) piperidine unit - and differ only in the presence and/or location of the N-based H-bond acceptor component(s) (Figure 4A). Importantly, all three ligands retain essentially the same geometry and volume as the parent compound, varenicline **1**, and do not present any additional or significant interactions associated with the periphery.

**Figure 4.**
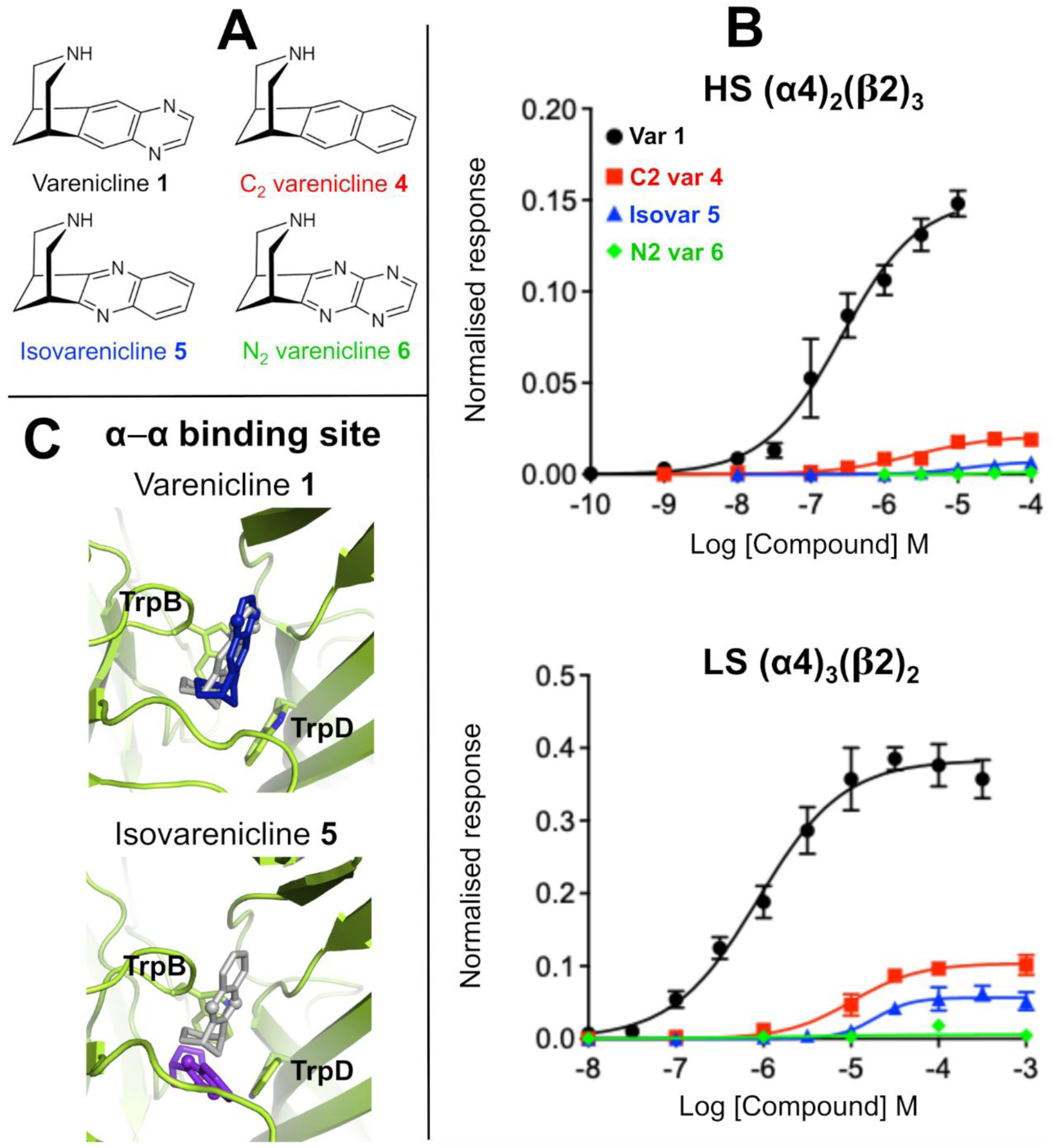
Functional profiles of varenicline 1 and its variants 4-6. **A**) Chemical structure of varenicline **1** (parent compound) and variants C_2_ varenicline **4**, isovarenicline **5** and N_2_ varenicline **6**. **B**) Concentration-response curves for varenicline **1**, C_2_ varenicline **4**, isovarenicline **5** and N_2_ varenicline **6** for the HS (α4)_2_(β2)_3_ and LS (α4)_3_(β2)_2_ receptors. Concentration-response curves for varenicline **1** and its variants were obtained at wild-type HS (α4)_2_(β2)_3_ and LS (α4)_3_(β2)_2_ nAChRs expressed heterologously in *Xenopus* oocytes. Data points in the concentration-response curves represent the mean ± SEM of 8-10 experiments using 6-8 different *Xenopus* donors. Current responses were measured using two-electrode voltage-clamping. Peak current amplitudes for all ligands tested were normalized to maximal ACh response (1 mM) before fitting the data with the Hill equation, as described in the Supporting Information. Estimated parameters EC_50_ and maximal relative efficacy (RE) are shown in Table 3. **C)** The different binding modes adopted by varenicline **1** and isovarenicline **5** in the α-α pocket after 300 ns of simulation. The grey sticks represent the starting binding mode (after energy minimization) for the agonists (see Supporting Information for a detailed description of how complexes were constructed). The final binding pose for varenicline **1** and isovarenicline **5** are colored in blue and purple respectively, with the nitrogen atoms of the quinoxaline moiety highlighted with spheres. TrpB and TrpD are shown with sticks.

The chemistry used to prepare C_2_ varenicline **4**, isovarenicline **5** and N_2_ varenicline **6** is outlined in Scheme 1, and full details are available in the Supporting Information. Each ligand was characterized as the corresponding ammonium salt, which was then used for affinity binding and functional pharmacology studies (Tables 2 and 3).

**Table 2.** Binding affinity constants (K_i_) for varenicline **1** and ligands **4** and **5**. The K_i_ values for the human α4β2, α3β4, and α7 nAChRs (as well as ratios relative to α4β2) are shown. K_i_ value for N_2_ varenicline **6** was not determined due to disruption (and subsequent closure) of the Milan lab as a consequence of the COVID-19 pandemic.

| $K_i$<br>Ligand | $\alpha 4\beta 2$<br>(nM) | $\alpha 3\beta 4$<br>(nM) | $\alpha 3\beta 4/\alpha 4\beta 2$ | $\alpha 7$<br>(nM) | $\alpha 7/\alpha 4\beta 2$ |
| --- | --- | --- | --- | --- | --- |
| Varenicline **1** | 0.46<br>$\pm 0.12$ | 171<br>$\pm 41$ | 372 | 75<br>$\pm 22.5$ | 163 |
| C<sub>2</sub> varenicline **4** | 14.3<br>$\pm 1.6$ | 267<br>$\pm 73$ | 18.7 | 1325<br>$\pm 595$ | 93 |
| Isovarenicline **5** | 5460<br>$\pm 1280$<br>(5.46 μM) | 67700<br>$\pm 37600$<br>(67.7 μM) | 12 | 126800<br>$\pm 86000$<br>(126.8 μM) | 23 |

**Table 3.** Agonist sensitivity of HS (α4)_2_(β2)_3_ and LS (α4)_3_(β2)_2_ to ligands varenicline **1**, C_2_ varenicline **4**, isovarenicline **5** and N_2_ varenicline **6**. Potency (EC_50_) and relative efficacy (RE) of varenicline **1** and its variants at wild-type HS (α4)_2_(β2)_3_ and LS (α4)_3_(β2)_2_ nAChR. EC_50_ values were estimated as described in Supporting Information. RE was determined by normalizing the responses of the varenicline ligands to the maximal current responses elicited by 1 mM ACh (ACh EC_100_) (I/I_max_ACh). Data shown represent the mean ± SEM of n= 8-10 experiments in 6 to 8 different batches of *Xenopus* oocytes. Statistical differences between varenicline **1** and ligands **4**, **5** and **6** were determined by one-way ANOVA followed by a post hoc Dunnett’s test and/or a posthoc Bonferroni multiple comparison test to determine the level of significance between varenicline **1** and ligands **4**, **5** or **6**. *denotes a statistically significant difference (p < 0.05) between varenicline **1** and ligands **4**, **5** or **6**. ND, not determined due to low levels of functional expression (less than 50 nA of ACh maximal currents). These data are also presented in a graphical format in Figure 4B.

| Ligand | HS ($\alpha 4$)<sub>2</sub>($\beta 2$)<sub>3</sub> | | LS ($\alpha 4$)<sub>3</sub>($\beta 2$)<sub>2</sub> | |
| --- | --- | --- | --- | --- |
| | EC<sub>50</sub> μM | RE | EC<sub>50</sub> μM | RE |
| Varenicline **1** | 0.286 ± 0.1 | 0.14 ± 0.012 | 1.01 ± 0.4 | 0.38 ± 0.025 |
| C<sub>2</sub> varenicline **4** | 2.64 ± 0.8* | 0.022 ± 0.0033* | 11.50 ± 1.74* | 0.102 ± 0.012* |
| Isovarenicline **5** | 13.27 ± 2.5* | 0.0071 ± 0.00086* | 26 ± 8.5* | 0.054 ± 0.013* |
| N<sub>2</sub> varenicline **6** | ND | 0.0010 ± 0.0001* | ND | 0.0032 ± 0.0023* |

**Scheme 1.**
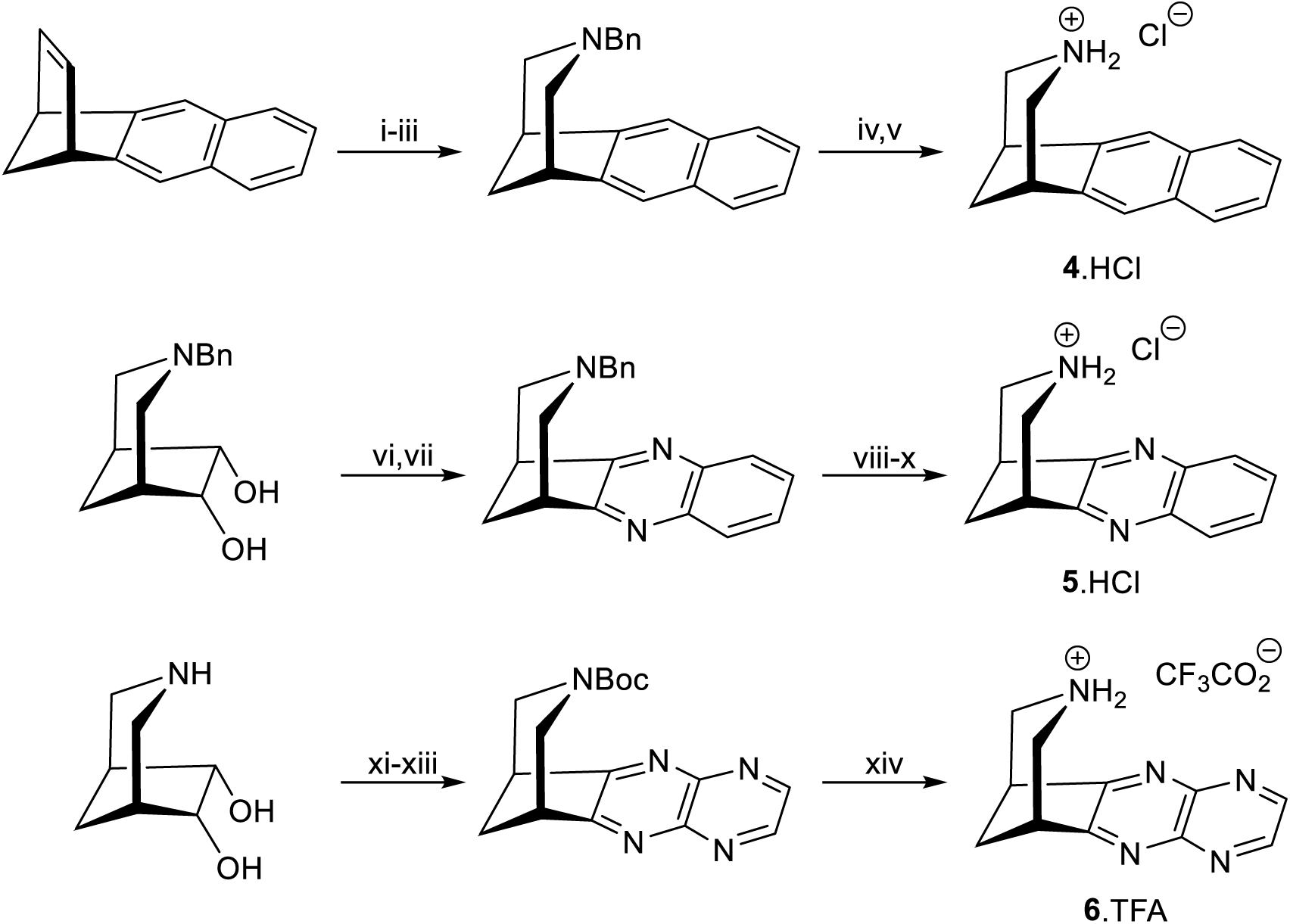
*Reagents*: C_2_ varenicline **4**: i, NMNO, OsO_4_ (cat), acetone, water (98%); ii, NaIO_4_, THF/water; iii, NaBH(OAc)_3_, BnNH_2_ (65% over 2 steps); iv, H_2_, Pd(OH)_2_ (20 wt% on C), Boc_2_O, MeOH/EtOAc (74%); v, HCl in MeOH (quantitative). Isovarenicline **5**: vi, DCC, DMSO, Cl_2_CHCO_2_H, rt, 20h; vii, 1,2-phenylenediamine, rt, 18h, (70% over 2 steps); viii, chloroethyl chloroformate, ClCH_2_CH_2_Cl 80 °C, 18h; ix, (a) MeOH, reflux 2h Boc_2_O, rt, 18h (86% over 2 steps); x, HCl in MeOH, quantitative. N_2_ varenicline **6**: xi, Boc_2_O, Na_2_CO_3_, THF/water, rt, 16h (90%); xii, TFAA, DMSO, CH_2_Cl_2_, then Et_3_N; xiii, 2,3-diaminopyrazine, MeOH, 65 °C, 16h (38% over 2 steps); xiv, 5% TFA in MeOH (69%).

In terms of exploring the profiles of novel variants **4**-**6** alongside the parent compound varenicline **1**, we have assessed: *(i)* ligand affinity constants (K_i_) to the native α4β2 nAChR (as well as the human α3β4 and α7 nAChR subtypes); *(ii)* function (full/partial agonist) profiles at the LS and HS isoforms of the α4β2 nAChR and compared these to varenicline **1**, nicotine **2** and cytisine **3**; *(iii)* the effect of the different aryl/heteroaryl moieties within **4-6** on the p*K*_a_ of the piperidine amine; and *(iv)* the optimized binding modes of variants **4** and **5**, and their dynamics in the LS and HS α4β2 isoforms relative to varenicline **1**.

*(i)* Given a requirement for initial recognition, we have determined the ligand affinity constants (K_i_) for ligands **4** and **5** (together with varenicline **1**) to various nAChR, including the human α4β2 (Table 2). Given that varenicline **1** also interacts with these subtypes, comparative K_i_ data for human α3β4 and human α7 receptors have also been included. C_2_ Varenicline **4**, while weaker than the parent compound, has a K_i_ value within the nanomolar (nM) range and is most potent at the α4β2 subtype; further support for binding of **4** to the α4β2 nAChR comes from the inhibition of ACh by **4** (see Figure S44; Table S2). Intriguingly, however, there is a step-change decrease in K_i_ associated with isovarenicline **5**, which binds to α4β2 in the micromolar (μM) range. Note that although not directly relevant here, but of wider interest, the relative differences observed at α4β2 for **4** and **5**, including reduced subtype-discrimination than observed with varenicline **1**, are also seen at the human α3β4 and α7 subtypes.^86^

Binding affinity data for N_2_ varenicline **6** was not determined due to the disruption caused by the COVID-19 pandemic and other external factors (see Table 2 caption); however, this ligand’s inability to bind was characterized via an indirect approach. Using α4β2 nAChRs expressed heterologously in *Xenopus* oocytes, agonist effects of ligand **6** were determined. As shown in Table 3 and Figure 4B, N_2_ varenicline **6** displayed very poor efficacy at both HS and LS α4β2 nAChR. If a ligand (even with very poor relative efficacy) is *binding* to a receptor target, this will influence the binding of other agonists.^29^ Figure S44 shows that N_2_ varenicline **6** does not affect ACh current response in oocytes heterologously expressing α4β2 nAChRs, which we interpret as indicating that N_2_ varenicline **6** does not bind appreciably to α4β2 nAChR.

*(ii)* To determine functional profile of the new varenicline variants, we have evaluated **1**, **4**-**6** against both the HS and HS stoichiometries of α4β2, with their concentration-response curves as well as EC_50_ and RE values shown in Figure 4B and Table 3.

These data show a similar trend to that associated with K_i,_ with similar patterns within the two α4β2 receptor stoichiometries. C_2_ Varenicline **4** shows a (very) weak agonist profile, which reduces significantly further in the case of isovarenicline **5**, and N_2_ varenicline **6** is essentially inactive.

*(iii)* Given the trends that emerged in Figure 4B, and, in particular, the obvious discontinuity between the profiles of **1** and **4** vs **5**, we have also determined the p*K*_a_ values of the basic piperidine amine center within this ligand series using a spectrophotometric titration method (full details are available in the Supporting Information and Figures S45-S52, with the results summarized in Table 4).

**Table 4.** Experimentally determined p*K*_a_ values for the piperidine amine center of varenicline **1**, C_2_ varenicline **4**, isovarenicline **5** and N_2_ varenicline **6**. The specific salt involved is indicated (based on the final deprotection method used in Scheme 1), and commercially available varenicline tartrate was used. p*K*_a_ values were determined as described in Supporting Information (Table S4).

| Ligand (salt used) | pK<sub>a</sub> | Literature values |
| --- | --- | --- |
| Varenicline **1** (tartrate) | 8.90 ± 0.1 | 9.2,<sup>87</sup> 9.3,<sup>88</sup> 9.22 ± 0.13<sup>89</sup> |
| C<sub>2</sub> varenicline **4** (HCl) | 9.63 ± 0.08 | - |
| Isovarenicline **5** (HCl) | 8.44 ± 0.09 | - |
| N<sub>2</sub> varenicline **6** (CF<sub>3</sub>CO<sub>2</sub><sup>-</sup>) | 7.31 ± 0.05 | - |

As anticipated, the basicity of the ligands decreased in the following order: C_2_ varenicline **4**, varenicline **1**, isovarenicline **5,** and N_2_ varenicline **6**, which correlates with the inductive influence of an increasingly electron-deficient heteroarene on the proton affinity of the basic amine group; the rigid σ-bond scaffold and its alignment to the aryl π-system may enhance the efficiency of this inductive process. Clearly, amine protonation within the binding site (as opposed to bulk solution) is a prerequisite for enabling both the cation-π and H-bond donor interactions (see Figure 1D) that contribute to stabilizing the receptor complex. That said, as discussed below, the basicity of the amine group in the ligands seems to only be relevant for the functional profile of N_2_ varenicline **6**.

*(iv)* Models for the complexes between the ECD of the wild-type HS (α4)_2_(β2)_3_ and LS (α4)_3_(β2)_2_ nAChR and varenicline variants **4** and **5** were constructed. One ligand molecule was placed in each α-α and α-β binding sites in poses corresponding to those observed in the cryo-EM structure of the human α4β2 nAChR-varenicline complex (PDB code: 6UR8^67^). Note that no model for the α4β2 nAChR-N_2_ varenicline **6** system was constructed due to the extremely low level of activity and potency observed for this ligand (Table 3). The complexes containing C_2_ varenicline **4** and isovarenicline **5**, in which the secondary amine is protonated (consistent with the p*K*_a_ determined for this group, see Table 4) to enable cation-π and H-bond donor interactions with TrpB, were relaxed by energy minimization (for details, see Supporting Information). Our models show that C_2_ varenicline **4** and isovarenicline **5** were optimally located within the α4β2 binding sites in essentially the same orientation as the parent compound varenicline **1** (see Figures S4-S5).

Equilibrium MD simulations were then performed to assess the dynamics of ligands **4** and **5** when bound to the wild-type HS and LS α4β2 nAChR and to compare their behavior to varenicline **1** (Figures S32-S33). Like varenicline **1**, variants **4** and **5** remained bound to both the α-β and α-α pockets, forming a stable cation-π interaction with TrpB and sporadic interactions with TyrA and TrpD (Figures S32 and S34). Varenicline **1** and C_2_ varenicline **4**, as expected from their rigid structures, exhibited limited mobility within the binding sites, consistently maintaining similar orientations throughout the simulation (Figures S35-S36). Given that C_2_ varenicline **4** has essentially the same size and shape and displayed similar dynamics as the parent compound varenicline **1**, the changes in activity and potency observed (Table 3) are likely due to the loss of the H-bond interaction between the ligand (as acceptor) and the (donor) protein, specifically with residues β2S133, α4T183 and α4T139. Unlike the quinoxaline unit in varenicline **1**, the naphthalene group of ligand **4** is not an H-bond acceptor.

Isovarenicline **5**, however, exhibited notable differences in dynamics compared to varenicline **1** (Figures S32 and S35). It also showed varying levels of mobility between the α-β and α-α pockets, with the ligand displaying high mobility in the latter (Figure S35) and adopting binding modes that were different from the initial one (Figures 4D and S36). This is likely, we posit, to be due to the distinct H-bond networks formed between the quinoxaline group of ligand **5** and β2S133 and α4T152 side-chains (Figures S37-S38). Specifically, the loss of H-bonding with the OH group of β2S133 in the α-β site and the formation of a new, stable interaction with the hydroxyl side-chain of α4T152 in the α-α pocket are noteworthy (Figures S37-38).

## CONCLUSION

An attractive pharmacological profile and strong track record as a smoking cessation agent^38^ made cytisine **3** an attractive lead for the development of a proprietary and potentially improved therapeutic, which in 2006 led to the launch of varenicline **1** (Chantix®).^18–21^ However, accumulating evidence now suggests that these two ligands are perhaps not as closely related as initially thought, particularly in terms of their functional mechanisms.^26, 29, 31, 49^ While structurally related, pharmacologically varenicline **1** and cytisine **3** should perhaps be considered more as “cousins” rather than “siblings”. Auerbach and co-workers have recently shown that varenicline **1** stands out as the first ligand in the ‘low-efficiency’ category (η = 33%), whereas typical nAChR ligands, such as cytisine **3**, usually show higher efficiencies (η = 41%).^90^ Efficiency (η), which is a measure of the proportion of binding energy converted to gating, likely reflects both ligand structure and the complexity of receptor-ligand interactions that drive binding and receptor activation.^91, 92^

Structurally, both varenicline **1** and cytisine **3** feature a rigid bicyclic core incorporating a piperidyl unit; however, the adjacent heteroaryl moieties are quite different, with varenicline **1** incorporating a quinoxaline and cytisine **2** a 2-pyridone group. Functionally, the two molecules also differ significantly. For instance, although both ligands have very similar full agonist profiles at the α7 nAChR,^26, 29^ their profiles differ at the α4β2 subtype, mainly at the HS (α4)_2_(β2)_3_ isoform.^26, 29^ While varenicline **1** acts as a weak partial agonist of (α4)_2_(β2)_3_, achieving approximately 13%-18% of the maximum efficacy relative to ACh, cytisine **3** is essentially inactive (Table 1).^26, 29^ Another significant feature that differentiates these two ligands is that varenicline **1** is a potent agonist at the human 5-HT_3_ serotonin receptor, with an efficacy of 80% relative to 5-HT whereas cytisine **3** is an antagonist. While the pharmacological requirements for an effective smoking cessation drug (e.g. profile at α7; roles of the (α4)_2_(β2)_3_ vs (α4)_3_(β2)_2_ stoichiometries) are still to be fully elucidated,^93, 94^ off-target effects, such as binding to the 5-HT_3_ receptor, are a significant factor in reducing end-user compliance.^95, 96^ Therefore, understanding the structure and dynamic basis underpinning functional differences between related ligands, such as varenicline **1** and cytisine **3**, and various receptors is paramount. The work of Dougherty and co-workers has demonstrated that, at the molecular level, varenicline **1** does not utilize the same full set of three receptor-ligand interactions as nicotine **2** or even the related compound cytisine **3**.^53, 54, 56^ Based on a selective backbone modification, Dougherty and co-workers showed that, unlike nicotine **2** and cytisine **3**, varenicline **1** does not make a *functional* interaction with β2Leu146.^56^ This important finding prompted our studies and led us to seek and identify alternative interaction(s) to that “missing” third component associated with varenicline **1** function. Our efforts have aimed to provide a more detailed, comprehensive description of the receptor-ligand interaction patterns involving varenicline **1**, in order to enhance our understanding of the mode of action of this ligand and shed light on the differences associated with, e.g., its functional profile at the 5-HT_3_ receptor vs that of cytisine **2**. To pursue this, we have employed a multidisciplinary strategy based on known and novel ligands integrating biomolecular modelling and simulation, chemical synthesis and physiochemical characterization, binding and comprehensive functional studies.

Initially, using the experimental structural data available, we constructed models for the complexes between the ECDs of the LS and HS isoforms of the human α4β2 nAChR and varenicline **1**, nicotine **2**, cytisine **3**, and ACh (Figures 2A and S2-S3). This was followed by extensive MD simulations to uncover potential new functional interactions involving the agonists, particularly varenicline **1**, within the α-β and α-α binding pockets of the α4β2 nAChR. The simulations revealed two new potential interactions associated with the H-bond hydroxyl donor in the side-chain of α4T183 and β2S133 within the principal and complementary faces of the α-β binding site (Figure 2B). In the simulations, these residues can form transient H-bonds with some ligands, including varenicline **1** and nicotine **2**. Note that, unlike α4T183 (which is present on the principal side of both binding sites), the β2S133 on the complementary face of the α-β site is replaced by α4T139 in the α-α pocket (Figure 1C). These three residues all had the potential to interact with the quinoxaline moiety of varenicline **1** and, thereby, provide the “missing” third H-bond (receptor donor/ligand acceptor) interaction (in addition to the cation-π and H-bond associated with the ammonium center of the ligand) characterized in the Dougherty model (Figure 1D). The interactions with α4T183, β2S133 and/or α4T139 also require participation of the ligand (as the H-bond acceptor) and using synthetic chemistry, we have explored further those structural features of varenicline **1** that mediate its binding and function.

Targeted mutagenesis, together with electrophysiological assays, allowed us to assess the relative importance of each of these hydroxyl-containing residues across ACh, varenicline **1** as well as nicotine **2** and cytisine **3**. For this, the polar residues α4T183, β2S133 and α4T139 were mutated to valine, thereby eliminating their side-chain hydrogen bonding potential. Note that by removing the hydroxyl group -an H-bond donor-from their side chains, the serine/threonine-to-valine mutations prevent α4T183, β2S133 and α4T139 from participating in H-bond interactions with the ligands and/or water molecules. However, the backbone amide NH in these residues, which also serves as an H-bond donor, may still engage in hydrogen bonding. The profiles of varenicline **1,** nicotine **2**, cytisine **3** and ACh were then determined in the mutant receptors (Figures 3 and S39-S43; Table 1). Please note that while the α4T139V and β2S133V mutations -both situated on the α4/β2 complementary face-affect the α-α and the α-β sites individually, the α4T183V impacts all binding sites simultaneously due to its position on the principal side of the pocket (Figure 1C).

These studies (Figure 3; Table 1) demonstrated that, overall, none of the mutations introduced (whether individually or in clusters) affect the ACh profile at either the HS (α4)_2_(β2)_3_ or LS (α4)_3_(β2)_2_ isoforms of the α4β2 receptor. This confirms that α4T183, β2S133 and α4T139 do not play a role in ACh function and further supports Dougherty’s observation that β2L146 (or α4T152 in the α-α pocket) likely accounts for the receptor H-bond donor interaction with ACh.^53, 54^

In contrast to the scenario described above for ACh, varenicline **1** exhibited a significantly different response at the β2S133V mutant, displaying a marked reduction in relative efficacy, showing a 3-fold and 21-fold decrease at the HS and LS β2S133V mutant, respectively, along with concomitant shifts towards higher EC_50_ values (Figure 3; Table 1). Note that at the HS (α4)_2_(β2)_3_ wild-type receptor, varenicline **1** is already a relatively weak partial agonist, with an efficacy of only 18% relative to ACh; however, upon introducing the serine-to-valine (β2S133V) mutation in the complementary face of the α-β sites, varenicline **1** shows negligible efficacy (Table 1). Additionally, the impact of the α4T183V and α4T139V mutations on the functional profile of varenicline **1** was generally less pronounced than that observed for the β2S133V mutation. At the α4T183V receptor, varenicline **1** had a modest loss of efficacy and potency, with about a 2-fold and 6-fold decrease, respectively, at HS (α4)_2_(β2)_3_ and LS (α4)_3_(β2)_2_ receptors (Table 1). For the HS form of the α4T139V nAChR, as expected, no effect on the functional profile of varenicline **1** (or any other ligands for that matter) was observed, as this isoform lacks an α-α site; in contrast, the α4T139V mutation had a moderate effect on the efficacy and potency of varenicline **1** in LS isoform, with a level of reduction similar to that observed for the LS α4T183V mutant (Table 1). At the two double mutants, β2S133Vα4T183V and β2S133Vα4T139V, varenicline **1** exhibited again a dramatic reduction in its functional profile (both efficacy and potency), with the magnitude of the changes generally in the same order as to those observed in the single point β2S133V mutant (Table 1).

Based on all the data presented above, we conclude that β2S133 (rather than α4T183 or α4T139), via the H-bonds formed by its side-chain hydroxyl group (as opposed to the backbone NH, which is still present in the mutants), is the dominant and (up until now unidentified) “third component” required to mediate the function of varenicline **1.** Regarding the role of β2S133, α4T183 and α4T139 in the function of nicotine **2** and cytisine **3**, the situation is less clear than varenicline **1**. The effects of the β2S133V, α4T183V and α4T139V mutations, when present, are markedly more muted compared to those observed for varenicline **1** (Table 1). However, for both nicotine **2** and cytisine **3**, the most impactful interaction still appears to be associated with β2S133, although the changes in the ligand functional profiles in the β2S133V-containing mutants were modest (Table 1). We interpret these results as an ability of nicotine **2** and cytisine **3** to access β2S133 in addition to (but not to the exclusion of) β2L146, which nevertheless remains the optimal partner for mediating the functional profiles of these two ligands.

As stated above, the α4T139V mutation, which is the residue associated with the complementary face of the α-α binding site, unsurprisingly had no effect on the functional profiles of the ligands studied at HS (α4)_2_(β2)_3_ and, at best, showed only a minimal impact at the α-α-containing LS (α4)_3_(β2)_2_ stoichiometry, depending on the ligand used (Table 1). This is consistent with the findings of, for example, Balle and co-workers, who have previously shown that α4T139A has only a modest impact on the maximal efficacy of NS9283.^79^

If we consider that, as with the α-β site, a third anchoring connection is required to potentiate activity via the α-α site, then the residue(s) responsible for mediating this interaction remain yet undefined. The α-α and α-β binding sites are structurally different on the complementary face, as the former is formed by a α4 subunit and the latter by a β2 one, with these differences resulting in distinct pharmacological profiles for each site.^47, 48^ Examples of these differences include the substitution of β2S133, β2V136, β2F144, and β2L146 located in loop E of the α-β site by α4T139, α4H142, α4Q150, and α4T152 in the α-α site, respectively.^48, 56, 76, 77^ The simulations performed in this study indicate that the hydroxyl moiety of α4T152 in the α-α site can directly interact with the H-bond acceptor groups of certain agonists (Figure S30), therefore suggesting that this residue may potentially serve as the missing “third connection” in this agonist binding site. Previous experimental studies have demonstrated that the α4T152A and α4T152V mutations can decrease the activity of NS9283 several fold,^79^ supporting the hypothesis that α4T152 can indeed sustain interactions with some ligands. Additionally, given that α-α site residues, such as α4W88 and α4H142, have been identified as key for the gating efficiency of the LS (α4)_3_(β2)_2_ receptor,^59, 79^ it would be prudent to say that the precise role of α4T152 in modulating the functional profile of different agonists within the LS α4β2 isoform requires further investigation.

The structure of varenicline **1**, along with its detailed comparison to cytisine **3**, also merits comment. As explained above, the bicyclic piperidine moiety is common to both varenicline **1** and cytisine **3**, but their adjacent heteroaryl units, quinoxaline and 2-pyridone, respectively, differ significantly in terms of the spatial relationships associated with these H-bond acceptor elements. However, given the relatively non-basic nature of the quinoxaline moiety, to fully comprehend how varenicline **1** exerts its action, two fundamental questions must be addressed: (*i*) is the quinoxaline unit as present in varenicline **1** required for both binding *and* function, or do those characteristics rely primarily on rigidity and overall shape?; (*ii*), if the quinoxaline group is essential for function, what evidence is available to support the role of the quinoxaline as an H-bond acceptor within the context of nAChRs?

To answer these questions, we have designed a targeted set of novel varenicline variants, namely C_2_ varenicline **4,** isovarenicline **5**, and N_2_ varenicline **6**. These new ligands, which all incorporate a basic piperidine center to retain the critical cation-π and H-bond donor characteristics, maintain the same size, shape and molecular volume as the parent compound, varenicline **1** (Figure 4A). This enabled us to avoid added complications associated with additional peripheral substituents to probe receptor-ligand recognition *and* function, and the role played by the aryl moiety in these two connected processes.

C_2_ Varenicline **4**, which differs from varenicline **1** in containing a naphthalene instead of a quinoxaline group (Figure 4A), displays a K_i_ of 14.3 nM, corresponding to an approximately 30-fold decrease in affinity compared to the parent compound (Table 2). In terms of function, C_2_ varenicline is a very weak partial agonist showing a 10-fold increase in EC_50_ at both HS and LS isoforms of α4β2, together with reduced efficacy, with a 6- and 4-fold reduction at HS (α4)_2_(β2)_3_ and LS (α4)_3_(β2)_2_ respectively (Table 3). We conclude from these results that while the quinoxaline moiety (as present in varenicline **1**) enhances *binding*, it is not a prerequisite, with size/rigidity/shape (and the interactions associated with the ammonium center) being essential determinants for recognition. Further, the MD simulations performed suggest that when bound to the α4β2 nAChR, C_2_ varenicline **4** exhibits comparable dynamics and adopts similar binding modes to those of varenicline **1** (Figures S35-S36), with the biggest differences between the two ligands arising from the lack of the H-bonds associated with the quinoxaline residue. However, the presence of a quinoxaline unit within varenicline **1** is an essential requirement for *function* (Table 3), therefore showing that this moiety contributes to an additional interaction(s) that is unavailable to C_2_ varenicline **4**.

In the case of isovarenicline **5**, where the orientation of the quinoxaline moiety has been reversed within the ligand scaffold (Figure 4A), a step change in both binding (K_i_) and function at α4β2 was observed (Tables 2 and 3). Ligand **5** only binds very weakly to α4β2 (high μM, which is an observation that carries across to both the α3β4 and α7 subtypes) and also shows markedly reduced efficacy, with a 20-fold and 7-fold decrease at HS (α4)_2_(β2)_3_ and LS (α4)_3_(β2)_2_, respectively. The simulations associated with isovarenicline **5** align well with the experimental findings, revealing that this ligand is less stable when bound to α4β2, adopting configurations markedly different from varenicline **1** (Figures 4C and S35-S36) and, importantly, forming alternative interaction networks within the binding pockets (Figures S37-S38). A clear distinction between the interaction networks for varenicline **1** and isovarenicline **5** is the loss of H-bonding to the side chain of β2S133 and the emergence of a frequent interaction with the side chain of α4T152 in the latter (Figures S37–S38). As previously discussed, the interaction between varenicline **1** and β2S133 is essential for the ligand functional activity (Table 1). The absence of this key interaction in isovarenicline **5** likely contributes to its significantly diminished efficacy. Taken together, the experimental and computational findings, allow us to answer the first of the two questions posed above, namely whether the quinoxaline in varenicline **1** is necessary for both binding and function. Our results indicate that both the incorporation and specific arrangement/orientation of the quinoxaline unit within the scaffold of varenicline **1** is essential for function.

The results reported here for C_2_ varenicline **4** and isovarenicline **5** shed light on the second question raised above regarding the role of the interactions involving the quinoxaline moiety of varenicline **1**. Our findings clearly demonstrate that H-bonding, either direct or water-mediated, between the quinoxaline group and the protein is indeed crucial for function. Further compelling experimental evidence that this “missing third” varenicline **1** interaction is based on quinoxaline acting as a H-bond acceptor comes from crystallographic data for the *Capitella teleta* AChBP-varenicline **1** complex (PDB code:4AFG),^58^ an analogous complex with the serotonin binding protein, 5-HTBP, from *Aplysia californica* (PDB code: 5AIN),^49^ as well as the crystal structure of varenicline **1** bound to the iCytSnFR cytisine sensor precursor binding protein (PDB code: 7S7X)^97^. In all of these structures, although they are not functionally significant, water (and also tyrosine)-varenicline **1** interactions involving the quinoxaline N atoms as a H-bond acceptor are evident. Furthermore, our findings here underscore the importance of the precise spatial positioning of the quinoxaline unit in varenicline **1** as its interactions with the protein must occur within a specific, well-defined region of the binding site. Altering the orientation of the quinoxaline group within a ligand (as done in isovarenicline **5**) substantially disrupts the ligand-receptor interaction network and subsequently, ligand function.

N_2_ Varenicline **6** incorporates a 1,4,6,8-tetraazanaphthalene moiety, which we anticipated to be significantly less available as an H-bond acceptor than the quinoxaline of varenicline **1** (Figure 4A).^98^ N_2_ Varenicline showed no evidence of binding, as there was no concentration-dependent inhibition by ligand **6** of ACh function (Figure S44), nor was activity detected at either HS (α4)_2_(β2)_3_ or LS (α4)_3_(β2)_2_ isoforms (Table 3). A likely explanation for this lack of binding is presented below, and lies in the experimentally determined p*K*_a_ of the piperidine group in N_2_ varenicline **6** (Table 4).

Given that piperidine protonation in varenicline **1** is a prerequisite for the cation-π and H-bond donor/acceptor interactions with TrpB in the principal side of the α-β pockets,^54^ the affinity constants (K_i_) and p*K*_a_ values for varenicline **1** and variants **4**-**6** were determined (Tables 2 and 4, respectively). A significant decrease in K_i_ was observed between C_2_ varenicline **4** (p*K*_a_=9.63) and isovarenicline **5** (p*K*_a_=8.44), despite no meaningful differences in the protonation state of their ammonium centers at physiological pH. Based on the experimentally determined p*K*_a_ values, both ligands remain ≥90% protonated at pH 7.5 (Table S5). Therefore, the observed ligand affinity of C_2_ varenicline **4** vs isovarenicline **5** is not adequately explained by the p*K*_a_ difference in the piperidine moieties. Moreover, isovarenicline **5** is more basic than either nicotine **2** (pyrrolidine p*K*_a_=7.80)^99^ or cytisine **3** (p*K*_a_=8.20),^100^ further supporting that protonation differences are not responsible for its reduced binding affinity. Rather, as explained above, the functional profile of C_2_ varenicline **4** can be attributed to the inability of the naphthyl moiety to provide the additional H-bond acceptor required for functional activity, despite exhibiting dynamics and binding poses similar to that of the parent compound varenicline **1**. In contrast, the behavior of isovarenicline **5** is intriguing. The decrease in its functional profile is associated with distinct interaction patterns that isovarenicline **5** forms within the binding sites compared to varenicline **1**. These encompass, for example, the formation of novel contacts with α4T152 and the loss of those involving β2S133 (Figures S37-S38). This finding highlights the importance of the precise H-bond acceptor position within varenicline **1**, where an interaction with β2L146 is precluded by distance, yet the ligand can compensate by engaging in an alternative functional mechanism via β2S133. Finally, the very poor H-bond acceptor characteristics of the 1,4,5,8-tetraazanaphthyl group in N_2_ varenicline **6** would suggest a functional profile resembling that of naphthyl-based C_2_ varenicline **4**. However, this is not the case; the very low binding affinity and negligible activity of N_2_ varenicline **6** are likely associated with the low p*K*_a_ for the piperidine amine (p*K*_a_=7.31), suggesting that protonation may be impaired.^101^

Given that the viability of varenicline **1** to participate as a H-bond acceptor within similar receptor binding environments is now firmly established, and based on the data for the mutants above (Table 1), we suggest that the key interactions of the quinoxaline moiety of varenicline **1** involve β2S133 in the complementary side and (to a lesser degree) α4T183 in the principal side of the α-β binding sites within the α4β2 nAChR. Furthermore, building on previous findings by Dougherty and colleagues showing that varenicline **1** does not rely on β2L146 to enable *function*,^56^ we posit that the binding model of varenicline **1** to the α-β sites of the α4β2 subtype, which retains both the cation-π and H-bond donor components derived from the protonated secondary amine center in the ligand, can now be expanded to include a H-bond donor-acceptor interaction (as the “missing” third component) and that this previously unrecognized interaction involves the quinoxaline heterocycle with chiefly β2S133 as illustrated in Figure 5.

**Figure 5:**
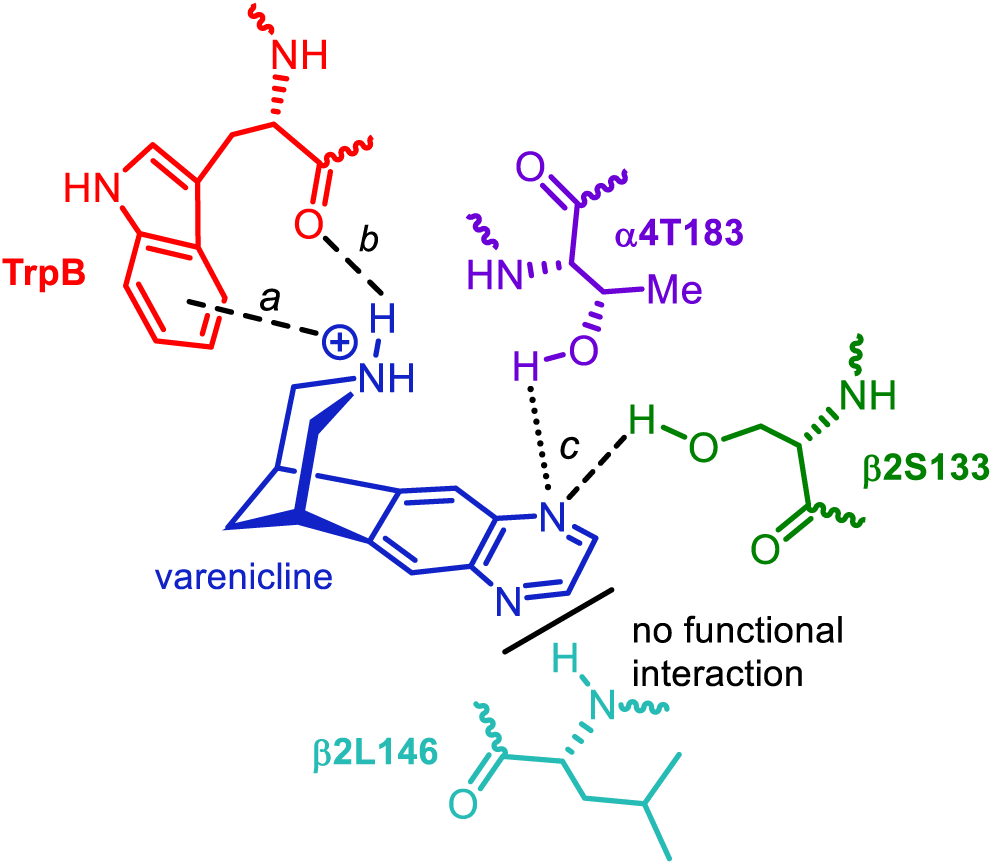
Expanded Dougherty-Lester nAChR binding model for varenicline **1** in the α-β pocket of α4β2 nAChR, highlighting the three key functional interactions: (*a*) cation-π, (*b*) backbone C=O as H-bond acceptor associated with TrpB within the α4 subunit (as defined previously by Dougherty and Lester),^51–53^ and the newly identified (*c*) H-bond donor-acceptor interaction between the quinoxaline moiety of varenicline **1** and the side-chain OH group of b2S133 (complementary subunit). The less prominent interaction involving the OH of a4T183 is also included in the scheme, as it can occur in the absence of a functional connection involving the backbone NH of b2L146. Note that although the new ligand-protein contacts involving b2S133 and a4T183 are depicted as being direct, the possibility of a bridging water molecule mediating the interactions cannot be excluded.

In conclusion, we have uncovered and characterized the to-date “missing interaction” that modulates the partial agonist profile of varenicline **1** at the α4β2 receptor, one of the most prevalent neuronal subtypes in humans and a primary target in nicotine addiction. We have demonstrated that, despite its limited basicity, the quinoxaline moiety within varenicline **1** does indeed engage in functionally relevant interactions with the α4β2 nAChR. In the α-β binding site of this subtype, these interactions are highly specific and well-defined, likely involving the hydroxyl side chains of β2S133 and α4T183. Given that H-bond donor groups at positions equivalent to β2S133 and α4T183 are highly conserved across human neuronal nAChR subunits (Figures 2D and S22), it is plausible to suggest that these newly characterized interactions contribute to the functional profile of varenicline **1** (and potentially other agonists) across a range of neuronal subtypes, such as α7 and α3β4. This, in our opinion, warrants further investigation. Uncovering the functional interactions of each agonist (and how these vary across receptor subtypes) will strengthen the foundation for rational drug design targeting these proteins. This is particularly relevant in the human 5-HT_3_ receptor, where cytisine **3** (IC_50_=0.5mM) is 2000-fold less potent than varenicline **1** (IC_50_=0.25 μM).^102^ The ability of varenicline **1** to define alternative and functionally relevant binding patterns with the receptors, distinct from that of its progenitor cytisine **3**, may directly account for its capacity, unlike cytisine **3**, to activate the 5-HT_3_ receptor.

## ACKNOWLEDGEMENTS

ASFO was supported at the University of Bristol by a BBSRC Discovery Fellowship ([BB/X009831/1]). FV was supported by ANID Becas Chile 72210124. We thank Achieve Life Sciences for a gift of (‒)-cytisine, EPSRC (EP/N024117/1) for financial support, and TG thanks Dr Jack Rogers for support and Professor Varinder Aggarwal for access to laboratory facilities at the University of Bristol. We thank EPSRC for providing ARCHER2 time via HECBioSim (hecbiosim.ac.uk). Data analysis was conducted using the facilities of the Advanced Computing Research Centre at the University of Bristol (https://www.bris.ac.uk/acrc/).

## ASSOCIATED CONTENT

### Supporting Information

The Supporting Information is available free of charge at https://pubs.acs.org/doi/xxxxx. Full experimental details for the synthesis and characterization of the varenicline variants **4**-**6** reported, including copies of ^1^H and ^13^C NMR (and where relevant 2D and ^19^F NMR) and the crystallographic determination of N-Boc isovarenicline **S18 (**and CIF); full details of the molecular dynamics simulations conducted and their analysis, together with supporting figures and tables; nAChR ligand binding affinity measurements; nAChR functional studies, including human α4β2 nAChR expression, single and double point mutation studies and details of electrophysiology studies plus supporting figures; p*K*a measurement studies via spectrophotometric titration.

## DATA AVAILABILITY

All MD data (including input and trajectory files) will be publicly available upon publication via the University of Bristol Research Data Repository (https://data.bris.ac.uk/).

## AUTHOR INFORMATION

### Corresponding Authors

**A. Sofia F. Oliveira** - School of Chemistry and Computational Chemistry Centre, School of Chemistry, University of Bristol, Bristol BS8 1TS, United Kingdom;

**Isabel Bermudez** - Department of Biological and Medical Sciences, Oxford Brookes University, Oxford OX3 0BP, United Kingdom;

**Timothy Gallagher** - School of Chemistry and Computational Chemistry Centre, School of Chemistry, University of Bristol, Bristol BS8 1TS, United Kingdom;

### Authors

**Sheenagh G. Aiken** - School of Chemistry and Computational Chemistry Centre, School of Chemistry, University of Bristol, Bristol BS8 1TS, United Kingdom;

**Daniele Fiorito** - School of Chemistry and Computational Chemistry Centre, School of Chemistry, University of Bristol, Bristol BS8 1TS, United Kingdom;

**Matthew Harper** - School of Chemistry and Computational Chemistry Centre, School of Chemistry, University of Bristol, Bristol BS8 1TS, United Kingdom;

**Grzegorz Pikus** - School of Chemistry and Computational Chemistry Centre, School of Chemistry, University of Bristol, Bristol BS8 1TS, United Kingdom;

**Juno Underhill** - School of Chemistry and Computational Chemistry Centre, School of Chemistry, University of Bristol, Bristol BS8 1TS, United Kingdom;

**Jacob Murray -** Department of Chemistry, Durham University, South Road, Durham DH1 3LE, United Kingdom;

**Joshua Rawlinson -** Department of Chemistry, Durham University, South Road, Durham DH1 3LE, United Kingdom;

**AnnMarie C. O’Donoghue** - Department of Chemistry, Durham University, South Road, Durham DH1 3LE, United Kingdom;

**Cecilia Gotti -** CNR, Institute of Neuroscience, University of Milan, I-20129 Milan, Italy;

**Sarah Lummis -** Department of Biochemistry, University of Cambridge, Cambridge, CB2 1QW, United Kingdom;

Teresa Minguez Viñas - Department of Biological and Medical Sciences, Oxford Brookes University, Oxford OX3 0BP, United Kingdom;

**Franco Viscarra -** Department of Biological and Medical Sciences, Oxford Brookes University, Oxford OX3 0BP, United Kingdom;

### NOTES

The authors declare no competing financial interest.

