## Supporting Information for "How Varenicline Works: Identifying Critical Receptor and Ligand-based Interactions"

### Table of Contents


**A. Synthetic Chemistry** **SI 3 – SI 40**

- *(i)* General information
- *(ii)* Synthetic procedures and characterization data; x-ray crystallographic details of N-Boc isovarenicline **S18**
- *(iii)* Supporting figures: $^{1}H$ and $^{13}C$ NMR spectra of key intermediates and final products

**B. Computational Modelling** **SI 41 – SI 85**

- *(i)* Molecular dynamics (MD) simulations
- *(ii)* Analysis of MD simulations
- *(iii)* Supporting figures and tables

**C. nAChR Ligand Binding Measurements** **SI 86 – SI 87**

- *(i)* Expression of human α4β2, α3β4 and α7 nAChR
- *(ii)* Radioligand binding assays
- *(iii)* Competition binding assays
- *(iv)* Statistical analysis

**D. nAChR Methods and Functional Studies** **SI 88 – SI 96**

- *(i)* Animals
- *(ii)* Human α4β2 nAChR expression in *Xenopus* oocytes
- *(iii)* Single and double mutations
- *(iv)* Electrophysiological recordings
- *(v)* Statistical analysis
- *(vi)* Supporting figures and tables

**E. p$K_a$ Determinations** **SI 97 – SI 106**

- (*i*) Experimental description of the materials and assays
- (*ii*) Theoretical background
- (*iii*) Spectrophotometric method validation: 4-dimethylaminopyridine (DMAP) and nicotine **2**
- (*iv*) Spectrophotometric titration of varenicline **1**, nicotine **2**, cytisine **3** and varenicline variants **4**-**6**

**F. Supporting Information References** **SI 107 – SI 110**

#### A. Synthetic Chemistry

##### *(i)* General information

Reactions requiring inert conditions were conducted under an $N_2$ atmosphere using standard Schlenk-line techniques. Anhydrous solvents were obtained from an Anhydrous Engineering alumina column drying system or from distillation following standard procedures. All other reagents were purchased from commercial suppliers and used as received. Thin layer chromatography was performed using aluminum backed 60 F254 silica plates. Visualization was achieved by UV fluorescence or a basic $KMnO_4$ solution and heat.

Infrared spectra were recorded using a Perkin Elmer Spectrum Two FT-IR spectrometer.

NMR spectra were recorded on Bruker Advance III HD 500 Cryo, Varian 400-MR, Jeol ECS 400 or JEOL ECZ 400 spectrometers. Chemical shifts (δ) are quoted in parts per million (ppm) and are referenced to the residual solvent peak, coupling constants (*J*) are given in Hz. Multiplicities are abbreviated as: br (broad), s (singlet), d (doublet), t (triplet), q (quartet), m (multiplet) or combinations thereof. Assignments (when indicated) were made with the aid of COSY, HSQC, HMBC experiments.

Mass spectrometry was performed by the University of Bristol mass spectrometry service by either ($EI^+$) using a VG Micromass Autospec spectrometer or by electrospray ionization (ESI+) using a Bruker Daltonics MicrOTOF II spectrometer.

Numbering system (shown here for varenicline) as used for $^1H/^{13}C$ NMR structural assignments of varenicline variants.

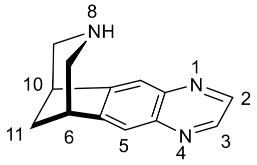

Varenicline

Structures are numbered based on the Schemes shown below except where a structure (varenicline variants) appears in the published manuscript, and the number used here corresponds to that used in the main paper.

#### *(ii)* Synthetic procedures and characterization data

##### 1. $C_2$ Varenicline 4

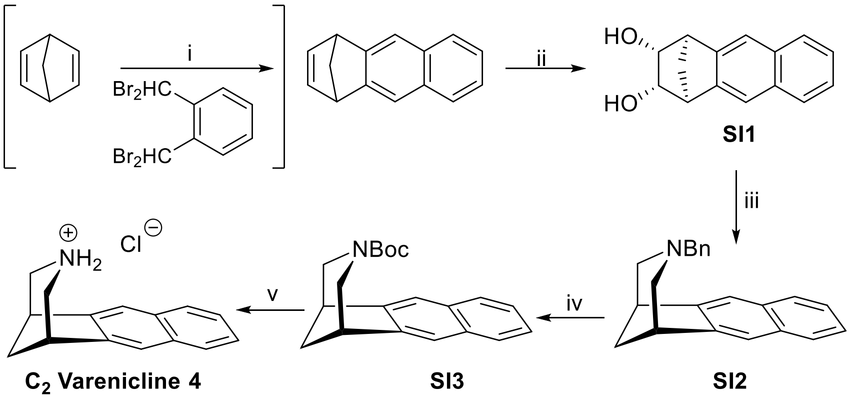


**SI Scheme 1**: *Reagents*: i, KI, xylene, 65 °C, 18h[1] (35%) [not included in main manuscript]; ii, NMNO, $OsO_4$ (cat), acetone, water (98%); iii, (a) $NaIO_4$, THF/water[2, 3] then (b) $NaBH(OAc)_3$, $BnNH_2$ (65% over 2 steps); iv, $H_2$, $Pd(OH)_2$ (20% wt on C), $Boc_2O$, MeOH/EtOAc (74%); v, HCl in MeOH (quantitative).

##### *N*-Bn $C_2$ Varenicline SI2

*exo*-1,4-Methano-1,2,3,4-tetrahydro-2,3-dihydroxyanthracenedihydroxyanthracene **SI1** (222 mg, 0.98 mmol) was dissolved in THF:$H_2O$ (2.5:1, 20 mL) then $NaIO_4$ (230 mg, 1.08 mmol) was added in a single portion. The mixture was stirred at rt for 30 minutes then $H_2O$ (40 mL) was added. The aqueous phase was extracted with $CH_2Cl_2$ (4 × 10 mL), the extracts were dried ($MgSO_4$) and concentrated to give the crude dialdehyde as a yellow oily paste. The dialdehyde was dissolved in anhydrous $CH_2Cl_2$ (20 mL) under an $N_2$ atmosphere then cooled to 0 °C. Sodium triacetoxyborohydride (830 mg, 3.92 mmol) was added followed by benzylamine (118 µL, 1.08 mmol) dropwise. The mixture was allowed to slowly warm to rt overnight, quenched with saturated aq. $Na_2CO_3$ (15 mL) and $H_2O$ (20 mL). The product was extracted with $CH_2Cl_2$ (3 × 20 mL), the combined extracts were washed with brine (30 mL), dried ($MgSO_4$) and concentrated. Purification by silica chromatography (Biotage; 5% to 15% EtOAc in hexane) gave the ***N*-Bn $C_2$ varenicline SI2** (190 mg, 65%) as a colorless oil. $^1H$ NMR (400 MHz, $CDCl_3$) δ 7.83 – 7.76 (m, 2 H, C10/13-H), 7.55 (s, 2 H, 5/12-H), 7.44 – 7.37 (m, 2 H, 2/3-H), 7.14 – 7.06 (m, 3 H, Ph), 6.90 – 6.81 (m, 2 H, Ph), 3.48 (s, 2 H, $PhCH_2$), 3.24 (t, *J* 4.4 Hz, 2 H, 6/10-H), 2.98 – 2.89 (m, 2 H, 7/9-H), 2.53 (d, *J* = 10.3 Hz, 2 H, 7/9-H), 2.27 (m, 1 H, 11-H), 1.79 (d, 1 H, *J* 10.5

Hz, 11-H); $^{13}$C NMR (126 MHz, $CDCl_3$) δ 146.0 (C5a), 138.8 (Ph), 133.5 (C4a), 128.5 (Ph), 128.1 (Ph), 127.8 (C1), 126.6 (Ph), 124.8 (C2), 119.3 (C5), 61.8 (PhCH2), 57.7 (C7), 43.6 (C11), 41.3 (C6); HRMS (ESI): calculated for $C_{22}H_{22}N$ $[M+H]^+$: 300.1747, found: 300.1733.

##### ***N*-Boc C$_2$ Varenicline SI3**

To a solution of *N*-Bn $C_2$ varenicline **SI2** (176 mg, 0.588 mmol) in MeOH:EtOAc (1:1, 12 mL) was added $Boc_2O$ (0.270 mL, 1.17 mmol) and $Pd(OH)_2$ (20 wt% on carbon, 83 mg). The mixture was stirred rapidly under an atmosphere of hydrogen at room temperature for 24 h, after which the mixture was filtered through Celite. The solids were washed with $CH_2Cl_2$ (50 mL), the filtrate was concentrated and purification by silica chromatography (Biotage; 2% to 30% EtOAc in hexane) gave ***N*-Boc C$_2$ varenicline SI3** (135 mg, 74%) as a colorless solid. FTIR $v_{max}$ / $cm^{-1}$ (neat): 1691; $^1$H NMR (400 MHz, $CDCl_3$; broadening and splitting of some signals due to amide resonance was observed) δ 7.80 – 7.75 (m, 2 H, 1/4-H), 7.64 (s, 1 H, 5/12-H), 7.61 (s, 1 H, 5/12-H), 7.43 – 7.37 (m, 2 H, 2/3-H), 4.07 (d, *J* 12.5 Hz, 1 H, 7/9-H), 3.94 (d, *J* 12.5 Hz, 1 H, 11-H), 3.36 – 3.25 (m, 3 H, 2 × 6/10-H, 7/9-H), 3.21 (d, *J* = 12.4 Hz, 1 H, 7/9-H), 2.35 (m, 1 H, 11-H), 1.94 (d, *J* 10.8 Hz, 1 H, C11-H), 1.16 (s, 9 H); $^{13}$C NMR (126 MHz, $CDCl_3$) δ 156.1 (C=O), 143.9/143.8 (C5a, rotamers), 133.64/133.59 (C4a, rotamers), 128.1/127.7 (C1, rotamers), 125.3/125.2 (C2, rotamers), 121.2/120.6 (C5, rotamers), 79.3 ($CCMe_3$), 50.5/49.4 (C7, rotamers), 41.6 (C11), 40.1/40.0 (C6, rotamers), 28.3 ($CCMe_3$); HRMS (ESI): calculated for $C_{20}H_{24}NO_2$ $[M+H]^+$: 310.1802, found: 310.1814.

##### **C$_2$ Varenicline hydrochloride salt 4**

*N*-Boc $C_2$ varenicline **SI3** (24 mg, 0.078 mmol) was dissolved in HCl (4 mL, 0.5 M in MeOH) and allowed to stand for 18 h at rt. The mixture was concentrated to afford the title compound **C$_2$ varenicline hydrochloride 4** (19 mg, quantitative) as an off-white solid. $^1$H NMR (400 MHz, $D_2O$), δ 8.02 – 7.96 (m, 2H, 1-H), 7.94 (s, 2H, 5-H), 7.63 – 7.57 (m, 2H, 2-H), 3.67 – 3.60 (m, 2H, 7-H), 3.52 (d, *J* = 12.3 Hz, 2H, 6/10-H), 3.37 (d, *J* = 12.3 Hz, 2H, 6/10-H), 2.44 (m, 1H, 11-H), 2.20 (d, *J* = 11.6 Hz, 1H, 11-H); $^{13}$C NMR (126 MHz, $D_2O$) δ 140.5 (C4), 133.6 (C6), 128.0 (C7), 126.2 (C8), 122.5 (C5), 47.8 (C1), 40.2 (C3), 38.0 (C2); HRMS (ESI+): Calculated for $C_{15}H_{16}N$ $[M+H]^+$: 210.1283, found: 210.1268.

#### 2. Isovarenicline 5

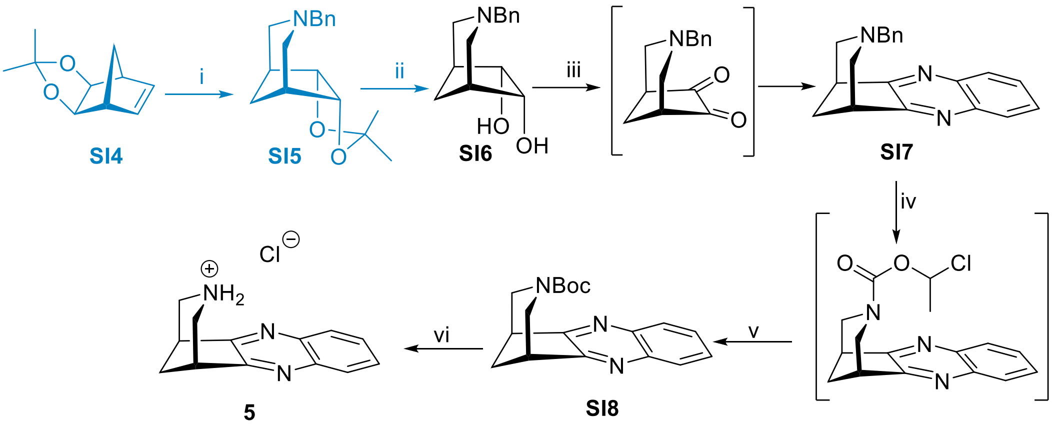


**SI Scheme 2**: *Reagents*: i, (a) $O_3$, DCM, -78 °C then $Me_2S$ (b) $NaBH(OAc)_3$, $BnNH_2$, DCM, rt, 35% overall; ii, HCl, THF, 81%; iii, (a) DCC, DMSO, $Cl_2CHCO_2H$, rt, 20h; iv, (b) 1,2-phenylenediamine, rt, 18h, 70% over 2 steps; vi, chloroethyl chloroformate, $ClCH_2CHCl$ 80 °C, 18h; v, (a) MeOH, reflux 2h (b) $Boc_2O$, 86% over 2 steps; vi, HCl in MeOH, quantitative OR (+)-tartaric acid, EtOH/acetone 76%

Note that diol **SI6** was available commercially but synthetic details for conversion of S14 to S16 are provided here given the specialized nature of the commercial supply used.

##### N-Benzyl 3-azabicyclo[3.2.1]octane-6,7-diol acetonide S15

To a solution of acetonide **SI4**[4] (1.0 g, 6.0 mmol) in DCM (12 mL), ozone was bubbling at -78 °C until the blue colour appeared/persisted. The excess of ozone was removed by a stream of nitrogen at -78°C, and after that, dimethylsulfide (0.95 mL, 12.4 mmol) was added. The reaction mixture warm to rt and stirred for 16h. The solvent was removed to give a transparent oil, which was dissolved in dry DCM (80 mL) and sodium triacetoxyborohydride (5.10 g, 24.18 mmol) was added. The mixture was cooled to 0 °C and a solution of benzylamine (0.71 g, 0.73 mL) in dry DCM (20 mL) was added dropwise over 30 minutes. The mixture was warmed to rt, stirred for 18h, then washed with water (30 mL) and brine (30 mL) and dried ($MgSO_4$). The solvent was removed and the crude product was purified by silica gel chromatography (ethyl acetate:hexane) to give **acetonide SI5** (575 mg, 35%) as a colorless oil, which was used without additional purification. $^1H$ NMR (400 MHz, $CDCl_3$): $\delta$ 7.25-7.13 (m, 5H), 4.44 (d, *J*=1.6 Hz, 2H),

3.33 (s, 2H), 2.67-2.63 (m, 2H), 2.08 (t, *J* 4 Hz, 2H), 1.99 (d, *J* 12 Hz, 2H), 1.87 (m, 1H), 1.36 (s, 3H), 1.28 (s, 3H), 1.09 (d, *J* = 12 Hz, 1H); $^{13}$C NMR (100 MHz, $CDCl_3$): $\delta$ 138.6, 128.6, 128.2, 126.9, 108.5, 83.4, 62.5, 56.1, 40.3, 31.6, 25.9, 23.8; $R_f$: 0.70 (30% ethyl acetate in hexane).

**N-Benzyl 3-azabicyclo[3.2.1]octane-6,7-diol SI6**

Acetonide **SI5** (663 mg, 2.43 mmol) was dissolved in THF (12 mL) and 4M HCl (12 mL) was added. The mixture was stirred at 80°C for 72h, after which time 10% aq. $NaHCO_3$(30 mL) was added and pH was adjusted to pH 10 by addition of aqueous $Na_2CO_3$ dropwise. The aqueous solution was extracted with ethyl acetate (3 x 30 mL), and the extracts were dried ($Na_2SO_4$) and concentrated. Purification of the residue by silica gel chromatography (hexane:ethyl acetate) gave **diol SI6** (459 mg, 81%) as a colorless oil that solidified, and could be further purified by recrystallization from ethyl acetate. $^{1}$H NMR (400 MHz, $CDCl_3$): $\delta$ 7.23-7.12 (m, 5H), 4.11 (s, 2H), 3.32 (s, 2H), 3.19 (bs, 2H), 2.69-2.65 (m, 2H), 2.02-2.00 (m, 2H). 1.94 (d, *J* = 8Hz, 2H), 1.90-1.52 (m, 1H), 1.07 (d, *J* = 8 Hz, 1H); $^{13}$C NMR (100 MHz, $CDCl_3$): $\delta$ 138.8, 128.6, 128.1, 126.9, 75.2, 62.4, 57.1, 43.6, 32.0; $R_f$: 0.41 (50% ethyl acetate in hexane); MS (ESI): calculated for $[C_{14}H_{20}NO_2]^+$: 234.1494, found $[M+H]^+$: 234.1490.

***N*-Benzyl isovarenicline SI7** (via Pfitzner–Moffatt oxidation).

A solution of N-benzyl 3-azabicyclo[3.2.1]octane-6,7-diol **SI6**[4] (250 mg, 1.07 mmol) in DMSO (2 mL) was added to a mixture of DCC (1.77 g, 8.57 mmol) in DMSO (10 mL) followed by dichloroacetic acid (221 mg, 140 μL, 1.72 mmol). The reaction mixture was stirred at rt for 20h, filtered through celite and the solids were washed with EtOAc (50 mL). EtOAc was removed under reduced pressure and to the resulting orange solution was added 1,2-phenylenediamine (116 mg, 1.07 mmol). The mixture was stirred at rt for 18h, water (150 mL) was then added and the product was extracted with EtOAc (3 x 50mL). The extracts were washed with water (150 mL) and brine (150 mL), dried ($Na_2SO_4$) and after removal of solvents, the residue was purified by chromatography (hexane: EtOAc 9:1 → 7:3) to give **N-benzyl isovarenicline SI7** (224 mg, 70%) as a pale orange solid. $^{1}$H NMR (400 MHz, $CDCl_3$): $\delta$ 7.99-7.95 (m, 2 H), 7.63-7.59 (m, 2 H), 7.04-6.89 (m, 3 H). 6.72-6.68 (m, 2 H), 3.38 (s, 2 H), 3.26 (t, 2 H, = 8 Hz), 3.15- 3.11 (m, 2 H), 2.61 (d, 2 H, = 8 Hz), 2.34 (m, 1 H), 1.92 (d, 1 H, = 12 Hz); $^{13}$C NMR

(100 MHz, $CDCl_3$): δ 163.3, 141.7, 137.4, 128.8, 128.4, 128.3, 128.0, 126.8, 61.5, 57.0, 41.1, 39.6; MS (ESI): calculated for $[C_{20}H_{20}N_3]^+$: 302.1652, found $[M+H]^+$: 302.1650.

[Using **SI6** and analogous Swern oxidation conditions (DMSO/TFAA/DCM at -78 °C), we isolated **SI7** in 44% yield.]

##### ***N*-Boc isovarenicline SI8**

Chloroethyl chloroformate (251 μL, 2.32 mmol) was added to a solution of N-benzyl isovarenicline **SI7** (100 mg, 0.332 mmol) in 1,2-dichloroethane (7 mL) and the mixture was stirred at 80°C for 20 h. After cooling to rt and removal of solvent, the residue was dissolved in MeOH (7 mL) and heated under reflux for 2h.** After cooling, di-*tert*-butyl dicarbonate (87 mg, 0.4 mmol) was added and the mixture was stirred at rt for 18h. After concentration, purification by chromatography (hexane:EtOAc 7:3 → 5:5) gave **N-Boc isovarenicline SI8** (89 mg, 86%) as a colorless solid. $^1$H NMR (400 MHz, $CDCl_3$; broadening due to amide resonance was observed): δ 7.99-7.96 (m, 2 H), 7.65-7.60 (m, 2 H), 4.23-4.10 (m 2 H), 3.40-3.25 (m, 4 H), 2.49-2.43 (m, 1 H), 2.05 (d, = 12 Hz, 1 H), 1.14 (s, 9 H); $^{13}$C NMR (100 MHz, $CDCl_3$): δ 161.4, 155.6, 142.2, 129.1, 80.1, 49.3, 48.3, 40.0, 38.1, 28.2; MS (ESI): calculated for $[C_{18}H_{21}N_3O_2Na]^+$: 334.1527, found $[M+Na]^+$: 334.1526.

** At this point, the solvent can be evaporated, and the residue was filtered through a plug of silica (EtOAc as eluent) to provide isovarenicline **5** (as the free base) that was judged (by TLC and $^1$H NMR) to be sufficiently pure to use directly to prepare the tartrate salt (see below). The advantage of the N-Boc intermediate **SI8** is its ease of purification.

The structure of *N*-Boc isovarenicline **SI8** was confirmed by X-ray crystallographic analysis (Figure S1) and the CIF is included as part of the Supporting Information package.

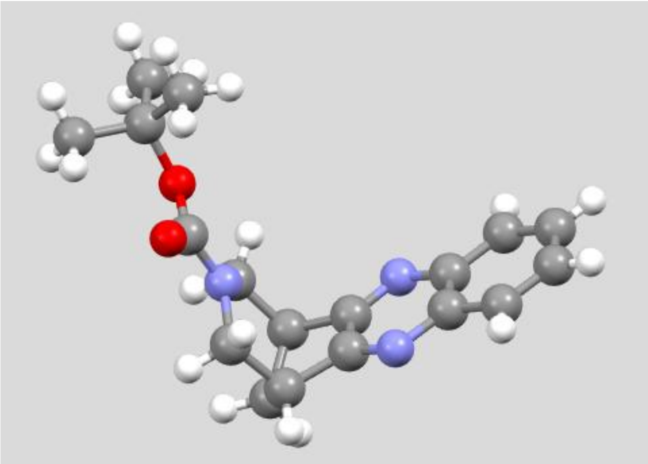

**Figure S1-** X-ray structure of *N*-Boc isovarenicline **SI8**. The grey, white, red and blue spheres correspond to carbon, hydrogen, oxygen and nitrogen atoms, respectively.

**Isovarenicline HCl salt 5.**

To a solution of *N*-Boc isovarenicline **SI8** (180 mg, 0.58 mmol) in MeOH (5 mL) was added HCl (4M in dioxane, 1 mL). After 8 h, the solvents were removed and the solid was triturated with cold EtOAc to give **isovarenicline HCl 5** (140 mg, quantitative) as a light tan solid $^{1}$H NMR (600 MHz,$D_2O$): δ 7.98 (m, 2 H, H1), 7.80 (m, 2 H, H2), 3.62 (br d, 2 H, *J* = 12 Hz, H7a), 3.58 (m, 2 H, H6), 3.44 (d, 2 H, *J* = 12 Hz, H7b), 2.60 (m, 1 H, H9a), 2.33 (d, 1 H, *J* = 12 Hz, H9b); $^{13}$C NMR (100 MHz, $D_2O$): δ 158.7, 141.5, 130.8, 128.1, 46.3, 38.2, 36.8. MS (ESI): calculated for $[C_{13}H_{14}N_3]^+$: 212.1188, found $[M]^+$: 212.1190.

**Alternative salt preparation**

**Isovarenicline <u>tartrate</u> salt**

Isovarenicline (from N-Bn isovarenicline **SI7** - see** above; 134 mg, 0.64 mmol) in EtOH:acetone (10:3, 1.5 mL) was added to a solution of (*L*)-(+)-tartaric acid (102 mg, 0.68 mmol) in EtOH:acetone (10:3, 1 mL). After 15 min, the mixture was cooled to 0$^{o}$C and the solid was isolated by filtration, washed with a small quantity of cold 10:3 EtOH:acetone and air-dried to give **isovarenicline tartrate** (174 mg, 76%) as an off-white powder. Spectroscopic properties were identical to the HCl salt with the exception of the signals associated with tartrate, but full details are provided below. $^{1}$H NMR (600 MHz, $D_2O$): δ 7.96 (m, 2 H, H1), 7.80

(m, 2 H, H2), 4.39 (s, 2 H, tartrate), 3.62 (br d, 2 H, *J* = 12 Hz, H7), 3.58 (m, 2 H, H6), 3.44 (d, 2 H, *J* = 12 Hz, H7), 2.60 (m, 1 H, H9), 2.33 (d, 1 H, *J* = 12 Hz, H9); $^{13}$C NMR (100 MHz, $D_2O$): δ 176.3 (C=O, tartrate), 158.7, 141.5, 130.8, 128.1 , 72.8, 46.3, 38.2, 36.8.

#### 3. $N_2$ Varenicline 6

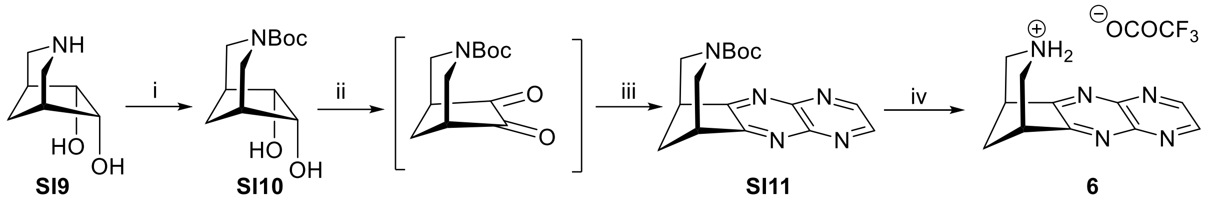


**SI Scheme 3.** *Reagents*: i, $Boc_2O$, $Na_2CO_3$, THF/water, rt, 16h (90%); ii, TFAA, DMSO, $CH_2Cl_2$, then $Et_3N$; iii, 2,3-diaminopyrazine, MeOH, 65 °C, 16h (38% over 2 steps); iv, 5% TFA in MeOH (69%).

##### *tert*-Butyl-6,7-dihydroxy-3-azabicyclo[3.2.1]octane-3-carboxylate SI10

To a solution of commercially available diol **SI9**·(as HCl salt; also available by hydrogenolysis of **SI6**) (0.89 g, 5.00 mmol) in THF (16 mL) was added di-tert-butyl dicarbonate (1.31 g, 6.00 mmol), followed by a solution of $Na_2CO_3$ (1.11 g, 10.5 mmol) in water (8 mL). The mixture was stirred 16 h at rt, then diluted with EtOAc (50 mL) and brine (20 mL). The phases were separated, and the aqueous phase was extracted with EtOAc (2 x 50 mL). The combined EtOAc extracts were washed with brine (50 mL), dried ($Na_2SO_4$), concentrated and the residue was purified by silica gel chromatography (pentane/EtOAc 20:80 to 100% EtOAc) to give **SI10** (1.10 g, 90%) as a colorless solid. $^1$H NMR (500 MHz, $CDCl_3$) δ 4.18 – 4.05 (m, 4H), 2.83 (d, *J* = 13.0 Hz, 2H), 2.26 (m, 2H), 1.58 – 1.49 (m, 2H), 1.45 (s, 9H). $^{13}$C NMR (126 MHz, $CDCl_3$) δ 156.2, 80.1, 70.3, 37.7, 29.4, 29.4, 28.5. HRMS (ESI) calculated for $C_{12}H_{21}NO_4$, $[M+H]^+$: 244.1543, found: 244.1546.

##### *N*-Boc $N_2$ Varenicline SI11

To a solution of DMSO (0.15 mL) in $CH_2Cl_2$ (1.2 mL) at -78 °C, was slowly added trifluoroacetic anhydride (0.21 mL, 1.5 mmol), followed by dropwise addition of a solution of diol **SI10** (0.12

g, 0.5 mmol) in $CH_2Cl_2$ (0.4 mL). The reaction mixture was stirred for 1 h at -78 °C, then triethylamine (0.35 mL, 2.5 mmol) was added. The resulting mixture was stirred 1 h at rt, then water (5 mL) was added and the mixture was extracted with $CH_2Cl_2$ (2 x 10 mL). The combined extracts were dried ($Na_2SO_4$) and the crude 1,2-diketone was used immediately in the next step.

To a solution of crude diketone (as above) in MeOH (1.0 mL) was added 2,3-diaminopyrazine[5] (55 mg, 0.5 mmol) The resulting heterogeneous mixture was then stirred at 65 °C in a closed Schenck tube for 16 h affording a tan-colored homogeneous solution. The mixture was concentrated and purification by chromatography (BIOTAGE, Sfar Silica HC D 10 g column, $CH_2Cl_2$/MeOH gradient 2 to 20%) and recrystallization from diethyl ether, afforded ***N*-Boc $N_2$ varenicline SI11** (60 mg, 38% over 2 steps) as a pale-yellow solid. $^1$H NMR (400 MHz, $CDCl_3$) δ 9.04 (s, 2H), 4.36 – 4.19 (m, 2H), 3.66 – 3.48 (m, 2H), 3.49 – 3.35 (m, 2H), 2.61 (m, 1H), 2.20 (d, *J* = 11.7 Hz, 1H), 1.23 (s, 9H). $^{13}$C NMR (101 MHz, $CDCl_3$) δ 167.3, 155.5, 147.0, 146.5, 80.5, 49.5, 48.5, 40.2, 38.0, 28.2. HRMS (ESI) calculated for $C_{16}H_{19}N_5O_2$, $[M+Na]^+$: 336.1431, found: 336.1428.

**$N_2$ Varenicline trifluoroacetate 6**

*N*-Boc $N_2$ varenicline **SI11** (40 mg, 0.13 mmol) was dissolved in a solution of trifluoroacetic acid (5% v/v in dichloromethane, 4 mL) and allowed to stand for 16 h at rt. The solvent was removed and toluene was added to facilitate removal of residual TFA under vacuum. The resulting brown oil was triturated with acetone affording a crystalline yellow solid, from which the solvent was decanted, and the solids were washed again with acetone, acetone decanted and the solid was dried under vacuum to give **$N_2$ varenicline trifluoroacetate 6** (30 mg, 69%) as a yellow crystalline solid. . $^1$H NMR (500 MHz, DMSO) δ 9.22 (s, 1H), 9.19 (s, 2H), 8.61 (s, 1H), 3.65 (dd, *J* = 4.9, 2.3 Hz, 2H), 3.62 (d, *J* = 11.0 Hz, 2H), 3.42 (d, *J* = 11.0 Hz, 2H), 2.63 (m, 1H), 2.42 (d, *J* = 11.7 Hz, 1H). $^{13}$C NMR (126 MHz, DMSO) δ 164.4, 158.3 (q, *J* = 35.8 Hz), 147.7, 146.4, 115.8 (q, *J* = 292.4 Hz), 45.9, 38.3, 36.5. $^{19}$F NMR (376 MHz, DMSO-*$D_6$*) δ -74.57. HRMS (ESI) calculated for $C_{11}H_{12}N_5$, $[M+H]^+$: 214.1087, found: 214.1089.

We also evaluated an alternative approach to $N_2$ varenicline **6** based on use of the N-benzyl intermediate by analogy to the chemistry used for **4** and **5**.

**N-Benzyl $N_2$ varenicline SI12**

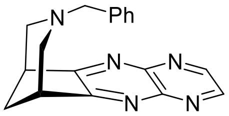

**SI12**

To mixture of DCC (1.77 g, 8.57 mmol) in DMSO (10 mL) a solution of benzyl-3-azabicyclo[3.2.1]octane-6,7-diol **SI6** (250 mg, 1.07 mmol) in DMSO (2 mL) was added followed by dichloroacetic acid (221 mg, 140 μL, 1.72 mmol). The reaction mixture was stirred at rt for 20h, then filtered through celite which was washed with ethyl acetate (50 mL). EtOAc was removed and to the orange residue was added 2,3-diaminopyrazine (118 mg, 1.072 mmol) and the mixture was stirred at 65 °C for 3h. Water (150 mL) was added, and aqueous phase was extracted with ethyl acetate (3 x 50mL). The combined extracts were washed with water (150 mL) and brine (150 mL), concentrated, and the orange residue was purified by silica chromatography (eluent 1% MeOH → 2.5% MeOH in DCM) to give **N-benzyl $N_2$ Varenicline SI12** (130 mg, 40%) as a pale-yellow solid. $^1H$ NMR (400 MHz, $CDCl_3$): δ 8.94 (s, 2H), 7.05 – 7.00 (m, 3H), 6.71 – 6.69 (m, 2H), 3.42 (bt, *J* = 4 Hz, 2H), 3.38 (s, 2H), 3.23 – 3.19 (m, 2H), 2.68 (d, *J* = 8 Hz, 2H), 2.44 (m, 1H), 2.02 (d, *J* = 12 Hz, 1H); $^{13}C$ NMR (100 MHz, $CDCl_3$): δ 169.3, 146.4, 146.1., 137.1, 128.4, 128.3, 127.1, 61.8, 57.4, 41.3, 39.5; Calculated for $[C_{18}H_{18}N_5]^+$: 304.1562, found $[M+H]^+$: 304.1560.

N-Benzyl $N_2$ varenicline **SI12** was of very limited synthetic utility because under a variety of conditions (essentially those approaches that had worked for debenzylation of **SI2** and **SI7**), we were unable to deprotect successfully **SI12**. We only observed substrate decomposition but $^1H$ NMR analysis of the crude product indicated heteroarene reduction/fragmentation had probably occurred.

#### *(iii)* Supporting figures: $^{1}$H and $^{13}$C NMR spectra of key intermediates and final products.

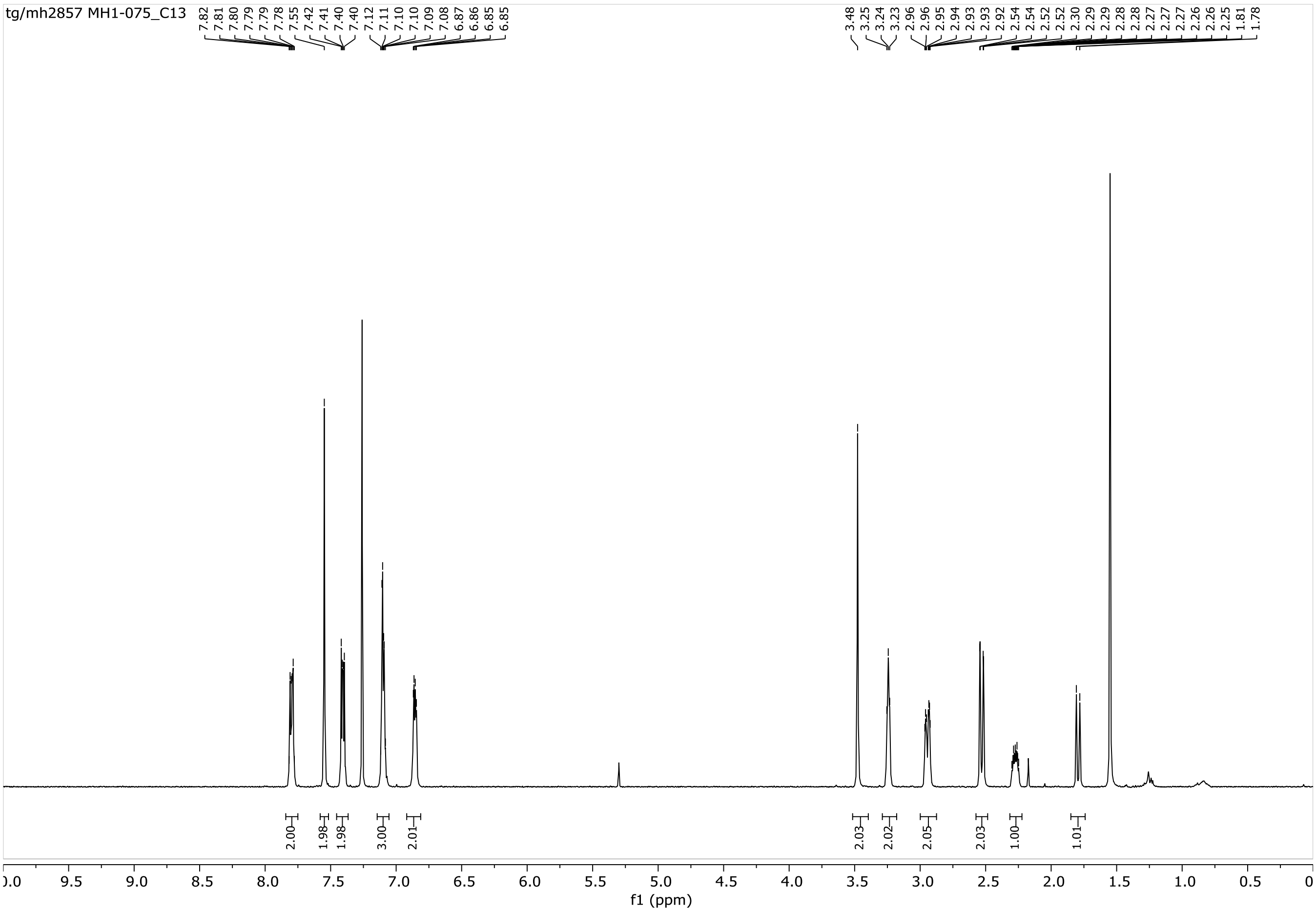


$^{1}$H NMR of *N*-Bn $C_2$ varenicline **SI2**

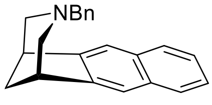


**SI2**

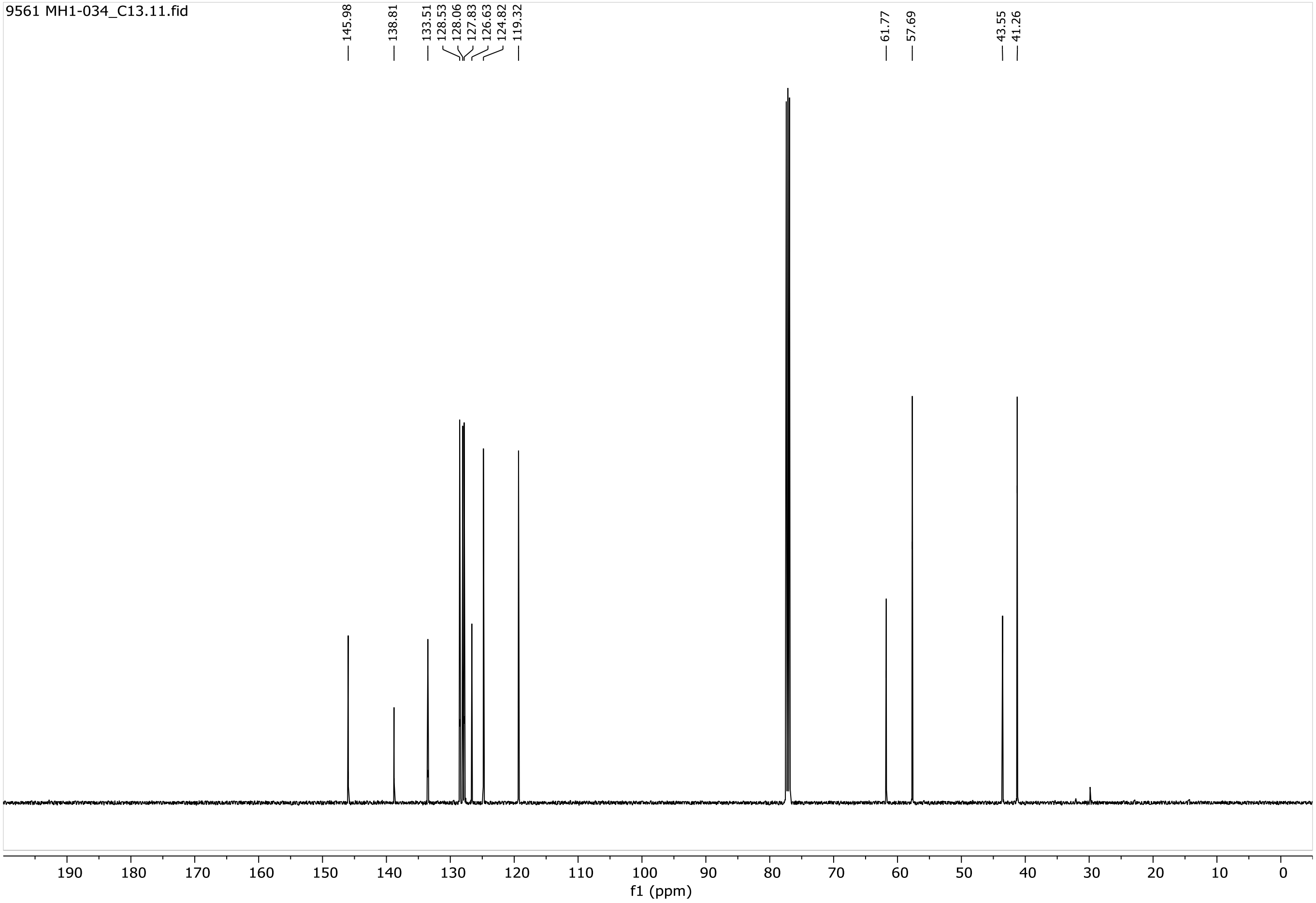


$^{13}C$ NMR of *N*-Bn $C_2$ varenicline **SI2**

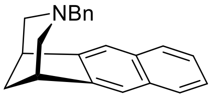


**SI2**

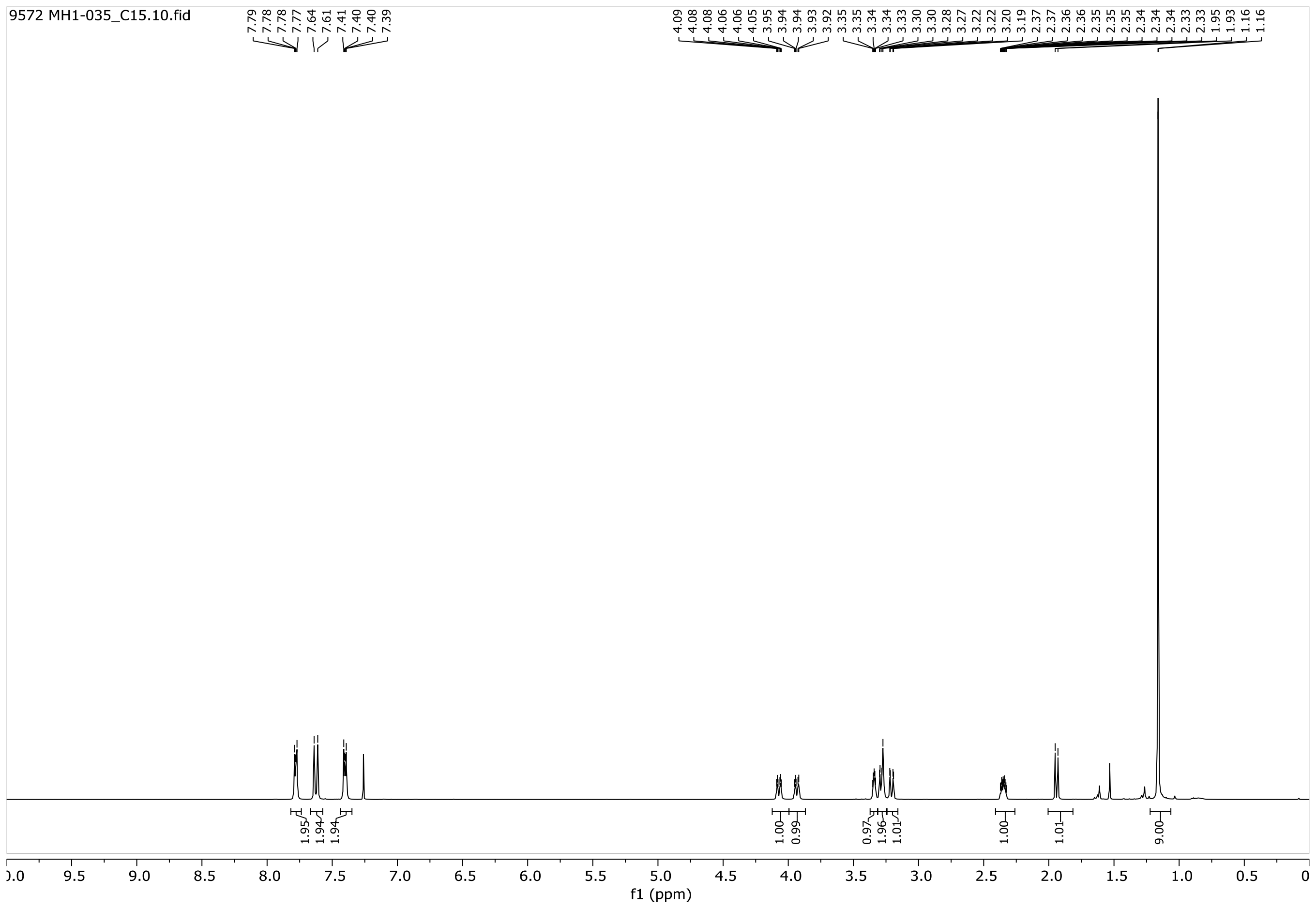


$^1$H NMR of *N*-Boc $C_2$ varenicline **SI3**

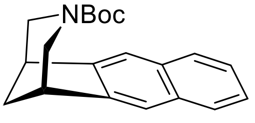


**SI3**

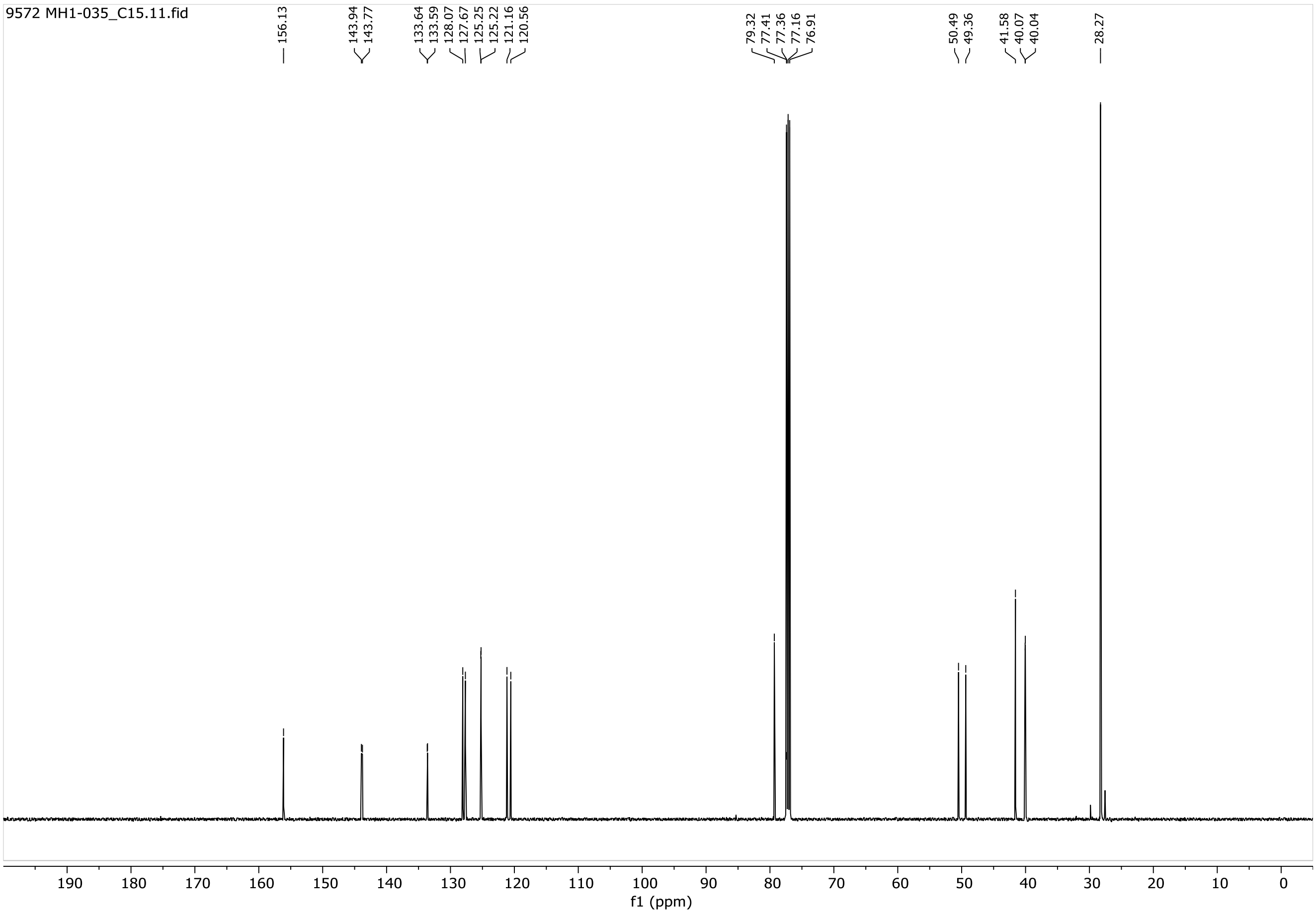


$^{13}$C NMR of *N*-Boc $C_2$ varenicline **SI3**

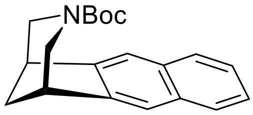


**SI3**

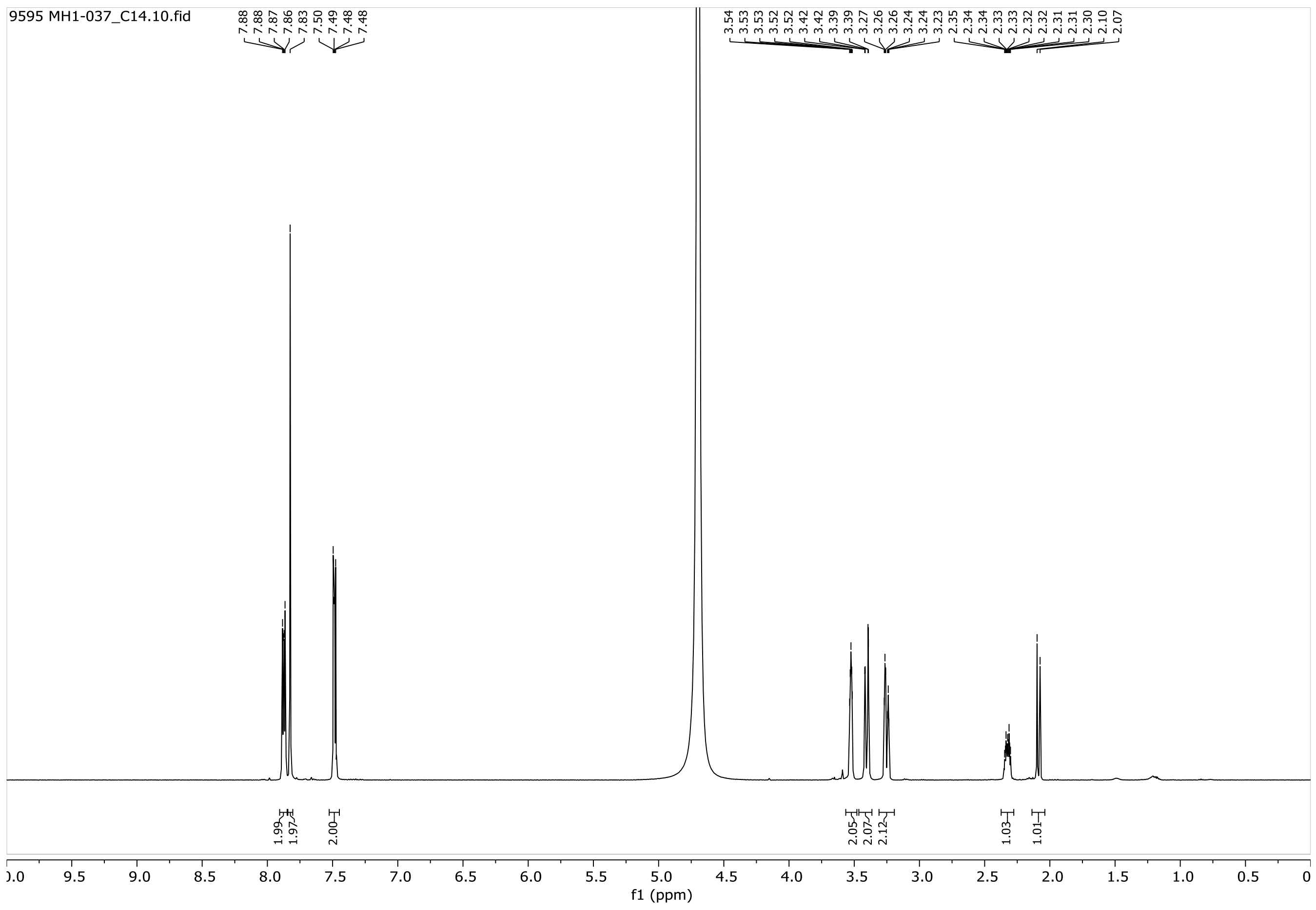

$^1H$ NMR $C_2$ varenicline.HCl **4**

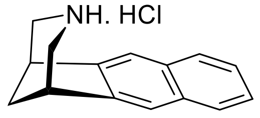

**4**

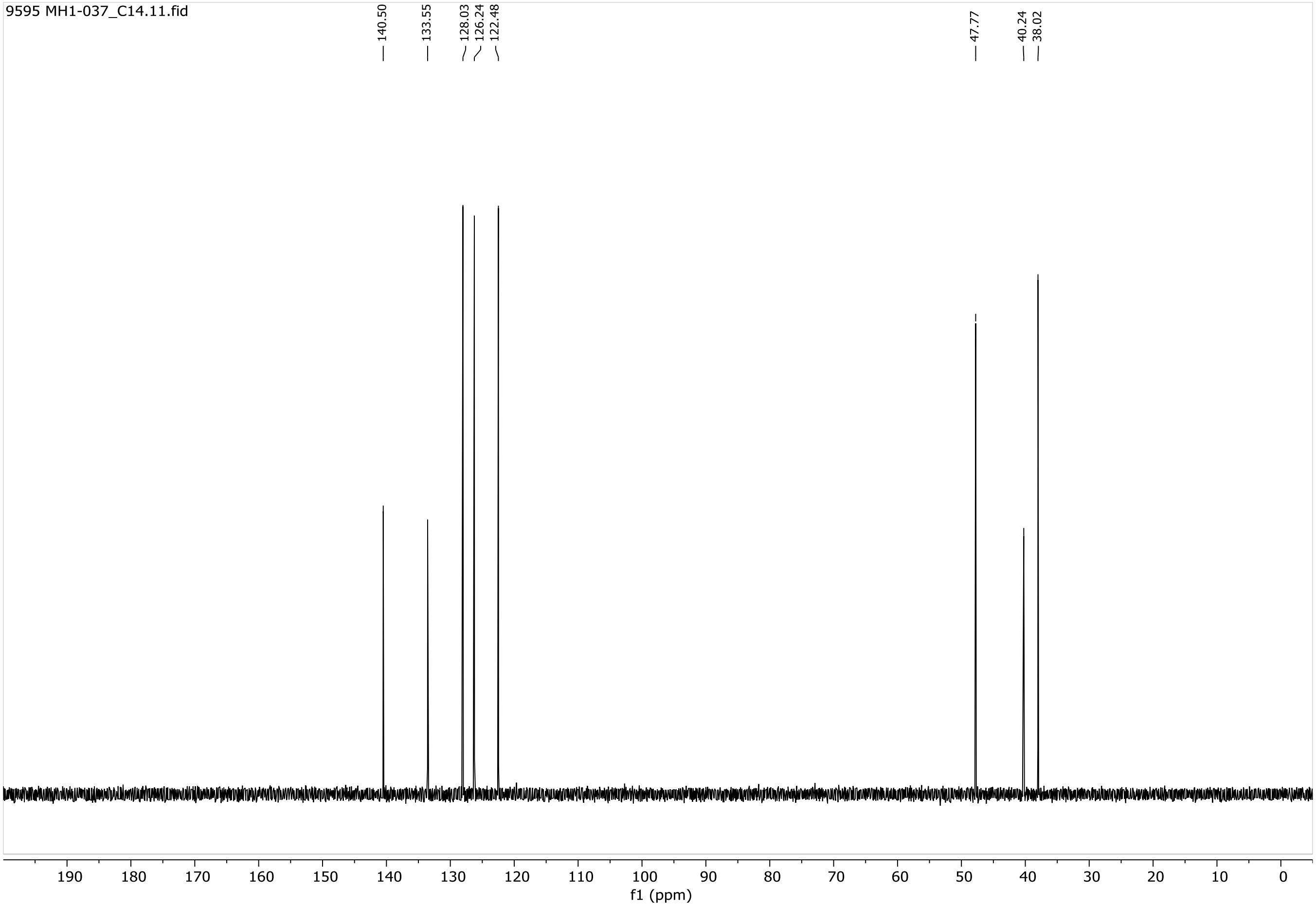


$^{13}C$ NMR $C_2$ varenicline.HCl **4**

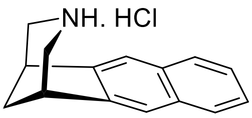


**4**

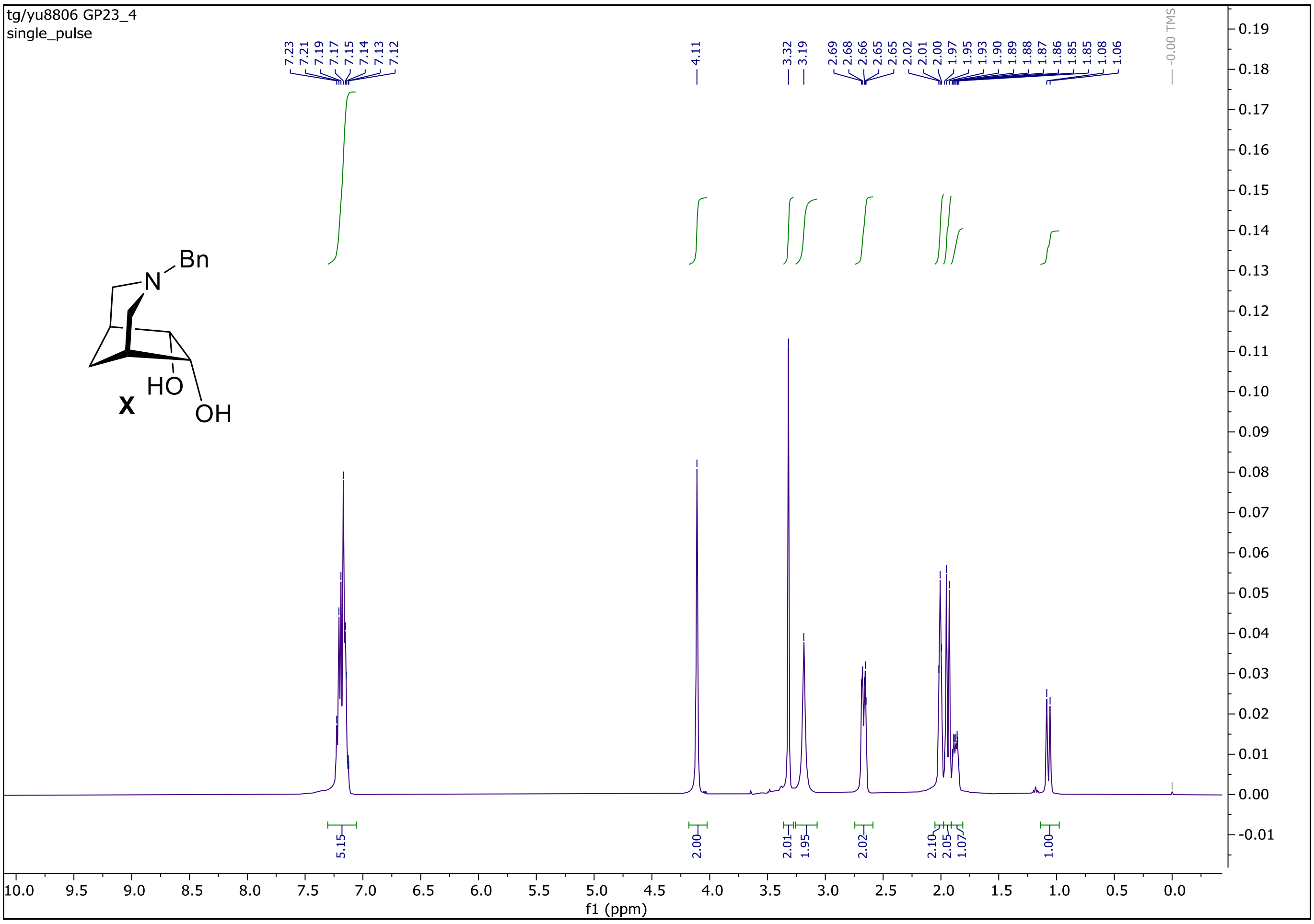


[1]H NMR of *N*-Bn diol **SI6**

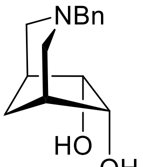


**SI6**

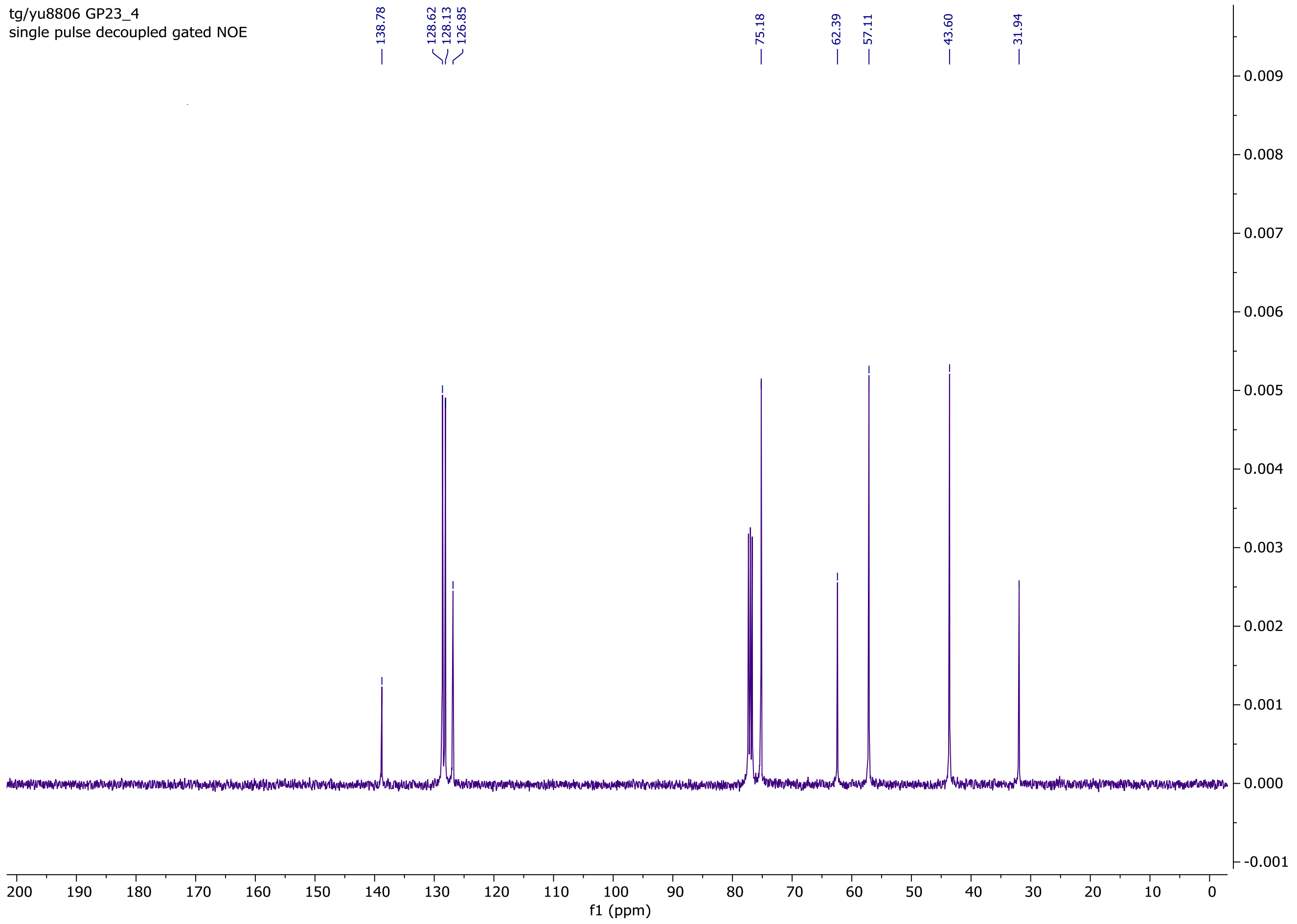


$^{13}$C NMR of *N*-Bn diol **SI6**

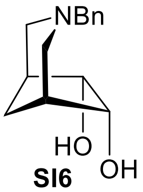


**SI6**

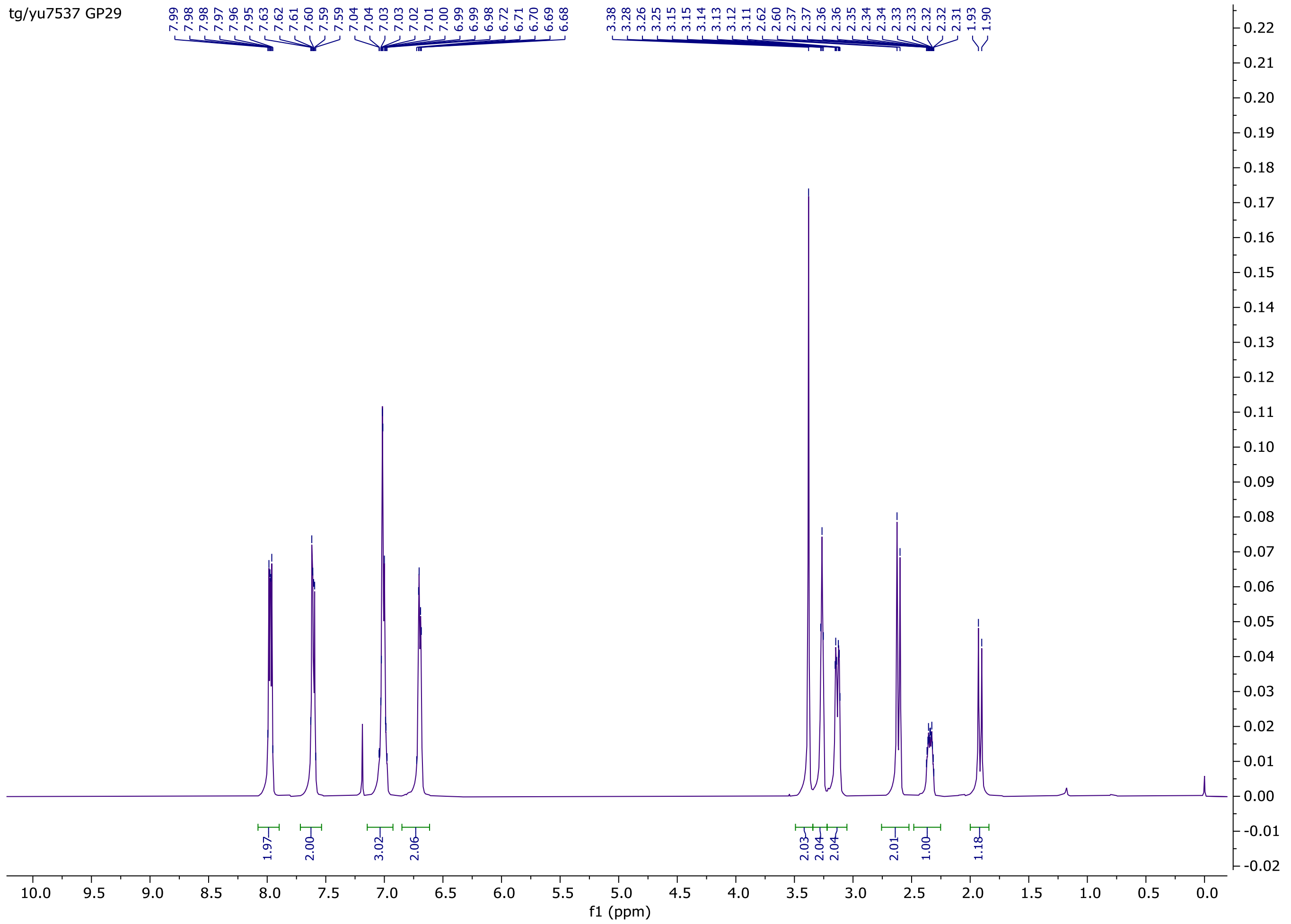



$^1$H NMR of *N*-Bn isovarenicline **SI7**

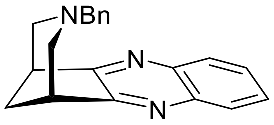

**SI7**

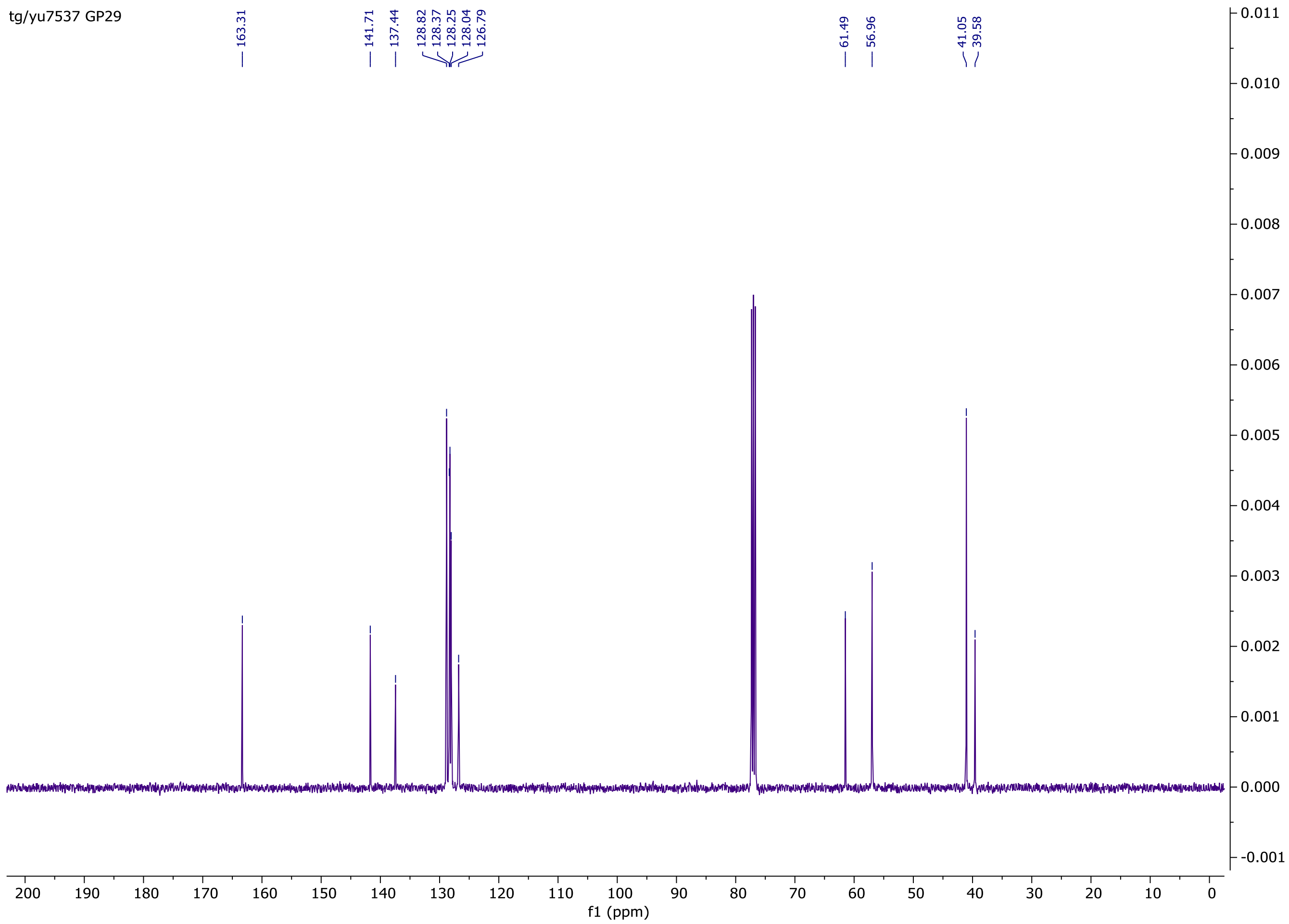


$^{13}$C NMR of *N*-Bn isovarenicline **SI7**

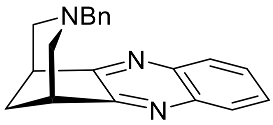


**SI7**

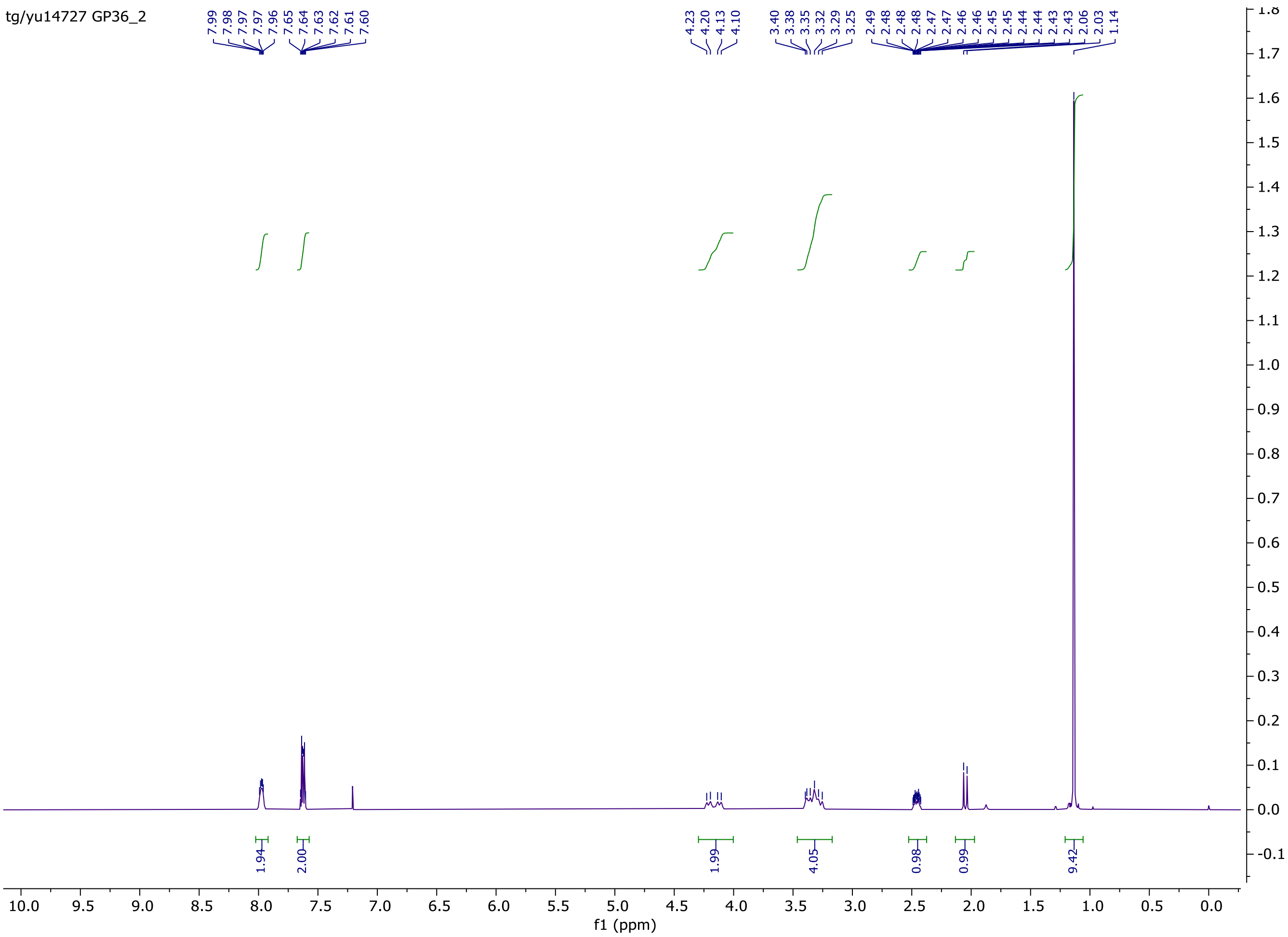


$^1$H NMR of *N*-Boc isovarenicline **SI8**

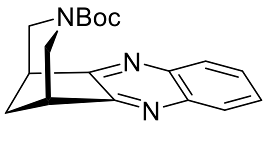


**SI8**

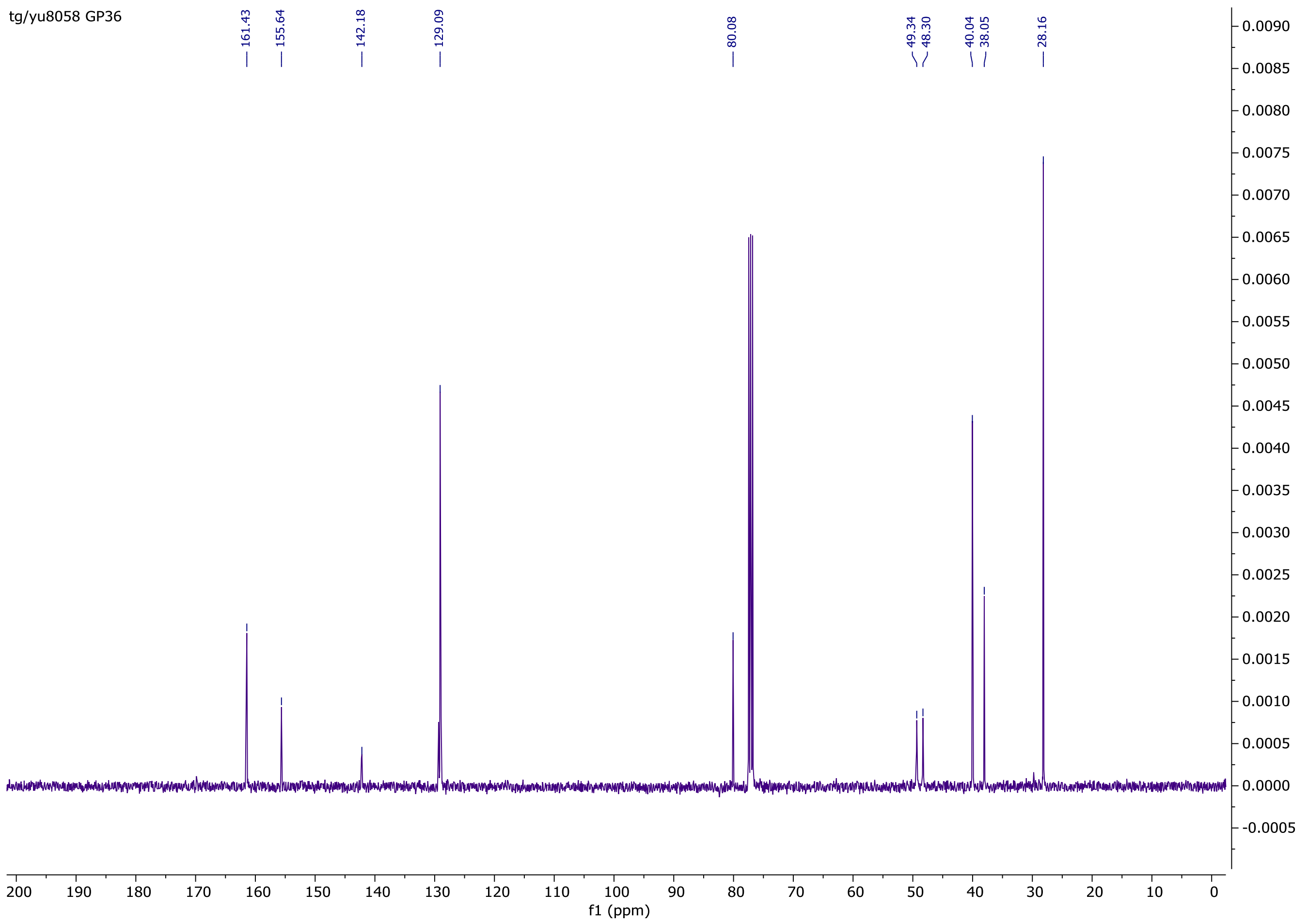


$^{13}$C NMR of *N*-Boc isovarenicline **SI8**

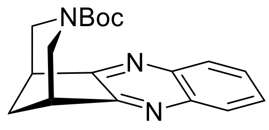


**SI8**

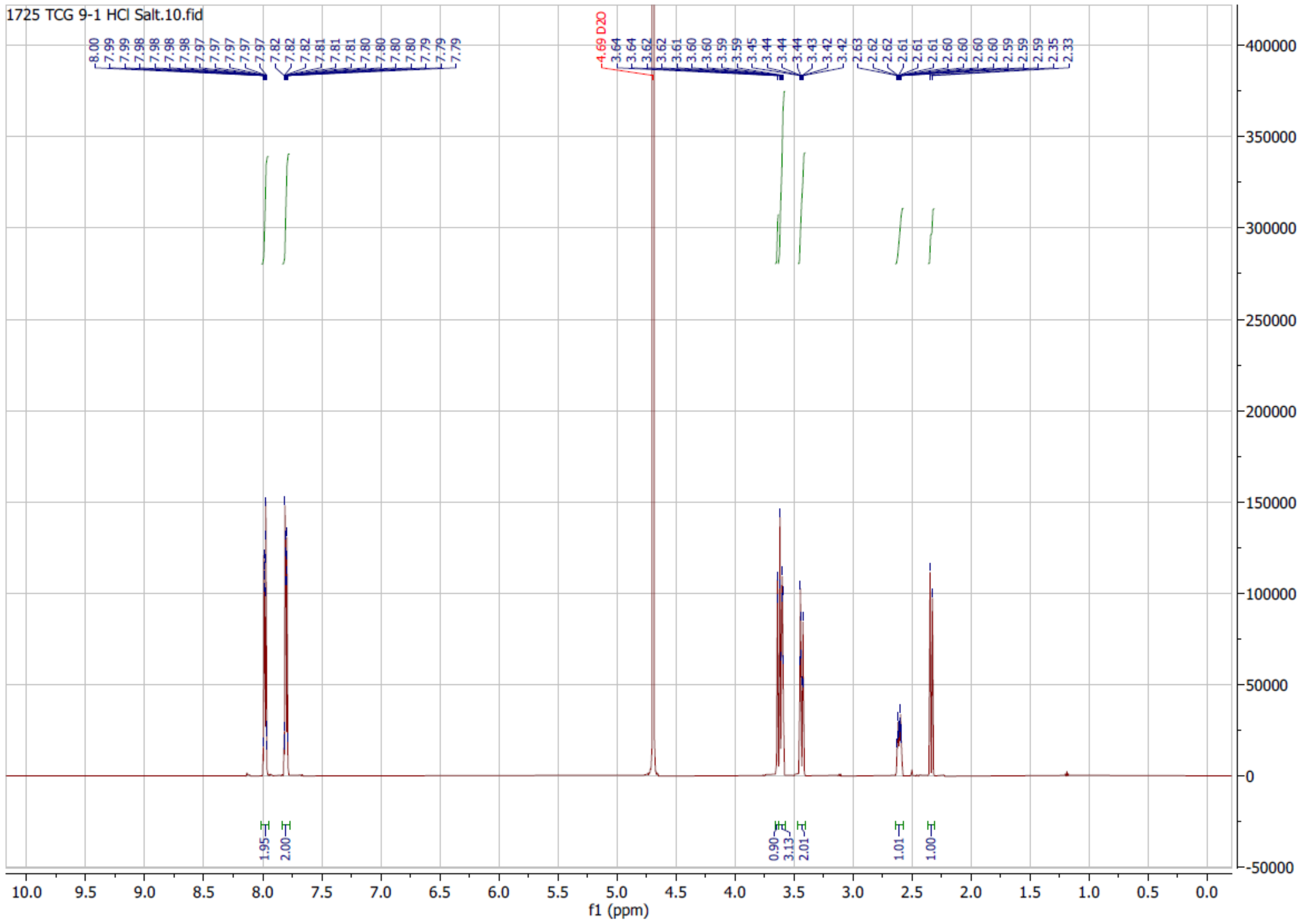
1725 TCG 9-1 HCl Salt.10.fid
4.69 D2O
f1 (ppm)

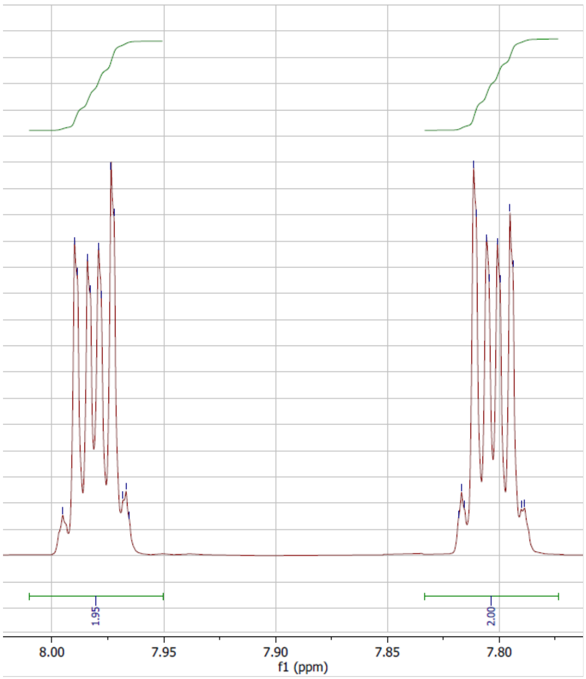
f1 (ppm)

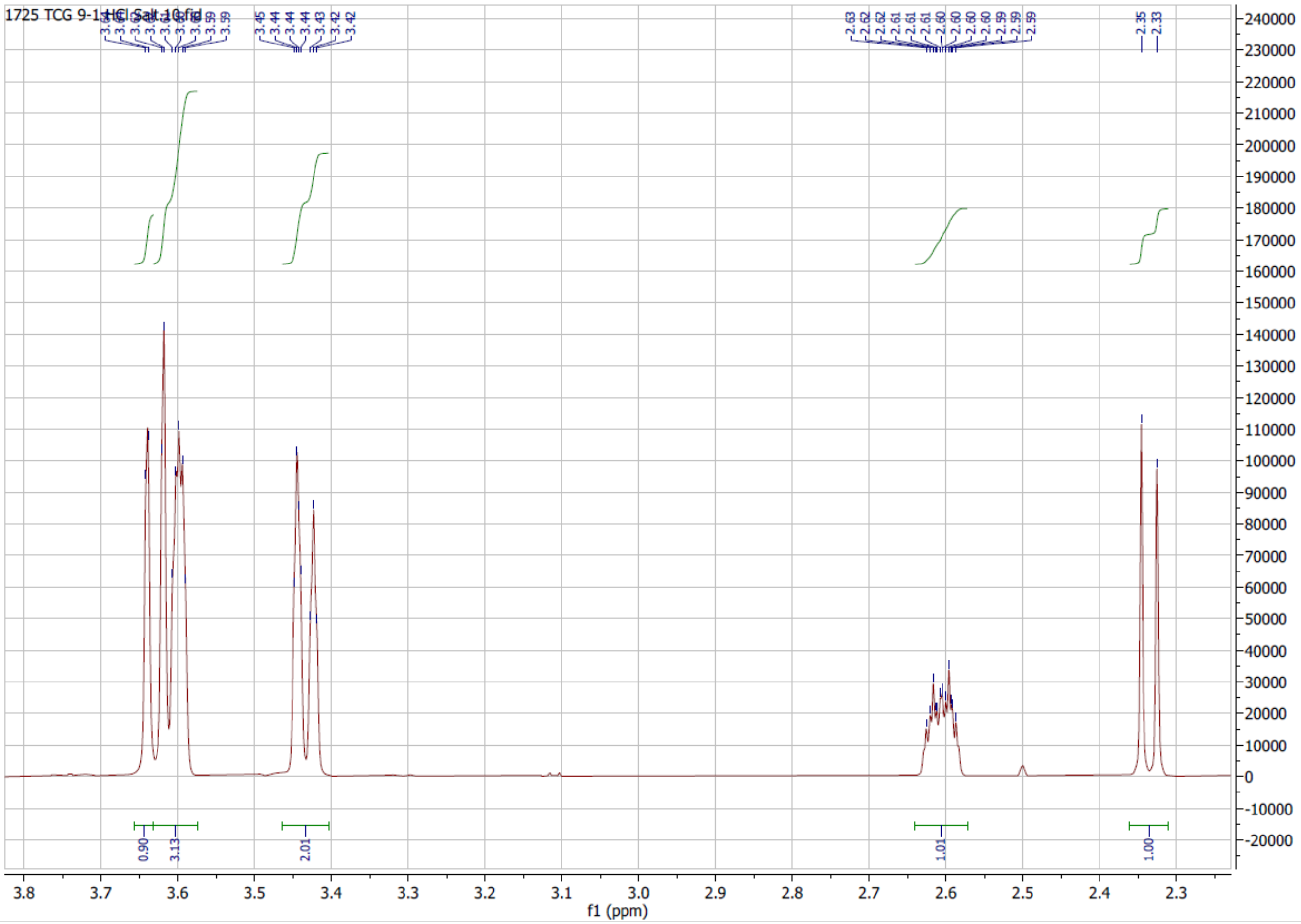


[1]H NMR of isovarenicline HCl **5**

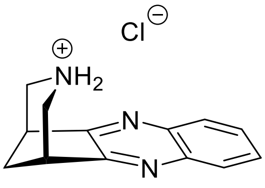

**5**

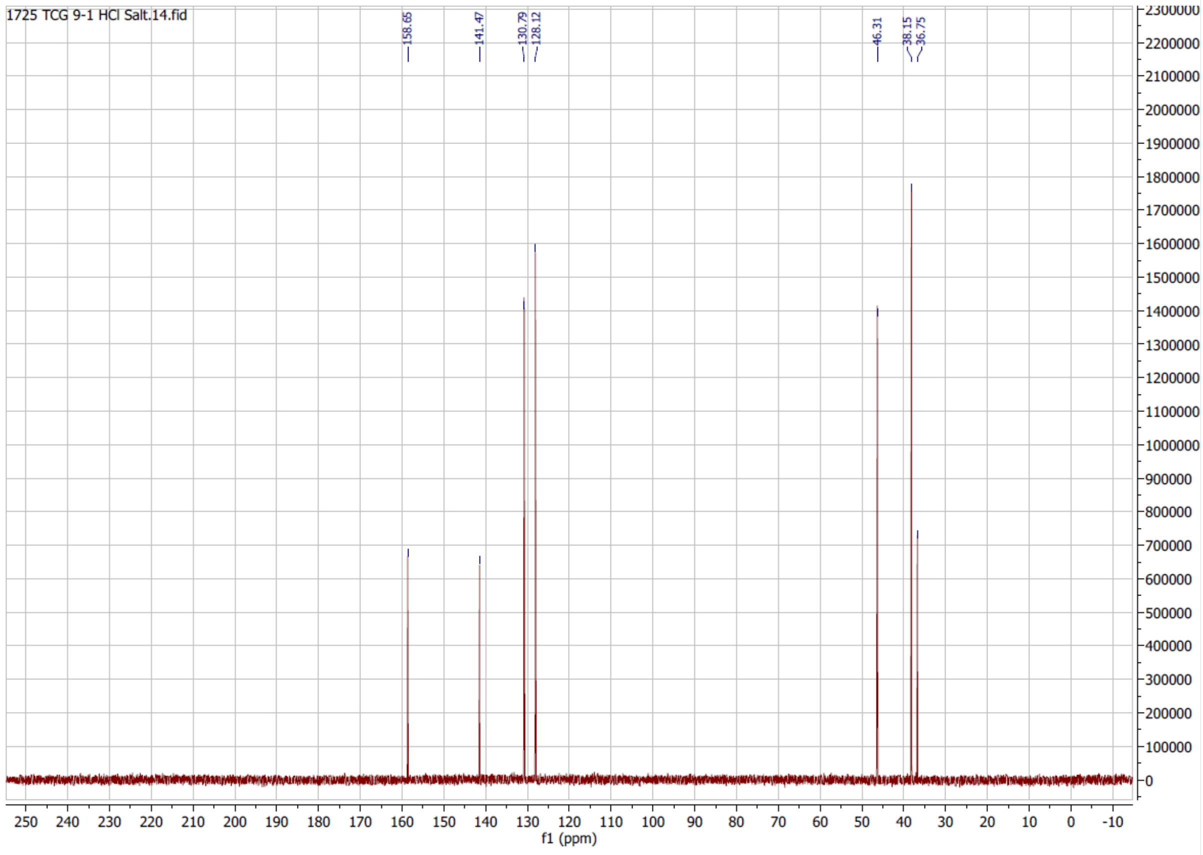


$^{13}$C NMR of isovarenicline HCl **5**

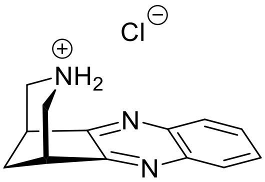


**5**

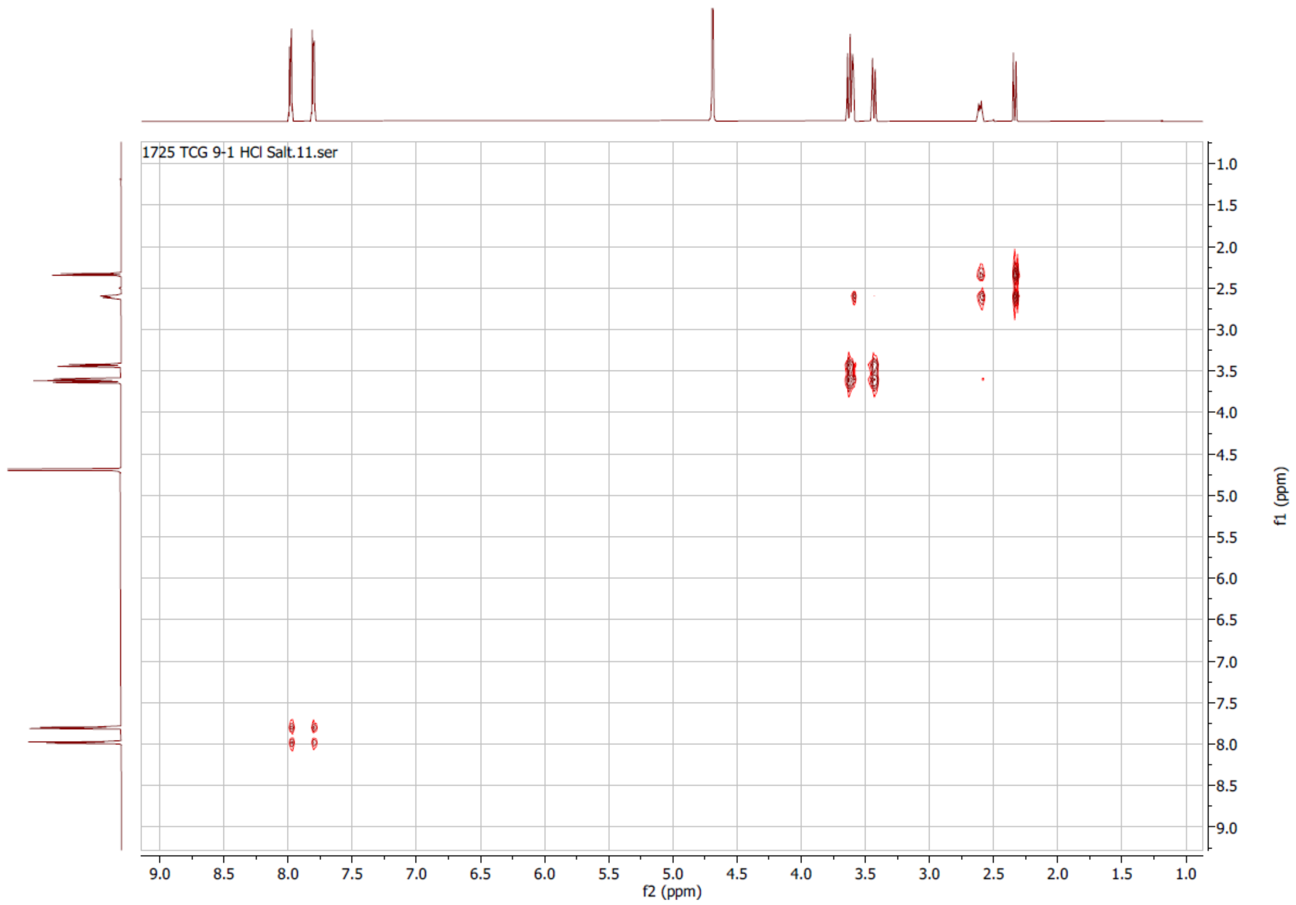
1725 TCG 9-1 HCl Salt.11.ser
f2 (ppm)
f1 (ppm)
9.0
8.5
8.0
7.5
7.0
6.5
6.0
5.5
5.0
4.5
4.0
3.5
3.0
2.5
2.0
1.5
1.0

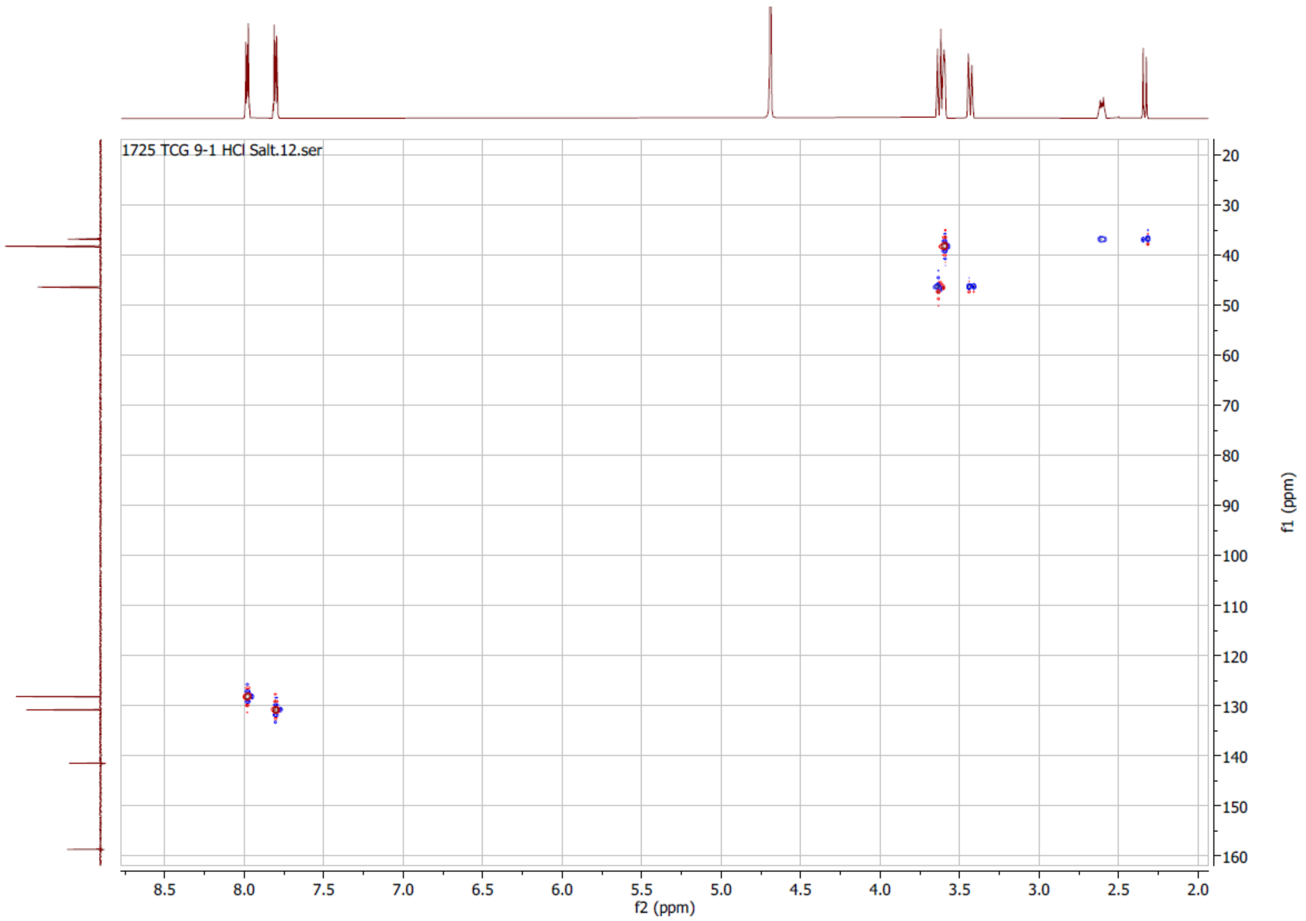

1725 TCG 9-1 HCl Salt.12.ser
8.5
8.0
7.5
7.0
6.5
6.0
5.5
5.0
4.5
4.0
3.5
3.0
2.5
2.0
f2 (ppm)
20
30
40
50
60
70
80
90
100
110
120
130
140
150
160
f1 (ppm)

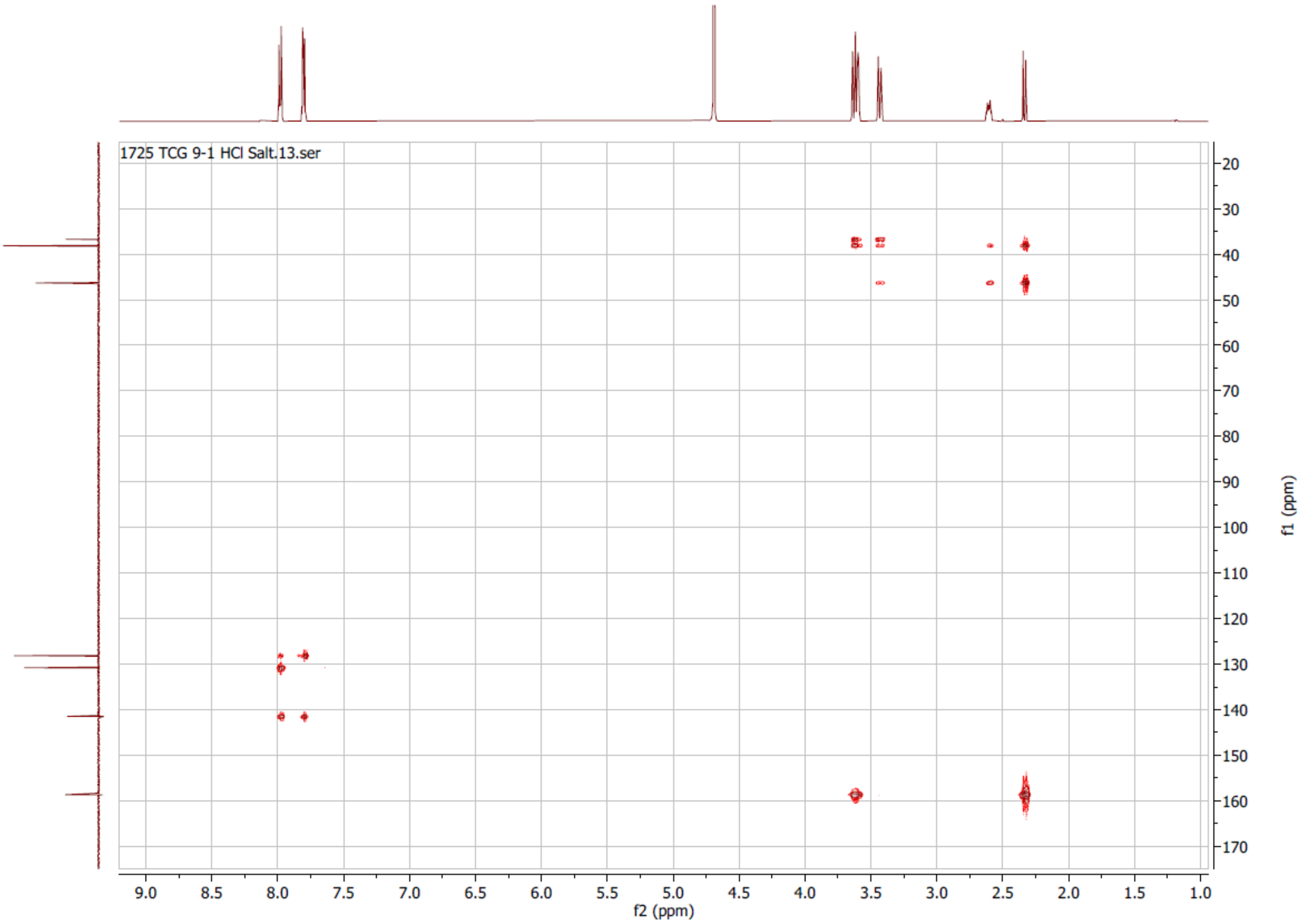


2D COSY. HSQC and HMBC spectra of isovarenicline HCl **5**

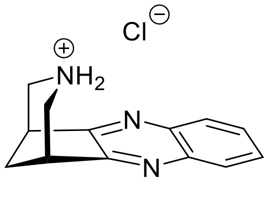


**5**

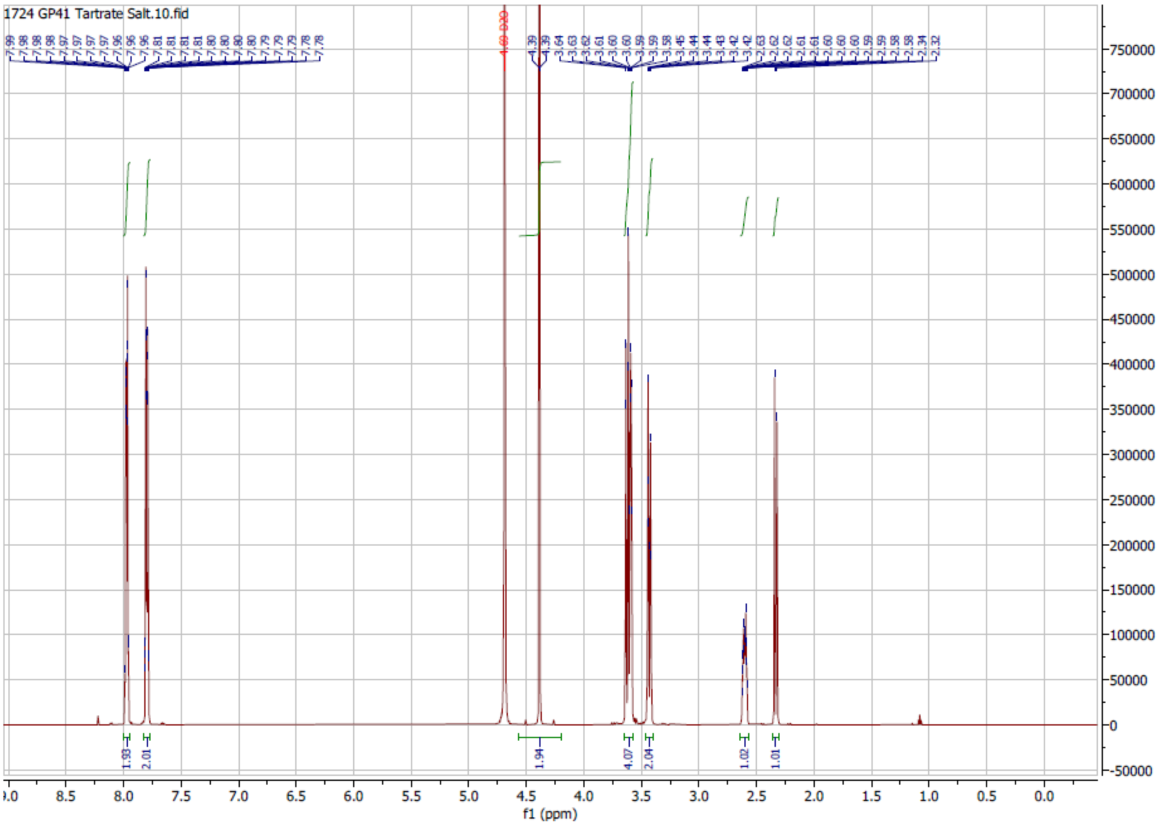


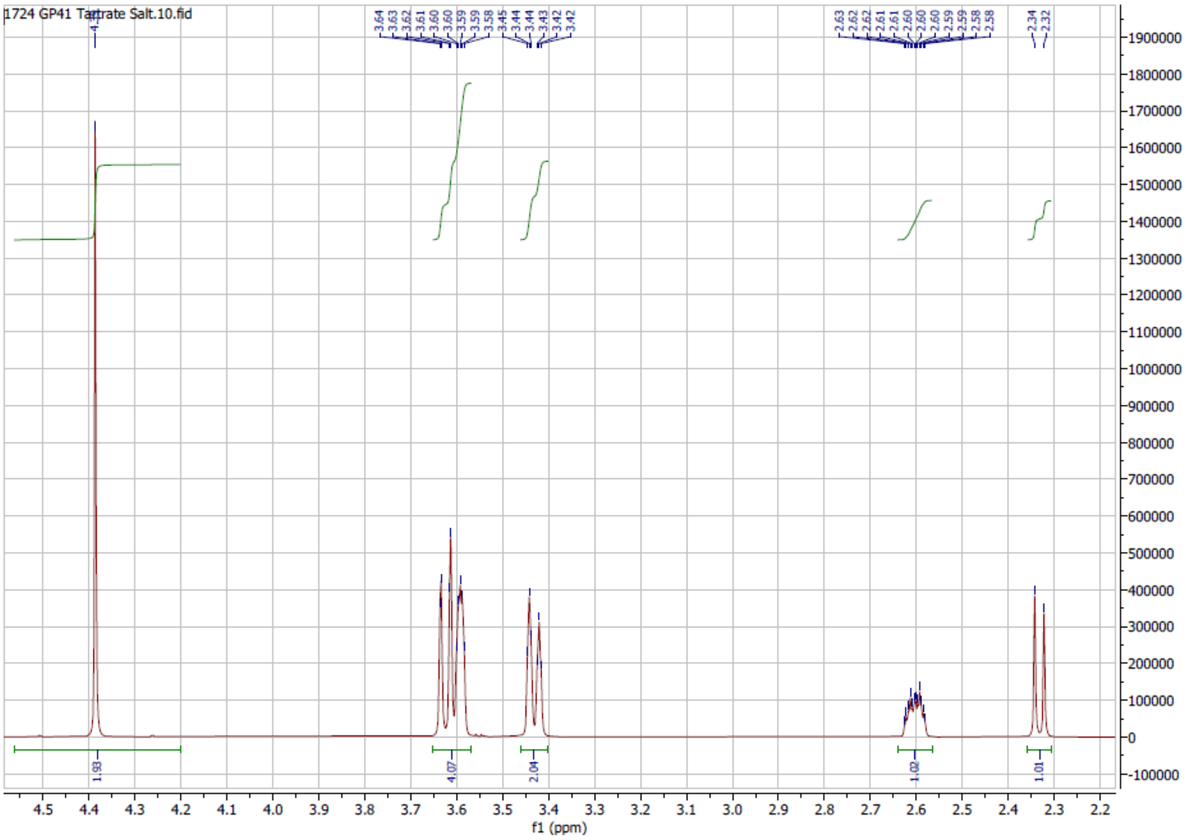


$^1$H NMR of isovarenicline tartrate

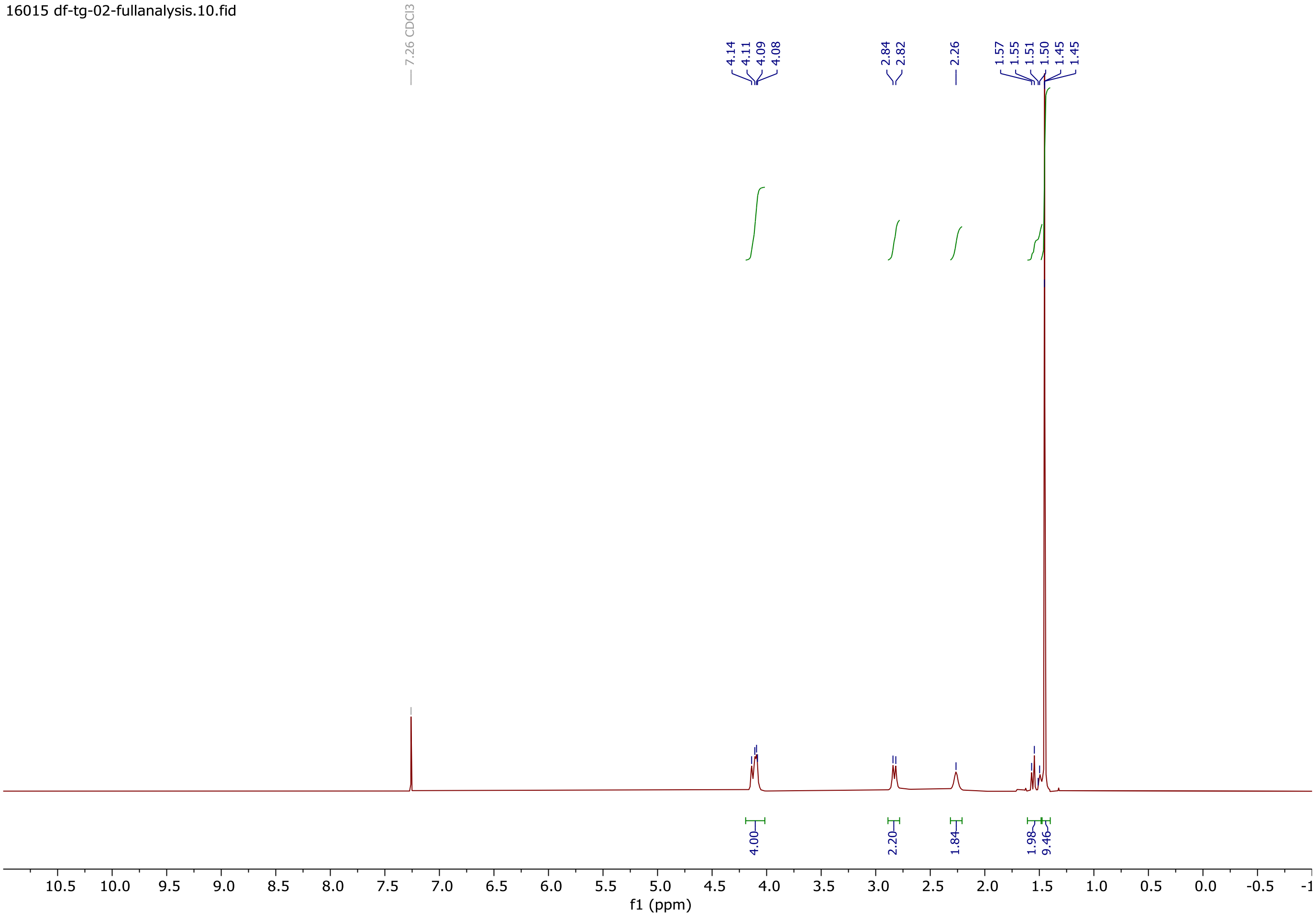


$^{1}$H NMR of N-Boc diol **SI10**

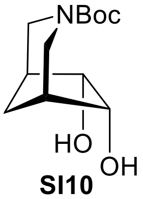


**SI10**

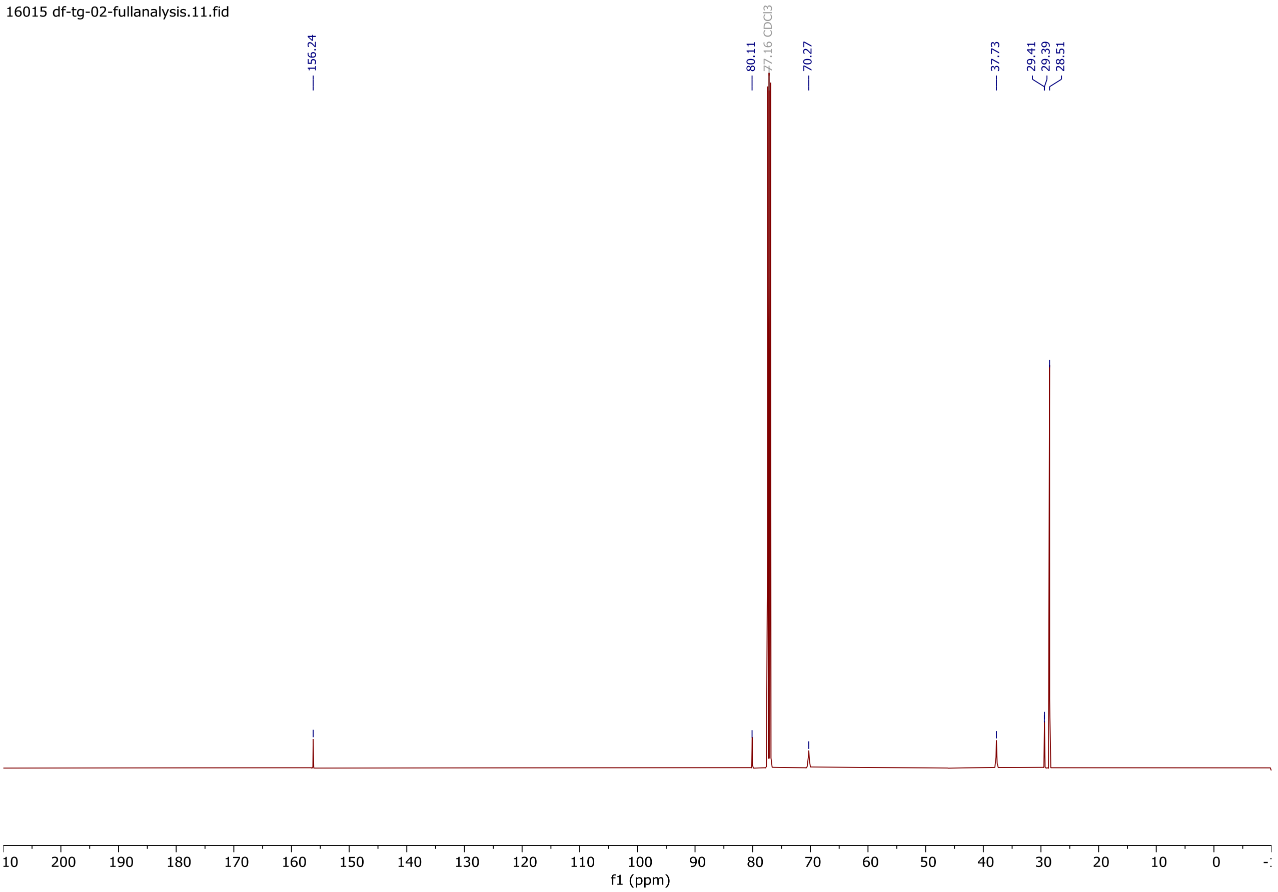


$^{13}C$ NMR of N-Boc diol **SI10**

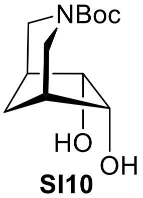


**SI10**

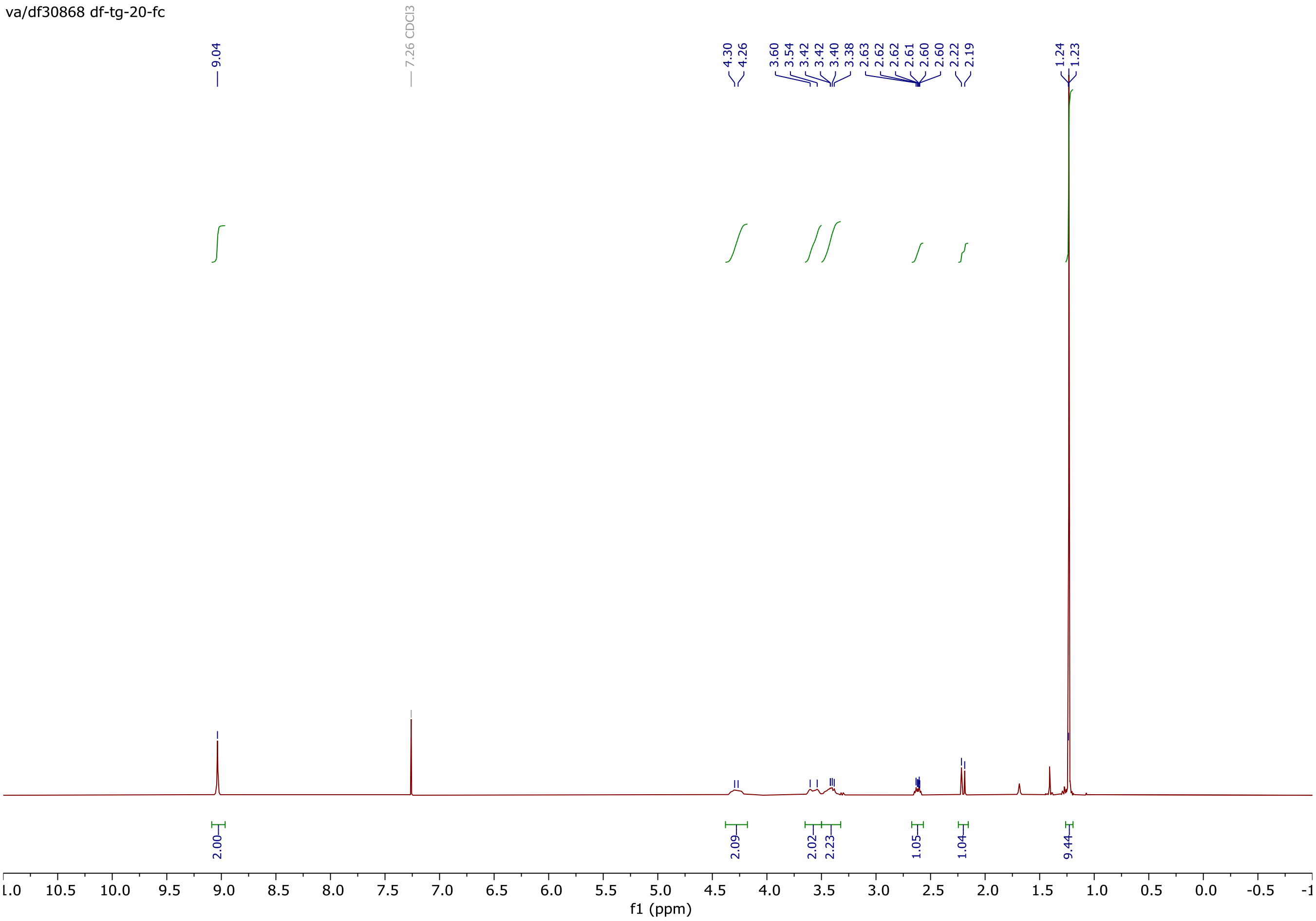


$^{1}$H NMR of N-Boc $N_2$ varenicline **SI11**

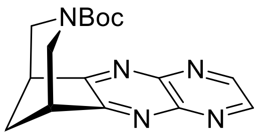


**SI11**

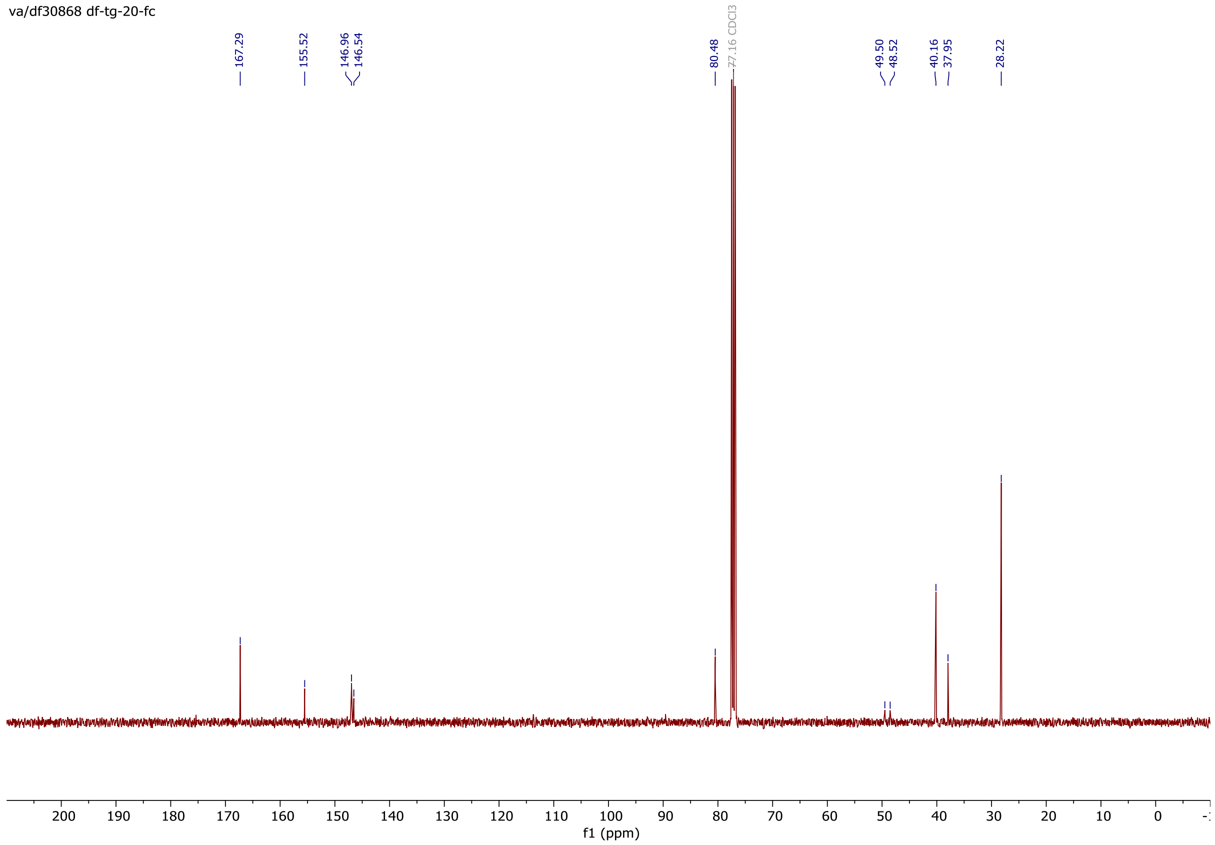


$^{13}C$ NMR of N-Boc $N_2$ varenicline **SI11**

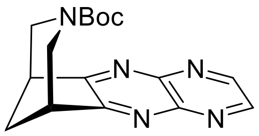


**SI11**

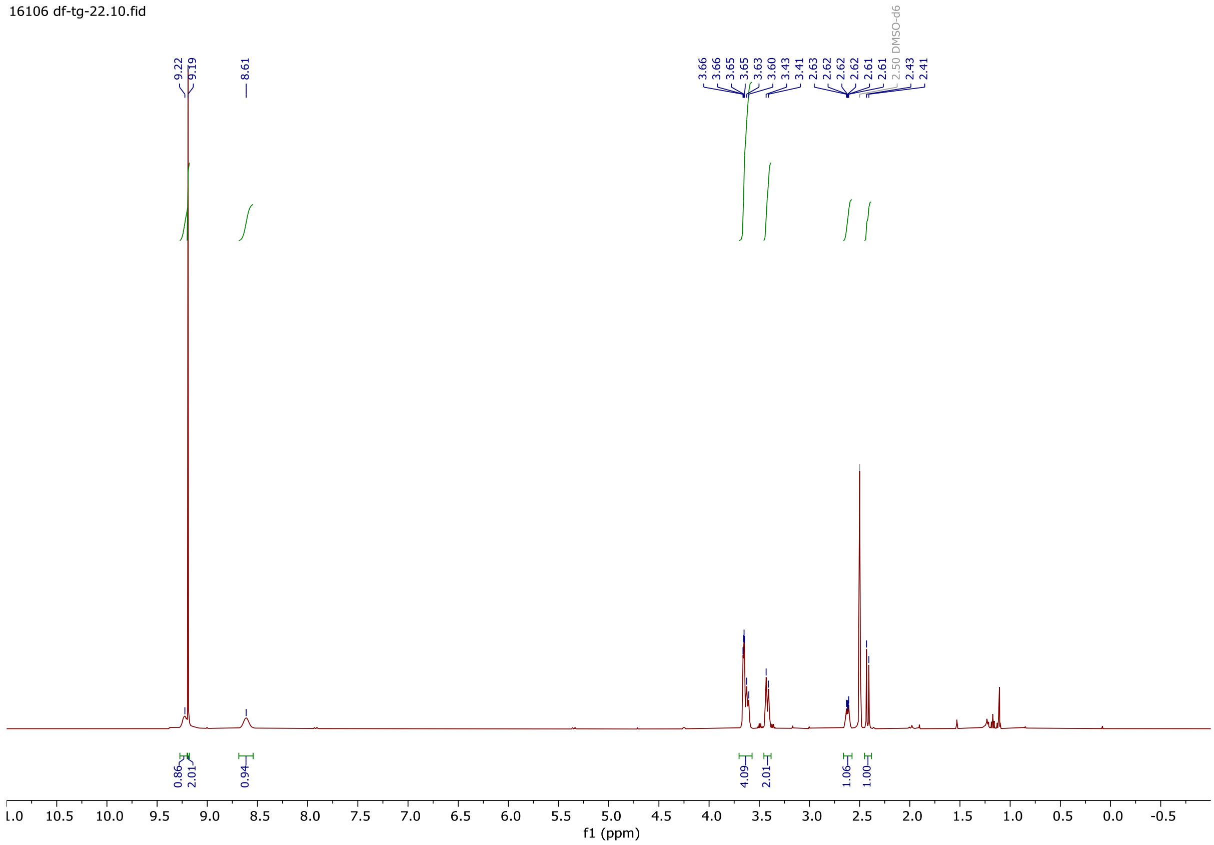


$^1$H NMR of $N_2$ varenicline TFA **6**

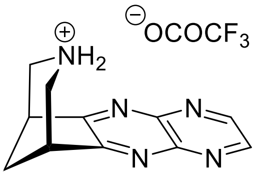

**6**

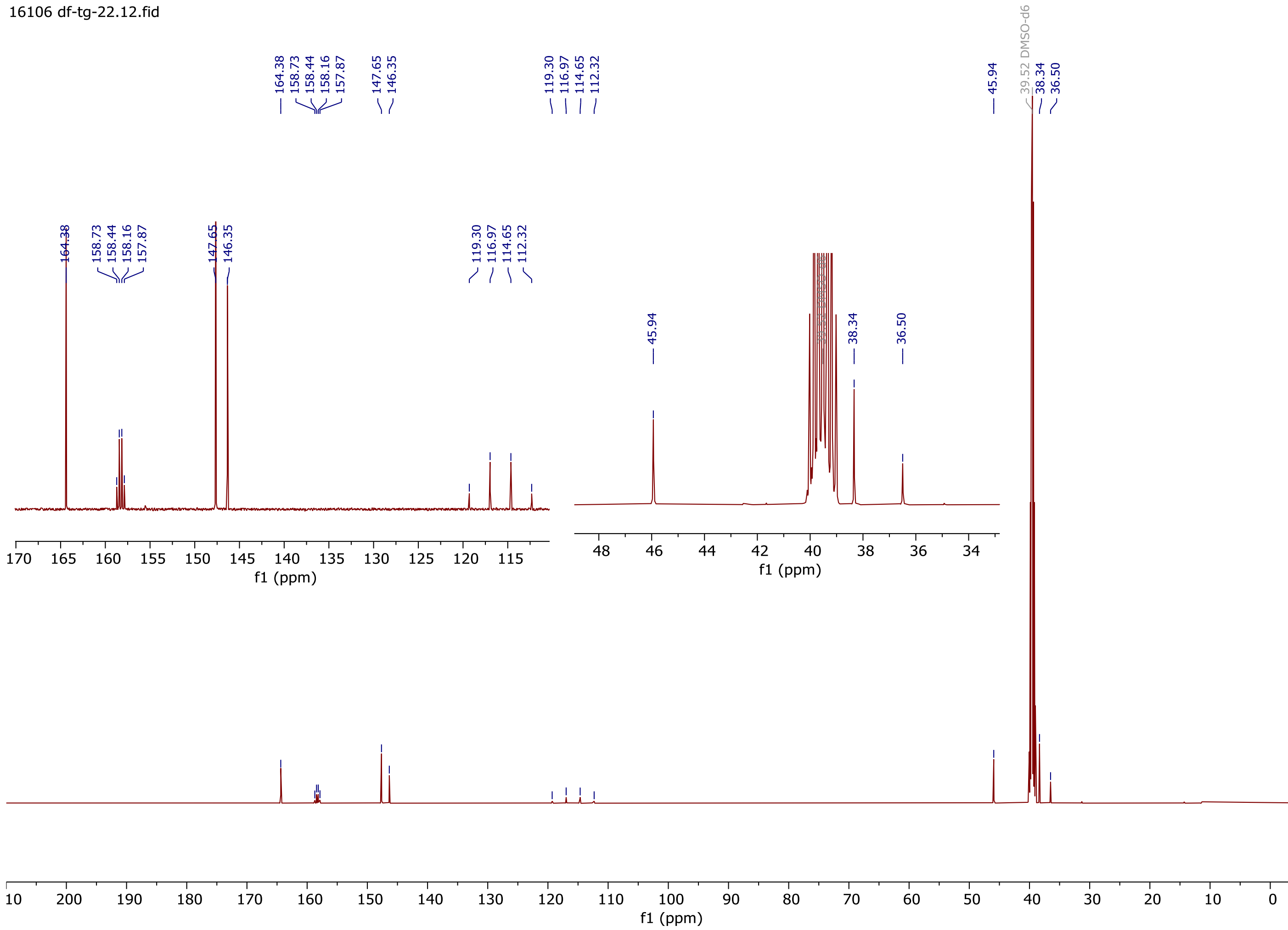


$^{13}C$ NMR (DMSO) of $N_2$ varenicline TFA **6**

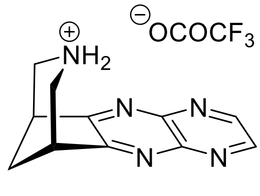


**6**

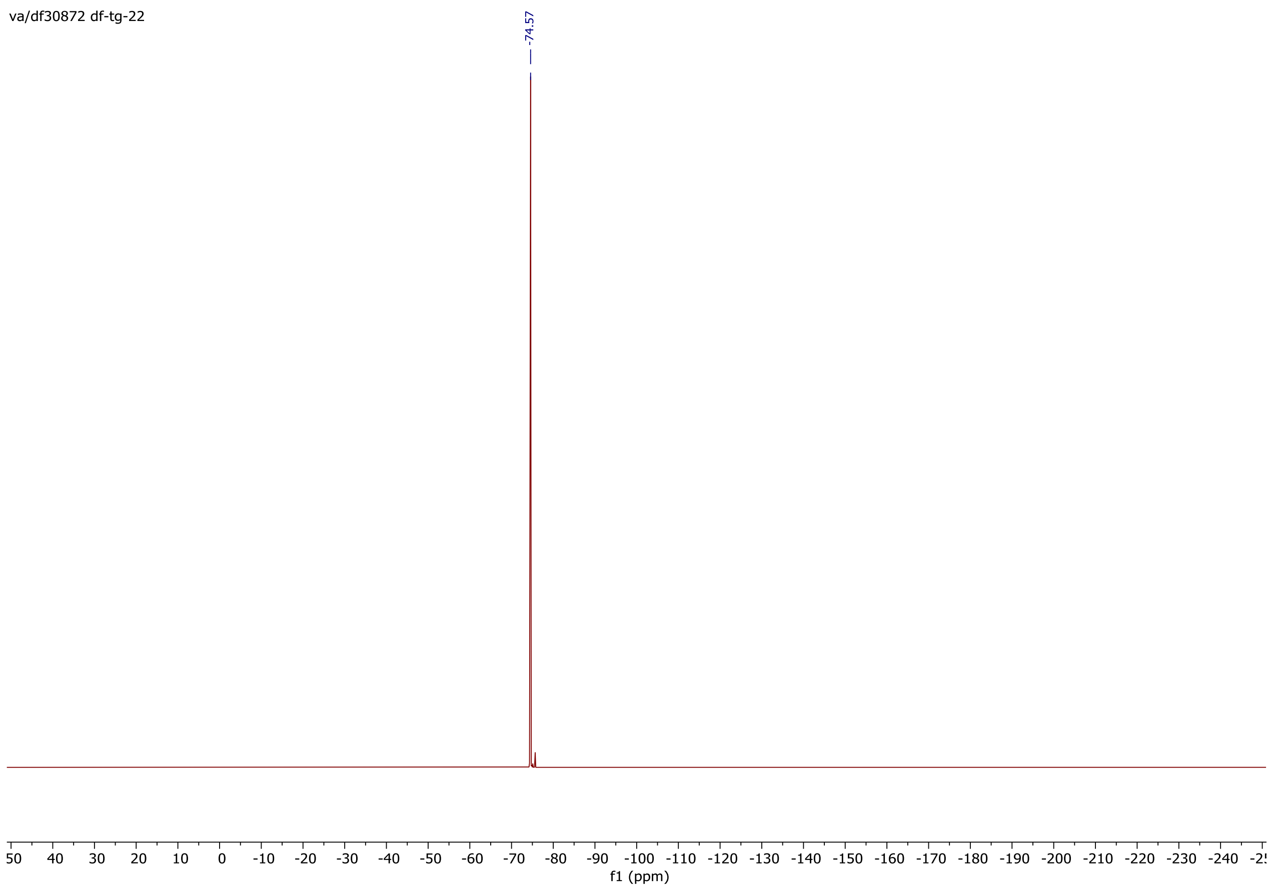


$^{19}F$ NMR of $N_2$ varenicline TFA **6**

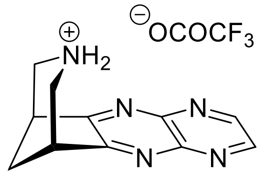


**6**

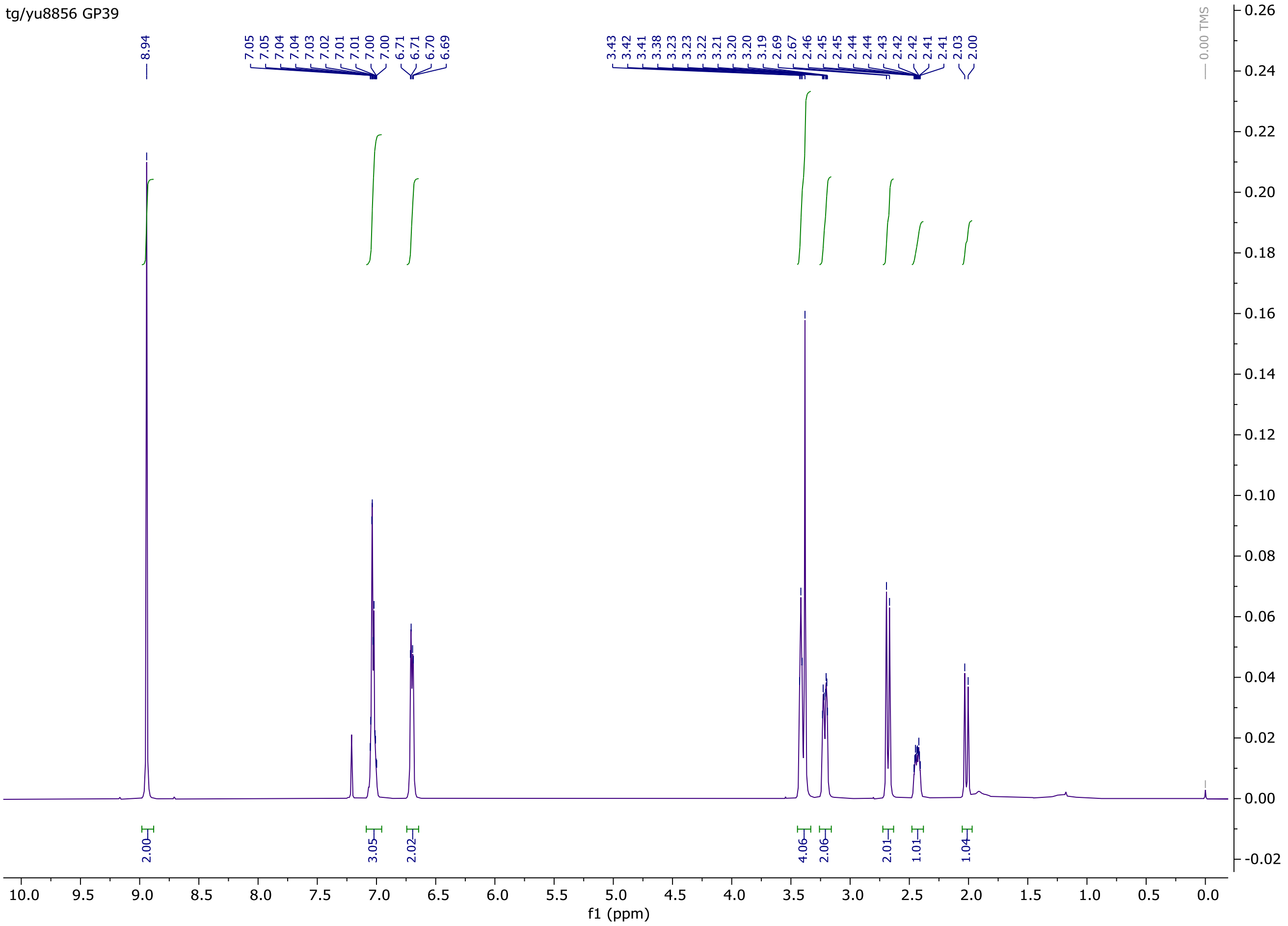


$^{1}$H NMR of N-Bn $N_2$ varenicline **SI12**

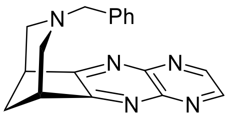


**SI12**

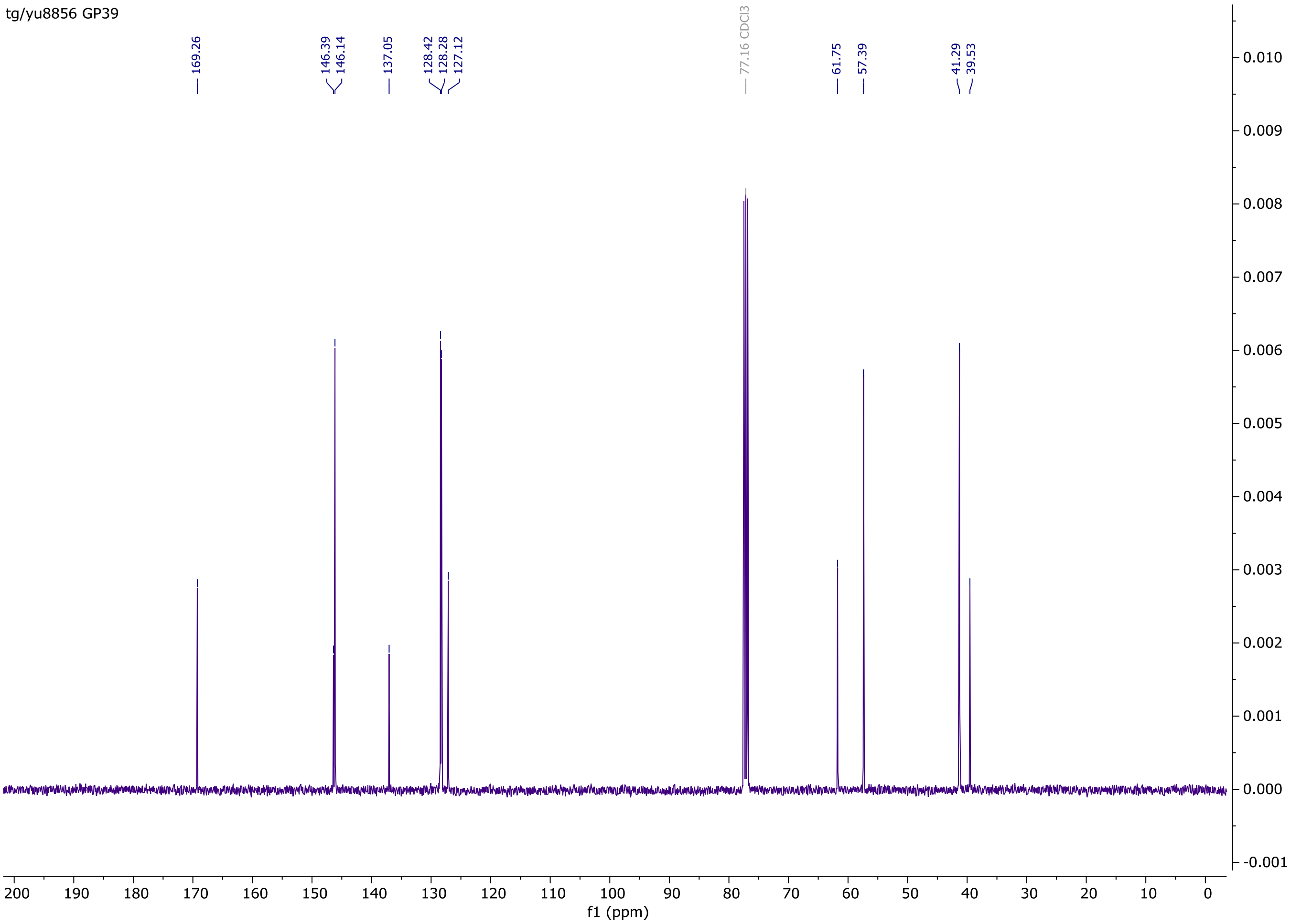


$^{13}$C NMR of N-Bn $N_2$ varenicline **SI12**

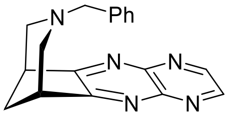


**SI12**

#### B. Computational Modelling

##### *(i)* Molecular dynamics (MD) simulations

Molecular dynamics (MD) simulations of the extracellular domain (ECD) of human wild-type $(\alpha4)_3(\beta2)_2$ and $(\alpha4)_2(\beta2)_3$ nicotinic acetylcholine receptors (nAChRs) were performed to identify the key interactions formed by different agonists within the α4α4 and α4β2 binding pockets. Since the agonists binding sites are exclusively located in the ECD, the transmembrane and intracellular domains were excluded from the simulations to reduce computational costs. The complexes between the ECD of the human low-sensitivity (LS) isoform of the α4β2 nAChR, $(\alpha4)_3(\beta2)_2$, and seven different agonists (varenicline **1**, **Var 1**; nicotine **2**, **Nct 2**; cytisine **3**, **Cyt 3**; $C_2$ varenicline **4**, **$C_2$ Var 4**; isovarenicline **5**, **Isovar 5**; $N_2$ varenicline **6**, **$N_2$ Var 6;** and, acetylcholine, **ACh**) were constructed using the cryo-EM structure of the complete $(\alpha4)_3(\beta2)_2$ receptor with nicotine bound (PDB code: 6CNK[6]). The binding modes for ACh and cytisine **3** were the same as those previously described in Minguez-Viñas *et.*[7] and Rego-Campello *et al.,*[8] respectively. The binding mode for varenicline **1** and its variants was the same as the one observed in the cryo-EM structure (PDB code: 6UR8[9]). Complexes between the ECD of the human high-sensitivity (HS) isoform of the α4β2, $(\alpha4)_2(\beta2)_3$, and varenicline **1** and the three new varenicline variants were also built based on the X-ray structure of the complete $(\alpha4)_2(\beta2)_3$ receptor with nicotine bound (PDB code: 5KXI[10]). All simulated systems contained one agonist bound to each agonist binding site: in the $(\alpha4)_3(\beta2)_2$ stoichiometry, the α-α and both α-β pockets were occupied; conversely, in the $(\alpha4)_2(\beta2)_3$ stoichiometry, each of the two non-consecutive α-β pockets contained one agonist molecule. The structures for the wild-type complexes (shown in Figures S2-S5) were used as the starting inputs for wild-type simulations.

Five mutant complexes formed by the ECD of the human $(\alpha4)_3(\beta2)_2$ nAChR and varenicline **1**, nicotine **2**, cytisine **3** and ACh were also built and simulated to investigate the dynamic and structural effect of the mutations (Table S1). The mutations introduced were: a serine-to-valine substitution in position 133 of the complementary β2 face of the α-β binding pockets (hereafter named β2S133V), a threonine-to-valine substitution in position 183 of the principal α4 face of the α-β and α-α pockets (α4T183V), a threonine-to-valine substitution in position 139 of the complementary α4 face of the α-α pocket (α4T139V) and two double mutants, one simultaneously co-expressing the α4T183V and β2S133V substitutions (β2S133Vα4T183V) and the other the α4T139V and β2S133V mutations (β2S133Vα4T139V). Note that the numbering here refers to Uniprot sequences P43681 and P17787 for the human α4 and β2 subunits, respectively. Starting structures for simulations of the mutants were created using the mutagenesis tool in PyMOL.[11]

In this work, a total of 32 different systems were investigated (see Table S1). The simulations for ECD of human $(\alpha4)_2(\beta2)_3$ nAChR with nicotine **2,** cytisine **3** and ACh were taken from our previous work.[7, 8]

All titratable residues were modelled in their standard state at a physiological pH (i.e. aspartates and glutamates were negatively charged, lysines and arginines were positively charged, and histidines were neutral), similarly to our previous work.[7, 8] All agonists (varenicline **1**, nicotine **2**, cytisine **3**, **$C_2$** varenicline **4,** isovarenicline **5** and ACh) were considered to be positively charged.

All MD simulations were performed using Gromacs.[12] The Amber ff99SB-ILDN forcefield[13] was used to describe the protein, whereas the parameters for varenicline **1**, nicotine **2,** cytisine **3** and ACh were taken from our previous work.[7, 8] Acpype[14] was used to generate Amber-compatible GAFF parameters for the **$C_2$** varenicline **4** and isovarenicline **5** ligands. All systems were solvated using the TIP3P water model.[15] The simulations were performed using a 2-fs time-step for the integration of the equations of motion. Non-bonded long-range electrostatic interactions were calculated using the smooth particle mesh Ewald method,[16] with a Fourier grid spacing of 0.16 Å and a 1.2 Å cutoff for direct contributions. A 12 Å cut-off was also used for the van der Waals interactions with long-range dispersion corrections for the energy and pressure. The LINCS algorithm[17] was used to constrain bonds in the protein and agonists, and SETTLE[18] was used to keep water molecules rigid.

Prior to the unrestrained simulations, all systems were energy minimized and initialized using the protocol described in Rego-Campello *et al.*[8] Briefly, this procedure involves a three-step energy minimization: in the first step, harmonic restraints were applied to all non-hydrogen atoms; in the second step, to Cα atoms only; and, in the third step, no restraints were used. After energy minimization, a short MD simulation step with all the non-hydrogen atoms restrained was performed, followed by a second short simulation in which position restraints were applied to Cα atoms only. All the unrestrained simulations started from these relaxed conformations.

All unrestrained simulations were performed using a constant temperature of 310 K using the velocity-rescaling thermostat,[19] with separate couplings for the solutes (protein and ligands) and solvent and a relaxation time constant of 0.1 ps. The pressure within these simulations was maintained at 1 bar using the Parrinello-Rahman barostat[20, 21] with a coupling constant of 1 ps. Each system was simulated three times, each 300 ns, leading to a total of 27 μs of simulation time across all systems. MD input and output files (including the simulation trajectories) will be made publicly available on publication *via* the University of Bristol Research Data Repository (https://data.bris.ac.uk/).

##### *(ii)* Analysis of MD simulations

The trajectories were analyzed using Gromacs tools.[12] The structural stability of the simulated systems was examined by monitoring the Cα root mean square deviations (RMSD) relative to the starting structures. All systems remained stable over the simulation time, with the average Cα RMSD profiles showing a plateau after ~20 ns (Figures S6 and S32). The receptor's secondary structure content was monitored using the DSSP software,[22] with minimal secondary structure loss after the simulation time (Figures S7 and S32).

Principal component analysis (PCA) was also performed to examine the sampling and equilibration of the replicates (Figure S8), as previously described.[23, 24] All replicates were combined before the analysis so that they all shared a common subspace, and their behaviour could be directly compared. Each trajectory used for PCA contained one conformation per nanosecond per replicate with the protein Cα atoms. The two principal components (PC) 1 and 2 were used to assess the sampling and equilibration of the simulations, and the extent to which the different replicates explore different regions of conformational space. As can be seen in Figure S8, generally, the different replicates sampled different regions of conformational, thus improving the overall sampling for each system and helping to mitigate sampling problems.

Cα root mean square fluctuations (RMSF) were calculated to characterize the dynamic behavior of the receptor across the various systems (Figures S9 and S33) and a Student's t-test was used to compare the RMSF between the wild type and mutants and to assess the significance of the differences observed (similarly to Oliveira *et al.*[23]) (Figure S10). A sample size of three was used for the t-test, which assumed the two samples were independent and the dependent variable was normally distributed.

The RMSD of the agonists was monitored to assess the stability of their initial binding poses (Figures S11-S14 and S32). Generally, despite the changes in binding mode observed for some ligands (e.g. ACh in the α-α binding pocket), all agonists remained bound to their respective binding sites in both the wild-type and mutant receptors. The only exception was the ACh molecule bound to the second α-β pocket of replicate 3 in the β2S133Vα4T139V-ACh system, which exited the pocket after about 112 ns (Figure S13G).

The statistical correlations between the protonated nitrogen atom of varenicline **1**, nicotine **2,** cytisine **3** and ACh and all the Cα atoms in the wild-type receptor was determined to identify the regions in the protein whose motions are coupled to the ligands (Figure S23). The correlations were determined by combining all replicates' trajectories for each individual system, with each trajectory containing a total of 90001 conformations.

Probability density maps were generated to visualize the spatial distribution and identify preferred positions of the agonists (Figures S15-S17) and the side-chain of α4T183, α4T139 and β2S133 (Figures S24-S25) for the wild-type and mutant complexes. For each agonist-receptor system, the maps were calculated by combining the entire trajectories for all replicates of that system.

##### *(iii)* **Supporting figures and tables**

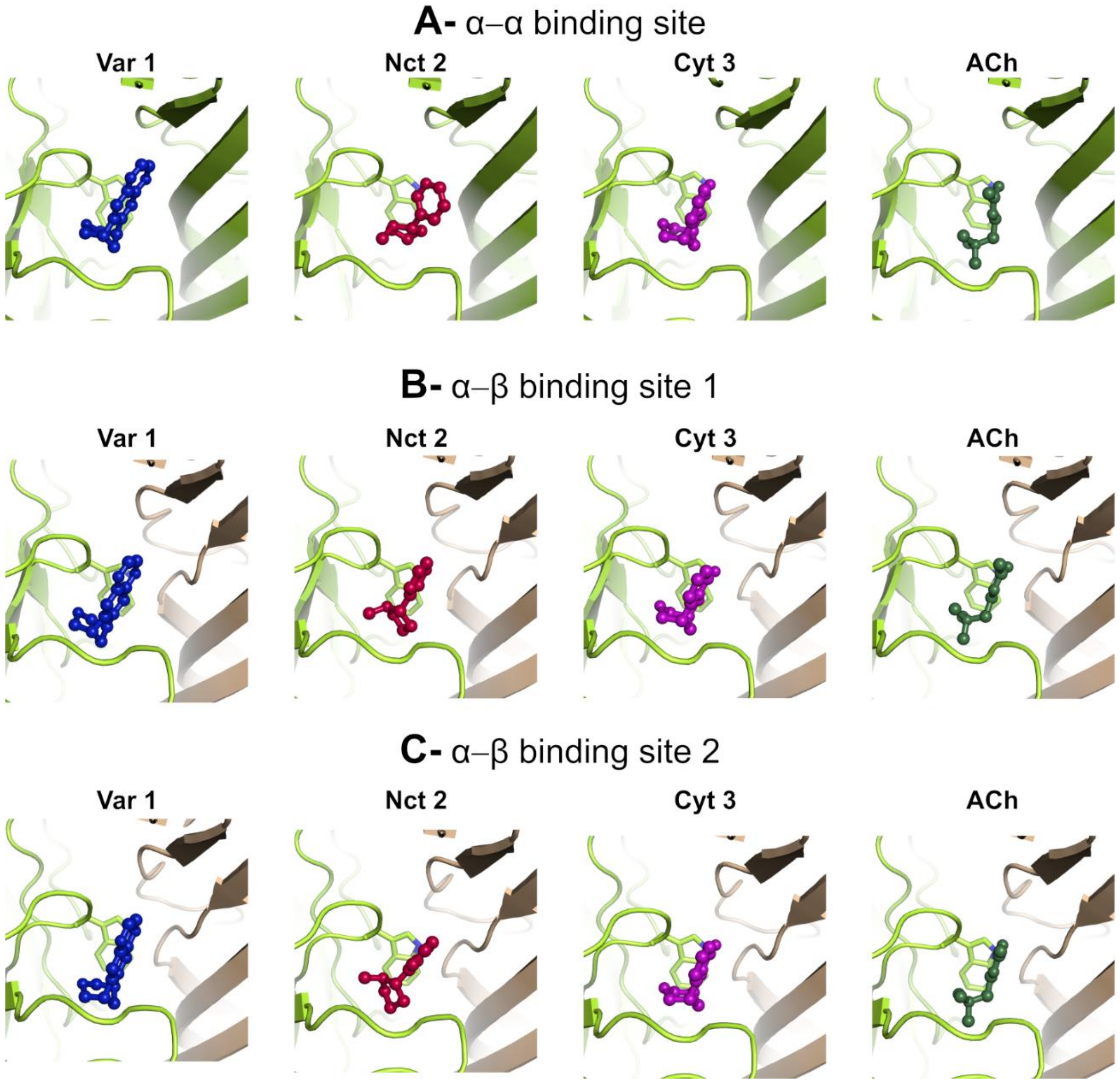


**Figure S2-** Agonist binding mode in the α-α and α-β pockets in the wild-type LS $(\alpha 4)_3(\beta 2)_2$ nAChR. (**A**) Binding mode of **1**, nicotine **2**, cytisine **3,** and ACh in the α-α pocket. (**B**) Binding mode of varenicline **1**, nicotine **2**, cytisine **3** and ACh in the first α-β pocket (**C**) Binding mode of varenicline **1**, nicotine **2**, cytisine **3** and ACh in the second α-β pocket. Agonists are represented as balls-and-sticks and TrpB (W182 in the principal α4 face), a tryptophan residue that provides the anchor point for the agonist, shown with sticks. The α4 and β2 subunits are colored  in yellow and light brown, respectively.

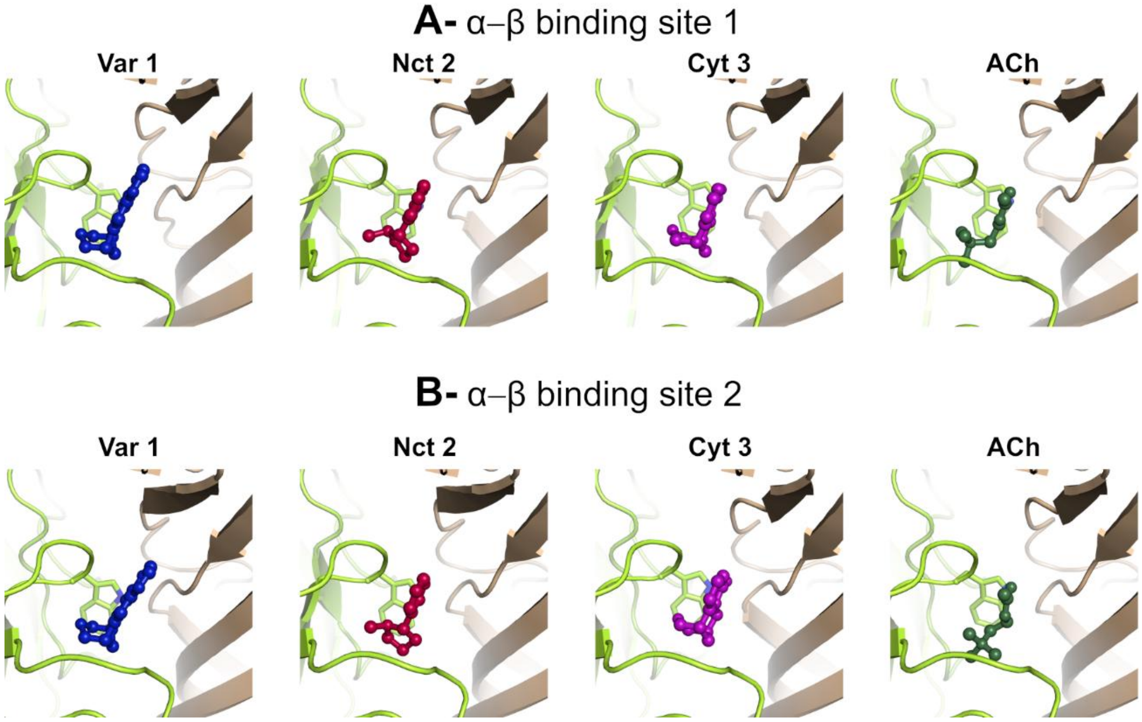


**Figure S3-** Agonist binding mode in the α-β pockets in the wild-type HS $(\alpha4)_2(\beta2)_3$ nAChR. (**A**) Binding mode of varenicline **1**, nicotine **2**, cytisine **3**, and ACh in the first α-β pocket (**B**) Binding mode of varenicline **1**, nicotine **2**, cytisine **3**, and ACh in the second α-β pocket. Agonists are represented as balls-and-sticks and TrpB (W182 in the principal α4 face), a tryptophan residue that provides the anchor point for the agonist, shown with sticks. The α4 and β2 subunits are colored in yellow and light brown, respectively.

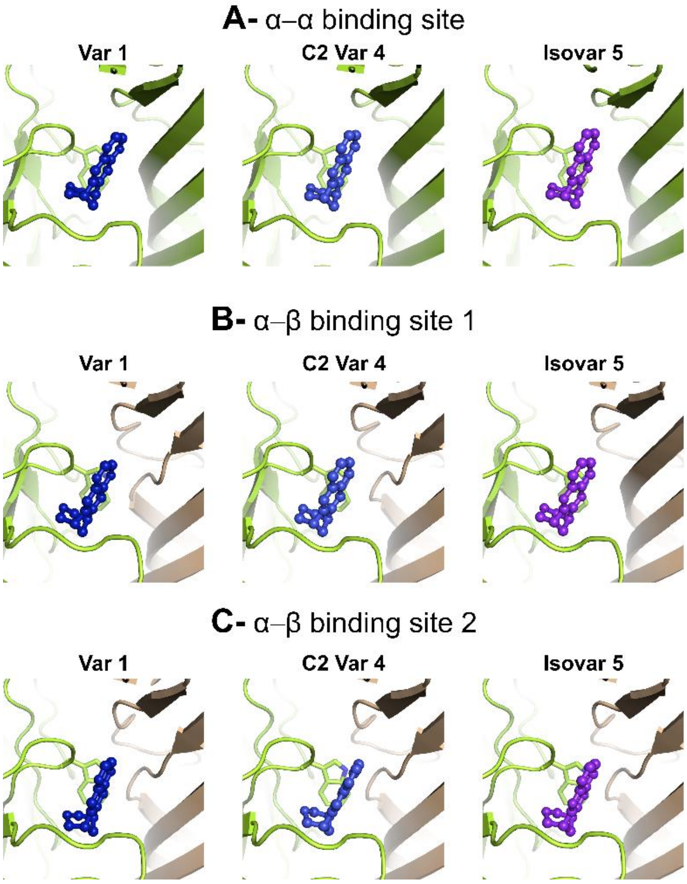


**Figure S4-** Binding mode of varenicline **1** and its variants in the α-α and α-β pockets of the wild-type LS $(\alpha4)_3(\beta2)_2$ nAChR. (**A**) Binding mode of varenicline **1**, $C_2$ varenicline **4,** and isovarenicline **5** in the α-α pocket. (**B**) Binding mode of varenicline **1**, $C_2$ varenicline **4**, isovarenicline **5** and $N_2$ varenicline **6** in the first α-β pocket (**C**) Binding mode of varenicline **1**, $C_2$ varenicline **4**, and isovarenicline **5** in the second α-β pocket. Agonists are represented as balls-and-sticks and TrpB (W182 in the principal α4 face), a tryptophan residue that provides the anchor point for the agonist, shown with sticks. The α4 and β2 subunits are colored in yellow and light brown, respectively.

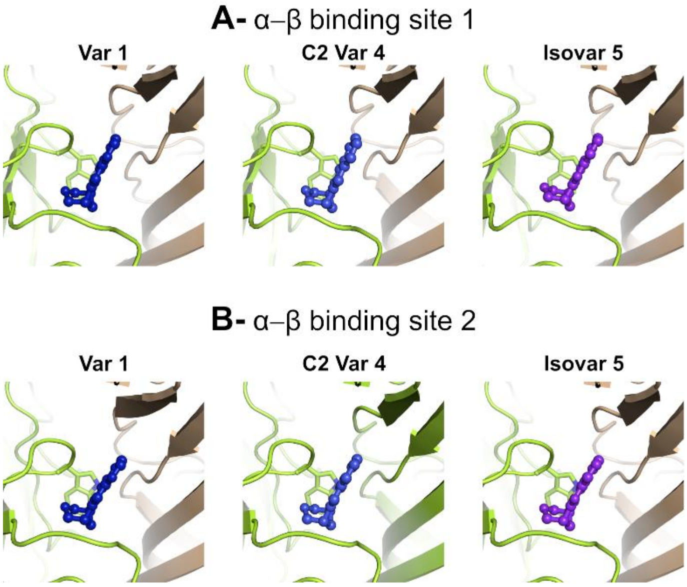


**Figure S5-** Binding mode of varenicline **1** and its variants in the α-β pockets of the wild-type HS $(\alpha4)_2(\beta2)_3$ nAChR. (**A**) Binding mode of varenicline **1**, $C_2$ varenicline **4**, and isovarenicline **5** in the first α-β pocket (**B**) Binding mode of varenicline **1**, $C_2$ varenicline **4**, and isovarenicline **5** in second α-β pocket. Agonists are represented as balls-and-sticks and TrpB (W182 in the principal α4 face), a tryptophan residue that provides the anchor point for the agonist, shown with sticks. The α4 and β2 subunits are colored in yellow and light brown, respectively.

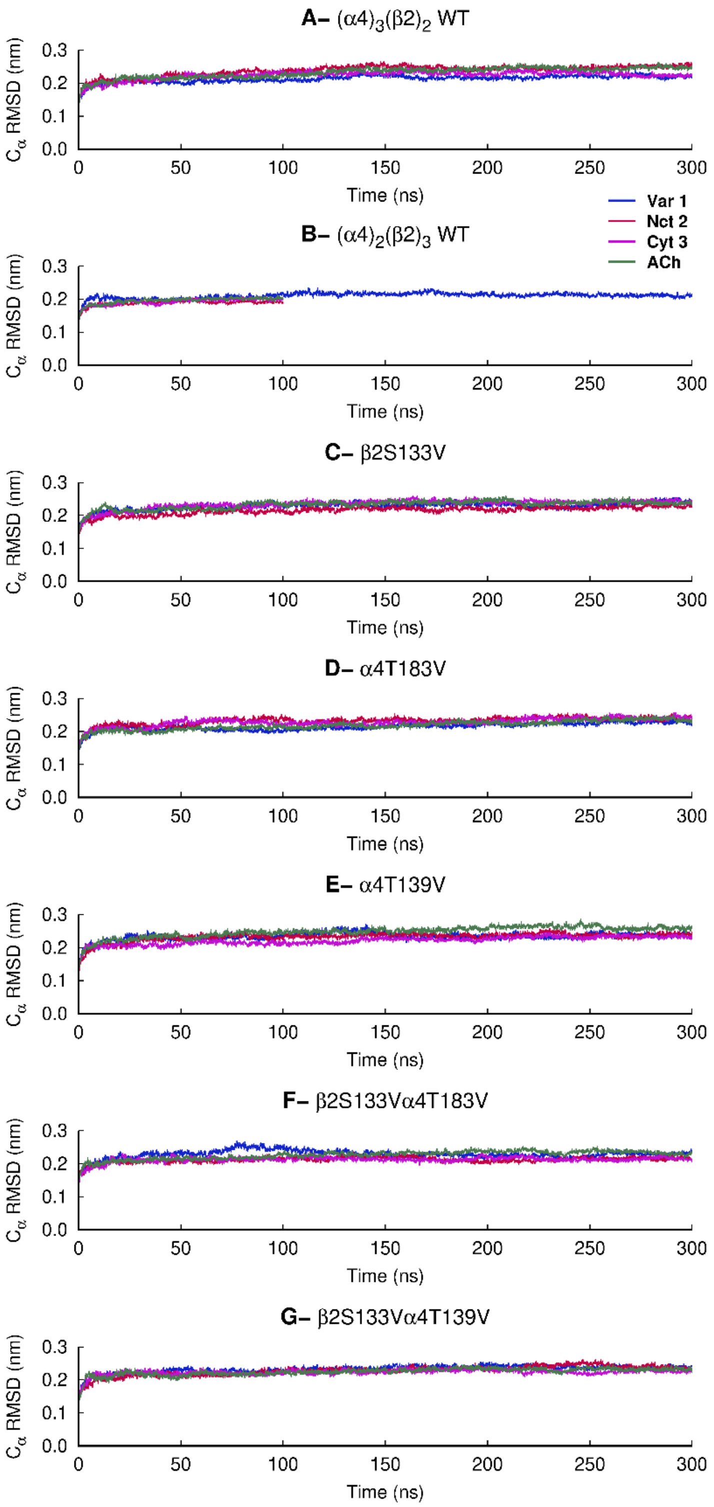


**Figure S6-** Temporal evolution of the average Cα RMSD for the varenicline **1**, nicotine **2**, cytisine **3** and ACh bound systems. The Cα RMSD was calculated relative to the starting structures, and the averages were obtained over all replicates for each system. Please note that simulations of the complexes between the HS $(\alpha4)_2(\beta2)_3$ wild-type receptor and nicotine **2**, cytisine **3** and ACh (panel B) were taken from our previous work.[7, 8] Please zoom in on the image for detailed visualization.

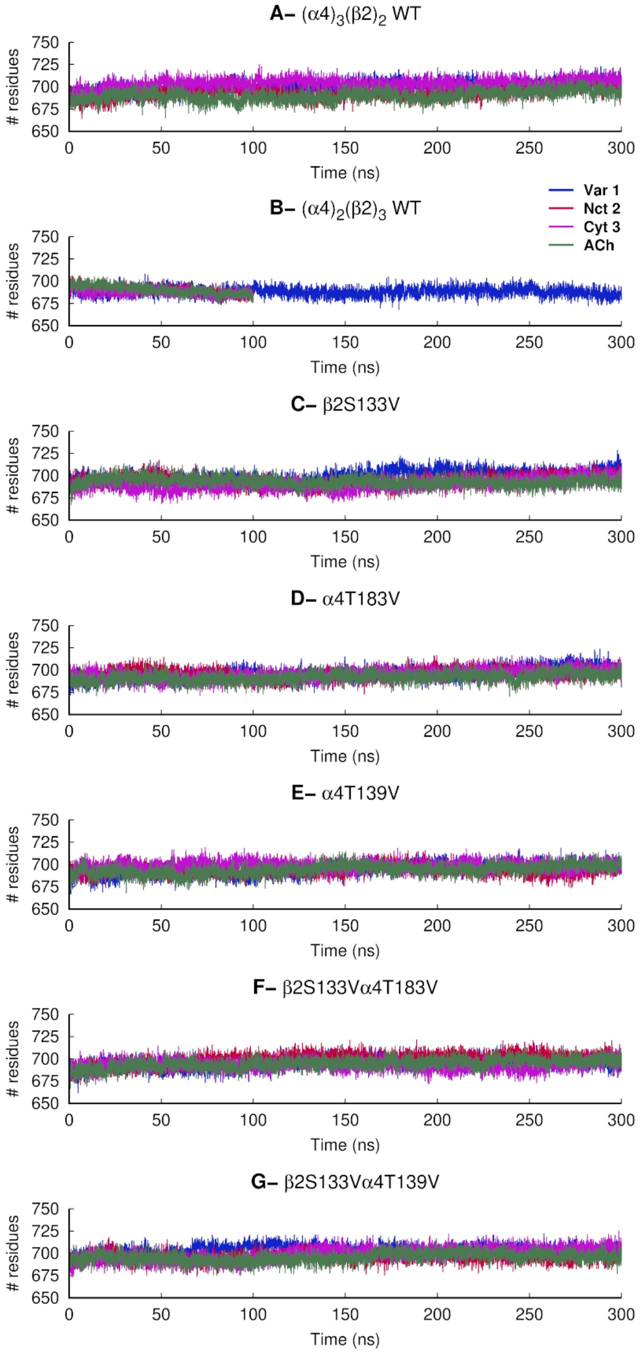


**Figure S7-** Temporal evolution of the number of residues involved in secondary structure features for the varenicline **1**, nicotine **2**, cytisine **3** and ACh bound systems. The secondary structure assignment was performed with the DSSP software[22] and includes all residues assigned to α-helix, π-helix, $3_{10}$- helix, 5-helix, β-sheet, β-strand and β-bridge secondary structure classes. The averages were obtained over all replicates for each system. The trajectories for the complexes HS $(\alpha4)_2(\beta2)_3$ wild-type receptor and nicotine **2**, cytisine **3** and ACh (in panel B) were taken from our previous work.[7, 8] Please zoom in on the image for detailed visualization.

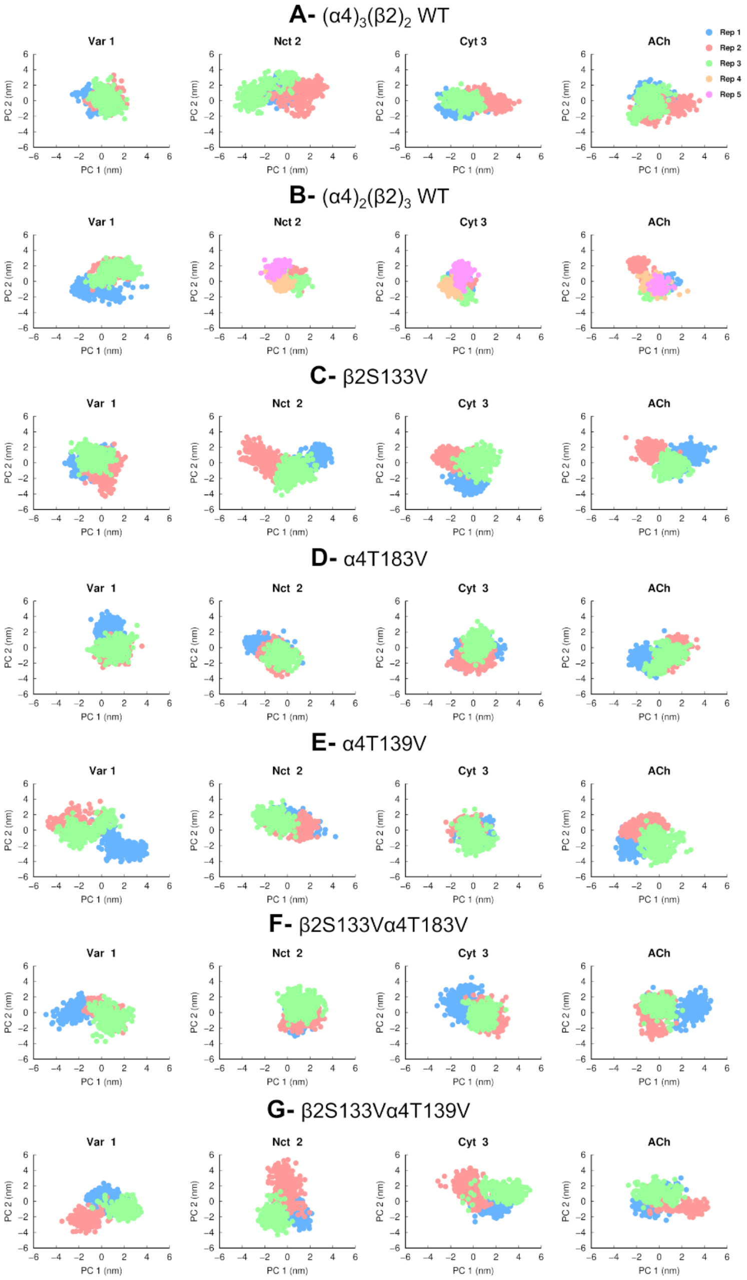


**Figure S8**- Principal component analysis (PCA) of the replicates for the ECD of the LS $(\alpha4)_3(\beta2)_2$ (**A**) and HS $(\alpha4)_2(\beta2)_3$ (**B)** wild-type and mutant (**C-G**) simulations. All replicates for each system were combined before the analysis so that they all shared a common subspace, and their behavior could be directly compared. Each trajectory used for PCA contained one conformation per nanosecond per replicate with the protein Cα atoms. PC 1 and 2 were used to assess the sampling of the simulations, and the extent to which the different replicates explore different regions of conformational space. This analysis shows that, generally, the different replicates sample different regions of the space, thus improving the overall sampling for each system. Please zoom in on the image for detailed visualization.

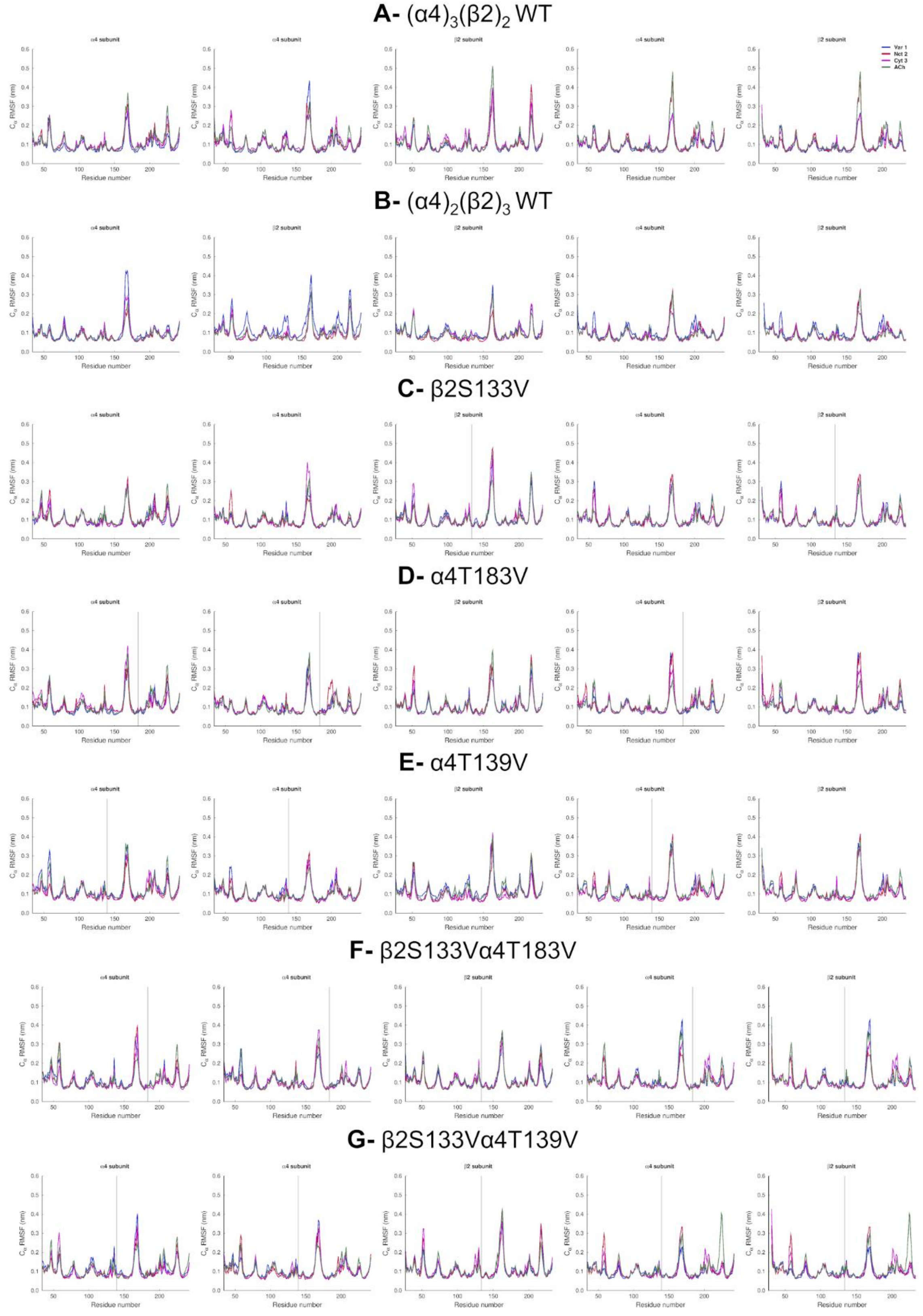


**Figure S9**- Average Cα RMSF for the LS $(\alpha4)_3(\beta2)_2$ (**A**) and HS $(\alpha4)_2(\beta2)_3$ (**B)** wild-type and mutant (**C-G**) complexes. The Cα RMSF was calculated using the entire trajectories and averaged across all replicates for each complex. The vertical grey lines highlight the position of the mutations. Please zoom in on the image for detailed visualization.

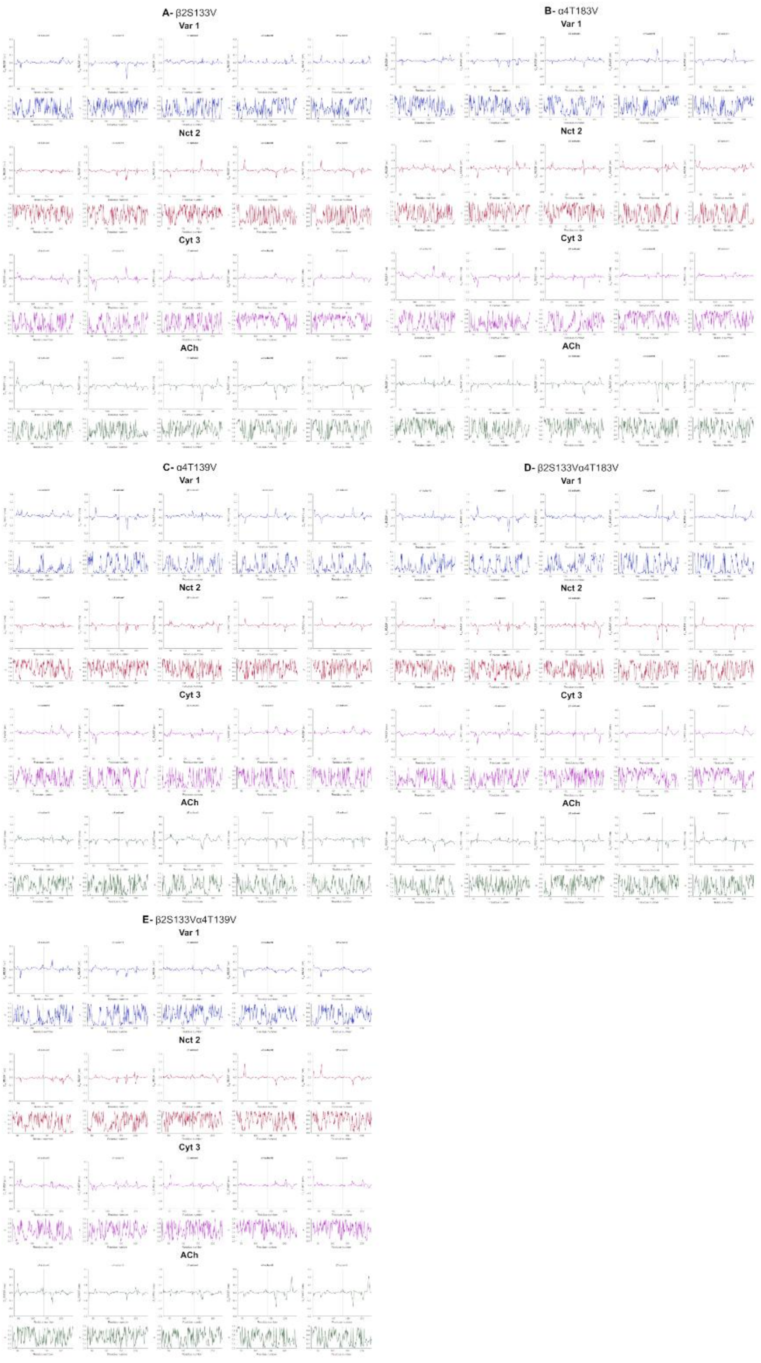


**Figure S10**- Average Cα RMSF difference between wild-type and mutant systems (**A-E**) and associated *p* values. A Student's t-test was used to compare the wild-type and mutant systems and to assess the significance of the differences. Positive values in the Cα RMSF difference plots correspond to a greater flexibility of the wild type during the simulations, whereas negative values correspond to an increased flexibility of the mutant. The vertical grey lines highlight the position of the mutations. Please zoom in on the image for detailed visualization.

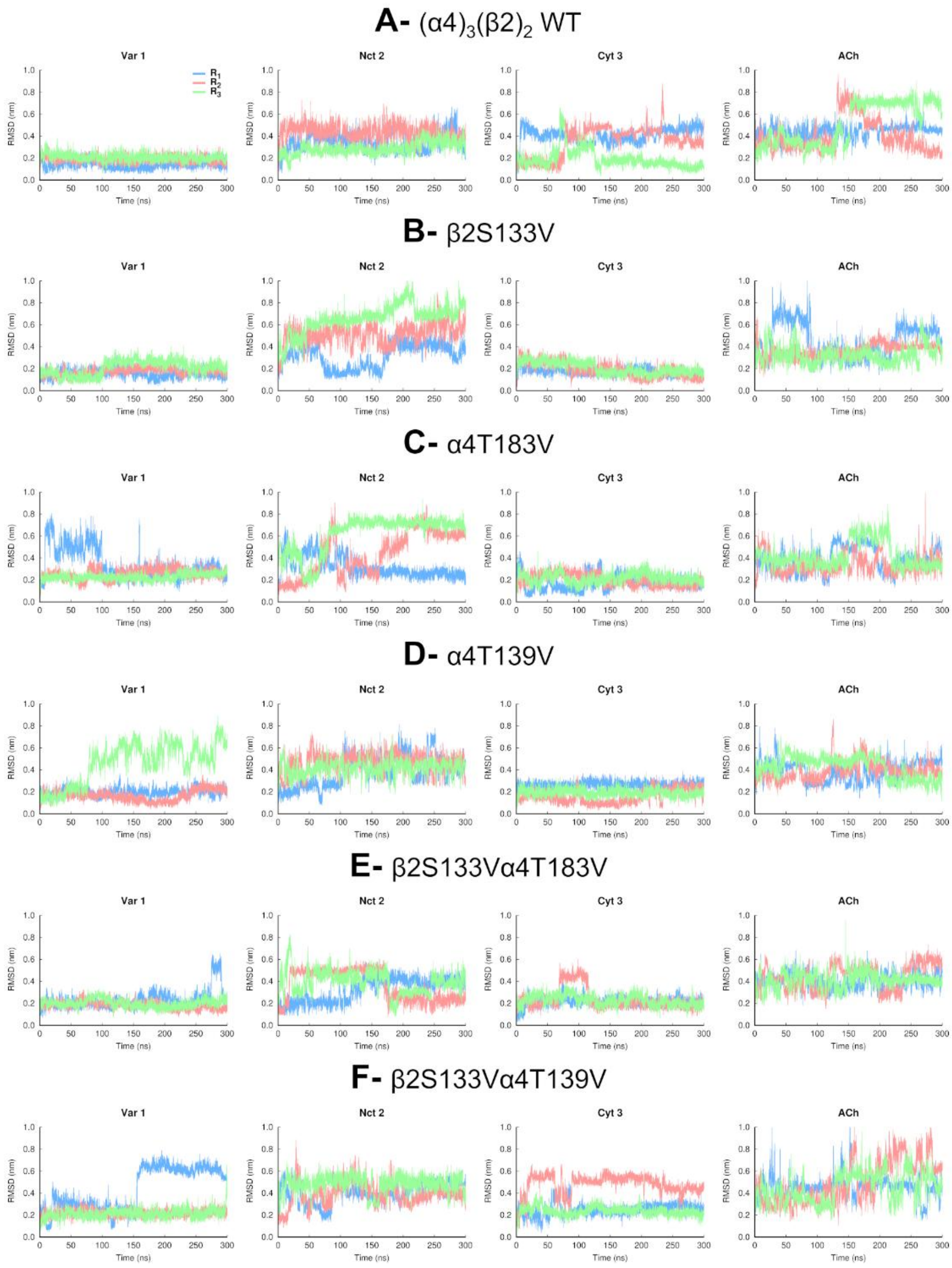


**Figure S11-** Temporal evolution of the RMSD for the agonists bound to the α-α binding pocket in the LS $(\alpha4)_3(\beta2)_2$ (**A**) wild-type and mutant (**B**-**F**) systems. The RMSD was determined with respect to the initial binding mode of the agonists at the start of the simulations. Note that the α-α binding pocket only exists in the LS $(\alpha4)_3(\beta2)_2$ isoform of the receptor. Note also that despite the changes in binding mode observed for some ligands, all agonists remained stably bound to their respective binding sites. Please zoom in on the image for detailed visualization.

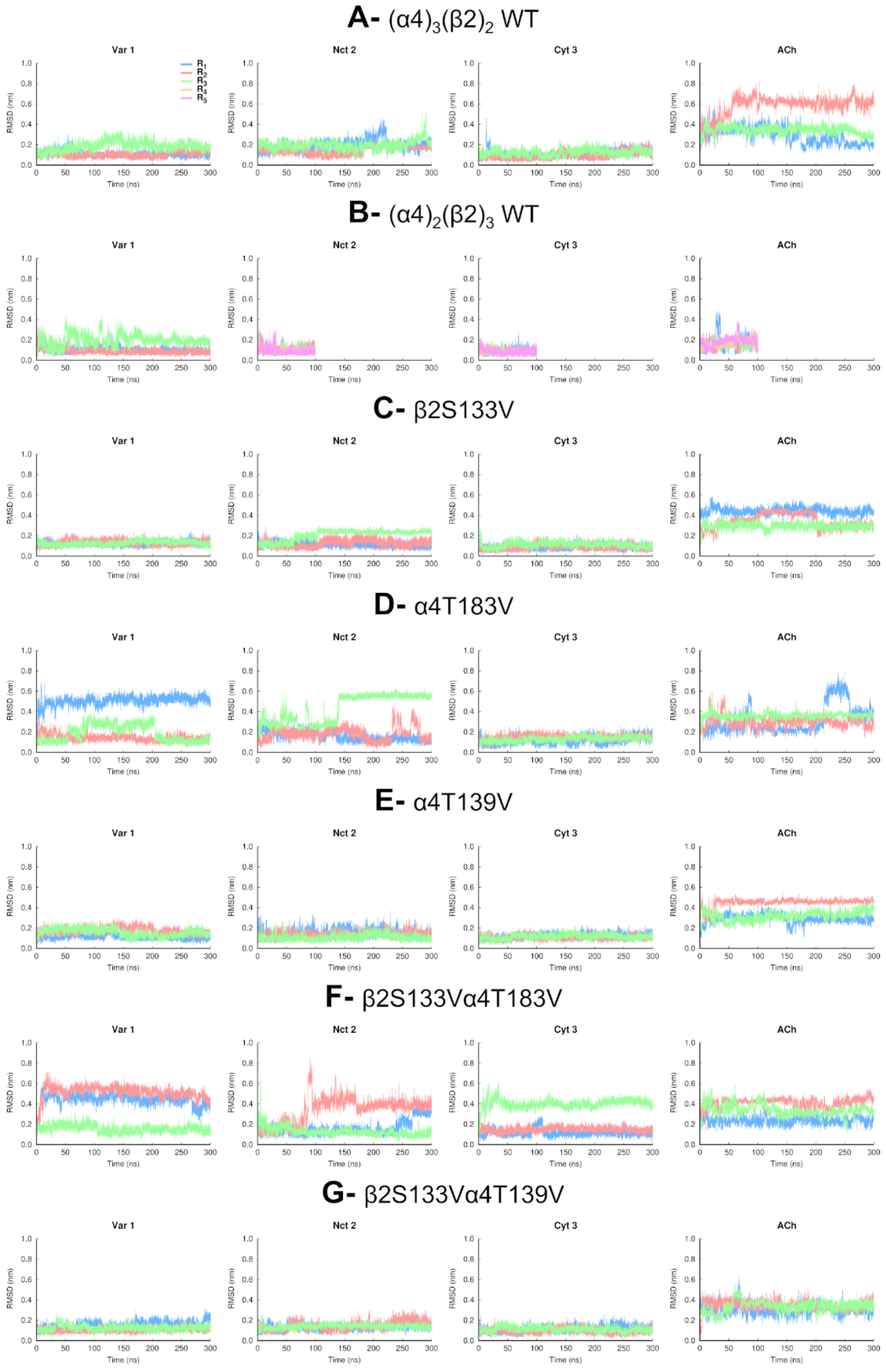


**Figure S12-** Temporal evolution of the RMSD for the agonists bound to the first α-β pocket in the LS $(\alpha4)_3(\beta2)_2$ (**A**) and HS $(\alpha4)_2(\beta2)_3$ (**B**) wild-type and mutant (**C**-**G**) systems. The RMSD was determined with respect to the initial binding mode of the agonists at the start of the simulations. Note that despite the changes in binding mode observed for some ligands, all agonists remained stably bound to their respective binding sites. Please zoom in on the image for detailed visualization.

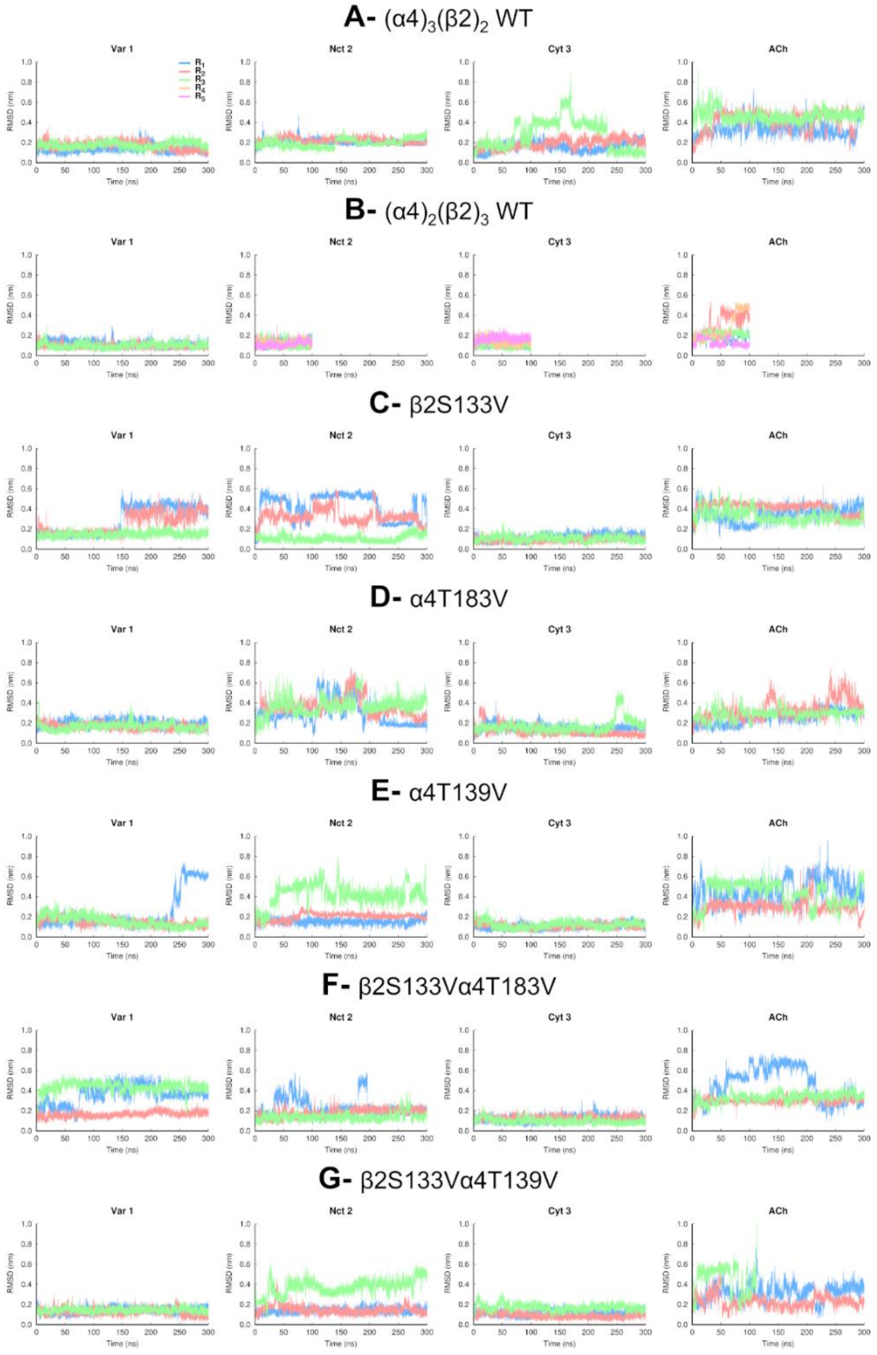


**Figure S13-** Temporal evolution of the RMSD for the agonists bound to the second α-β pocket in the LS $(\alpha4)_3(\beta2)_2$ (**A**) and HS $(\alpha4)_2(\beta2)_3$ (**B**) wild-type and mutant (**C**-**G**) systems. The RMSD was determined with respect to the initial binding mode of the agonists at the start of the simulations. Note that all agonists remained stably bound to their respective binding sites, except for the ACh molecule in the second α-β pocket for replicate 3 in the β2S133Vα4T139V-ACh complex, which exited the pocket after 112 ns (as indicated by the green line in the rightmost plot of panel G). Please zoom in on the image for detailed visualization.

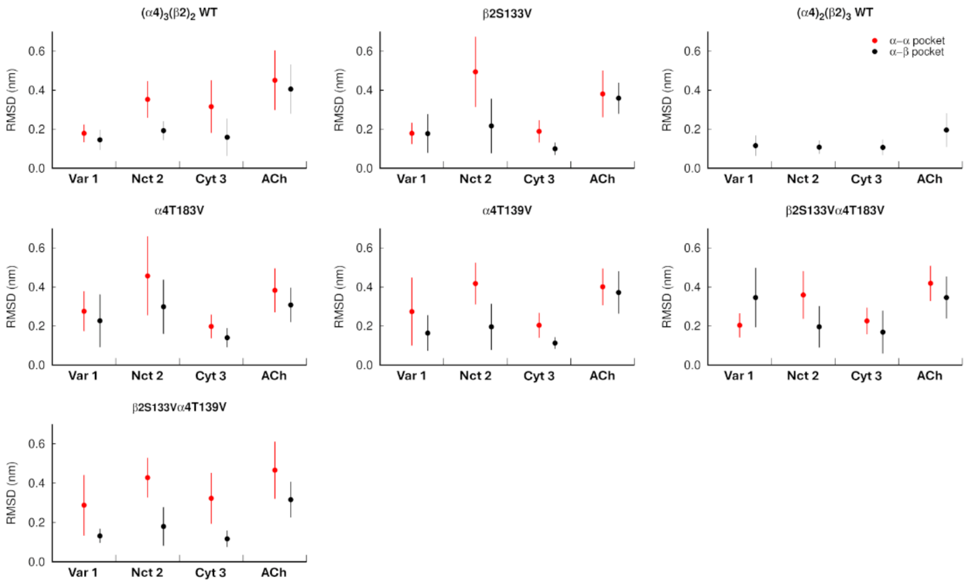


**Figure S14**- Average RMSD values for the different agonists (namely varenicline **1**, nicotine **2**, cytisine **3** and ACh) in the LS $(\alpha4)_3(\beta2)_2$ and HS $(\alpha4)_2(\beta2)_3$ wild-type and mutant simulations. Plots show the average RMSD values for the agonists in the α-α (red) and α-β (black) binding pockets. The RMSDs were calculated relative to the agonists' initial position. For each system, RMSDs were averaged over the entire trajectories for all replicate simulations. In the plots, the average RMSD values are represented by spheres, while the vertical lines indicate the standard deviation (SD) within the dataset. Please note that the LS $(\alpha4)_3(\beta2)_2$ isoform of the receptor contains both α-α and α-β binding sites, while the HS $(\alpha4)_2(\beta2)_3$ one only contains α-β sites. The average value for the α-β pocket shown was calculated over the two α-β pockets present in each complex. Note also that the average RMSD value and SD for the α-β pocket of β2S133Vα4T139V-ACh complex excludes the data for the second site from 112 ns onwards for replicate 3, as the agonist exits this binding pocket in this simulation.

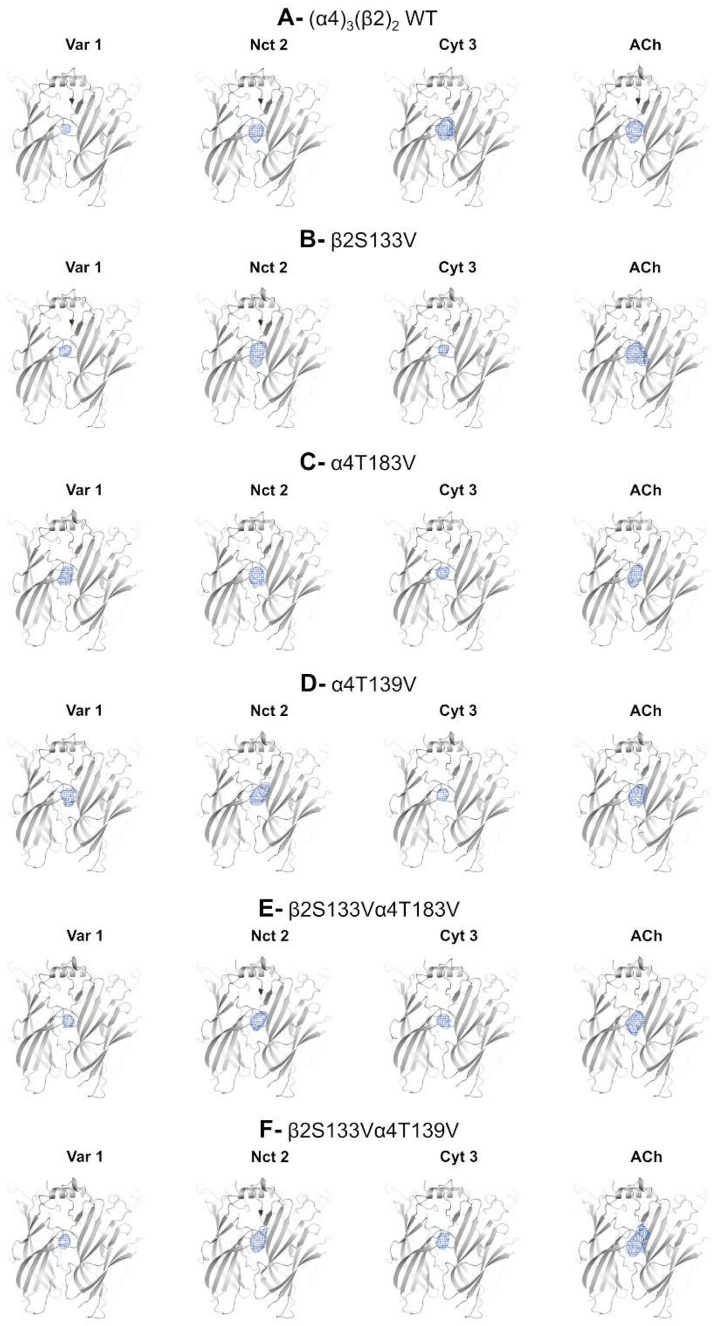


**Figure S15-** Probability density maps for the agonists bound to the α-α binding pocket in the wild-type (**A**) and mutant (**B-F**) LS $(\alpha4)_3(\beta2)_2$ simulations. The contours at 0.00001 $Å^{-3}$ for the protonated nitrogen atoms of the agonist are depicted as a blue mesh. For all systems, the maps were calculated combining the entire trajectories for each one of the three replicates of that system. Please zoom in on the image for detailed visualization.

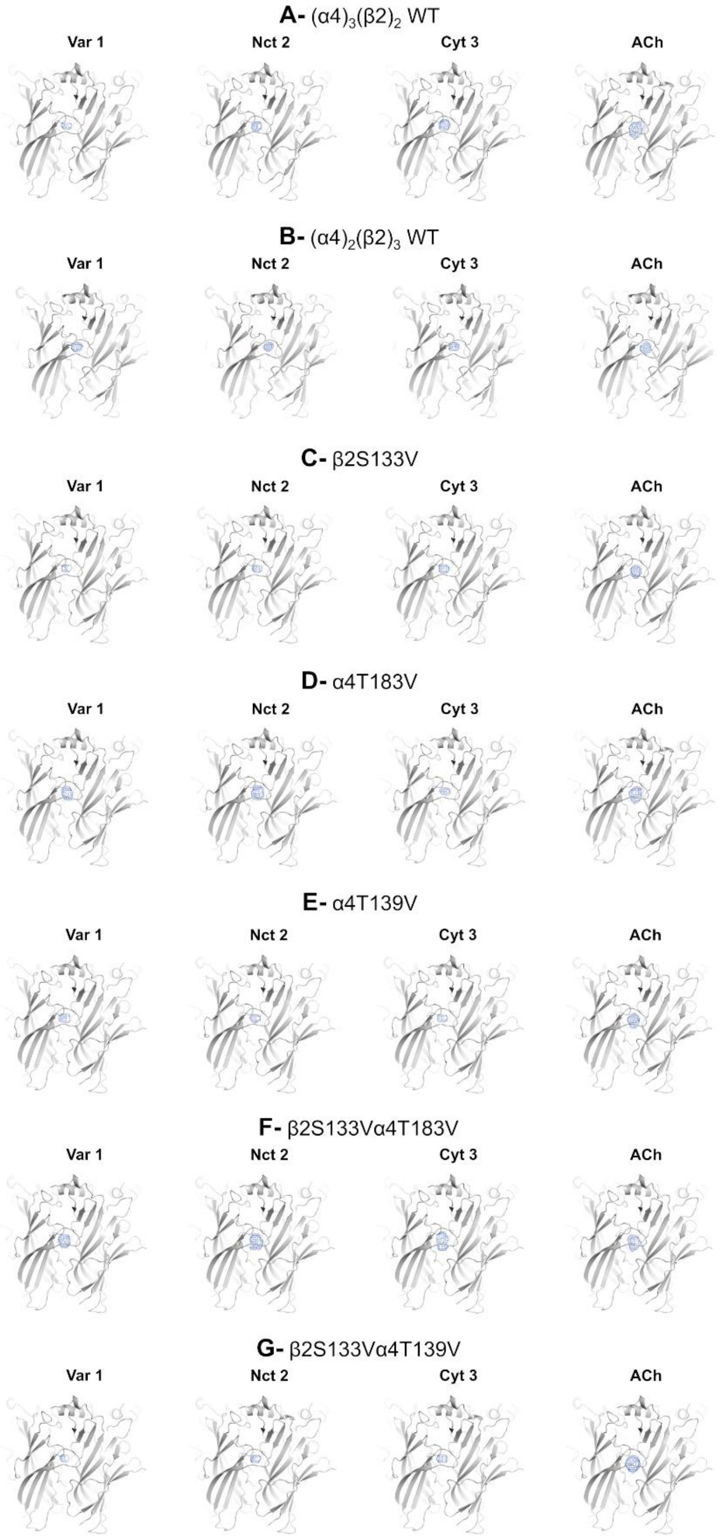


**Figure S16-** Probability density maps for the agonists bound to the first α-β binding pocket in the LS $(\alpha4)_3(\beta2)_2$ and HS $(\alpha4)_2(\beta2)_3$ wild-type (**A-B**) and mutant (**C-G**) simulations. The contours at 0.00001 $Å^{-3}$ for the protonated nitrogen atoms of the agonist are depicted as a blue mesh. For all systems, the maps were calculated combining the entire trajectories for each one of the three replicates of that system. Please zoom in on the image for detailed visualization.

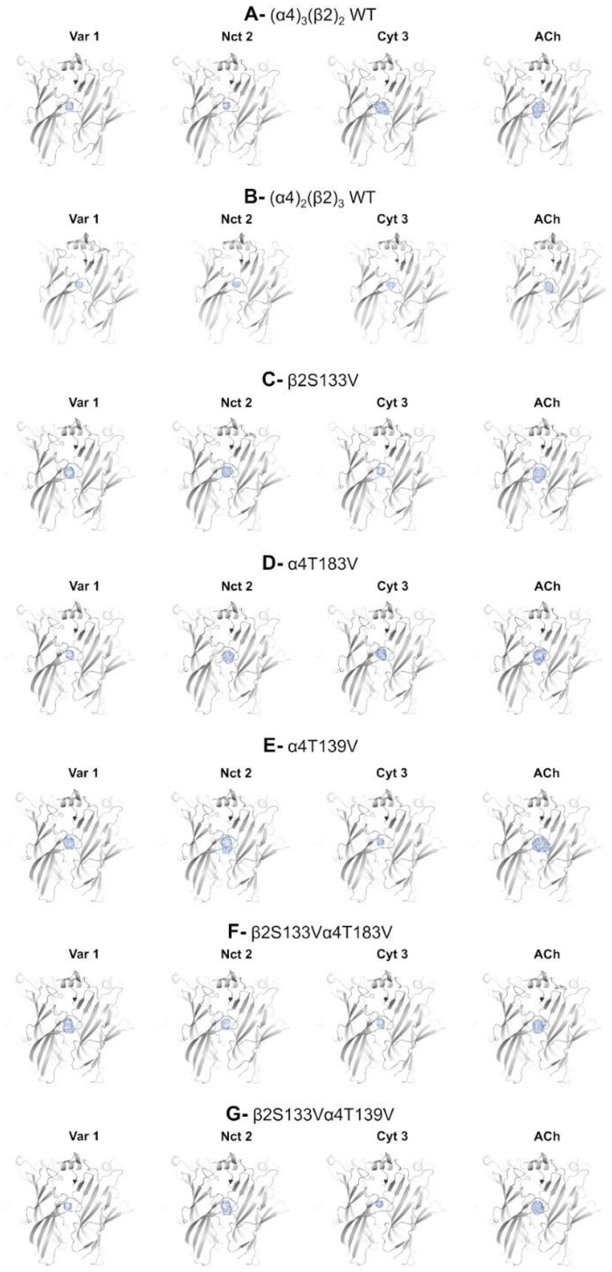


**Figure S17-** Probability density maps for the agonists bound to the second α-β binding pocket in the LS $(\alpha4)_3(\beta2)_2$ and HS $(\alpha4)_2(\beta2)_3$ wild-type (**A-B**) and mutant (**C-G**) simulations. The contours at 0.00001 $Å^{-3}$ for the protonated nitrogen atoms of the agonist are depicted as a blue mesh. For all systems, with the exception of β2S133Vα4T139V-ACh, the maps were calculated combining the entire trajectories for each one of the three replicates of that system. For the β2S133Vα4T139V-ACh complex, the map was obtained using the entire trajectories for replicates 1 and 2 and the first 112 ns for replicate 3 (as ACh exits the second α-β binding pocket after 112 ns of simulation). Please zoom in on the image for detailed visualization.

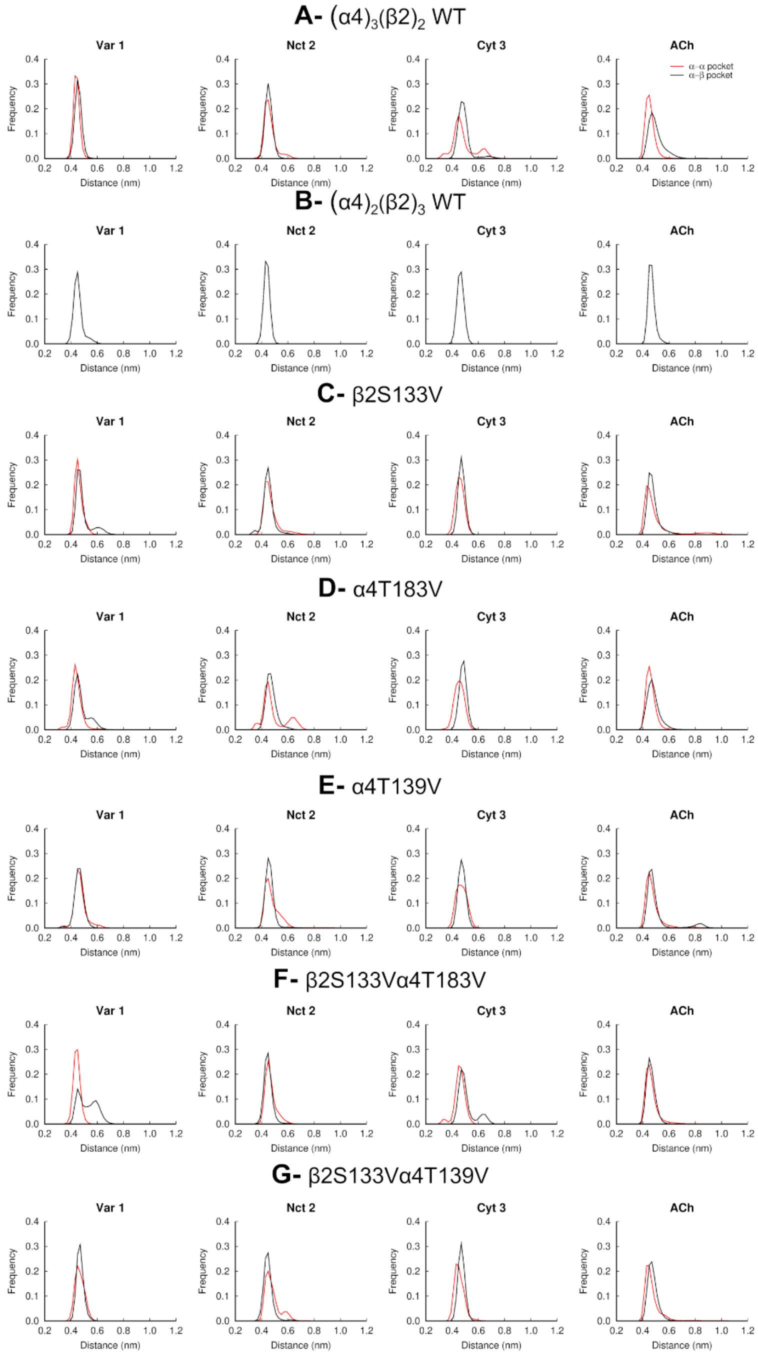


**Figure S18**- TrpB-agonist distance for the LS $(\alpha4)_3(\beta2)_2$ (**A**) and HS $(\alpha4)_2(\beta2)_3$ (**B**) wild-type and mutant (**C-G**) systems. The distance between the side-chain of TrpB (W182 in the principal α4 subunit) and the protonated nitrogen atom of varenicline **1**, nicotine **2**, cytisine **3** and ACh for the α-α (red line) and α-β (black line) binding pockets is shown. The histogram for the α-β pocket reflects the distances over the two α-β binding pockets present in both the LS and HS isoforms of the α4β2 nAChR. Please note that the histogram for the α-β binding pocket of β2S133Vα4T139V-ACh complex excludes the data for the second α-β pocket from 112 ns onwards for replicate 3, as the agonist exits this binding pocket in the simulation after 112 ns.

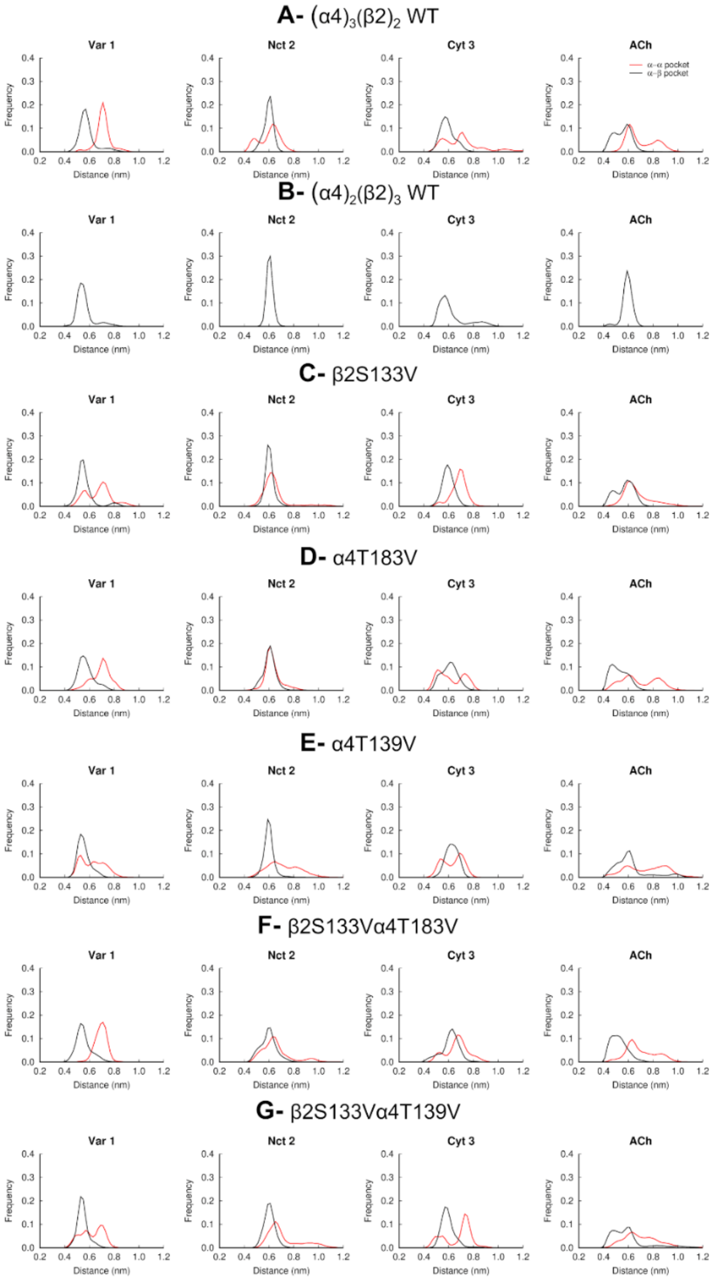


**Figure S19**- TyrA-agonist distance for the LS $(\alpha4)_3(\beta2)_2$ (**A**) and HS $(\alpha4)_2(\beta2)_3$ (**B**) wild-type and mutant (**C-G**) systems. The distance between the side-chain of TyrA (Y126 in the principal α4 subunit) and the protonated nitrogen atom of varenicline **1**, nicotine **2**, cytisine **3** and ACh for the α-α (red line) and α-β (black line) binding pockets is shown. The histogram for the α-β pocket reflects the distances over the two α-β binding pockets present in both the LS and HS isoforms of the α4β2 nAChR. Please note that the histogram for the α-β binding pocket of β2S133Vα4T139V-ACh complex excludes the data for the second α-β pocket from 112 ns onwards for replicate 3, as the agonist exits this binding pocket in the simulation after 112 ns.

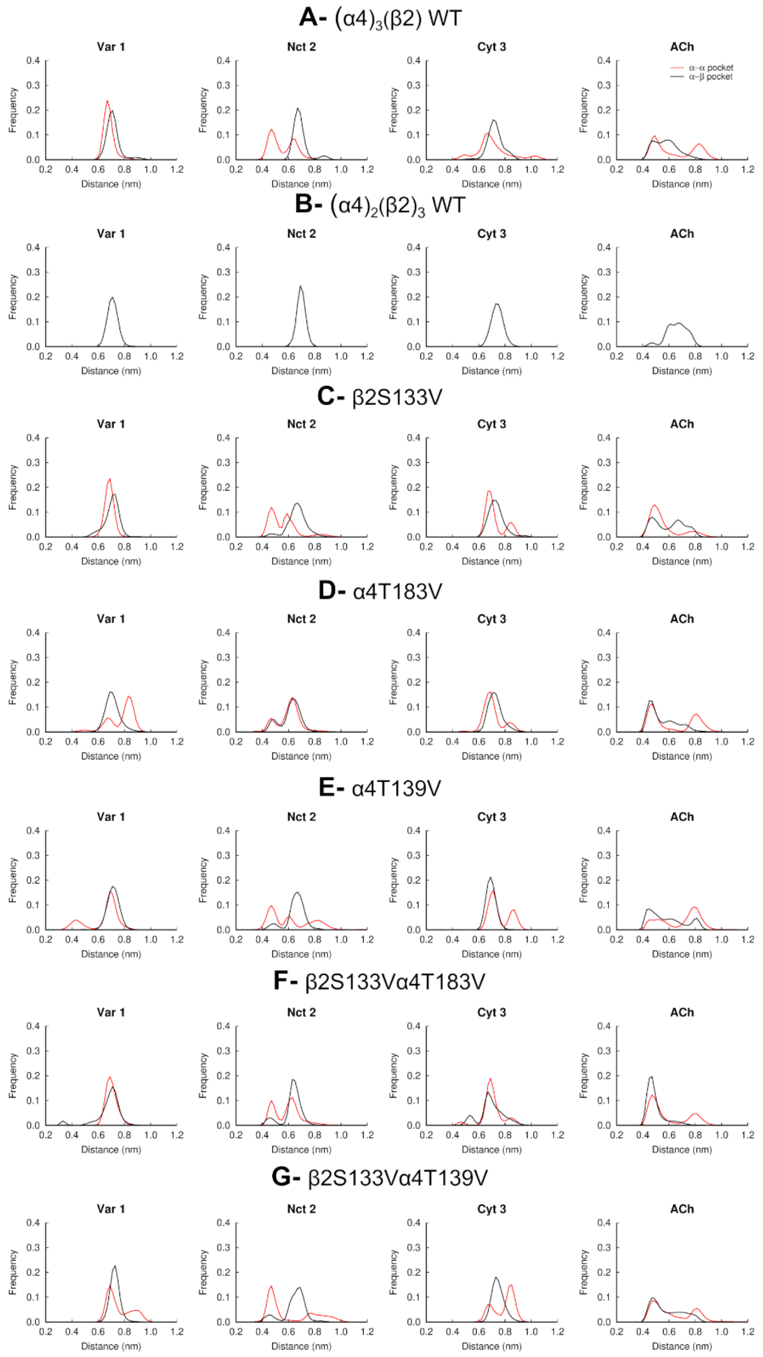


**Figure S20**- TrpD-agonist distance for the LS $(\alpha4)_3(\beta2)_2$ (**A**) and HS $(\alpha4)_2(\beta2)_3$ (**B**) wild-type and mutant (**C-G**) systems. The distance between the side-chain of TrpD (W88 in the complementary α4 subunit of the α-α binding pocket and W82 in the complementary β2 subunit of the α-β binding pocket) and the protonated nitrogen atom of varenicline **1**, nicotine **2**, cytisine **3** and ACh for the α-α (red line) and α-β (black line) binding pockets. The histogram for the α-β pocket reflects the distances over the two α-β binding pockets present in both the LS and HS isoforms of the α4β2 nAChR. Please note that the histogram for the α-β binding pocket of β2S133Vα4T139V-ACh complex excludes the data for the second α-β pocket from 112 ns onwards for replicate 3, as the agonist exits this binding pocket in the simulation after 112 ns.

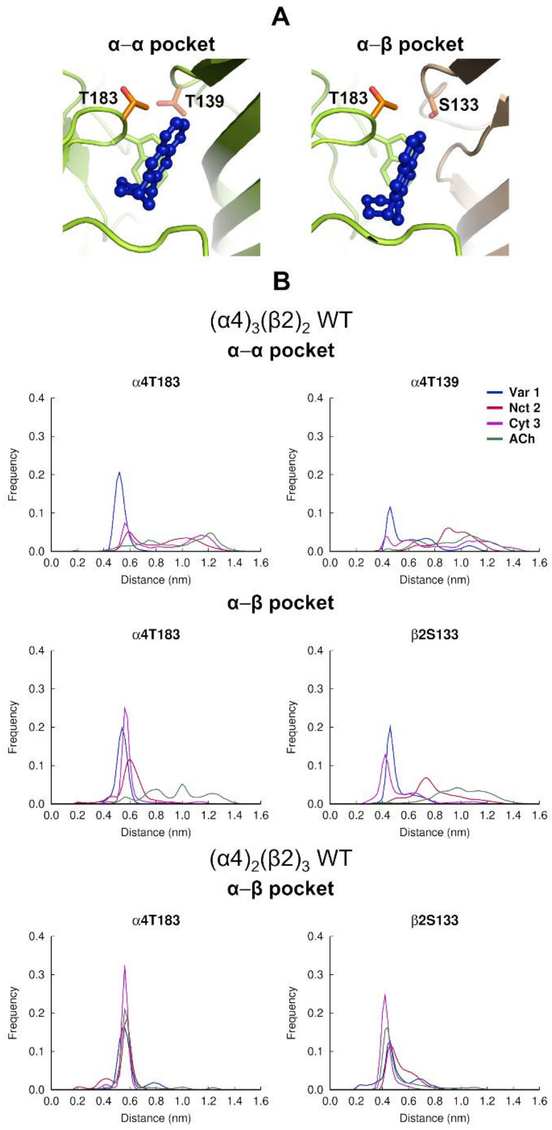


**Figure S21-** Agonist interactions with α4T183, α4T139 and β2S133 in the LS $(\alpha4)_3(\beta2)_2$ and HS $(\alpha4)_2(\beta2)_3$ wild-type systems. (**A**) Location of α4T183 and α4T139 in the α-α pocket (left panel) and α4T183 and β2S133 in the α-β (right panel) binding site. The α4 and β2 subunits are colored in yellow and light brown, respectively. Varenicline **1** is highlighted in dark blue. The side-chain of α4T183, α4T139 and β2S133 are represented with orange sticks, whereas TrpB is shown with yellow sticks. Note that α4T183 is located in the principal α4 face of the pockets whereas α4T139 and β2S133 are in the complementary face of the α-α and α-β pockets, respectively. (**B**) Distribution of the minimum distance between the agonist and α4T183, α4T139 and β2S133 in the α-α and α-β binding pockets of the LS $(\alpha4)_3(\beta2)_2$ and HS $(\alpha4)_2(\beta2)_3$ receptor. The reported values correspond to the minimum distances between the agonist (specifically, the closest pyrazine nitrogen in the quinoxaline moiety of varenicline **1**, the pyridine nitrogen of nicotine **2,** the pyridone carbonyl oxygen of cytisine **3** and the closest oxygen in the ester group of ACh), and the hydroxyl group of α4T183, α4T139 and β2S133 in all the MD trajectories for each complex. The histogram for the α-β pocket reflects the distances over the two α-β binding pockets present in the LS and HS isoforms of the α4β2 nAChR. These distance profiles indicate that some agonists can closely approach the H-bond donors in the side-chain of α4T183, α4T139, and β2S133, thus suggesting the possibility of transient interactions occurring between them.

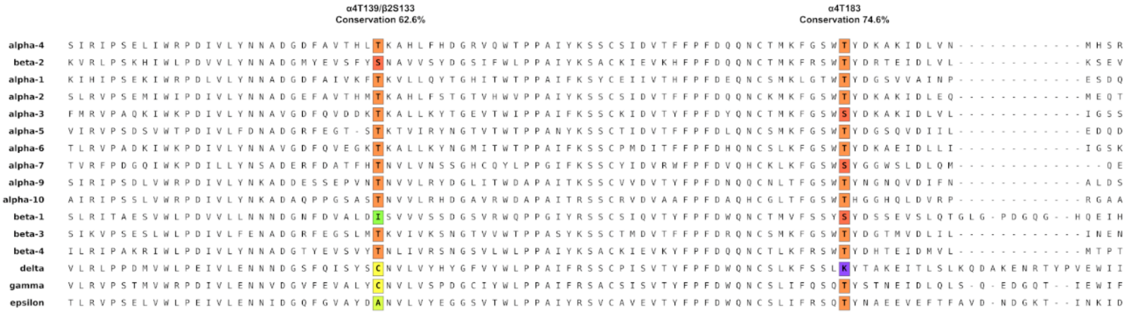


**Figure S22-** Sequence alignment for the human α1-α7, α9-α10, β1-β4, δ, γ, and ε nAChR subunits. The sequence alignments were performed using the Muscle server.[25] The colored boxes highlight the locations of α4T183, α4T139 and β2S133, with threonine, serine, isoleucine, cysteine, alanine, and lysine residues represented by orange, red, green, yellow, light green, and purple, respectively. Conservation percentages showed are expressed as 100×(1−H/$H_{max}$), where H is the Shannon entropy of the residue at the alignment position.[26] The sequences shown correspond to the following UniProt codes: P43681 (human α4), P17787 (human β2), P02708 (human α1), Q15822 (human α2), P32297 (human α3), P30532 (human α5), Q15825 (human α6), P36544 (human α7), Q9U6MI (human α9), Q9GZZ6 (human α10), P11230 (human β1), Q05901 (human β3), Q07001 (human δ), P07510 (human γ), and Q04844 (human ε).

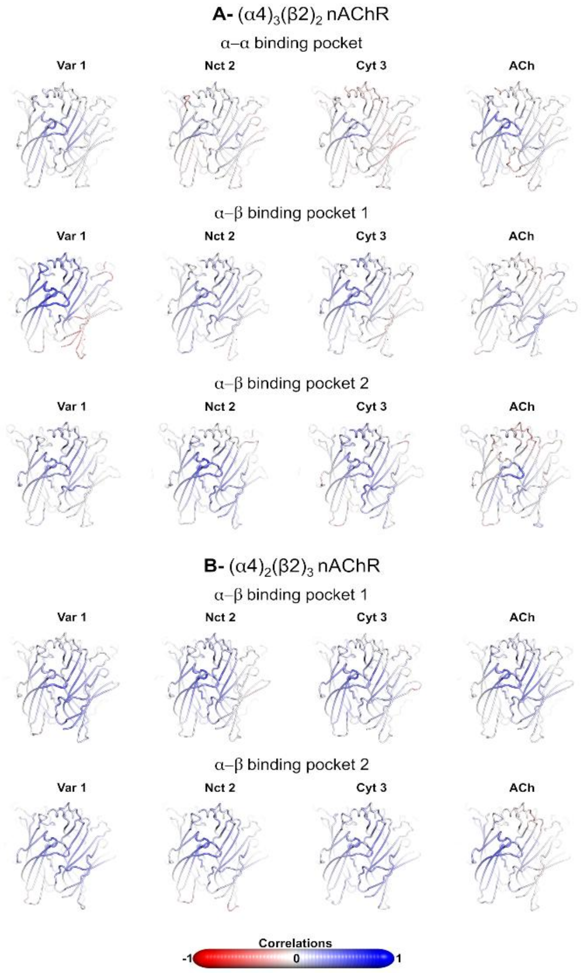


**Figure S23-** Statistical correlations for the different agonists when bound to ECD of the (**A**) LS $(\alpha4)_3(\beta2)_2$ and (**B**) HS $(\alpha4)_2(\beta2)_3$ nAChRs. Correlated motions for the agonist in the α-α and α-β binding pockets of the wild-type system. The correlations between the protonated nitrogen atom of the agonists and all the Cα atoms in the receptor are shown. Note that the atoms that systematically move along the opposite direction as the agonist have a correlation value of -1 whereas those systematically moving along the same direction show a correlation of 1. The atoms whose movements relative to the agonist are uncorrelated present a correlation value of 0. Please zoom in on the image for detailed visualization.

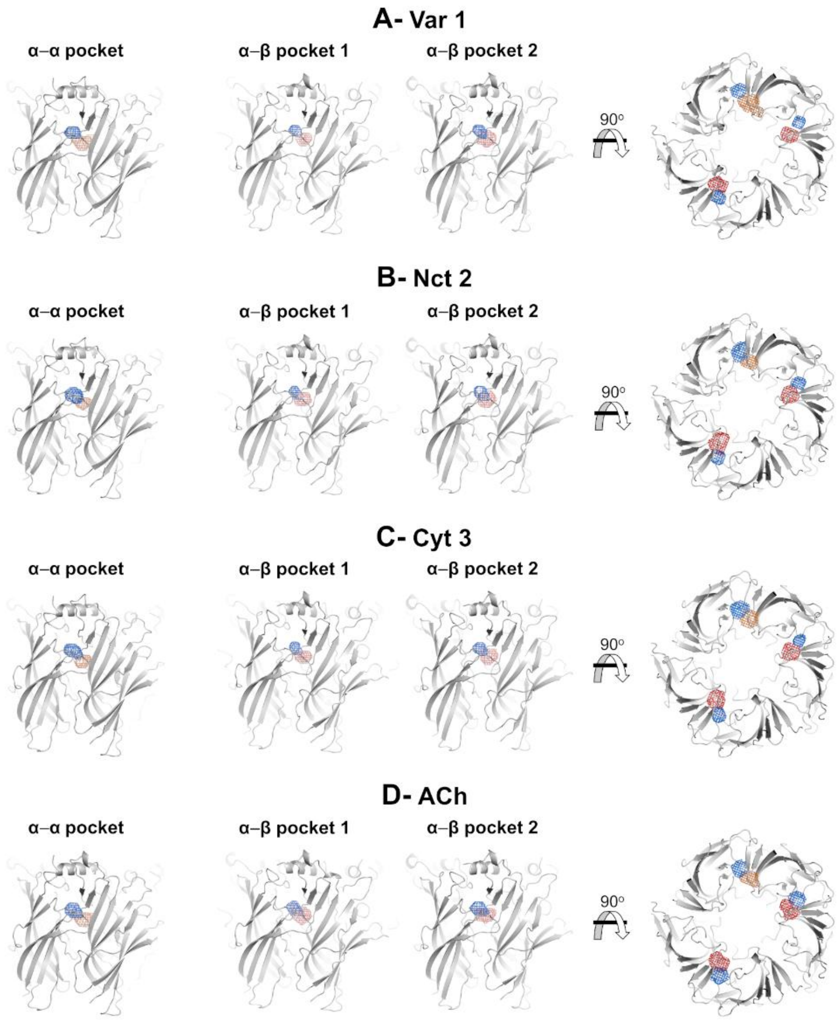


**Figure S24-** Probability density maps for the hydroxyl group of α4T183 (principal α4 side), α4T139 (complementary α4 side) and β2S133 (complementary β2 side) in the α-α and α-β binding pockets for the $(\alpha4)_3(\beta2)_2$ wild-type simulations. (**A-D**) Probability density maps for the varenicline **1**, nicotine **2**, cytisine **3** and ACh bound simulations. The oxygen atom in the hydroxyl group of α4T183, α4T139 and β2S133 were used to create the probability density maps. The contours at 0.00001 $Å^{-3}$ for α4T183, α4T139 and β2S133 are depicted as a blue, orange and red mesh, respectively. No noticeable differences are observed between the maps for the two α-β binding sites. However, in the α-α binding pocket, the side-chain of α4T183 and α4T139 exhibit greater mobility, more effectively exploring the binding site. Please zoom in on the image for detailed visualization.

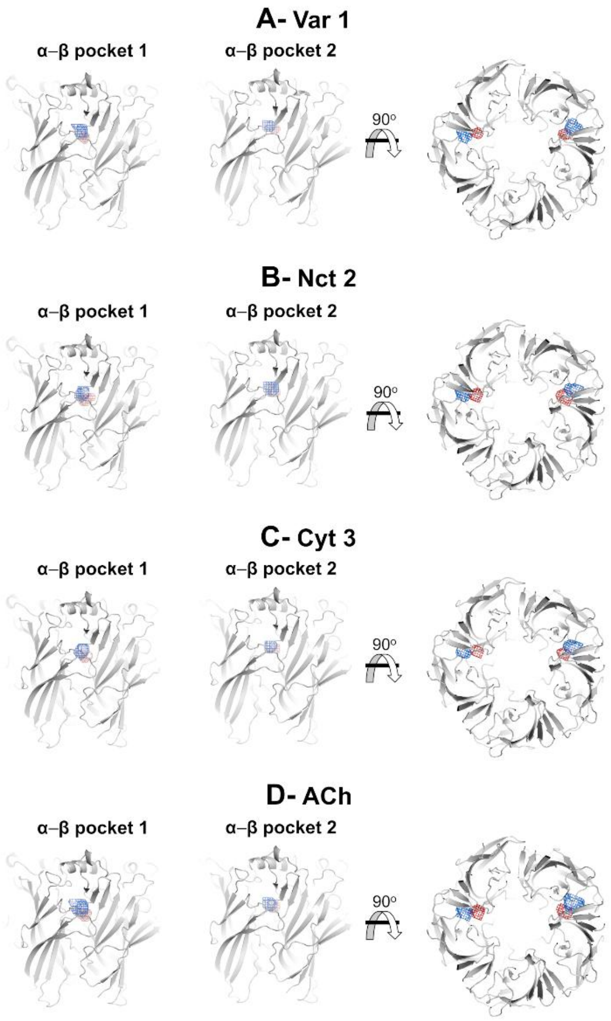


**Figure S25-** Probability density maps for the hydroxyl group of α4T183 (principal α4 side) and β2S133 (complementary β2 side) in the α-β binding pockets for the $(\alpha4)_2(\beta2)_3$ wild-type simulations. (**A-D**) Probability density maps for the varenicline **1**, nicotine **2**, cytisine **3** and ACh bound simulations. The oxygen atom in the hydroxyl group of α4T183 and β2S133 were used to create the probability density maps. The contours at 0.00001 $Å^{-3}$ for α4T183 and β2S133 are depicted as a blue and red mesh, respectively. No noticeable differences are observed between the maps for the two α-β binding sites.

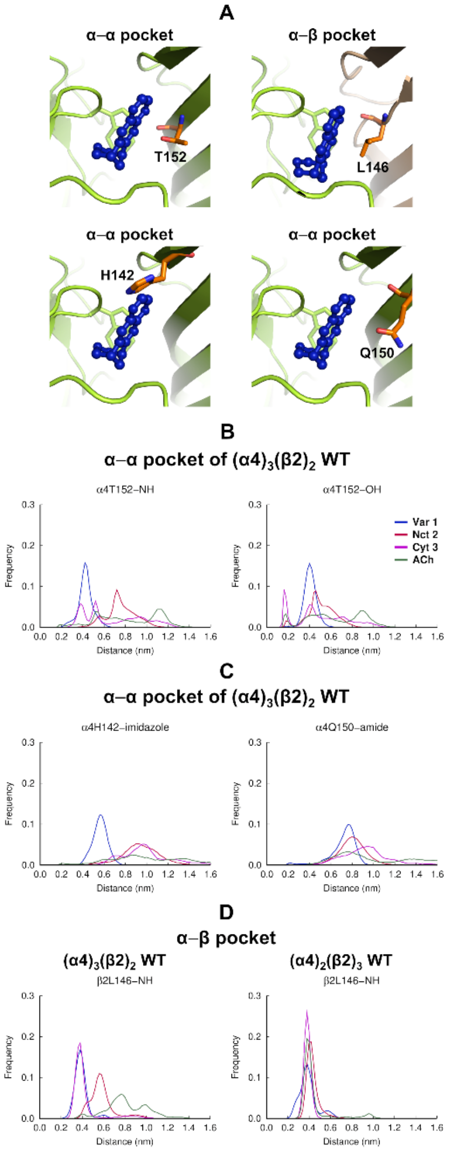


**Figure S26-** Interactions between the agonists and α4H142, α4Q150, α4T152, and β2L146 in the LS $(\alpha4)_3(\beta2)_2$ and HS $(\alpha4)_2(\beta2)_3$ wild-type systems. (**A**) Location of α4H142, α4Q150 and α4T152 in the α-α pocket and β2L146 in the α-β pocket. The α4 and β2 subunits are colored in yellow and light brown, respectively. Varenicline **1** is highlighted in dark blue. α4H142, α4Q150, α4T152 and β2L146 are represented with orange sticks while TrpB is shown with yellow sticks. Note that α4H142, α4Q150 and α4T152 are located in the complementary face of the α-α pocket whereas and β2L146 is situated in the complementary side of the α-β pockets. (**B**) Distribution of the minimum distance between the agonist and the backbone NH and side-chain hydroxyl group of α4T152 in the α-α pocket of the LS $(\alpha4)_3(\beta2)_2$ isoform. The values reported correspond to the minimum distance between the agonist (namely, the closest pyrazine nitrogen in the quinoxaline group of varenicline **1**, the pyridine nitrogen of nicotine **2**, the pyridone carbonyl oxygen of cytisine **3**, and the closest oxygen in the ester group of ACh) and the NH and OH group of α4T152 in all the MD trajectories for each complex. (**C**) Distribution of the minimum distance between the agonist and the H-bond donors in the side-chain of α4H142 and α4Q150 in the α-α pocket of the LS $(\alpha4)_3(\beta2)_2$ isoform. The values reported correspond to the minimum distance between the agonist (namely, the closest pyrazine nitrogen in the quinoxaline group of varenicline **1**, the pyridine nitrogen of nicotine **2**, the pyridone carbonyl oxygen of cytisine **3**, and the closest oxygen in the ester group of ACh) and the imidazole NH and amide NH2 group in the side-chain of α4H142 and α4Q150 in all the MD trajectories for each complex. (**D**) Distribution of the minimum distance between the agonist and the backbone NH of β2L146 in the α-β pockets of the LS $(\alpha4)_3(\beta2)_2$ (left panel) and HS $(\alpha4)_2(\beta2)_3$ (right panel) receptor. The values reported correspond to the minimum distance between the agonist (namely, the closest pyrazine nitrogen in the quinoxaline group of varenicline **1**, the pyridine nitrogen of nicotine **2**, the pyridone carbonyl oxygen of cytisine **3**, and the closest oxygen in the ester group of ACh) and the NH group of β2L146 in all the MD trajectories for each complex. The histograms reflect the distances over the two α-β binding pockets present in the LS and HS isoforms of the α4β2 nAChR. Note that the distance profiles above indicate that some agonists can directly interact with the α4T152 hydroxyl donor in the α-α pocket.

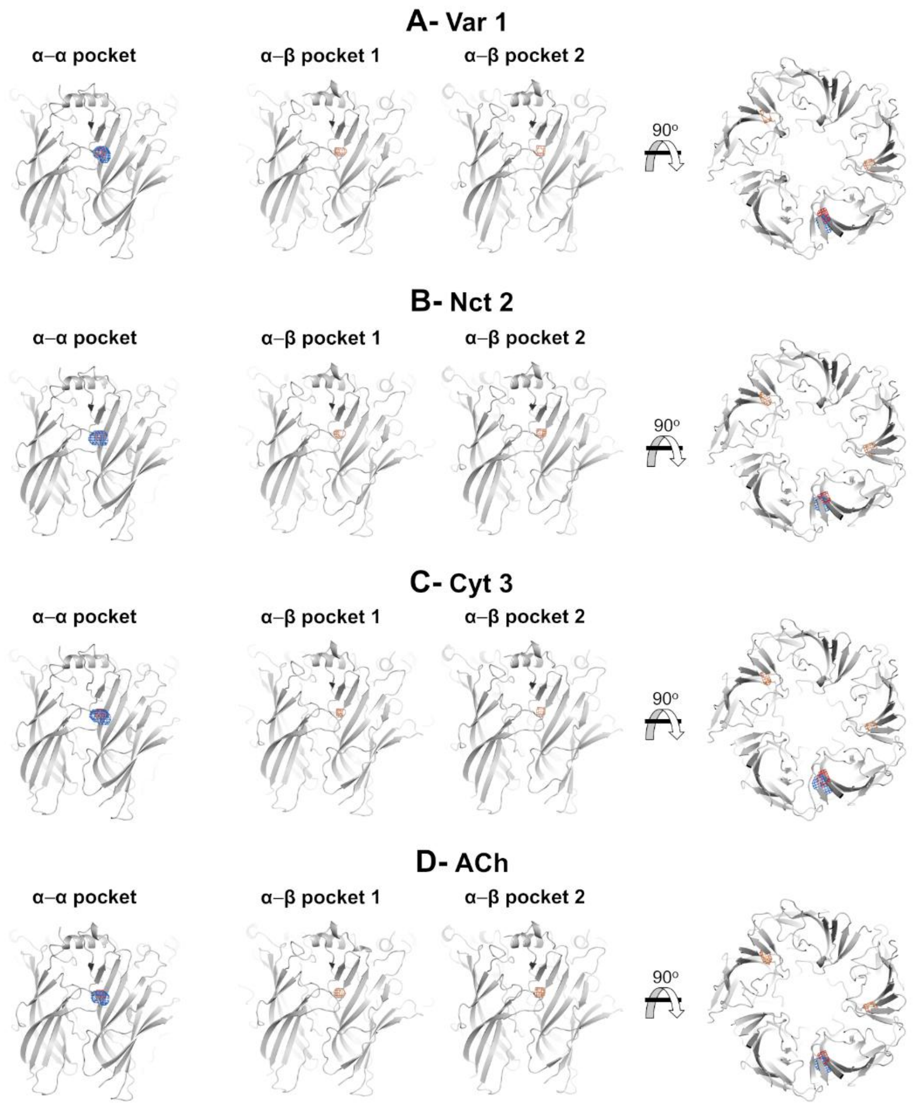


**Figure S27-** Probability density maps for the side-chain OH of T152 (complementary α4 side) and backbone NH group of T152 and L146 (complementary β2 side) in the α-α and α-β binding pockets for the LS $(\alpha4)_3(\beta2)_2$ wild-type simulations with varenicline **1** (**A**), nicotine **2** (**B**), cytisine **3** (**C**), and acetylcholine (**D**). The hydrogen atom in the backbone NH group of T152 and L146 and the oxygen atom in the OH group of T152 were used to create the probability maps. The contours at 0.00001 Å$^{-3}$ for NH and OH of T152 and NH of L146 are depicted as red, blue and orange meshes, respectively. Note that no noticeable differences are observed between maps across the various agonist-receptor complexes. Please zoom in on the image for detailed visualization.

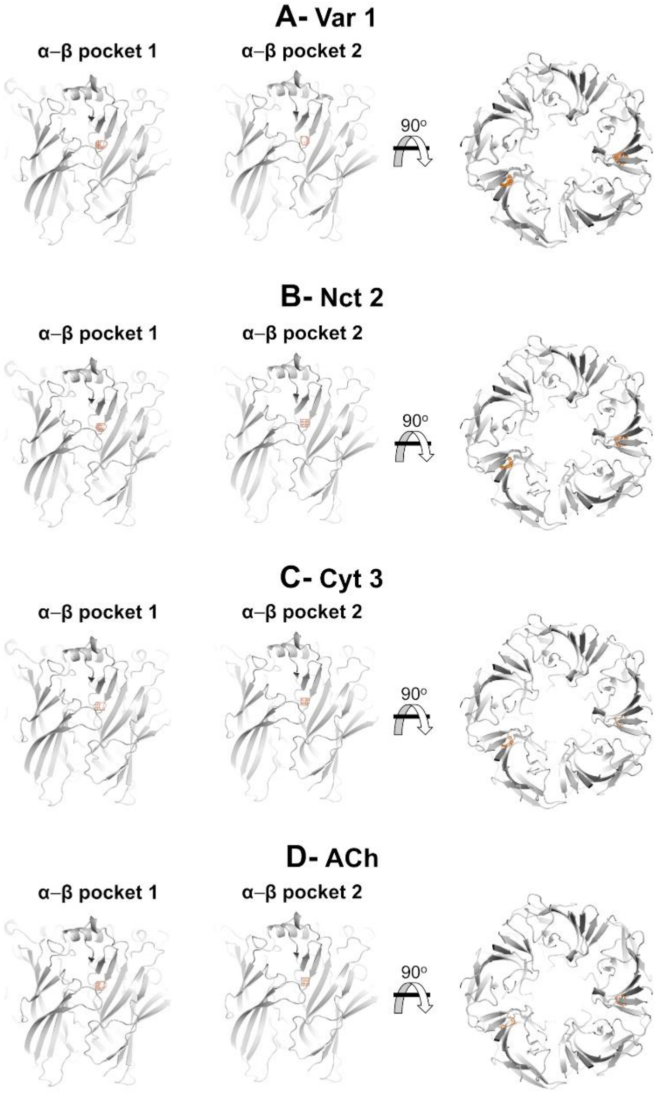


**Figure S28-** Probability density maps for the backbone NH group of L146 (complementary β2 side) in the α-β binding pockets for the HS $(\alpha4)_2(\beta2)_3$ wild-type simulations with varenicline **1** (**A**), nicotine **2** (**B**), cytisine **3** (**C**), and acetylcholine (**D**). The hydrogen atom in the backbone NH group of L146 were used to create the probability maps. The contours at 0.00001 $Å^{-3}$ for NH of L146 are depicted as orange meshes. Note that no noticeable differences are observed between maps across the various agonist-receptor complexes. Please zoom in on the image for detailed visualization.

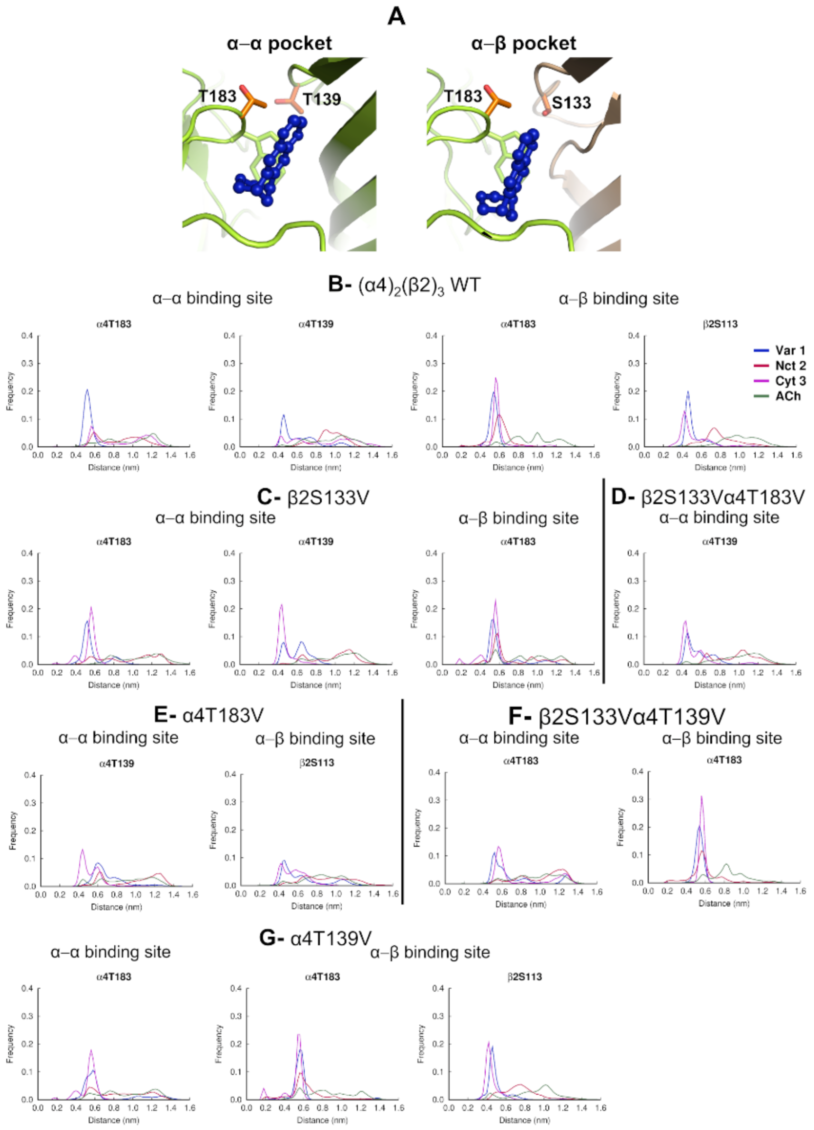


**Figure S29-** Agonist interactions with α4T183, α4T139 and β2S133 in the LS $(\alpha4)_3(\beta2)_2$ wild-type and mutant systems. (**A**) Location of α4T183, α4T139 and β2S133 in the α-α (left panel) and α-β (right panel) binding pockets of the LS $(\alpha4)_3(\beta2)_2$ wild-type receptor. The α4 and β2 subunits are colored in yellow and light brown, respectively. Varenicline **1** is highlighted in dark blue. The side-chain of α4T183, α4T139 and β2S133 are represented with orange sticks whereas TrpB is shown with yellow sticks. (**B-G**) Distribution of the minimum distance between the agonist and α4T183, α4T139 and β2S133 in the α-α and α-β binding pockets of the wild-type and β2S133V, α4T183V, α4T139V, β2S133Vα4T183V and β2S133Vα4T139V mutants. The reported values correspond to the minimum distances between the agonist (specifically, the closest pyrazine nitrogen in the quinoxaline group of varenicline **1**, the pyridine nitrogen of nicotine **2**, the pyridone carbonyl oxygen of cytisine **3**, and the closest oxygen in the ester group of ACh) and the hydroxyl group of α4T183, α4T139 and β2S133 in the MD trajectories for each complex. The histogram for the α-β pocket reflects the distances over the two α-β binding pockets present in the LS isoform of the α4β2 nAChR. Note that the substitution of α4T183, α4T139, and β2S133 by valine (which lacks the hydroxyl group in its side-chain) prevents potential hydrogen bonding with the agonists. The distance profiles indicate that, as anticipated, the mutations altered the interaction patterns between the H-bond donors in the receptor binding sites and the agonists acceptor groups.

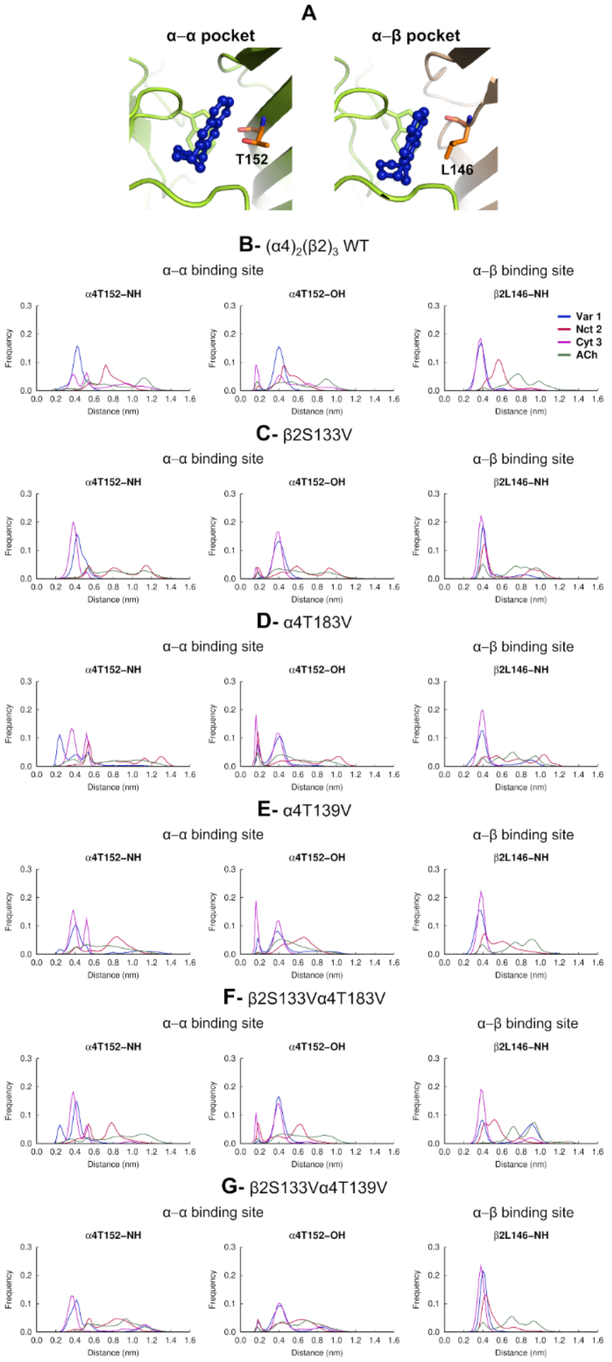


**Figure S30-** Agonist interactions with α4T152 and β2L146 in the LS $(\alpha4)_3(\beta2)_2$ wild-type and mutant systems. (**A**) Location of α4T152 and β2L146 in the α-α (left panel) and α-β (right panel) binding pockets of the $(\alpha4)_3(\beta2)_2$ wild type receptor. The α4 and β2 subunits are colored in yellow and light brown, respectively. Varenicline **1** is highlighted in dark blue. The side-chain of α4T152 and β2L146 are represented with orange sticks, whereas TrpB is shown with yellow sticks. (**B-G**) Distribution of the minimum distance between the agonist and α4T152 and β2L146 in the α-α and α-β binding pockets of the wild-type (**B**) and β2S133V (**C**), α4T183V (**D**), α4T139V (**E**), β2S133Vα4T183V (**F**) and β2S133Vα4T139V (**G**) mutants. The reported values correspond to the minimum distances between the agonist (specifically, the closest pyrazine nitrogen in the quinoxaline group of varenicline **1**, the pyridine nitrogen of nicotine **2**, the pyridone carbonyl oxygen of cytisine **3**, and the closest oxygen in the ester group of ACh) and the backbone NH of α4T152 and β2L146 and the side-chain OH of α4T152 in the MD trajectories for each complex. The histogram for the α-β pocket reflects the distances over the two α-β binding pockets present in the LS isoform of the α4β2 nAChR. Overall, the distance profiles indicate that the mutations mainly altered the interaction pattern between the agonists and α4T152 within the α-α pocket.

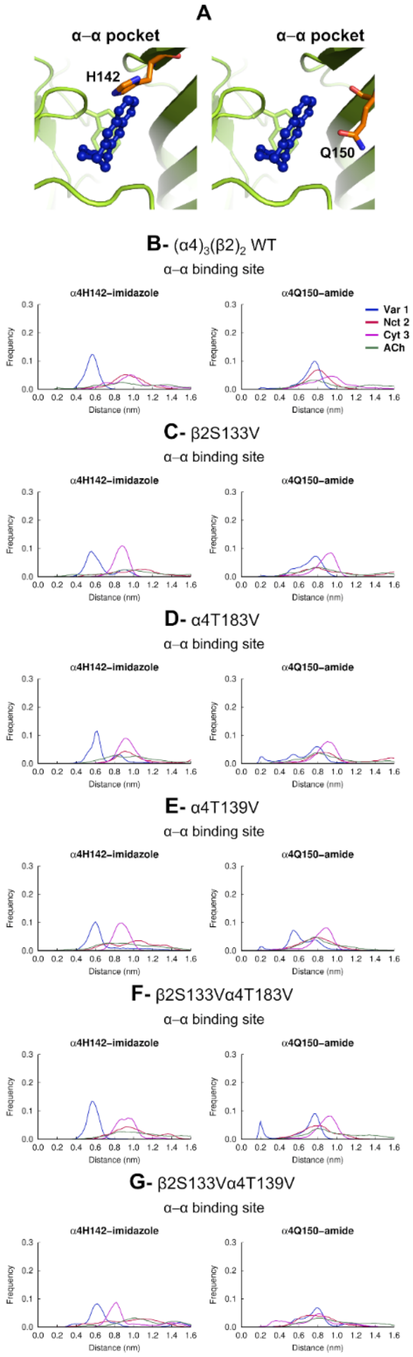


**Figure S31-** Agonist interactions with α4H142 and α4Q150 in the LS $(\alpha4)_3(\beta2)_2$ wild-type and mutant systems. (**A**) Location of α4H142 (left panel) and α4Q150 (right panel) in the α-α binding pocket of the LS $(\alpha4)_3(\beta2)_2$ wild-type receptor. The α4 and β2 subunits are colored in yellow and light brown, respectively. Varenicline **1** is highlighted in dark blue. The side-chain of α4H142 and α4Q150 are represented with orange sticks, whereas TrpB is shown with yellow sticks. (**B-G**) Distribution of the minimum distance between the agonists and α4H142 and α4Q150 in the α-α binding pocket of the wild-type (**B**) and β2S133V (**C**), α4T183V (**D**), α4T139V (**E**), β2S133Vα4T183V (**F**) and β2S133Vα4T139V (**G**) mutants. The reported values correspond to the minimum distances between the agonist (specifically, the closest pyrazine nitrogen in the quinoxaline group of varenicline **1**, the pyridine nitrogen of nicotine **2**, the pyridone carbonyl oxygen of cytisine **3**, and the closest oxygen in the ester group of ACh) and the NH in the imidazole side-chain of α4H142 and the side-chain NH2 amide of α4Q150 in the MD trajectories for each complex. Overall, the distance profiles indicate that the mutations mainly altered the interaction patterns between the agonists and α4Q150 within the α-α pocket.

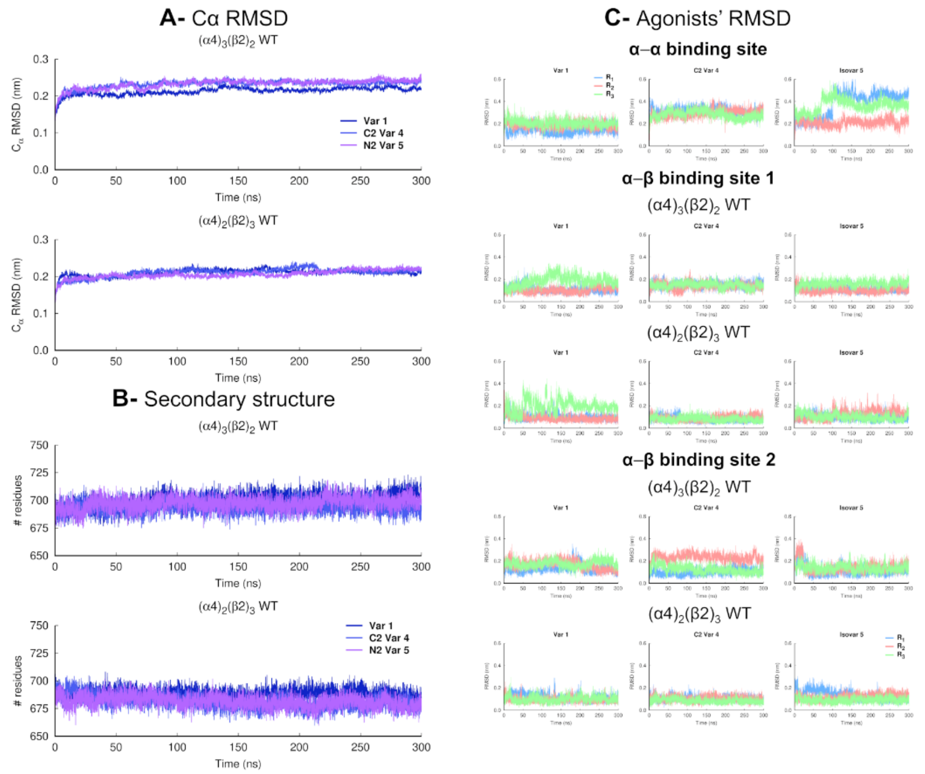


**Figure S32-** (**A**) Temporal evolution of the average Cα RMSD for the varenicline **1,** $C_2$ varenicline **4** and isovarenicline **5** bound to LS $(\alpha4)_3(\beta2)_2$ and HS $(\alpha4)_3(\beta2)_2$ wild-type systems. The Cα RMSD was calculated relative to the starting structures, and the averages were obtained over all replicates for each system. (**B**) Time evolution of number of residues involved in secondary structure motifs for the varenicline **1**, $C_2$ varenicline **4** and isovarenicline **5** bound systems. The secondary structure assignment was performed with the DSSP software[22] and includes all residues assigned to α-helix, π-helix, $3_{10}$-helix, 5-helix, β-sheet, β-strand and β-bridge secondary structure classes. The averages were obtained over all replicates for each system. (**C**) Time evolution of the RMSD for varenicline **1,** $C_2$ varenicline **4** and isovarenicline **5** when bound to the α-α and α-α binding pockets of the LS $(\alpha4)_3(\beta2)_2$ and HS $(\alpha4)_2(\beta2)_3$ wild-type system. The RMSD was determined with respect to the initial binding mode of the agonists at the start of the simulations. Note that the α-α binding pocket only exists in the LS $(\alpha4)_3(\beta2)_2$ isoform of the receptor. Please zoom in on the image for detailed visualization.

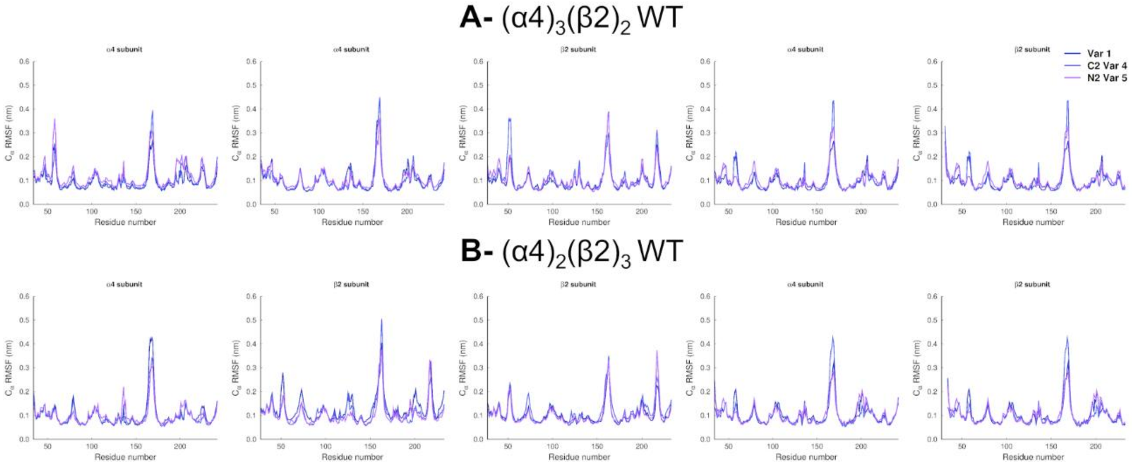


**Figure S33-** Average Cα RMSF for the LS $(\alpha4)_3(\beta2)_2$ (**A**) and HS $(\alpha4)_2(\beta2)_3$ **(B)** wild-type receptors with varenicline **1**, $C_2$ varenicline **4** and isovarenicline **5** bound. The Cα RMSF was calculated using the entire trajectories and averaged across all replicates for each complex. Please zoom in on the image for detailed visualization.

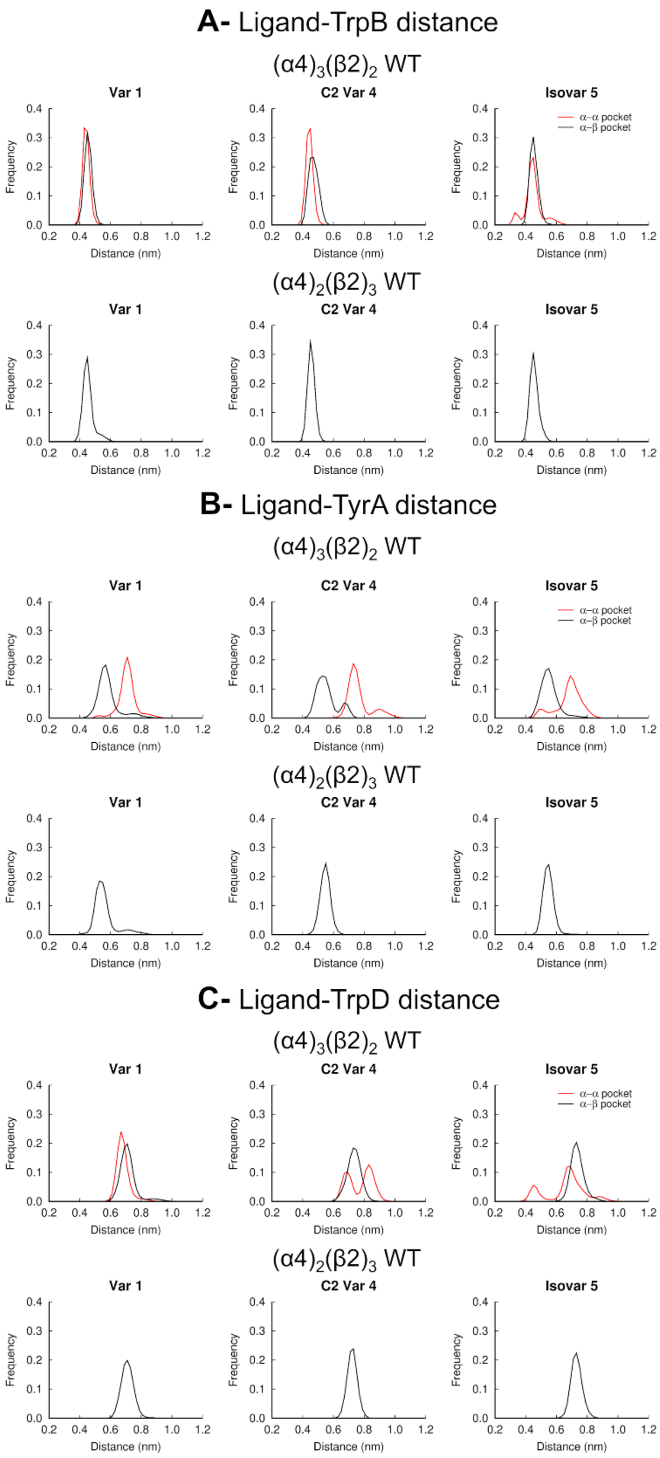


**Figure S34**- Distance profiles between varenicline **1**, $C_2$ varenicline **4** and isovarenicline **5** and TrpB **(A)**, TyrA (**B)** and TrpD (**C**) for the LS $(\alpha4)_3(\beta2)_2$ and HS $(\alpha4)_2(\beta2)_3$ wild-type systems. Distance between the side chain of TrpB (W182 in the principal α4 subunit), TyrA (Y126 in the principal α4 subunit) and TrpD (W88 in the complementary α4 subunit of the α-α binding pocket and W82 in the complementary β2 subunit of the α-β binding pocket) and the protonated (piperidine) nitrogen atom of varenicline**1**, $C_2$ varenicline **4** and isovarenicline **5** for the α-α (red line) and α-β (black line) binding pockets. The histogram for the α-β pocket reflects the distances over the two α-β binding pockets present in the LS $(\alpha4)_3(\beta2)_2$ and HS $(\alpha4)_2(\beta2)_3$ nAChRs.

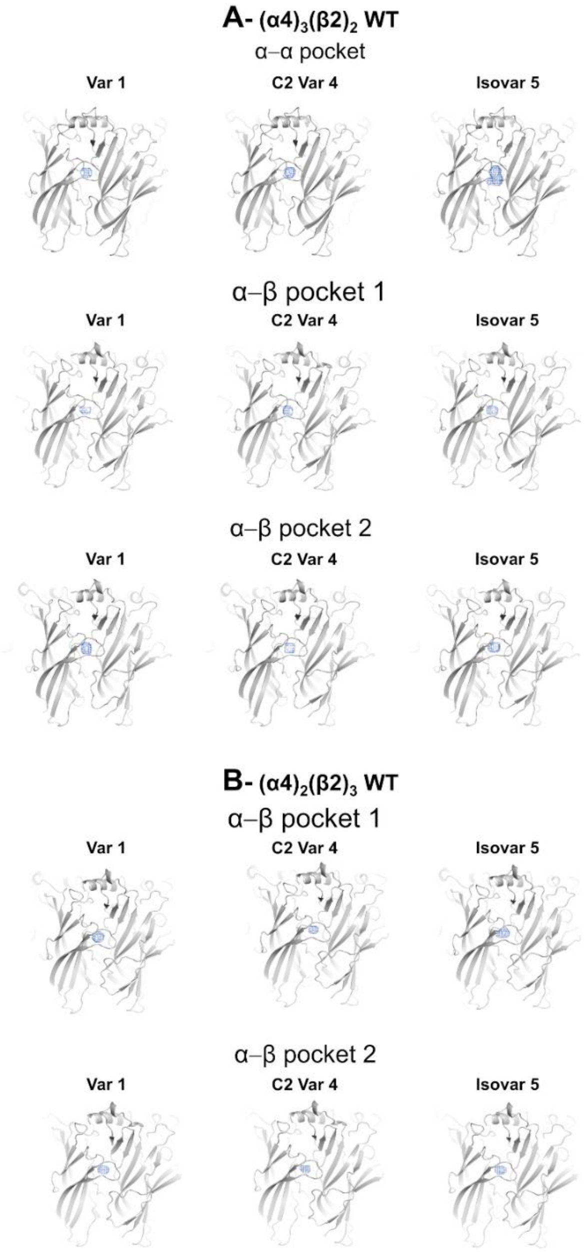


**Figure S35-** Probability density maps for varenicline **1**, $C_2$ varenicline **4** and isovarenicline **5** bound to the α-α and α-β binding pocket in the LS $(\alpha4)_3(\beta2)_2$ (**A**) and HS $(\alpha4)_2(\beta2)_3$ (**B**) wild-type simulations. The contours at 0.00001 $Å^{-3}$ for the protonated nitrogen atom of the agonists are depicted as a blue mesh. The maps were calculated combining the entire trajectories (0-300 ns) for each one of the three replicates of that system. Please zoom in on the image for detailed visualization.

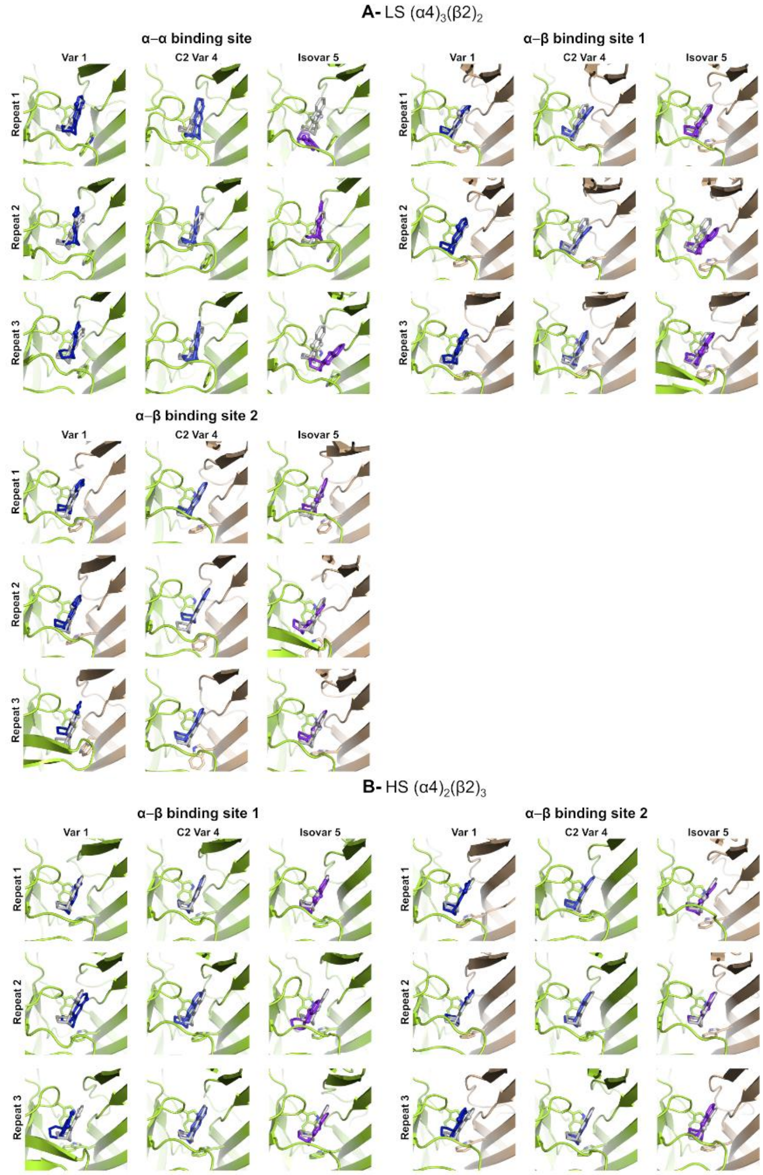


**Figure S36**- Binding mode of varenicline **1,** $C_2$ varenicline **4**, and isovarenicline **5** after 300 ns for the wild-type α4β2 complexes. (**A**) Binding mode of varenicline **1**, $C_2$ varenicline **4** and isovarenicline **5** in the α-α and α-β pockets of the wild-type LS $(\alpha4)_3(\beta2)_2$ nAChR. (**B**) Binding mode of varenicline **1**, $C_2$ varenicline **4** and isovarenicline **5** in the α-β pockets of the wild-type HS $(\alpha4)_2(\beta2)_3$ nAChR. The α4 and β2 subunits are colored in yellow and light brown, respectively. Agonists are depicted in blue, with the nitrogen atoms of the quinoxaline moiety (which serve as H-bond acceptors) highlighted by spheres. Please note that $C_2$ varenicline **4** contains a naphthalene residue instead of a quinoxaline unit, and thus lacks the nitrogen atoms needed to form hydrogen bonds. The grey sticks represent the starting binding mode for the agonists. TrpB and TrpD are shown with sticks. Please zoom in on the image for detailed visualization.

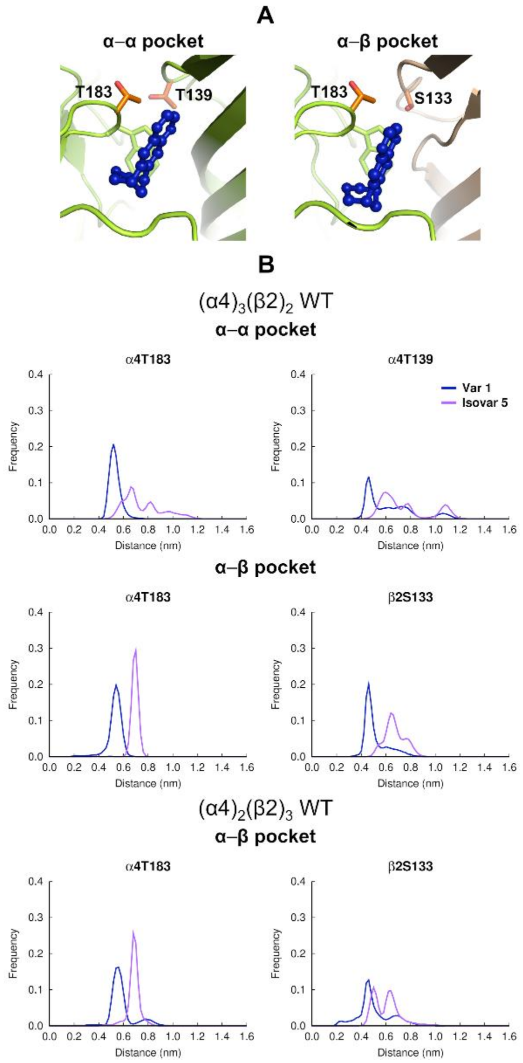


**Figure S37-** Varenicline **1** and isovarenicline **5** interactions with α4T183, α4T139 and β2S133 in the LS $(\alpha4)_3(\beta2)_2$ and HS $(\alpha4)_2(\beta2)_3$ wild-type systems. (**A**) Location of α4T183, α4T139 and β2S133 in the α-α (left panel) and α-β (right panel) binding pockets. The α4 and β2 subunits are colored in yellow and light brown, respectively. Varenicline **1** is highlighted in dark blue. The side-chain of α4T183, α4T139 and β2S133 are represented with orange sticks, whereas TrpB is shown with yellow sticks. (**B**) Distribution of the minimum distance between the closest pyrazine nitrogen in the quinoxaline group of varenicline **1** and isovarenicline **5** and α4T183, α4T139 and β2S133 in the α-α and α-β binding pockets of the $(\alpha4)_3(\beta2)_2$ and $(\alpha4)_2(\beta2)_3$ receptor in all the MD trajectories for each complex. Note that $C_2$ varenicline **4**, instead of a quinoxaline unit, possesses a naphthalene residue, and is therefore unable to form hydrogen bonds. The histogram for the α-β pocket reflects the distances over the two α-β binding pockets present in the LS and HS of the α4β2 nAChR. The distance profiles clearly demonstrate that, as expected, the distance between the hydrogen acceptor group in isovarenicline **5** and the residues side-chain OH group is too large, thereby preventing any direct interaction between the two.

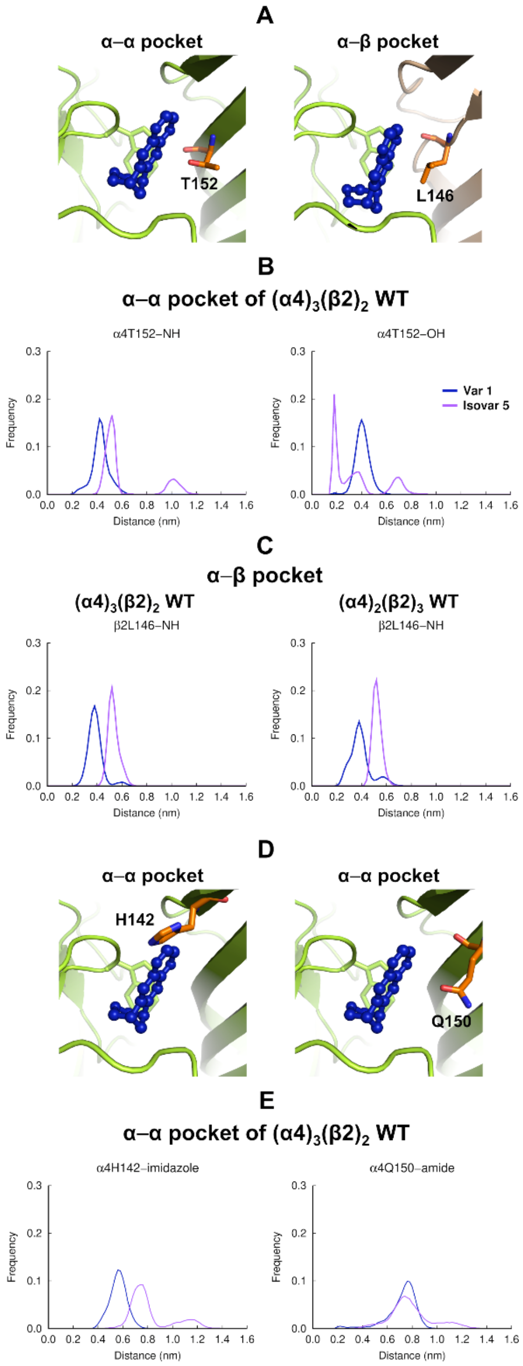


**Figure S38-** Varenicline **1** and isovarenicline **5** interactions with α4T152 and β2L146 in the LS $(\alpha4)_3(\beta2)_2$ and HS $(\alpha4)_2(\beta2)_3$ wild-type systems. (**A**) Location of α4T152 and β2L146 in the α-α (left panel) and α-β (right panel) binding pockets of the $(\alpha4)_3(\beta2)_2$ wild-type receptor. See caption of Figure S37 for more details. (**B**) Distribution of the minimum distance between the closest pyrazine nitrogen in the quinoxaline group of varenicline **1** and isovarenicline **5** and the backbone NH and side-chain OH of α4T152 in the α-α pocket of the LS isoform of α4β2. Please note that **$C_2$** varenicline **4**, instead of a quinoxaline unit, possesses a naphthalene residue, and is therefore unable to form hydrogen bonds. (**C**) Distribution of the minimum distance between the closest pyrazine nitrogen in the quinoxaline group of varenicline **1** and isovarenicline **5** and the backbone NH of β2L146 in the α-β pockets of the $(\alpha4)_3(\beta2)_2$ (left panel) and $(\alpha4)_2(\beta2)_3$ (right panel) receptor. The histograms reflect the distances over the two α-β binding pockets present in the LS and HS isoforms of the α4β2 nAChR. Surprisingly, the distance profiles above show that isovarenicline **5** can make a persistent interaction with the α4T152 hydroxyl donor in the α-α pocket. (**D**) Location of α4H142 (left panel) and α4Q150 (right panel) in the α-α binding pockets of the LS $(\alpha4)_3(\beta2)_2$ wild-type receptor. See caption of Figure S37 for more details. (**E**) Distribution of the minimum distance between the closest pyrazine nitrogen in the quinoxaline group of varenicline **1** and isovarenicline **5** and the NH group in the imidazole side-chain of α4H142 (left side) and the NH2 in the amide side-chain of α4Q150 (right side) in the α-α pocket of LS $(\alpha4)_3(\beta2)_2$. The distance profiles show isovarenicline **5** is unable to directly H-bond to α4H142 and α4Q150.

**Table S1-** Summary of the simulations performed for the ECD of the human α4β2 nAChR. *The simulations for the complexes between HS $(\alpha4)_2(\beta2)_3$ receptor with nicotine **2**, cytisine **3** and ACh were taken from our previous work.[7, 8]

Abbreviations used: varenicline **1** - **Var 1**; nicotine **2** - **Nct 2**; cytisine **3** - **Cyt 3**; acetylcholine - **ACh**; $C_2$ varenicline **4** - **C2 Var 4**; isovarenicline **5** - **Isovar 5**.

| | | System | Agonist | Simulation length (ns) | Number of replicates |
|---|---|---|---|---|---|
| 1 | $(\alpha4)_3(\beta2)_2$ | Wild type | **Var 1** | 300 | 3 |
| 2 | $(\alpha4)_3(\beta2)_2$ | Wild type | **Nct 2** | 300 | 3 |
| 3 | $(\alpha4)_3(\beta2)_2$ | Wild type | **Cyt 3** | 300 | 3 |
| 4 | $(\alpha4)_3(\beta2)_2$ | Wild type | **ACh** | 300 | 3 |
| 5 | $(\alpha4)_3(\beta2)_2$ | β2S133V | **Var 1** | 300 | 3 |
| 6 | $(\alpha4)_3(\beta2)_2$ | β2S133V | **Nct 2** | 300 | 3 |
| 7 | $(\alpha4)_3(\beta2)_2$ | β2S133V | **Cyt 3** | 300 | 3 |
| 8 | $(\alpha4)_3(\beta2)_2$ | β2S133V | **ACh** | 300 | 3 |
| 9 | $(\alpha4)_3(\beta2)_2$ | α4T183V | **Var 1** | 300 | 3 |
| 10 | $(\alpha4)_3(\beta2)_2$ | α4T183V | **Nct 2** | 300 | 3 |
| 11 | $(\alpha4)_3(\beta2)_2$ | α4T183V | **Cyt 3** | 300 | 3 |
| 12 | $(\alpha4)_3(\beta2)_2$ | α4T183V | **ACh** | 300 | 3 |
| 13 | $(\alpha4)_3(\beta2)_2$ | α4T139V | **Var 1** | 300 | 3 |
| 14 | $(\alpha4)_3(\beta2)_2$ | α4T139V | **Nct 2** | 300 | 3 |
| 15 | $(\alpha4)_3(\beta2)_2$ | α4T139V | **Cyt 3** | 300 | 3 |
| 16 | $(\alpha4)_3(\beta2)_2$ | α4T139V | **ACh** | 300 | 3 |
| 17 | $(\alpha4)_3(\beta2)_2$ | α4T183Vβ2S133V | **Var 1** | 300 | 3 |
| 18 | $(\alpha4)_3(\beta2)_2$ | α4T183Vβ2S133V | **Nct 2** | 300 | 3 |
| 19 | $(\alpha4)_3(\beta2)_2$ | α4T183Vβ2S133V | **Cyt 3** | 300 | 3 |
| 20 | $(\alpha4)_3(\beta2)_2$ | α4T183Vβ2S133V | **ACh** | 300 | 3 |
| 21 | $(\alpha4)_3(\beta2)_2$ | α4T139Vβ2S133V | **Var 1** | 300 | 3 |
| 22 | $(\alpha4)_3(\beta2)_2$ | α4T139Vβ2S133V | **Nct 2** | 300 | 3 |
| 23 | $(\alpha4)_3(\beta2)_2$ | α4T139Vβ2S133V | **Cyt 3** | 300 | 3 |
| 24 | $(\alpha4)_3(\beta2)_2$ | α4T139Vβ2S133V | **ACh** | 300 | 3 |
| 25 | $(\alpha4)_2(\beta2)_3$ | Wild type | **Var 1** | 300 | 3 |
| 26* | $(\alpha4)_2(\beta2)_3$ | Wild type | **Nct 2** | 100 | 5 |
| 27* | $(\alpha4)_2(\beta2)_3$ | Wild type | **Cyt 3** | 100 | 5 |
| 28* | $(\alpha4)_2(\beta2)_3$ | Wild type | **ACh** | 100 | 5 |
| 29 | $(\alpha4)_2(\beta2)_3$ | Wild type | **C2 Var 4** | 300 | 3 |
| 30 | $(\alpha4)_2(\beta2)_3$ | Wild type | **Isovar 5** | 300 | 3 |
| 31 | $(\alpha4)_2(\beta2)_3$ | Wild type | **C2 Var 4** | 300 | 3 |
| 32 | $(\alpha4)_2(\beta2)_3$ | Wild type | **Isovar 5** | 300 | 3 |

#### C. nAChR Ligand Binding Measurements

##### *(i)* Expression of human α4β2, α3β4 and α7 nAChR

*Heterologously expressed α4β2 and α3β4 nAChR.* HEK 293 cells were grown in Dulbecco's modified Eagle medium supplemented with 10% fetal bovine serum (FBS), 1% L-glutamine, 100 units/mL penicillin G and 100 µg/mL streptomycin in a humidified atmosphere containing 10% $CO_2$. cDNAs encoding human α3 and β4 or α4 and β2 were transfected into the HEK 293 cells at 30% confluency.

*Heterologously expressed α7 nAChR.* The SH-SY5Y cells were grown in RPMI medium (Lanza) supplemented with 10% fetal bovine serum (FBS), 1% of penicillin-streptomycin and 1% of L-glutamine. cDNA encoding human α7 was transfected into the SH-SY5Y cells at 30% confluency. The cells were maintained in an environment of 37°C containing 5% $CO_2$. The cell transfections were carried out in 100 mm Petri dishes using 30 mL of JetPEI$^{TM}$ (Polypus, France) (1 mg/mL, pH 7.2) and 10 µg of cDNAs. After 48 h transfection, the cells were collected, washed with PBS by centrifugation and frozen or used for binding analysis.

##### *(ii)* Radioligand binding assays

(±)-[$^3$H]Epibatidine (specific activity of 56-60 Ci/mmol) and [$^{125}$I]α-bungarotoxin (α-Bgtx) (specific activity of 200-213 Ci/mmol) were purchased from Perkin Elmer (Boston MA). Non-radioactive α-Bgtx, nicotine and epibatidine were purchased from Sigma-Aldrich.

*[$^3$H]Epibatidine binding.* Details of the binding experiments to the nicotinic subtypes have been previously reported by Tasso *et al.*[27] Saturation experiments were performed by incubating aliquots of membranes from HEK cells expressing α4β2 or α3β4 nAChR with 0.01-2.5 nM concentrations of (±)-[$^3$H]epibatidine overnight at 4°C. Nonspecific binding was determined in parallel by incubation in the presence of 100 nM unlabelled epibatidine. At the end of the incubation, the samples were filtered on GFC filters soaked in 0.5% polyethyleneimine and washed with 15 mL ice-cold phosphate buffered saline (PBS) and the filters were counted for radioactivity in a β counter. The affinity ($K_d$ in nM) of [$^3$H]epibatidine for the α4β2 and α3β4 nAChR subtypes were 0.075, and 0.194 respectively and were derived from the average value of three independent [$^3$H]epibatidine binding saturation experiments.

*[$^{125}$I]α-Bgtx binding.* Saturation binding experiments were performed using membranes of α7-transfected SHSY5Y incubated overnight with 0.1-1.0 nM concentrations of [$^{125}$I] α–Bgtx at rt. Nonspecific binding was determined in parallel by incubation in the presence of 1 µM unlabelled α–Bgtx. After incubation, the samples were filtered as described above and the bound radioactivity was directly counted in a γ counter. Specific radioligand binding was defined as total binding minus the nonspecific binding determined in the presence of 1 µM unlabelled α–Bgtx. Nonspecific binding was ~20-30% of total binding. The $K_d$ of [$^{125}$I] α–Bgtx for the α7 subtype was 1.2 nM and was derived from the average value of three independent [$^{125}$I]α–Bgtx binding saturation experiments.

***(iii)* Competition binding assays**

The ability of varenicline variant ligands to compete for the agonist binding sites of α4β2, α3β4 or α7 nAChR was determined by inhibition of [$^3$H]epibatidine and [$^{125}$I] α–Bgtx binding. Membranes from cells transfected with the appropriate nAChR subtype were incubated with increasing concentrations of test compound for five minutes, followed by overnight incubation at 4 °C, with [$^3$H]epibatidine: 0.1 nM (for α4β2 nAChR) or 0.25 nM (for α3β4 nAChR), or at rt with [$^{125}$I] α–Bgtx: 2-3 nM (for α7 nAChR); radioligand concentrations approximate to their experimentally determined Kd values (see below). After incubation, the membranes were washed five times with ice-cold PBS. [$^3$H]epibatidine binding was determined by liquid scintillation counting in a β counter, and [$^{125}$I] α–Bgtx binding by means of direct counting in an γ counter.

***(iv)* Statistical analysis**

Data from competition binding assays were evaluated by one-site competitive binding curve-fitting procedures using GraphPad Prism version 6 (GraphPad Software, Inc, CA, USA). Half maximal inhibition concentrations ($IC_{50}$) for varenicline variant ligands were obtained by fitting three independent competition binding experiments, each performed in duplicate for each compound on each subtype. Inhibition constants ($K_i$) were estimated by reference to the $K_d$ of the radioligand, according to the Cheng-Prusoff equation.

In the saturation binding assay, the maximum specific binding ($B_{max}$) and the equilibrium binding constant ($K_d$) values were calculated using one site–specific binding with Hill slope – model.

#### D. nAChR Methods and Functional Studies

##### *(i)* Animals

Adult female *Xenopus laevis* were purchased from the European *Xenopus* Resource Center (Portsmouth, UK). *Xenopus laevis* toads were housed and cared for following the UK Home Office code of practice guidelines for the species. The collecting of oocytes from *Xenopus* toads was carried in a regulated room in the Biomedical Services facility in Oxford University, where the toads were housed.

##### *(ii)* Human α4β2 nAChR expression in *Xenopus* oocytes

Compounds reported were tested for effects on the function of human α4β2 nACh receptors expressed heterologously in *Xenopus* oocytes, which were isolated from adult female *Xenopus laevis* toads as previously described.[7] Human α4β2 receptors were expressed as either $(\alpha4)_3(\beta2)_2$ (low sensitivity for ACh) or $(\alpha4)_2(\beta2)_3$ (high sensitivity for ACh) receptors. Expressions in oocytes was obtained as follows. Human cDNA for α4 or β2 were subcloned into plasmid pCI from Promega and injected into the nucleus of *Xenopus* oocytes as described previously.[28] To express $(\alpha4)_3(\beta2)_2$ nACh receptors, a mixture of 10 α4 : 1 β2 cDNAs was injected into the nucleus of oocytes, whereas for $(\alpha4)_2(\beta2)_3$ receptors the cDNA ratio injected was 1 α4 : 10 β2.

##### *(iii)* Single and double mutations

Mutations were introduced in the α4 or β2 nAChR subunits using the Stratagene QuikChange Site-Directed Mutagenesis Kit (Agilent, UK). The presence of the mutation and the absence of unwanted mutations were confirmed by sequencing the entire cDNA insert (Eurofins, UK). Note that we present the numbering of the residues according to the full length of the following UniProt sequence codes for human α4 and β2 subunits, respectively: P43681 (α4 subunit) and P17787 (β2 subunit). To obtain the position in the mature form, subtract 28 from the number for α4 and 25 for β2 subunit.

##### *(iv)* Electrophysiological recordings

Electrophysiological recordings were performed 2-5 days post-injection, as previously described (Moroni et al., 2006). Current responses were obtained by two-electrode voltageclamp recording at a holding potential of -60 mV using an Oocyte Clamp OC-725C amplifier (Warner Instruments, USA).

Concentration-response curves for agonists assayed were obtained by normalizing agonist-induced responses to the control ACh responses induced by 1 mM, a maximum effective ACh concentration at both α4β2 nAChR stoichiometries. A minimum interval of 5 minutes was allowed between agonist applications to ensure reproducible recordings. The agonist concentration-response relationship was characterized for data from each cell using non-linear regression in GraphPad (Prism 5, GraphPad, USA) by fitting the Hill equation ($Y([compound]) = Y_{max}\ (1 / (1 + (EC_{50}/[compound])\hat{}n^{Hill}))$), where Y is the response to a concentration of compound, $Y_{max}$ is the maximal response, $EC_{50}$ is the concentration producing half-maximal activation, and $n^{Hill}$ is the Hill coefficient. Concentration-Response data were collected for an individual cell, and data were normalized to the response to 1 mM ACh. The fit was rejected if the estimated error in any fit parameter was greater than 60% of the fit value, and all parameter estimates for that fit were discarded. For compounds that elicited less than 10% of the maximal ACh response, the $EC_{50}$ was not determined, and the relative efficacy was established by using the equation: maximal response to test compound/maximal ACh response. Data points represent the mean ± standard error of the mean (SEM) of 8-10 experiments carried out in at least three different batches of oocytes donors.

***(v)* Statistical analysis**

For functional assays, the final data sets were assembled from a minimum of 5 independent recordings (i.e. n = 5) conducted on oocytes obtained from at least 5 different *Xenopus* donors. Data obtained from the same batch of oocytes were considered replicates. The data sets represent full concentration-response relationships obtained from individual oocytes (i.e., incomplete experiments were discarded). The data from each experiment were fitted separately and the estimated $EC_{50}$ values were used to obtain the mean $EC_{50}$ or $IC_{50}$ (95% CI) reported in the manuscript or Supplementary Information. $LogEC_{50}$ values for agonist were analyzed using one-way ANOVA, followed by a post hoc Dunnett's test and/or a posthoc Bonferroni multiple comparison test to determine the level of significance between wild type and mutant receptors. Prior to the ANOVA analysis, the data were tested for normality using the D'Agostino and Pearson normality test in PRISM and were normally distributed. Post hoc tests were run only if F achieved $P < 0.05$ and there was no significant variance in homogeneity.

##### *(vi)* Supporting figures and tables

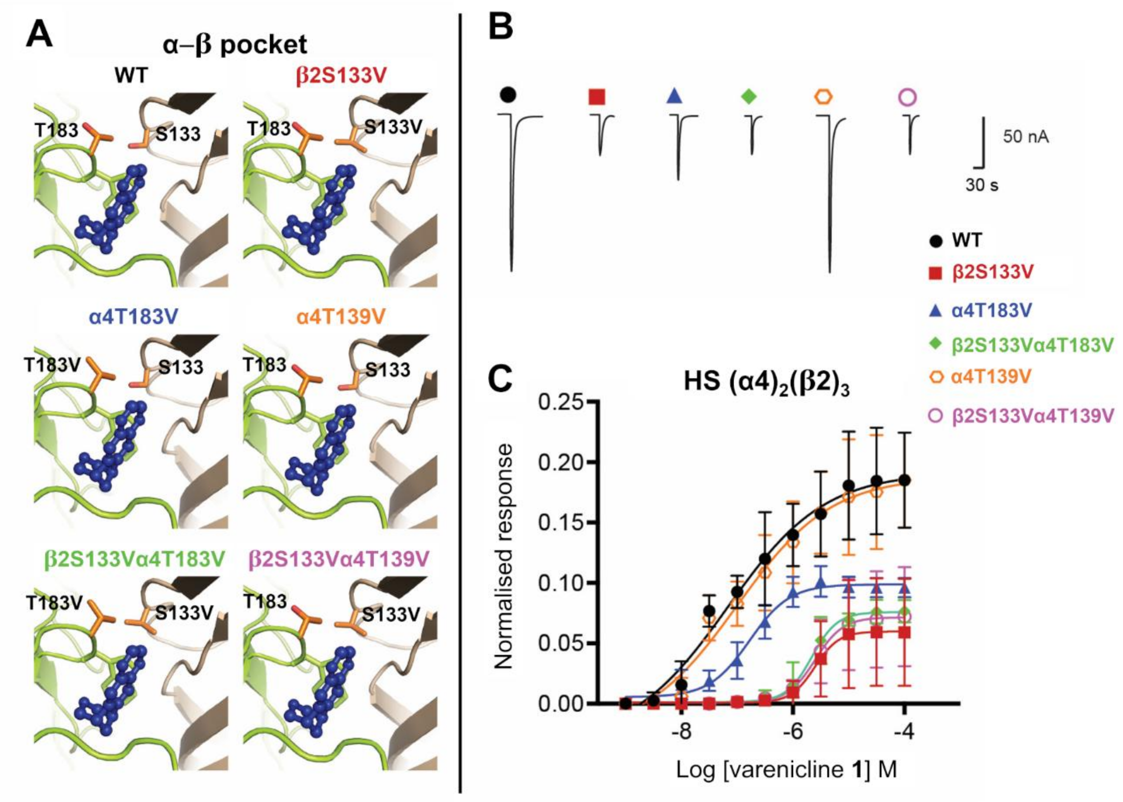


**Figure S39-** Effects of side-chain hydroxyl mutations on varenicline **1** agonism at HS $(\alpha4)_2(\beta2)_3$ nAChR. (**A**) Location of α4T183 and β2S133 and corresponding mutations to valine in the α-β binding pockets of the WT and mutants. The α4 and β2 subunits are colored in yellow and light brown, respectively. Varenicline **1** is highlighted in dark blue. The side-chain of α4T183 and β2S133 are represented with orange sticks, whereas TrpB is shown with yellow sticks. (**B**) Representative current traces elicited by maximal concentration of varenicline **1** (100 μM) applied to *Xenopus* oocytes expressing wild-type (WT) or mutant HS $(\alpha4)_2(\beta2)_3$ nAChR. Full concentration-responses curves are shown in panel **C**. (**C**) Concentration-response curves for varenicline **1** at wild-type (WT) and mutant $(\alpha4)_2(\beta2)_3$ nAChRs. Data points in the concentration-response curves represent the mean ± SEM of 8-10 experiments carried out using 6-8 different *Xenopus* donors. Current responses were measured using two-electrode voltage-clamping from *Xenopus* oocytes heterologously expressing WT or mutant $(\alpha4)_2(\beta2)_3$ nAChRs. Peak current amplitudes for varenicline **1** were normalized to maximal ACh response (1 mM) and then fitted with the Hill equation, as described in the Materials and Methods section above. Estimated parameters $EC_{50}$ and maximal relative efficacy (RE) are shown in Table 1 in the main text.

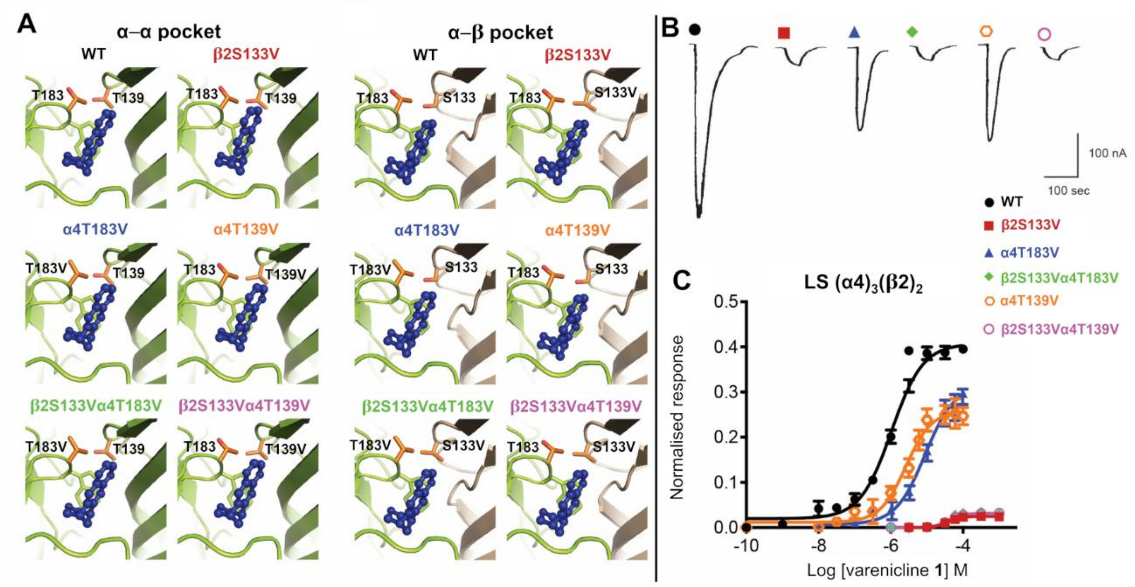


**Figure S40-** Effects of side-chain hydroxyl mutations on varenicline **1** agonism at LS $(\alpha4)_3(\beta2)_2$ nAChR. (**A**) Location of α4T183, α4T139 and β2S133 and corresponding mutations to valine in the α-α and α-β binding pockets of the WT and mutants. The α4 and β2 subunits are colored in yellow and light brown, respectively. Varenicline **1** is highlighted in dark blue. The side-chain of α4T183, α4T139 and β2S133 are represented with orange sticks, whereas TrpB is shown with yellow sticks. (**B**) Representative current traces elicited by maximal concentration of varenicline **1** (100 μM) applied to *Xenopus* oocytes expressing wild-type (WT) or mutant LS $(\alpha4)_3(\beta2)_2$ nAChR. Full concentration-responses curves are shown in panel **C**. (**C**) Concentration-response curves for varenicline **1** at wild-type (WT) and mutant $(\alpha4)_3(\beta2)_2$ nAChRs. Data points in the concentration-response curves represent the mean ± SEM of 8-10 experiments carried out using 6-8 different *Xenopus* donors. Current responses were measured using two-electrode voltage-clamping from *Xenopus* oocytes heterologously expressing WT or mutant $(\alpha4)_3(\beta2)_2$ nAChRs. Peak current amplitudes for varenicline **1** were normalized to maximal ACh response (1 mM) and then fitted with the Hill equation, as described in the Materials and Methods section above. Estimated parameters $EC_{50}$ and maximal relative efficacy (RE) are shown in Table 1 in the main text.

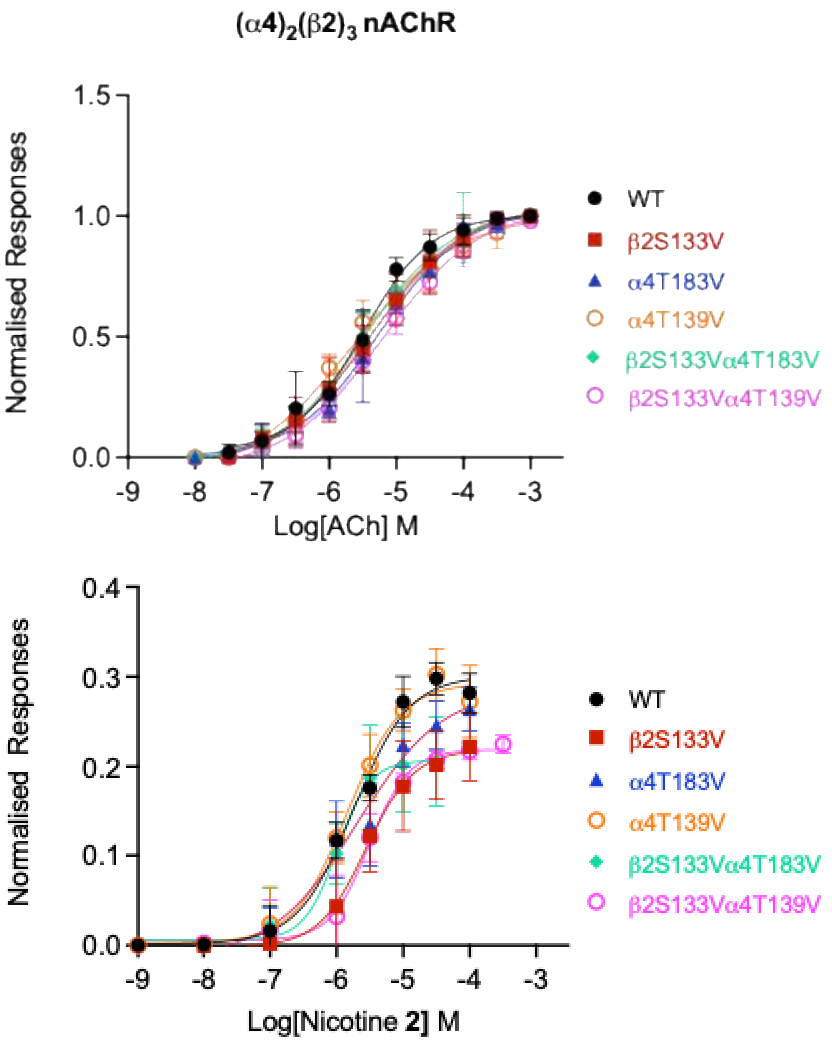


**Figure S41-** Effects of side-chain hydroxyl mutations on ACh or nicotine **2** agonism at HS $(\alpha4)_2(\beta2)_3$ nAChR. Concentration-response curves for ACh and nicotine **2** at wild-type (WT) and mutant $(\alpha4)_2(\beta2)_3$ nAChRs. Data points in the concentration-response curves represent the mean ± SEM of 8-10 experiments carried out using 6-8 different *Xenopus* donors. Current responses were measured using two-electrode voltage-clamping from *Xenopus* oocytes heterologously expressing WT or mutant $(\alpha4)_2(\beta2)_3$ nAChRs. Peak current amplitudes for ACh or nicotine **2** were normalized to maximal ACh response (1 mM) and then fitted with the Hill equation, as described in the Materials and Methods section above. Estimated parameters $EC_{50}$ and maximal relative efficacy (RE) are shown in Table 1 in the main text.

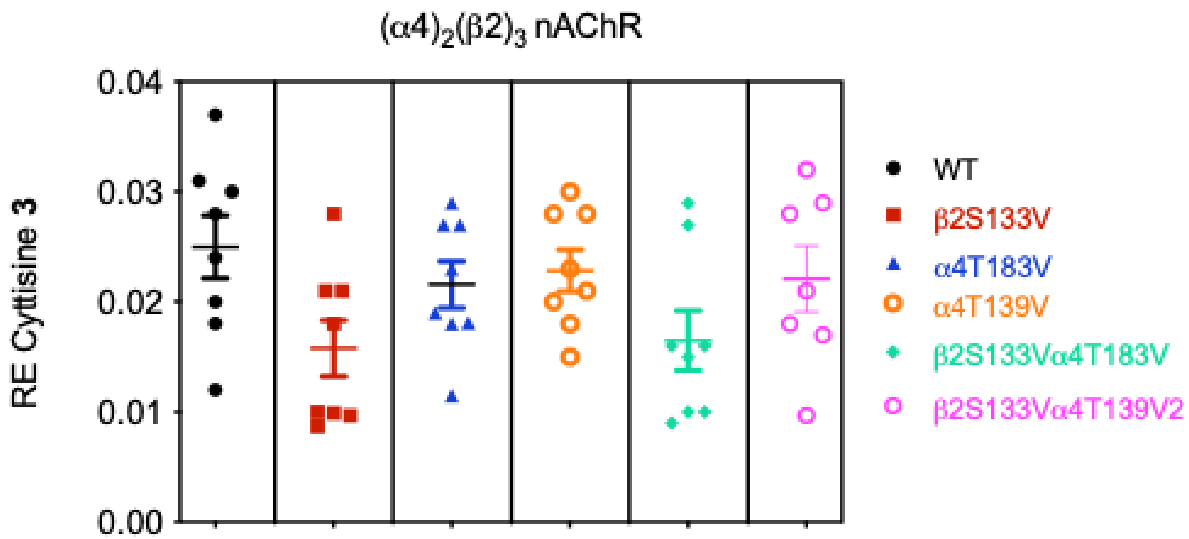


**Figure S42-** Relative efficacy of cytisine **3** at wild-type (WT) or mutant $(\alpha4)_2(\beta2)_3$ nAChRs. Cytisine **3** displays poor agonist efficacy at $(\alpha4)_2(\beta2)_3$ nAChRs, making it difficult to generate concentration-response curves. To assess the functional effects of side-chain hydroxy mutations on cytisine agonism, we estimated the relative efficacy of cytisine using the equation $I_{max}/I_{max}$ACh. Oocytes were challenged with increasing concentrations of cytisine until the responses reached a plateau, which was considered as the maximal current response to the agonist. Data are shown as a box and whisker plot. Current responses from wild-type (WT) or mutant receptors were measured using two-electrode clamping, as described in Materials and Methods section above. Statistical comparisons between WT and mutant receptors were performed using One Way ANOVA followed by a post hoc Dunnett's and/or Bonbeferroni multiple comparison tests. The estimated means ± SEM are shown in Table 1 in the main text.

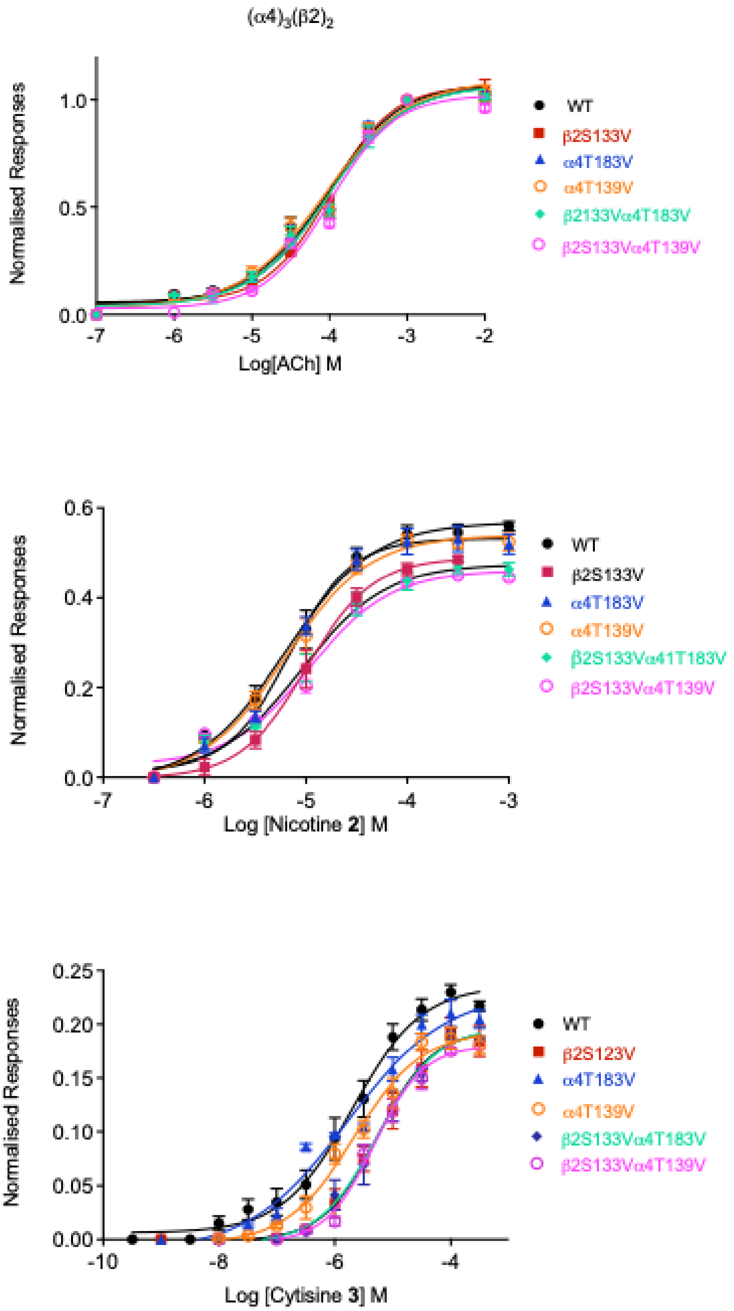


**Figure S43-** Effects of side-chain hydroxyl mutations on agonist sensitivity of $(\alpha4)_3(\beta2)_2$ nAChRs. Concentration-response curves for ACh, nicotine **2** and cytisine **3** were obtained at wild type (WT) or mutant $(\alpha4)_3(\beta2)_2$ nAChRs. Data points in the concentration-response curves represent the mean ± SEM of 8-10 experiments carried out using 6-8 different *Xenopus* donors. Current responses were measured using two-electrode voltage-clamping from *Xenopus* oocytes heterologously expressing WT or mutant $(\alpha4)_3(\beta2)_2$ nAChRs. Peak current amplitudes for all agonists tested ACh were normalized to maximal ACh response (1 mM) prior fitting the data with the Hill equation, as described in the Materials and Methods section above. Estimated parameters $EC_{50}$ and maximal relative efficacy (RE) are shown in Table 1 in the main text.

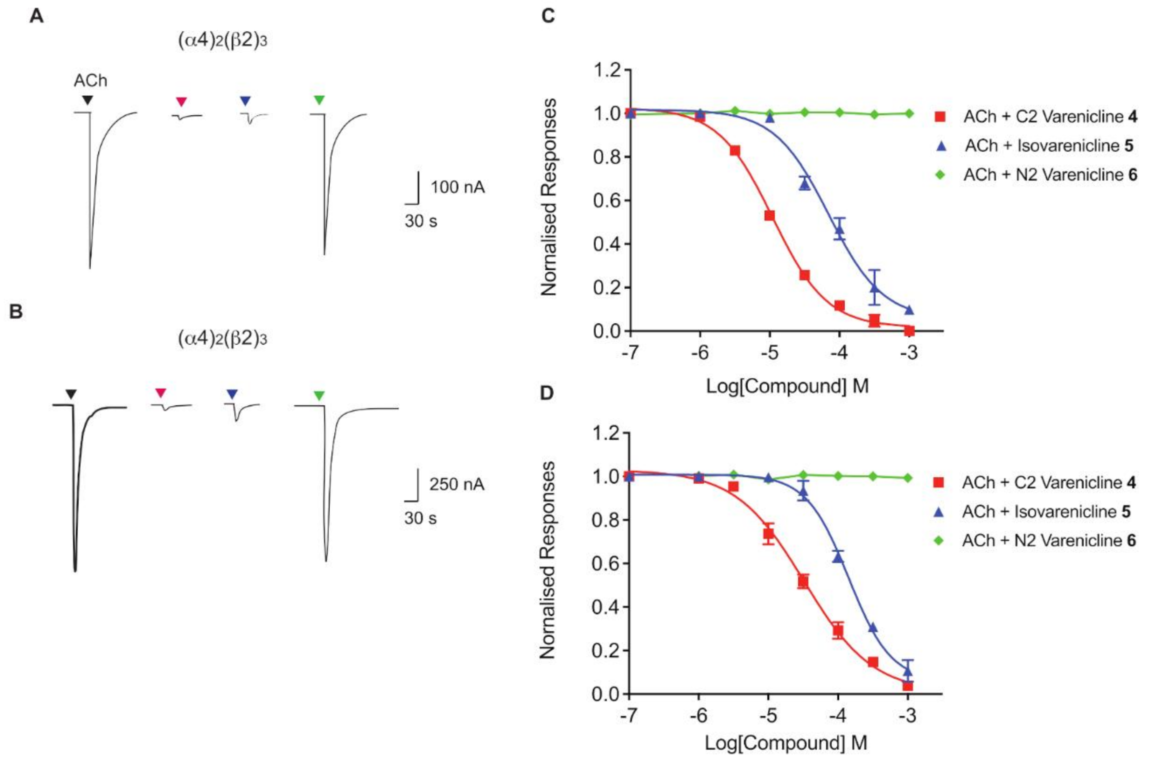


**Figure S44-** Inhibition of α4β2 varenicline **1**, $C_2$ varenicline **4**, isovarenicline **5** and $N_2$ varenicline **6** on α4β2 nAChR. $C_2$ varenicline **4** and isovarenicline **5** behave as partial agonist at α4β2 nAChRs and as such should be able to inhibit the responses to ACh. To obtain concentration-response curves data for the inhibitory effects of the varenicline ligands on the alternate forms of the α4β2 nAChR, the compounds (1 nM to 1 mM range) were co-applied with ACh $EC_{80}$: 30 μM for $(\alpha4)_2(\beta2)_3$ and 300 μM for $(\alpha4)_3(\beta2)_2$ receptors. The peak of the current responses obtained in this manner were then normalized to the peak of the responses elicited by ACh $EC_{80}$ alone. The normalized data were then fit by non-linear regression to the Hill equation, as described in the Materials and Methods section above. As shown in the concentration response curves for $C_2$ varenicline **4** and isovarenicline **5**, these ligands inhibited the responses to ACh in a concentration dependent manner. In contrast, $N_2$ varenicline **6**, which had no agonist effect at α4β2 nAChR, did not inhibit the responses to ACh, indicating that this ligand does not bind the agonist sites present on $(\alpha4)_2(\beta2)_3$ or $(\alpha4)_3(\beta2)_2$ nAChR.

**Table S2-** $IC_{50}$ values for the inhibition of $\alpha4\beta2$ nAChR by varenicline variants **4**-**6**. The $IC_{50}$ values shown were estimated non-linearly from the concentration-response data shown above, as described in the part (*iv*) of this section above. NE, no effect.

| **Ligand** | **$IC_{50}$ at $(\alpha4)_2(\beta2)_3$** | **$IC_{50}$ at $(\alpha4)_3(\beta2)_2$** |
|---|---|---|
| $C_2$ varenicline **4** | 11±1.2 | 33±14 |
| Isovarenicline **5** | 70±12 | 142±21 |
| $N_2$ varenicline **6** | NE | NE |

#### E. p*K*a Determination

##### *(i)* Experimental description of the materials and assays

**Table S3-** Summary of all buffers used in spectrophotometric titrations of compounds reported here. The ionic strength of the aqueous component of each sample was maintained at I = 0.3 M with KCl. Aqueous solutions of HCl and KOH were used to access pH values outside of the range accessible by buffers.

| Entry | Aqueous Solution | Total Concentration / M | %fb range | pH range |
|---|---|---|---|---|
| **1** | Hydrochloric acid (HCl) | 0.01 – 0.3 | - | 0.66 – 2.08 |
| **2** | Formic acid buffer (HCOOH/ HCOOK) | 0.1 | 10 – 40 | 2.77 – 3.55 |
| **3** | Acetic acid buffer ($CH_3COOH$/ $CH_3COOK$) | 0.1 | 10 – 90 | 3.75 – 5.98 |
| **4** | Phosphate buffer ($KH_2PO_4$/$K_2HPO_4$) | 0.1 | 10 – 30 | 5.89 – 6.96 |
| **5** | Triethanolammonium buffer ($N(CH_2CH_2OH)_3$.HCl) | 0.1 | 10 – 90 | 6.96 – 9.06 |
| **6** | Carbonate buffer ($KHCO_3$/$K_2CO_3$) | 0.1 | 10 – 90 | 9.19 – 11.15 |
| **7** | Triethylammonium buffer ($NEt_3$.HCl) | 0.1 | 80 – 90 | 11.40 – 11.92 |
| **8** | Potassium hydroxide (KOH) | 0.1 – 0.3 | - | 12.98 – 13.39 |

Measurements of the pH of the buffer solutions used were performed using a Radiometer Analytical MeterLab® PHM210 Standard pH Meter with a Radiometer Analytical XC161 Combination pH electrode containing 3 M KCl solution saturated with AgCl.

UV-Vis absorbance spectra were obtained using a Varian Cary 100 Bio UV-Vis spectrophotometer with a temperature regulated cuvette holder and attached heating unit. All absorbance data were obtained at 25 °C. Spectra were obtained for each substrate for wavelengths in the range 800 nm to 200 nm at 600 nm $min^{-1}$ (1 nm interval, 0.1 s average time, UV/Vis source change over at 350 nm) with baseline correction to account for absorbance due to the buffer present.

For single wavelength absorbance experiments the mean absorbance at a chosen wavelength was measured over the course of 1 min, with correction of the absorbance due to the buffer present.

Margins of error reported for values of $K_a$ and p$K_a$ constants were taken from the standard error in the respective spectrophotometric titration.

***(ii)* Theoretical background**

Due to the comparatively small amounts of material available (e.g. varenicline **1** and the variants described in the main text), a UV-Vis spectrophotometric titration method was employed for p$K_a$ determination. To access the p$K_a$ of a given compound, the change in absorbance at a chosen wavelength, $\lambda_{obs}$, is measured as the pH is changed. The observed absorbance at a given pH, $A_{obs}$, at $\lambda_{obs}$ is determined by the concentrations of the protonated and deprotonated forms of the analyte.

Absorbance-pH data are fitted to Equation (1):

$$K_a = \frac{10^{-pH}(A_{max} - A_{obs})}{(A_{obs} - A_{min})} \quad (1)$$

Equation **Error! Reference source not found.** is used when $A_{obs}$ decreases with increasing pH at $\lambda_{obs}$.

$$A_{obs} = \frac{A_{max}.10^{-pH} + A_{min}.K_a}{10^{-pH} + K_a} \quad (2)$$

Where $A_{obs}$ increases with increasing pH at $\lambda_{obs}$, the data are instead fitted to Equation (2).

$$A_{obs} = \frac{A_{min}.10^{-pH} + A_{max}.K_a}{10^{-pH} + K_a} \quad (3)$$

Owing to the dependence of p$K_a$ on temperature, ionic strength and solvent, in the present study, the following conditions were maintained for each titration: 25 °C, buffer ionic strength, I.S. = 0.3 M and 10 v/v% acetonitrile co-solvent in aqueous solution.

##### *(iii)* Spectrophotometric method validation: 4-dimethylaminopyridine (DMAP) and nicotine 2

As a control, a spectrophotometric titration of DMAP was performed under the experimental conditions described above (

**Figure** S45).

**Figure S45-** Spectrophotometric titration of DMAP

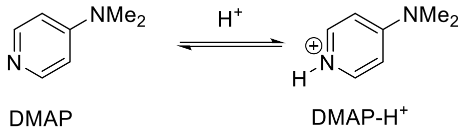


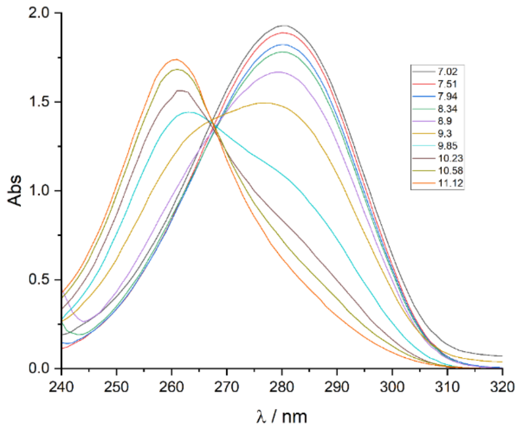


Analytical wavelengths ($\lambda_{obs}$) of 261 nm and 280 nm were chosen on either side of the isosbestic point at 268 nm. At 261 nm, DMAP is more absorbing than DMAP-$H^+$, therefore the data were fit to Equation (2). At 280 nm, the opposite is the case, so the data were fit to Equation **Error! Reference source not found.**. The titrations at both wavelengths are shown in Figure S46.

**A**

**B**

$$A_{obs} = \frac{A_{max} \cdot 10^{-pH} + A_{min} \cdot K_a}{10^{-pH} + K_a}$$

$A_{max} = 1.88 \pm 0.01$

$A_{min} = 0.586 \pm 0.003$

$K_a = (1.92 \pm 0.09) \times 10^{-10}$

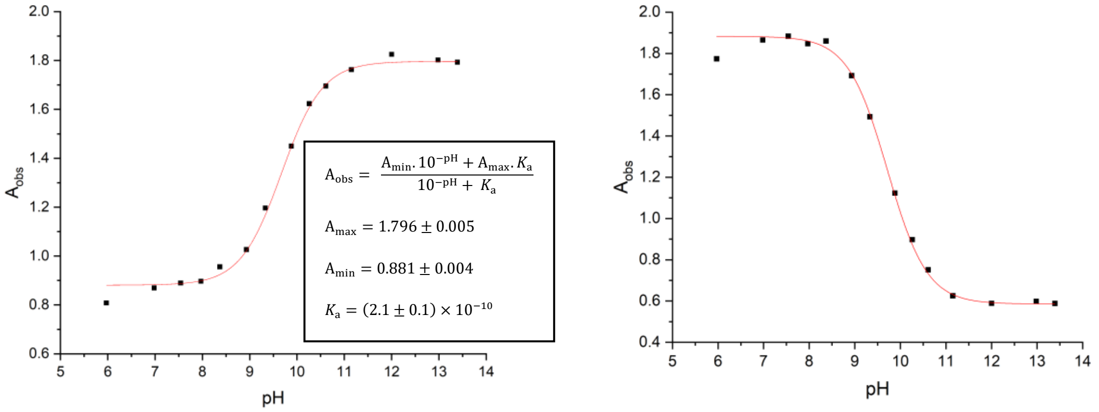


**Figure S46-** Spectrophotometric titration of DMAP at 261 nm (**A**) and 280 nm (**B**).

For the spectrophotometric titration of DMAP at 261 nm, the acid dissociation constant $K_a$ of DMAP-$H^+$ was determined as (2.1 ± 0.1) x $10^{-10}$ M (p$K_a$ = 9.67 ± 0.02) via equation (3). By analysis of data at 280 nm, $K_a$ = (1.92 ± 0.09) x $10^{-10}$ M (p$K_a$ = 9.72 ± 0.02) was obtained via Equation (2). The close similarity in p$K_a$ values determined at both wavelengths and the independent literature value of 9.6, asserts the validity of this method.[29] The small 0.1 unit increase in p$K_a$ of the conjugate acid of DMAP versus the literature value can possibly be attributed to the presence of 10 vol% acetonitrile in aqueous solution, necessary for the solubility of the other substrates in the present study. Increases in p$K_a$ values of neutral and cationic acids are normally observed in the pure weak donor solvent MeCN versus more polar protic media.

The p$K_a$ values of the conjugate acids of nicotine **2** were also determined under these experimental conditions to further validate the UV-Vis spectrophotometric method versus available literature values. The first acid dissociation, p$K_{a1}$ is attributed to the pyridinyl moiety and the second, p$K_{a2}$, to the *N*-methylpyrrolidinyl moiety. The analytical wavelengths for the spectrophotometric titrations were 259 nm and 269 nm, respectively.

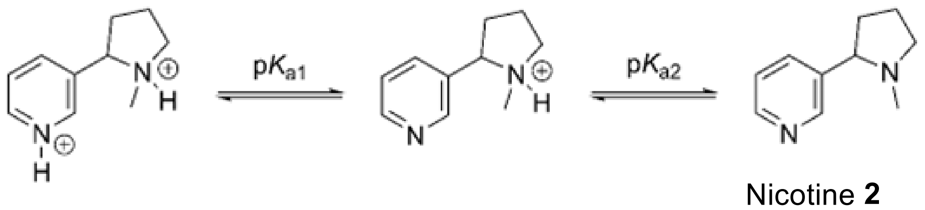


Nicotine **2**

Using the same protocol as described for DMAP, the observed absorbance change due to the deprotonation of the pyridinium moiety is larger than that for the *N*-methylpyrrolidinium component hence a greater concentration of nicotine **2** was necessary to determine $K_{a2}$ (3.75 x $10^{-4}$ M) than for $K_{a1}$ (1.67 x $10^{-4}$ M). Values of p$K_{a1}$ = 3.27 $\pm$ 0.02 and p$K_{a2}$ = 8.19 $\pm$ 0.03 were obtained by UV-Vis spectrophotometric titration under our experimental conditions. These p$K_a$s values are closely similar to reported literature values in water which range from 3.04-3.41 for p$K_{a1}$ and 7.94-8.02 for p$K_{a2}$.[30-33]

##### *(iv)* Spectrophotometric titration of varenicline 1, nicotine 2, cytisine 3 and varenicline variants 4-6

The p$K_a$ values of varenicline **1**, nicotine **2**, cytisine **3**, and varenicline derivatives **4-6** were determined. Spectrophotometric titration curves are shown in Figures S47-S52 and the resultant p$K_a$ values are summarized in Table S4.

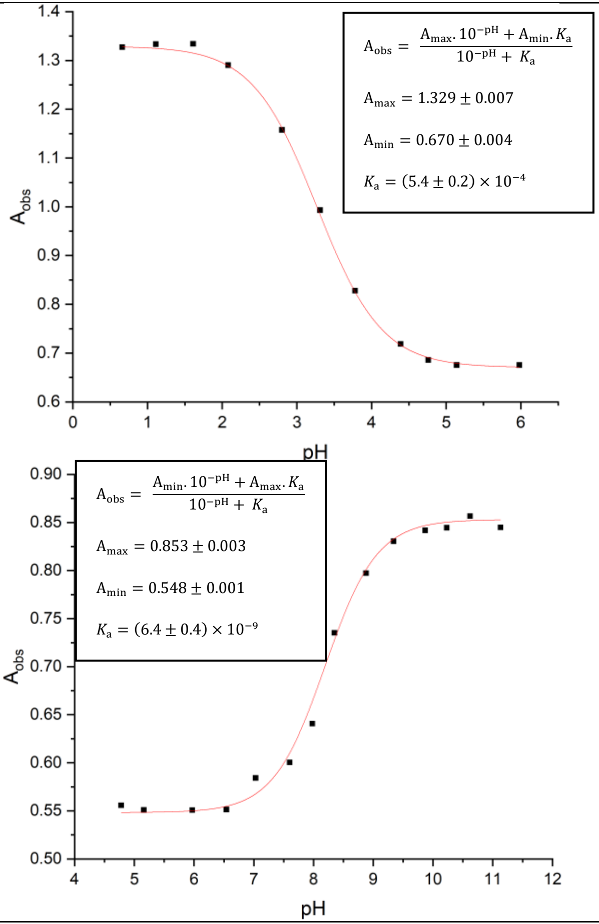


**Figure S47-** Spectrophotometric titrations of nicotine **2**

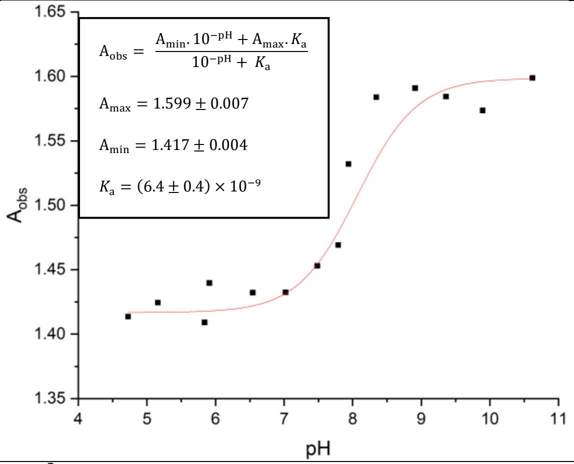


**Figure S48**- Spectrophotometric titration of cytisine **3**

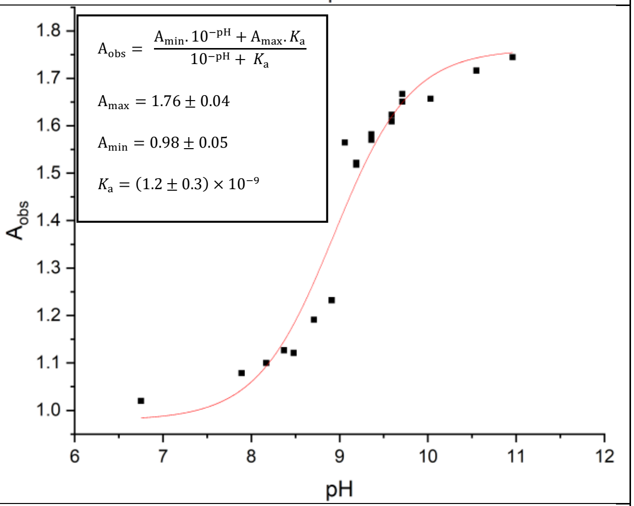


**Figure S49-** Spectrophotometric titration of varenicline tartrate **1**

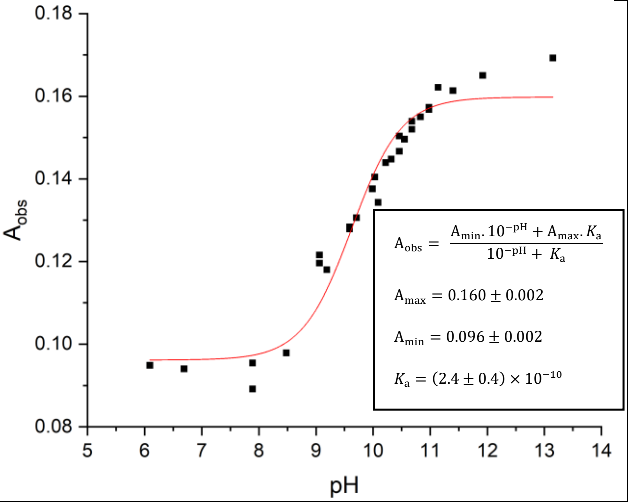


**Figure S50**- Spectrophotometric titration of $C_2$ varenicline hydrochloride **4**

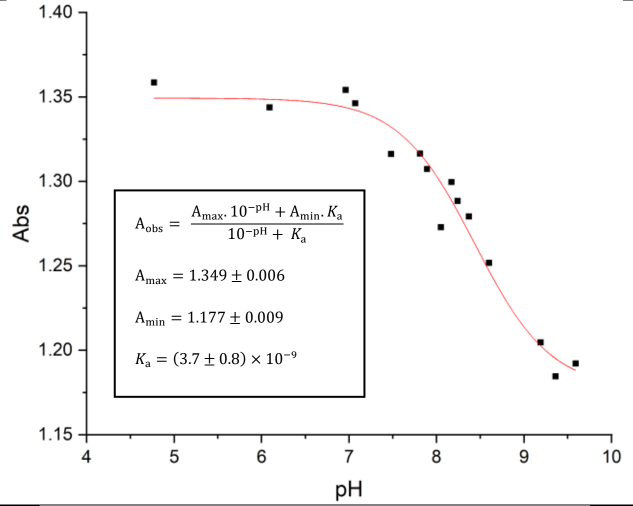


**Figure S51-** Spectrophotometric titration of isovarenicline hydrochloride **5**

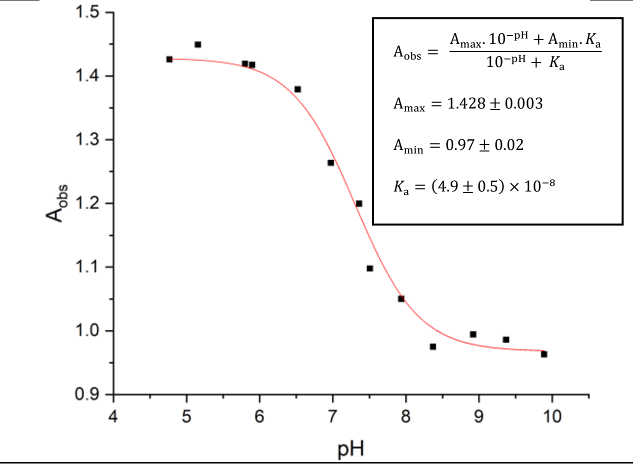


**Figure S52-** Spectrophotometric titration of $N_2$ varenicline trifluoroacetate **6**

**Table S4-** Summary of experimentally-determined p$K_a$ values. [a]Determined at 25 °C, buffer ionic strength, I.S. = 0.3 M and 10 v/v% acetonitrile co-solvent in aqueous solution.

| Compound | p$K_a$ value[a] | Literature value |
|---|---|---|
| 4-Dimethylaminopyridine (DMAP) | 9.67 ± 0.02 | 9.6[29] |
| Nicotine **2** | 3.27 ± 0.02<br>8.19 ± 0.03 | 3.04 – 3.41[30-33]<br>7.94 – 8.02[30-33] |
| 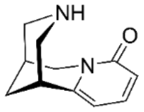<br>Cytisine Error! Reference source not found. | 8.07 ± 0.07 | 7.8[34] |
| 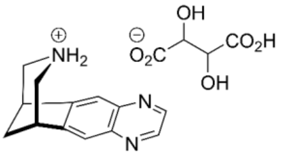<br>Varenicline tartrate **1** | 8.90 ± 0.1 | 9.3[34]<br>9.2 ± 0.1[35] |

| | | |
|---|---|---|
| 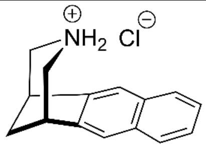<br>C2 Varenicline hydrochloride **4** | 9.63 ± 0.08 | - |
| 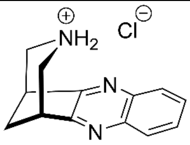<br>Isovarenicline hydrochloride **5** | 8.44 ± 0.09 | - |
| 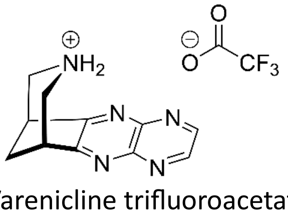<br>N2 Varenicline trifluoroacetate **6** | 7.31 ± 0.05 | - |

The p$K_a$ value for varenicline tartrate **Error! Reference source not found.** (8.9 ± 0.1) is ~0.3 units lower than the literature values (9.3[6] and 9.22 ± 0.13[7]). The literature value of 9.3 is provided without a quoted error or discussion of how the value was reached. The second literature value of 9.22 ± 0.13 was determined by a potentiometric titration in 100% water and at a different ionic strength of 0.15 M which likely accounts for the observed difference of 0.29 units. The p$K_a$ determined for cytisine **Error! Reference source not found.** is 0.3 units greater than the literature value (7.8); again, this literature value is provided without a quoted error or discussion of how the value was reached.[34] The literature p$K_a$ of the conjugate acid of piperidine is reported in the range 11.06-11.18.[30, 36-39] Annelation of the aliphatic piperidinyl ring to the aromatic moiety in varenicline tartrate **1** reduces the p$K_a$ of the piperidinyl moiety by 2.19 units in part owing to ammonium destabilization by the electron-deficient quinoxaline ring. The conjugate acid p$K_a$ of isovarenicline **5** is lower than that of varenicline **Error! Reference source not found.**, due to the closer proximity of the isomeric variant of the quinoxaline to the piperidinyl group, and this decrease reflects an increased electron-withdrawing (destabilizing) effect. $N_2$ Varenicline trifluoroacetate **6** has the lowest p$K_a$ of this series due to the presence of the highly electron deficient 1,4,5,8-tetraazanaphthalene ring system. On the other hand, the naphthalene-based $C_2$ varenicline hydrochloride **4** has the highest p$K_a$ of this series (9.61), although the naphthalene moiety does exert a "reduced basicity" effect relative to piperidine.

The data for varenicline **1** and its derivatives are summarized in Figure S53 and the effect of "local pH" on the distribution of protonated (B⁺H) and free base (B) forms of the key ligands associated with this study is shown in Table S5.

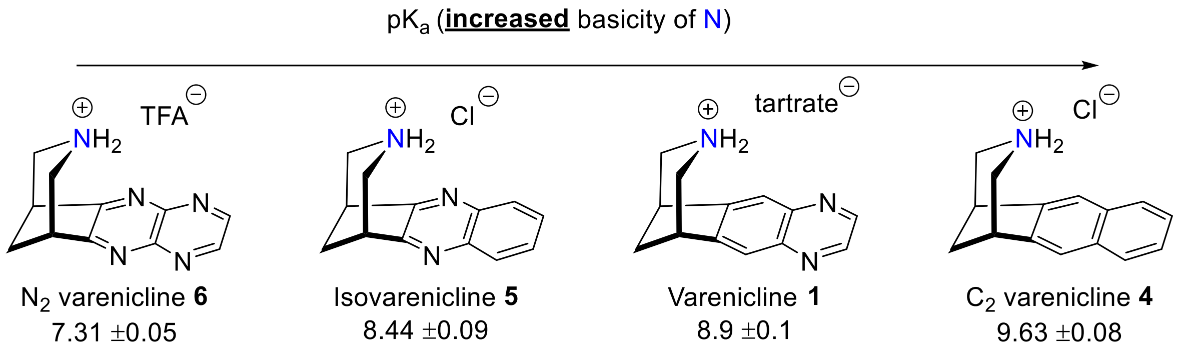


**Figure S53-** p$K_a$ trends of varenicline **1** and derivatives **4-6**.

**Table S5-** pH-Based distribution of protonated ($B^+H$) and free base (B) forms of varenicline **1**, cytisine **3**, and varenicline variants **4-6**.

| | % of **protonated Base⁺H** vs **free Base** | | | | |
|---|---|---|---|---|---|
| Ligand ranked by p*Ka*; **from most to least basic** | **pH 7.5** | **pH 7.8** | **pH 8.0** | **pH 8.3** | **pH 8.8** |
| $C_2$ Var **4**<br>pKa 9.63 | 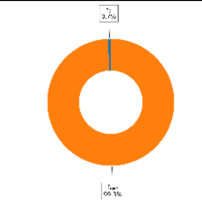 | 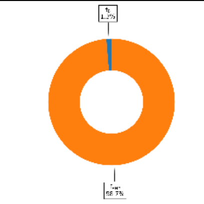 | 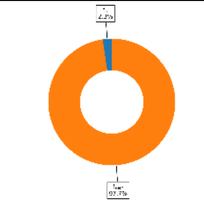 | 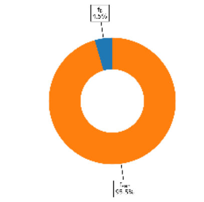 | 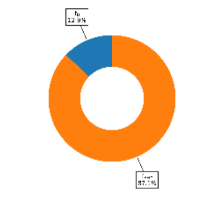 |
| Var **1**<br>pKa 8.90 | 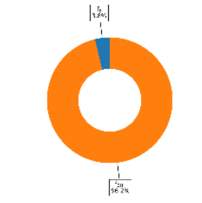 | 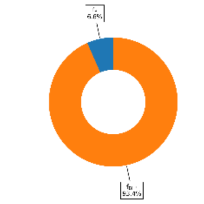 | 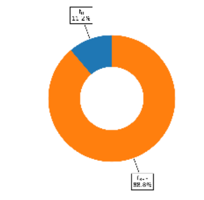 | 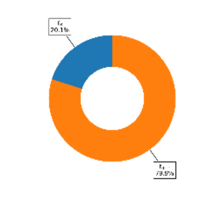 | 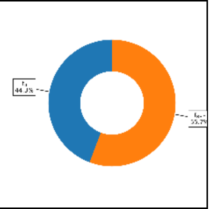 |
| Isovar **5**<br>pKa 8.44 | 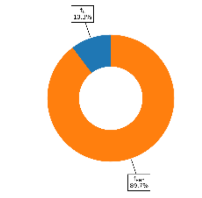 | 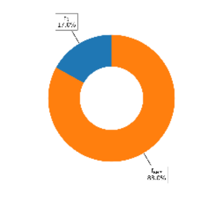 | 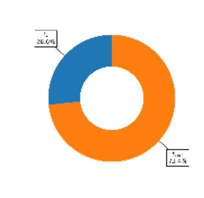 | 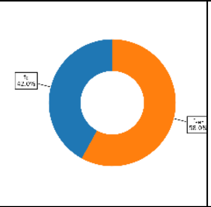 | 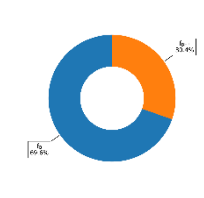 |

| | | | | | |
|---|---|---|---|---|---|
| Cyt **3**<br>pKa 8.20 | 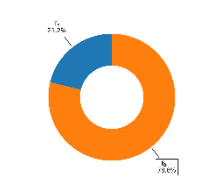 | 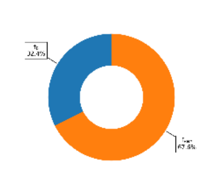 | 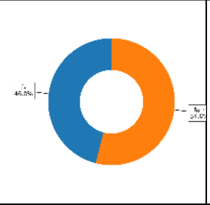 | 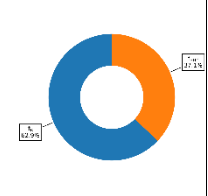 | 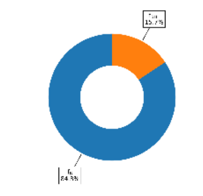 |
| $N_2$ var **6**<br>pKa 7.31 | 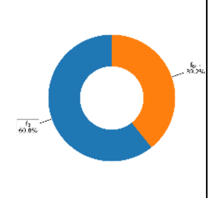 | 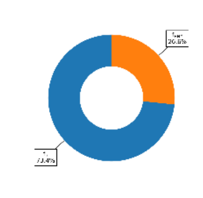 | 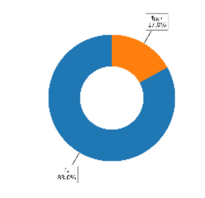 | 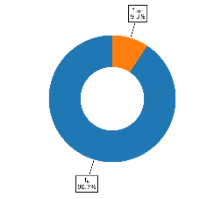 | 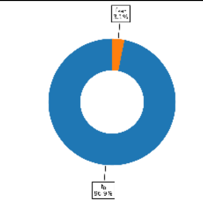 |

| |
|---|
| Ligands showing higher binding affinity |
| Ligands showing lower binding affinity |

#### F. Supporting Information References

(30) Perrin, DD. *Dissociation Constants of Organic Bases in Aqueous Solution*; Butterworths, 1965.
